# High-throughput protein binder discovery by rapid in vivo selection

**DOI:** 10.1101/2025.01.06.631531

**Authors:** Matthew J. Styles, Joshua A. Pixley, Tongyao Wei, Christopher Basile, Shannon S. Lu, Bryan C. Dickinson

**Affiliations:** Department of Chemistry, University of Chicago, 5735 S. Ellis Ave., Chicago, IL 60637; Chan Zuckerberg Biohub, Chicago, IL 60642

## Abstract

Proteins that selectively bind to a target of interest are foundational components of research pipelines^1,2^, diagnostics^3^, and therapeutics^4^. Current immunization-based^5,6^, display-based^7–14^, and computational approaches^15–17,18^ for discovering binders are laborious and time-consuming – taking months or more, suffer from high false positives – necessitating extensive secondary screening, and have a high failure rate, especially for disordered proteins and other challenging target classes. Here we establish Phage-Assisted Non-Continuous Selection of Protein Binders (PANCS-binders), an *in vivo* selection platform that links the life cycle of M13 phage to target protein binding though customized proximity-dependent split RNA polymerase biosensors, allowing for complete and comprehensive high-throughput screening of billion-plus member protein variant libraries with high signal-to-noise. We showcase the utility of PANCS-Binders by screening multiple protein libraries each against a panel of 95 separate therapeutically relevant targets, thereby individually assessing over 10^11^ protein-protein interaction pairs, completed in two days. These selections yielded large, high-quality datasets and hundreds of novel binders, which we showed can be affinity matured or directly used in mammalian cells to inhibit or degrade targets. PANCS-Binders dramatically accelerates and simplifies the binder discovery process, the democratization of which will help unlock new creative potential in proteome-targeting with engineered binder-based biotechnologies.

---

Affinity reagents - molecules that bind to a target protein of interest – are critical as basic research tools for measuring or tracking biomolecules^1^, as probes for studying biological regulation through induced proximity^2^, as core elements of diagnostics^3^, and as therapeutics, such as neutralizing antibodies and antibody-drug conjugates^4^. However, even for very well-studied organisms, including *homo sapiens,* antibody-based binders do not exist for many proteins of interest and, when available, are notorious for heterogenous quality control, function, and specificity^5^.

Binders can be developed by generating antibodies to a target of interest through animal immunization, molecular display methods, or by computational design^6^. Immunization-mediated binder generation typically costs thousands of dollars, takes many months, and is limited to creating antibody-based reagents^6^. *In vitro* selection approaches, such as display-based methods (e.g. phage display, mRNA display), cell sorting methods (e.g. FACS, MACS), and growth-based methods (e.g. bacterial-2-hybrid selections) can be used to mine high diversity libraries of protein variants to identify binders to a target of interest^7–11^. However, these selection-based methods take several months to complete, primarily due to high false positive rates necessitating time-consuming secondary screening^12–14^. Finally, while rapidly improving, computational approaches require significant computing capacities, expertise, and subsequent display-based selections, affinity maturation, and/or screening^15,16^.

Creating a novel binder to a protein generally requires months of highly specialized work, thousands of dollars, and often results in failure. Collectively, the costs and time associated with protein binder generation restrict production to labs with a significant focus and expertise in these techniques, prohibiting exploratory work. In comparison, creating selective binders to DNA or RNA is as simple as designing complementary oligos, which can be synthesized and delivered in a matter of days for ~$10. The programmable nature of nucleic acid binders has led to the rapid explosion of diverse CRISPR technologies and other genetic tools. To address this critical bottleneck, we sought to develop a platform for protein binder discovery that can find novel binders to protein targets of interest in a matter of days with high fidelity, that has the capacity to perform multiplexed screens, and that is easy enough any lab can do it. Such a platform would both accelerate and democratize the binder discovery process.

In this work, we establish <u>P</u>hage-<u>A</u>ssisted <u>N</u>on-<u>C</u>ontinuous <u>S</u>election of protein <u>B</u>inders (PANCS-Binders; **Fig. 1a**), a viral life cycle-based selection platform that can comprehensively screen high diversity (10^10+^) libraries of M13 phage-encoded protein variants and identify binders to panels of dozens or more proteins of interest in a matter of days.

**Fig. 1:**
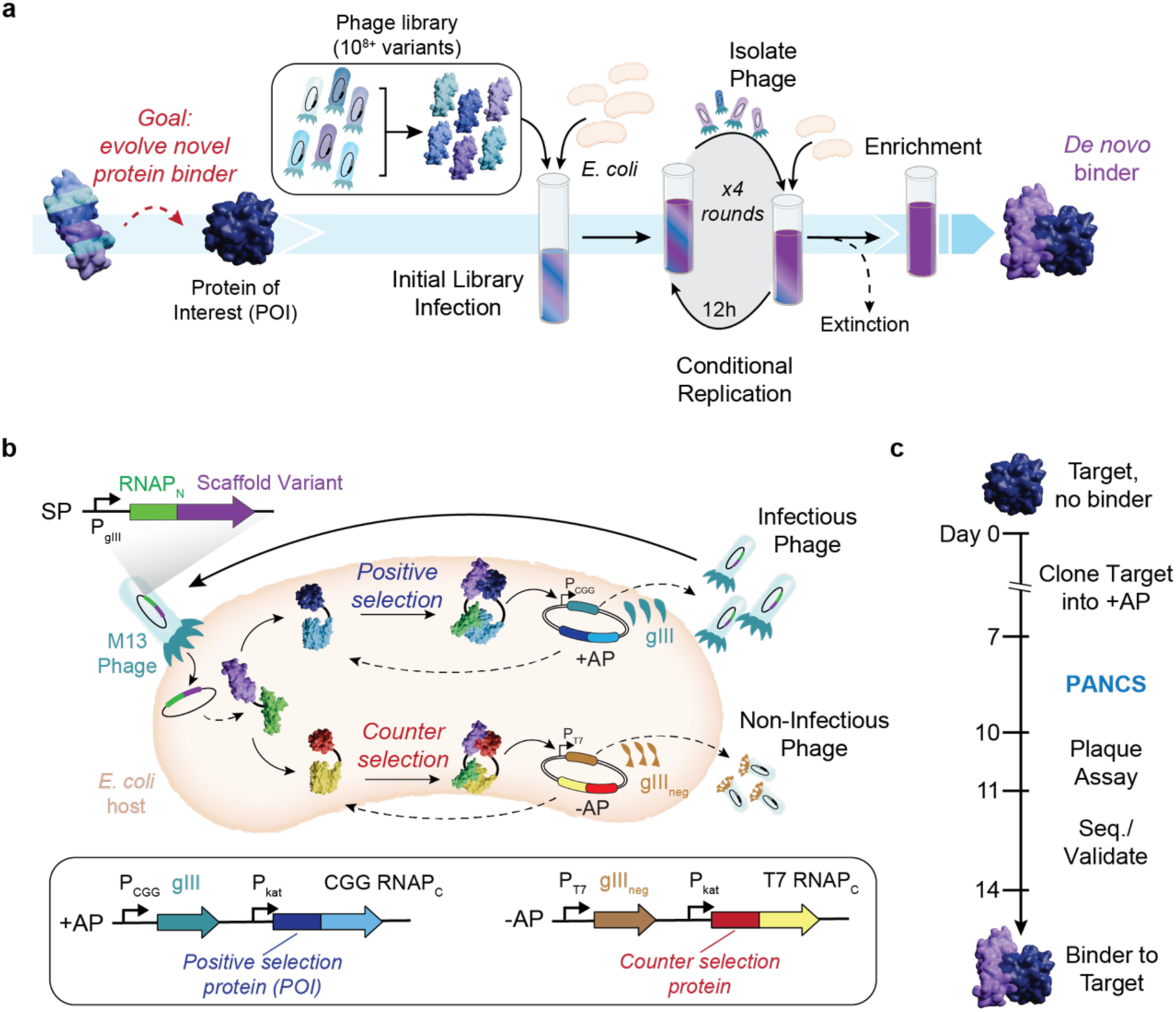
<u>P</u>hage-<u>A</u>ssisted <u>N</u>on-<u>C</u>ontinuous <u>S</u>election of protein <u>B</u>inders (PANCS-Binders) **a,** Schematic of PANCS-Binders process. Serial passaging of *de novo*, high diversity libraries of protein variants encoded in phage for the rapid discovery of binding variants via conditional phage replication on an *E. coli* selection strain. Selections include the extinction of inactive variants and enrichment of the active variants. **b,**Schematic of PANCS-binder molecular biology. A split RNAP biosensor is used for *in vivo* selection of binder variants. *gIII* is removed from the M13 phage and placed on the positive selection plasmid (+AP). Phage encode the N-terminal portion of the split RNAP (RNAP_N_) fused to a protein variant (potential binder). The +AP encodes the target fused to the C-terminal portion of RNAP (RNAP_C_). If the binder variant interacts with the target, then RNAP_N_ and RNAP_C_ recombine and transcribe *gIII*. A simultaneous counterselection is performed using a negative selection plasmid (−AP). The −AP encodes an off-target protein fused to an orthogonal RNAP_C_. If the binder variant interacts with the off-target or RNAP_C_, then the RNAP_N_ and RNAP_C_ recombine and transcribe *gIII*_neg_ – a dominant negative variant of *gIII* that poisons phage amplification by preventing release of phage from the *E. coli* host. **c,** Timeline for 2-week binder discovery using PANCS-Binders: clone the desired target(s) into a +AP(s) and construct the selection strain, 4-6 passages over 2-3 day of PANCS, a plaque assay or qPCR to assess endpoint titer, and then subcloning and sequencing to validate specific enriched variants.

PANCS-Binders uses replication-deficient phage that encode protein variant libraries tagged with one half of a proximity-dependent split RNA polymerase (RNAP_N_) biosensor (**Fig. 1b**)^19^. *E. coli* host cells are engineered to express a target protein of interest tagged with the other half of the split RNA polymerase (RNAP_C_). Protein-protein interaction (PPI) between a phage encoded variant and the target reconstitutes the RNA polymerase (RNAP) and triggers expression of a required phage gene, allowing phage encoding that variant to replicate, in line with the basic principles of PACE^20,21^. After optimization and trial selections, we demonstrated the versatility of PANCS-Binders by performing selections on 95 different protein targets with two *de novo* phage-encoded protein variant libraries, each encoding ~10^8^ unique protein variants, thereby completing 190 independent selections in 2 days. The hit rate of this screen was 55%, resulting in new binders for 52 diverse targets. We scaled up our library size 100-fold (~10^10^), which expanded the hit rate to 72% and dramatically improved the affinity of hits from PANCS – a 40-2000x improvement with affinities as low as 206 pM. Additionally, we showcased how hits can be quickly affinity matured though PACE, resulting in >20x improvement in affinity (to 8.4 nM). Finally, we demonstrated that the binders for two targets, Mdm2 and KRAS, engage their targets in mammalian cells: our Mdm2 binders inhibit the Mdm2-p53 interaction and fusion of our KRAS binder with an LIR motif leads to LC3B mediated degradation of endogenous KRAS^22^. The ease-of-use, speed, and reliability of PANCS-Binders will facilitate a transition of binder generation from an expensive specialty requiring months of work with high failure to a laboratory tool requiring less than 2 weeks (**Fig. 1c**) and available to any researcher.

## Optimizing Binder-PANCS

Recently, we established a split RNAP-based PPI-PACE platform for reprogramming the binding specificity of proteins^23^ (**Fig. 1b**), which we demonstrated could swap the binding specificity of BCL2 and MCL1 using continuous evolution. In general, PACE has been shown to be powerful for altering or tuning existing functions of molecules^24–27^, primarily from initial variants with minimal or closely related function, rather than *de novo* discovery of function. We aimed to adapt the components of our PPI-PACE platform for the use of mining high diversity libraries for *de novo* discovery of binders. To accomplish this, we cloned a phage-encoded, RNAP_N_-tagged 10^8^ unique variant affibody library (**Supplementary Fig. 1**)^28^. We then performed PACE with this library on two targets, the RAS binding domain of RAF (RAF) and IFNG (see **Table S3** for target details). Both evolutions went extinct (**Supplementary Fig. 2**). Prior efforts have established that PACE can enrich active phage from pools of inactive phage (1:1000 active-to-inactive ratio)^21,27^; however, *de novo* libraries are likely to have an active-to-inactive ratio closer to 1:10^7+^. To assess if the PACE evolution process itself led to phage extinction (as opposed to no binders being present in our library), we constructed a mock library selection system that included known, active variants. We performed PACS (Phage-Assisted Continuous Selection^21^), PACE without the mutagenesis plasmid, using KRAS as a protein target (+AP) and a mock library of containing a mixture of active phage encoding RAF that binds KRAS and inactive phage encoding an affitin, evolved to bind SasA, that does not bind KRAS (**Fig. 2a**)^29^. From these mock selections, we found that PPI-PACS could successfully enrich active phage from mock libraries of 1:10^5^ (active:inactive phage; **Supplementary Figure 3**), but failed to do so at any lower ratio (1:10^6–9^). This indicates that continuous selection does not sample every variant in the mock library in the initial infection step. PANCE, non-continuous passaging, has frequently been used as a less stringent version of PACE, and we suspected that part of this lower stringency could be due to a higher percentage of phage that infect cells prior to being washed away in the continuous flow versus in passaging^30,31^. We hypothesized that by extending the incubation time of phage with selection cells, we could more completely sample every variant in our *de novo* library, and therefore succeed in *de novo* selections.

**Fig. 2:**
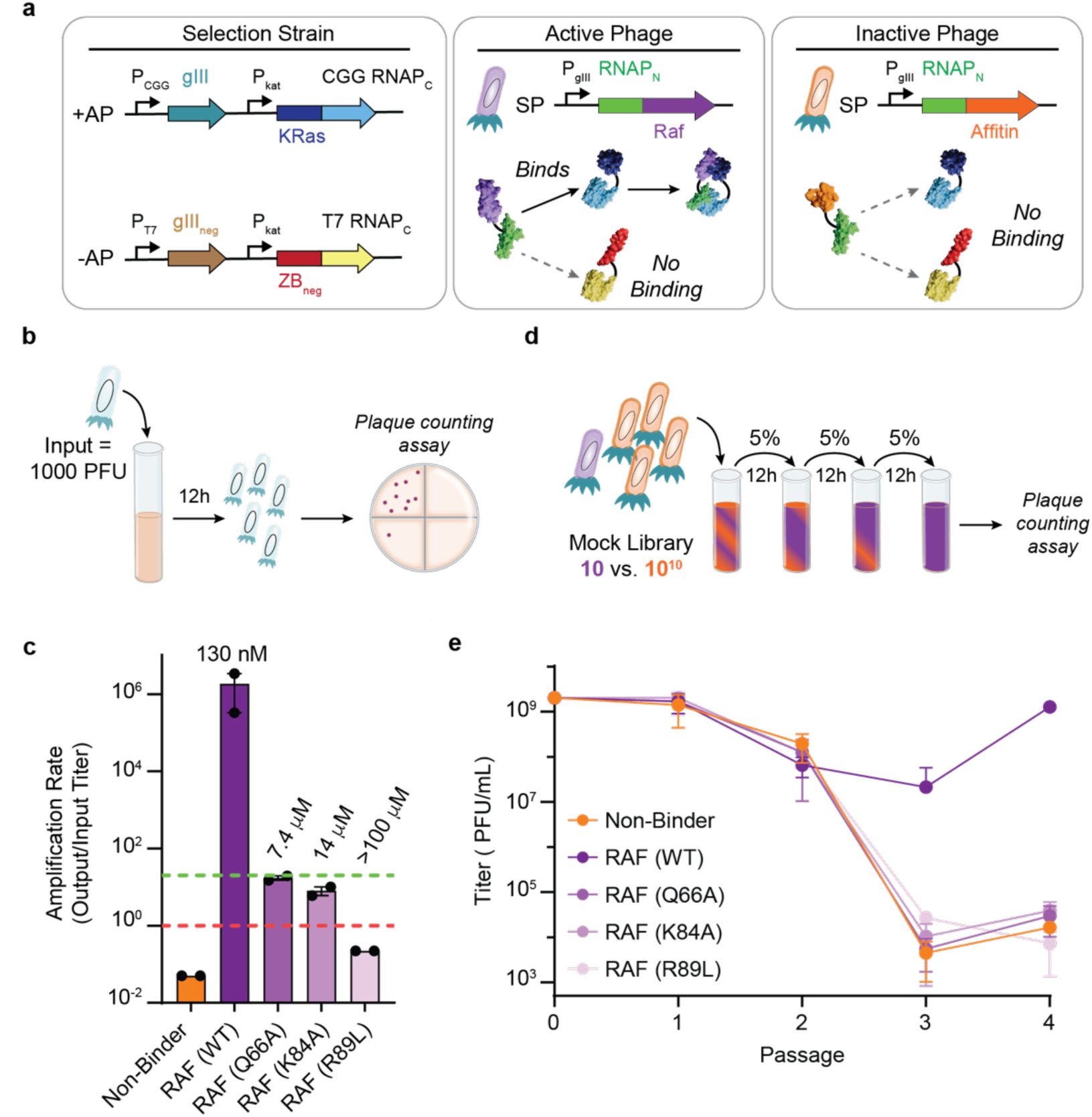
Development of PANCS-Binders for selection from *de novo*-like mock libraries. **a,** The selection system for mock libraries consists of an *E. coli* selection strain with KRAS4b (WT)-RNAP_C,CGG_ as the positive selection target (+AP) and ZB_neg_-RNAP_C,T7_ as the counter selection target (−AP). The mock library consisted of a mixture of two selection plasmids (SP): an active phage with RNAP_N_-RAF(RBD) and an inactive phage with RNAP_N_-Affitin (SasA). **b,** Phage amplification assay: 1000 PFU of each phage are incubated with 1 mL of a selection strain for 12 hours and then the titer is determined: amplification rate is output titer/input titer. **c,** Amplification rate for RAF variants and the non-binding affitin (SasA) phage on the KRAS selection strain. Published affinities (K_d_) are listed above each RAF variant amplification rate^32^. Each amplification rate was obtained in duplicate and error bars indicate SD. The green line indicates an amplification rate of 20 (no enrichment if passaging at 5%) and the red line indicates an amplification rate of 1 (no amplification). **d,** Four passage PANCS starting from a mock library (10 PFU active phage with 10^10^ PFU inactive phage (affitin (SasA)) in 5 mL KRAS4b (WT) selection strain (**Fig. 2a**) with a 12 hr passage outgrowth and 5% transfer of supernatant phage into fresh cells to seed each passage. **e,** Titers at the end of each passage (passage 0 indicates the initial titer to start PANCS). The limit of detection (LOD) in our plaque assays is 5*10^2^ PFU/mL (1 PFU); if a titer had 0 PFU, it was set to 0.2 PFU for calculation purposes (Affitin (SasA) in C). Each phage was passaged in triplicate and error bars indicate SD.

To test this hypothesis, we optimized a non-continuous selection procedure. First, we established 6 hours as a minimum time for incubating phage and cells to obtain nearly complete infection of our phage sample by monitoring the rate at which phage infect our cells (**Supplementary Fig. 4**); 12 hour incubations were chosen for convenience. To determine how quickly active phage would enrich and how quickly inactive phage would de-enrich, we measured the rate of amplification for active (RAF variants with known affinities) and inactive phage (Affitin (SasA)) in a KRAS selection strain (**Fig. 2a, b**). The amplification rates spanned 8 orders of magnitude: 10^6^ for high affinity WT RAF, 10^1–2^ for low affinity RAF mutants, and 10^-1–2^ for non-binders (**Fig. 2c**).

Based on the replication rates, we predicted that serial passaging with 5% of phage transferred between passages would result in selective enrichment of the high affinity WT RAF from 10 phage to >10^9^ phage in just 2 passages and the complete de-enrichment of the inactive phage from 10^9^ to 0 in just 4 passages (**Supplementary Fig. 5**). We tested this prediction by passaging mock libraries of 10 phage of each RAF variant spiked into 10^10^ inactive Affitin (SasA) phage (**Supplementary Fig. 2d**). As expected, over 2 days, the high affinity WT RAF variant enriched (a >10^15^-fold relative enrichment) during the four-passage selection; all weaker binders and inactive phage went extinct (**Fig. 2e**). We performed additional mock PANCS to understand the effects of several variables on this relative enrichment rate: +AP selection stringency (**Supplementary Fig. 6**), −AP selection stringency (**Supplementary Fig. 7**), transfer rate (**Supplementary Fig. 8**), and initial cell-to-phage ratio (**Supplementary Fig. 9**). Finally, we tested mock selections using several published binder-target pairs using our optimized AP strengths, transfer rate, and initial cell-to-phage ratios (**Supplementary Fig. 10**). Collectively, these mock selections indicate that this new system, which we named <u>P</u>hage-<u>A</u>ssisted <u>N</u>on-<u>C</u>ontinuous <u>S</u>election of Protein <u>B</u>inders (PANCS-Binders), can perform *de novo* selections of up to 10^10+^ variant libraries (above the typical 10^9–10^ *E. coli* transformation limit) in 2 days, using simple serial phage out-growths in culture tubes or even 96-well plates. Therefore, we next performed pilot selections with a *de novo* phage-encoded binder library to demonstrate that PANCS-Binders can be used to discover novel binders.

## Pilot *de novo* library PANCS-Binders

We selected six protein targets to attempt to discover novel binders for, each with varying architectures and degrees of structural order: KRAS4b(G12D), RAF (RBD), Mdm2 (1-188), IFNG, Myc DNA binding domain (DBD), and Sos1 disordered domain (**Fig. 3a**, **Supplementary Table 3**). We simply cloned each target into the +AP as a RNAP_C_ fusion and transformed into *E. coli* host cells with the ZB_neg_ −AP used in our final mock selections (**Supplementary Fig. 10**) to prepare the selection materials. We passaged the 10^8^ affibody library, which had gone extinct in PACE-based selections, on each *E. coli* selection strain in culture tubes for 12-hour outgrowths and 5% transfer between passages (**Fig. 3a** and **Supplementary Fig. 1**). After 4 rounds of passaging (48 hours total), we measured the titer in each condition. KRAS(G12D), RAF (RBD), IFNG, and Mdm2 had high titers (>10^8^), indicating successful selections, while Sos1 and Myc (DBD) selections had titers near the limit of detection (<10^5^), indicating failure to enrich binders (**Supplementary Table 4**).

**Fig. 3:**
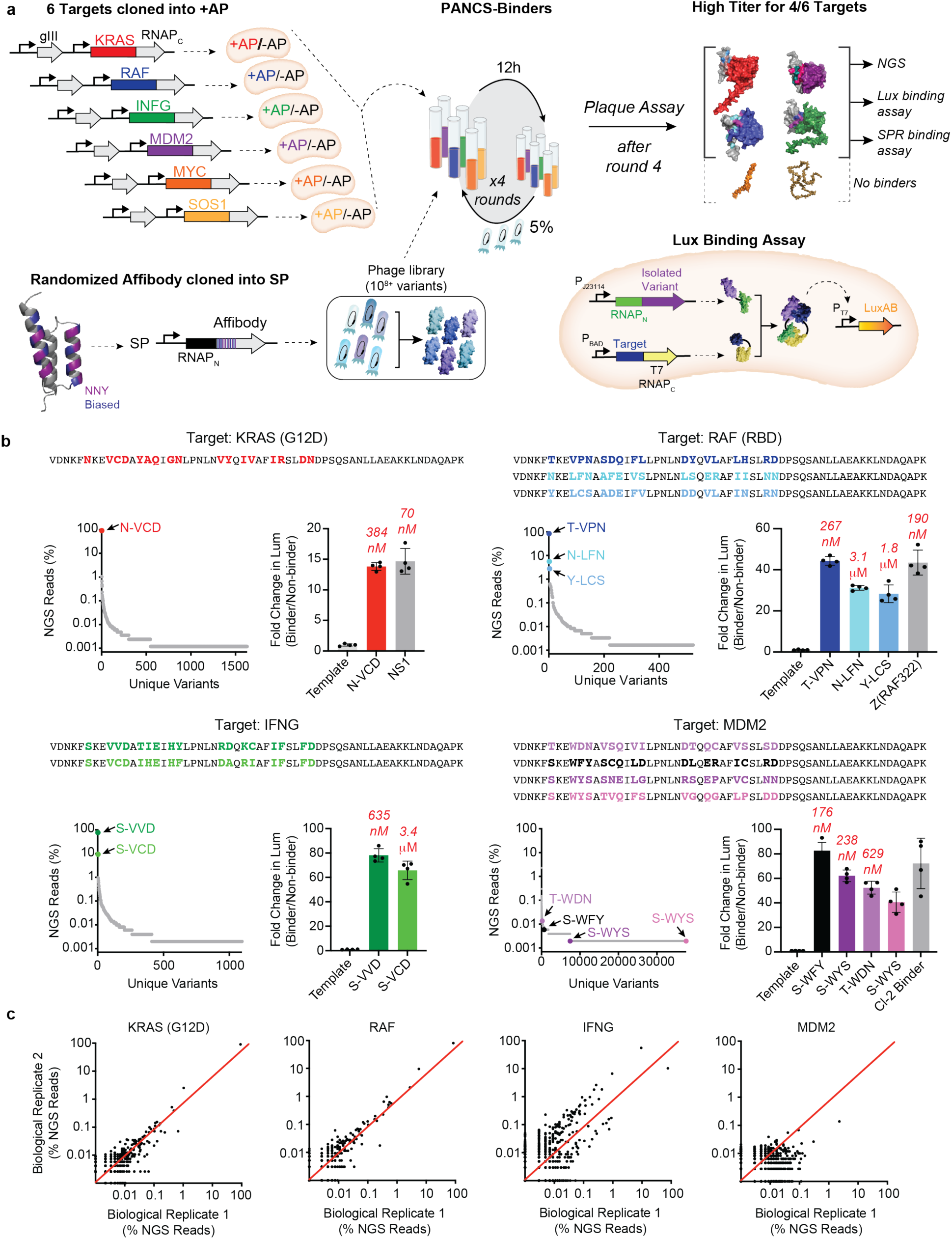
PANCS-Binders to discover novel binders from *de novo* libraries. **a,** Cloning a target +AP panel, transformation of selection strain, parallel 4-passage PANCS of each selection strain with a 10^8^ variant affibody library. Titer assessed after the fourth passage indicated which selections enriched binders (indicated as AlphaFold predictions of the binder-target pair (if titer was high) or target alone (if titer was low), see **Supplementary Table 4** for titers). The affibody sequence from passage 4 phage was then PCR amplified for NGS, subcloned into a luciferase assay system (shown bottom right), and subcloned into pET vectors for protein purification for SPR. Binding validation luciferase assay (bottom right): the target is fused to the RNAP_C_ on an expression plasmid, the binder is fused to the RNAP_N_ on a separate expression plasmid, and a reporter plasmid where LuxAB expression is determined by PPI dependent recombination of the spRNAP. **b,** Top variant amino acid sequences identified from the NGS (top) and the percentage of reads for each unique variant in the NGS of passage 4 (bottom left) for each successful selection. Variants shown in color with the associated sequence were examined further in the luciferase and SPR assays. The second S-WYS variant for Mdm2 isolated for testing in lux (pink) did not have a single read in the NGS but is indicated as having a single read for plotting on a log scale. Fold-change in luciferase signal (bottom right) for select variants from the *de novo* screens over the template affibody used in cloning the library (a non-binder), and when available, previously published binders to these targets (NS1 Monobody (70 nM K_d_^33^), Z(RAF322) Affibody (190 nM K_d_^34^), Cl2-based binder (K_d_ not determined^35^). Each condition has four independent data points, and error bars indicate SD. The *in vitro* binding affinity, K_d_, is reported as text for select variants (as measured by SPR, see **Supplementary Fig. 15**). **c,** Percentage of reads for each unique variant in the original PANCS compared to the percentage of reads for each unique variant in a biological replicate of the PANCS at passage 4 (comparison of parallel replicates in **Supplementary Fig. 17**). If a variant was present in one NGS sample but not in another, it was coded as 0.01%. of reads.

We performed next generation sequencing (NGS) on the four selections with a high titer (**Fig. 3b**), which revealed that the selections on KRAS(G12D), RAF (RBD), and IFNG each converged onto a single sequence (i.e., >80% of reads belonged to that sequence). The most dominant Mdm2 variant comprised only 2.6% of the population; in retrospect, this lack of convergence is unsurprising as Mdm2 binds an FXXXWF/Y motif common to ~3% of library variants, and therefore, many variants were enriched. For KRAS G12D, RAF (RBD), and IFNG selections, we also sequenced the library, passage 2, and passage 3, which revealed that the relative ratio between active variants is set by passage 2 (**Supplementary Fig. 11**), in line with our Mock PANCS results. Finally, we used AlphaFold3 to predict the binding interface for each hit, which, as expected, showed the randomized region of the affibody at the predicted interaction interface (**Fig. 3a** and **Supplementary Fig. 12**).

We subcloned the top variants from each selection (those >1% of reads in KRAS G12D, RAF (RBD), and IFNG selections and 4 random variants from the Mdm2 selection) into an expression plasmid (Lux-N) for measuring binding in a previously established *E. coli* luciferase assay (**Fig. 3a**)^19,21,23^. Reconstitution of the proximity dependent split RNAP, measured by the production of luminescence, was observed to be induced following co-expression of each affibody variant and its respective selection target. This indicated variant:target binding in *E. coli* (**Fig. 3b**). Positive binding controls, previously published binders discovered by ribosome display (Z(RAF322)^34^, phage display (NS1 Monobody)^33^, and rational engineering (Cl2 12.1A)^35^, produced comparable signal to the newly selected affibodies for RAF (RBD), KRAS G12D, and Mdm2, respectively. We confirmed specificity of binding by assaying for binding between one binder from each successful selection with the four targets that gave hits (**Supplementary Fig. 13**), which revealed high specificity of each binder (only S-VVD had off-target binding to Mdm2). We purified each of the top binders (**Supplementary Fig. 14**) and performed surface-plasmon resonance (SPR) binding assays, revealing binding between top variants and their target of interest with *in vitro* affinities between 176 and 635 nM (**Fig. 3b** and **Supplementary Fig. 15**). These results confirmed that the selections successfully enriched binder variants.

To assess reproducibility of the selections, we repeated the entire 6-target selection four additional times in parallel several months later. This yielded highly consistent results in terms of the extinction events and endpoint phage titers (**Supplementary Fig. 16**). We performed NGS on each of the replicate selections and observed high reproducibility (r = 0.72; average of each pairwise Pearson’s Correlation for variants >0.1% of NGS reads; variants enriched >5% were identical) between biological replicates (**Fig. 3c**) and within parallel replicates (**Supplementary Fig. 17**; r = 0.95). These results demonstrate that PANCS-Binders can rapidly - in just 48 hours, comprehensively, and reproducibly screen and isolate binder variants from *de novo* libraries without the need for replicates or additional screening.

## High-Throughput PANCS

We next sought to challenge the PANCS-Binders technology in a multiplexed high-throughput selection by attempting to simultaneously identify binders for a large panel of diverse protein targets in 96-well plate format (**Fig. 4a**). In addition to scaling down the selection volumes to 1 mL for plate compatibility, we made three additional adjustments for this selection: we extended the linker length between the target and RNAP_C_ to ensure that the position and orientation of the binder was not constrained (**Supplementary Fig. 18**), we reduced the selection stringency from 5% to 10% transfer, and we created a second ~10^8^ phage library based on an affitin scaffold to have two different scaffold libraries to compare to one another (**Supplementary Fig. 19**). We simply cloned each protein of interest (a total of 95 targets) into +APs without additional optimization, as well as a negative control no-fusion +AP consisting of a start codon followed by the 60-amino acid GS linker and RNAPc, to establish a 96-well plate of target selection strains. The 95 targets (**Supplementary Table 3**) vary in origin (mammalian, bacterial, or viral), localization (secreted, extracellular domains of membrane proteins, membrane proteins, cytosolic, or nuclear), function, and structure (fully ordered, significant disordered regions, fully disordered). We reasoned that such a diverse target panel should assess the capability for performing PANCS-Binders in a high-throughput manner and provide a realistic estimate of the expected hit rate for 10^8^ libraries. To confirm that the selection +APs were functional, we performed an amplification assay (**Supplementary Fig. 20**) with phage encoding RNAP_N_ WT, which is not proximity dependent (recombines with RNAP_C_ independent of target and binder interacting), with the −AP (demonstrating sufficient counterselection to prevent non-selective replication) and without the −AP (indicating sufficient target expression for binding induced replication; only four targets did not amplify >10-fold). Notably, this panel has many targets that are difficult to purify from *E. coli*, which highlights the superior properties of targets expression plasmids over *in vitro* selection methods.

**Fig. 4:**
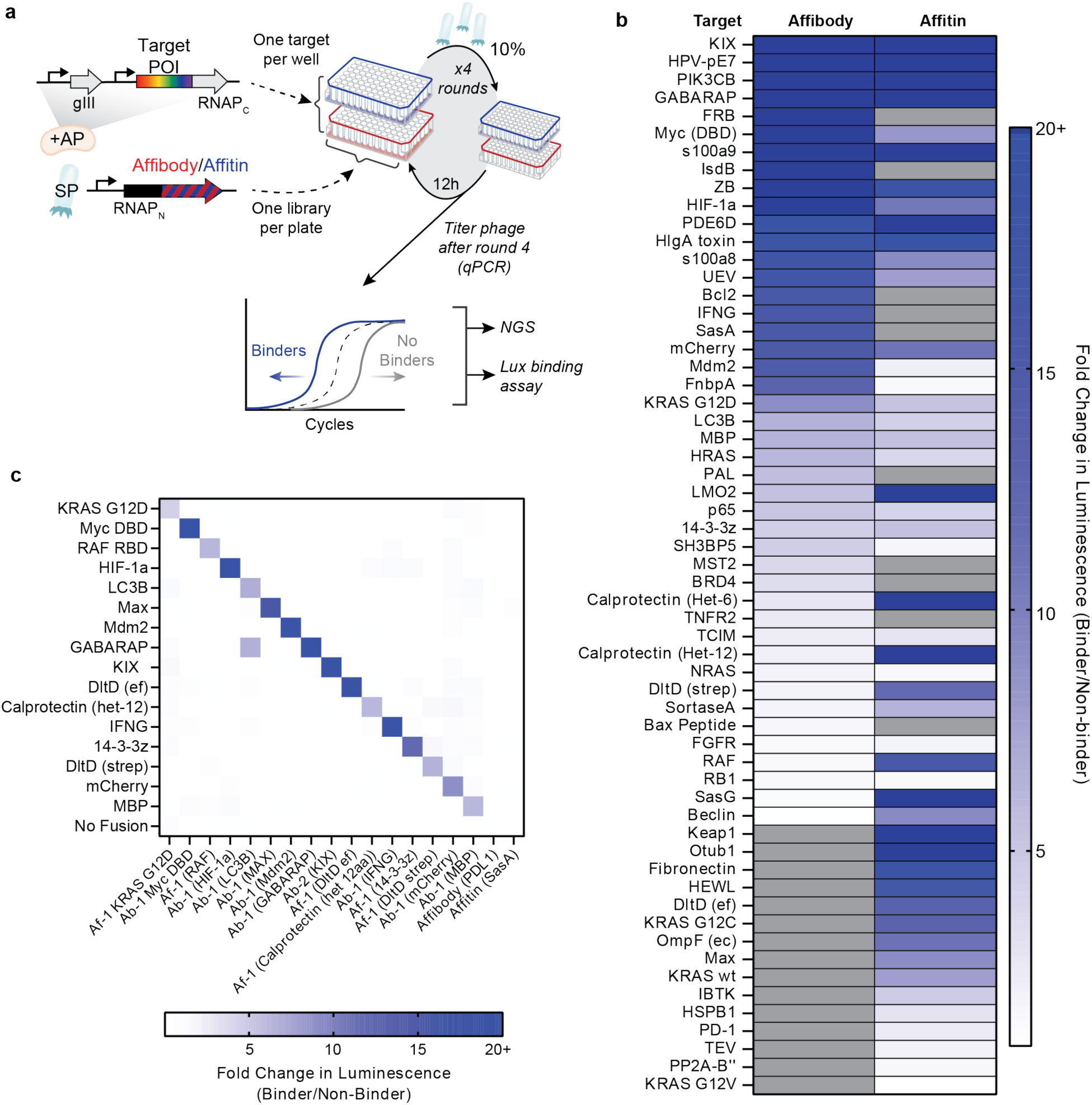
High-throughput PANCS-Binders for 95 targets. **a,** Cloning a 96-panel set of target selection strains, 96-deep well plate-based parallel PANCS-binder selection of two 10^8^ libraries (affibody (**Supplementary Fig. 1**) and affitin (**Supplementary Fig. 18**) using binding assays and qPCR to measure the endpoint titer in a high-throughput manner. **b,** *E. coli* spRNAP complementation luciferase assay heatmap (**Fig. 3a**) on all preliminary hit variants (titer > 10^7^ PFU/mL when top variant is full-length protein (see Note 1 in supporting information and **Supplementary Fig. 23** for individual plots). Fold/change in binding >20 is set equal to 20; selections which were not preliminary hits are indicated in grey. **c,** Fold-change in luciferase signal for selection of 15 variants from the *de novo* screens assessed across 15 targets and additional controls to assess selectivity. Within each target (y-axis) each binder is normalized to the signal for two non-binders (set to 1): an affibody that was evolved to bind PD-L1^36^ and an affitin that was evolved to bind SasA^29^. Only one off-diagonal had a fold-change >2.

After preparing the selection cells, we performed the 4 passage PANCS-Binders over the course of 48 hours and collected endpoint titers using qPCR to identify preliminary hits – titers >10^7^ PFU/mL (**Supplementary Fig. 21** and **Supplementary Table 5**; see **Supplementary Note 1** for a detailed discussion of how we selected this threshold including **Supplementary Figs. 22-26**). For all preliminary hit wells, we collected NGS, performed AlphaFold2 multimer predictions for the top variants (NGS and AlphaFold compiled in **Supplementary Fig. 22**), and subcloned variants from passage 4 phage and collected luciferase binding assay for the top variant(s) cloned (**Supplementary Fig. 23**). Overall, we validated 79 new binders to 52 targets (**Fig. 4b**). These results further demonstrate the high correlation between endpoint titer and binder enrichment. We used 16 of these binders to investigate the specificity of our selected binders (**Fig. 4c**), identifying only one off-target interaction between GABARAP and our LC3B binder which is not surprising given the high sequence similarity between GABARAP and LC3B (both of which bind LIR domains). With this dataset of 288 pairwise binding measurements, we sought to evaluate the ability of AlphaFold3^37^ to identify binding vs non-binding pairs; iPTM values were not well correlated with binding (**Supplementary Fig. 27**).

## Improving affinity and hit rate in PANCS-Binders

While the initial 55% hit rate and 100s of nM K_d_ showcase the viability of PANCS-Binders, we wanted to assess whether we could improve the hit rate and identify higher affinity binders by simply using larger libraries in PANCS-Binders. We cloned 10^10^ variant libraries for both the affibody and affitin scaffolds using parallel transformations (**Fig. 5a**) – a 100-fold improvement of our initial library size (**Supplementary Table 6**). We selected 8 targets that did not give hits in our initial screens, including VHL, TRIM, and PNCA, and 4 targets that previously generated hits, including KRAS G12D and RAF, and performed a large library PANCS-Binders screen (**Fig. 5a**) with 6 rounds of passaging over 72 hours (see Methods for additional details and protocol differences). In addition to getting new hits for the targets that also initially gave hits with smaller libraries, 3 out of the 8 previously failed targets now also yielded hits, as confirmed by the *E. coli* luciferase assay (38%; **Fig. 5a**, **Supplementary Table 7**). We analyzed these hits by NGS (**Supplementary Fig. 28**) and compared the new hits for KRAS G12D both using the *E. coli* luciferase assay and *in vitro* (**Fig. 5b**, binders for other targets shown in **Supplementary Figs. 29** and **31**): the affinity of the best binder obtained from this selection improved from 384 nM to 0.2 nM, representing a ~2000x improvement in affinity. These results confirm that PANCS-Binders is capable of quickly mining 10^10^ variant libraries for novel binders, improving our hit rate (predicted from 55% to 72%), and identifying higher affinity binders, all within a 72-hour selection.

**Fig. 5.**
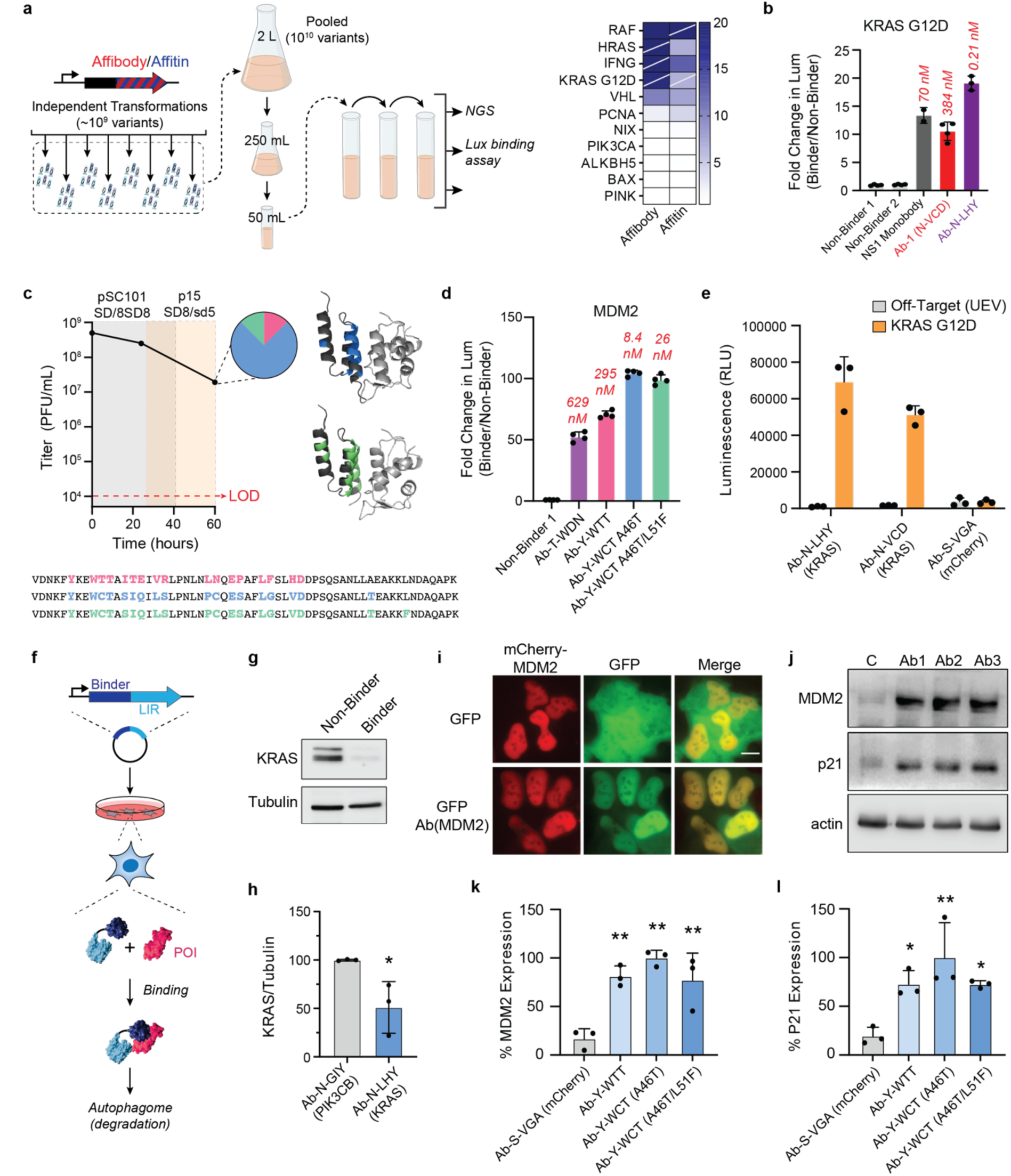
Larger libraries, affinity maturation, and mammalian applications of PANCS-binder hits. **a,** Large, 10^10^, libraries were prepared using parallel transformations (**Supplementary Table 6**) that were pooled to initiate large library PANCS-binder (**Supplementary Table 7**), which required starting at an initial volume of 500 mL and decreasing the volume of each passage slowly as the selection progressed (**Supplementary Fig. 28** for NGS results). *E. coli* spRNAP complementation luciferase assay heatmap shown for successful selections, with slashes indicating selections that also gave hits in the 95 panel selections (**Fig. 4**). **b,** Luciferase assay results for large KRas G12D selection (**Supplementary Fig. 29** for others) with *in vitro* affinity listed above each bar, comparing literature reference binder in grey, original binder from smaller scale selection in red, and new binder in purple (**Supplementary Fig. 30** for SPR). **c,** Affinity maturation of Mdm2 Affibody hit by PACE. Passage 4 from the Mdm2 selection with the 10^8^ affibody library (**Fig. 3b**) was used to seed PACE. PACE was initiated on a +AP selection strain with lower Mdm2 expression than the original PANCS. After 24 hours, there was a 12-hour mixing step with an even lower expression level strain for an additional 36 hours (total of 60 hours). Titers were monitored by activity independent plaque assay, and variants were subcloned at 60 hours (isolated variants shown here). **d,** Variants were tested in the luciferase binding assay alongside T-WDN and the non-binding Affibody (PDL1) as in **Fig. 3n**. Affinities measured by SPR (**Supplementary Fig. 28**) are shown above each luciferase bar graph. **e,** Initial KRAS G12D (**Fig. 3b**) and high affinity KRAS G12D (**Fig. 5b**) binders were tested for binding in mammalian cells using a split-Nano-luciferase assay in HEK293T cells.

One advantage of the PANCS-Binders technology is that the same selection strains and phage can quickly be adapted to PACE by transforming the selection strain with a mutagenesis plasmid to allow mutations to accrue during a directed evolution campaign. To test this, we performed PACE-based directed evolution on the passage 4 phage from our initial Mdm2-affibody selection (**Fig. 3b**) using adaptations of our previous method^23^. First, we identified higher stringency positive APs with reduced propagation of passage 4 phage (as measured by activity dependent plaque assays). Then we used these two strains to perform PACE over the course of 60 hours of evolution, after which the phage populations converged (6/8 subclones sequenced) on an affibody variant Y-WCT with an additional A46T mutation and then an additional L51F mutation (in 1/8 with A46T; **Fig. 5c**). Assessment using the *E. coli* luciferase binding assays showed the evolved variants have improved affinity for their targets (**Fig. 5d**).

We confirmed this *in vitro*: K_d_ = 8.4 nM for A46T, and 26 nM for A46T/L51F (**Fig. 5d** and **Supplementary Fig. 31**) compared with 176 nM for the highest affinity Mdm2 binder identified from the initial PANCS (a >20x improvement). Critically, the mutations arising from the supplemental PACE are not in the initial randomization sites, nor are they predicted to make direct contacts with the target protein (**Fig. 5c**). This is not altogether surprising; the power of unbiased directed evolution via PACE for optimizing existing function though non-intuitive mutations is well-established.

Transfections were performed in triplicate; error bars indicate SD. **f,** Schematic showing LC3B recruiting element (LIR motif) fused to binders for targeted degradation by autophagy.^38^ **g,** Representative WB of endogenous KRAS degradation by Ab N-LHY – LIR (**Supplementary Fig. 33** for full blots and replicates). **h,** Quantification of replicate degradation results as shown in **g** (**Supplementary Fig. 33**). **i,** Mdm2/Mdm2 binder co-localization: mCherry-tagged Mdm2 and GFP-tagged Mdm2 binder co-localize in the nucleus, while a control GFP does not (replicates in **Figure S34**). **j,** Representative WB of Mdm2-p53 PPI inhibition; inhibition of this PPI activates p53 transcription resulting in increased expression of p21 and Mdm2 (**Supplementary Fig. 35** for full blots and replicates). Quantification of replicate WBs for Mdm2-p53 PPI inhibition in U2OS cells: **k,** Mdm2 expression and **l,** p21 expression (**Supplementary Fig. 35**).

For Western blot quantification, statistical analyses were performed using one-way ANOVA with Dunnett’s multiple comparison test binder vs. control. *P < 0.05; **P < 0.01.

## Binders Function in Mammalian Cells

Lastly, we sought to test whether the novel binders could be taken directly from PANCS-Binders into mammalian cells and bind their targets in a functionally relevant manner. We cloned the KRAS G12D and KRAS binders Ab-N-VCD (**Fig. 3b**) and Ab-N-LHY (**Fig. 5b**) into a split nano-luciferase complementation assay^39^ and demonstrated robust binding in HEK293T cells (**Fig. 5e**). Next, we sought to convert Ab-N-LHY into a KRAS degrader by fusing it to an LIR (LC3B interacting region) domain for targeted protein degradation through authophagy^22^ (**Fig. 5f**). We observed robust degradation of endogenous KRAS in U2OS as assessed by WB in a binder0dependent manner (**Fig. 5g** and **Supplementary Fig. 33**). We then demonstrate that the high affinity Mdm2 binder (Ab-Y-WCT A46T) co-localizes with Mdm2 in the nucleus (**Fig. h** and **Supplementary Fig. 34**), confirming binding in mammalian cells. We then overexpressed AB-Y-WCT A46T and several other Mdm2 binders in U2OS cells to see if the binders could inhibit the MDM2-p53 interaction, by monitoring expression levels of Mdm2 and p21, which are both transcriptionally regulated by p53. We observed a strong induction of both targets upon MDM2 binder expression, which indicates robust inhibition of Mdm2 and activation of p53 (**Fig. 5i** and **Supplementary Fig. 35**). These results demonstrate that PANCS-Binders can produce binder variants with functional binding activity in mammalian cells.

## Discussion

PANCS-Binders is a rapid, reproducible, and reliable method for discovering protein binders. The entire process of cloning a target into the +AP/RNAP_C_ expression plasmids, selection, assessment, and secondary validation assays can routinely be performed in 2 weeks without the requirement for highly specialized expertise or equipment. The speed of PANCS-Binders comes from its strong de-enrichment of weak and non-binders and high enrichment of binders, which abrogates the need for secondary screening campaigns common in display-based selection techniques. We propose that two fundamental aspects of such techniques limit the relative enrichment of binding variants over non-binding variants commonly achieved in display methods: threshold selection (bound or not bound) and activity-independent amplification. PANCS-Binders utilizes a split RNAP-based biosensor with a large dynamic range that links the degree of variant function to the degree of phage replication for that variant, both creating a gradient rather than threshold selection and removing the activity-independent amplification step. Notably, neither Alphafold2 Multimer (**Supplementary Fig. 24**) nor AlphaFold3 (**Supplementary Fig. 27** and **32**) could predict binding (false negatives) or non-binding (false positives) from our screen, illustrating the enduring importance and value of real experimentation and limitations still inherent in computational modeling. Furthermore, the high-quality binding data generated through PANCS can be used in improving computational modeling and AI-based design techniques.

The ability to utilize high diversity libraries in phage-assisted selections and evolutions is a powerful tool in the directed evolution arsenal and should expand the range of evolutions possible. PACE and PANCE have been applied to alter or tune the specificity of a variety of protein functions^8,19–21,23–25,27,31,40^; however, because mutations accumulate incrementally, these campaigns require a nearly continuous evolutionary pathway starting from low or non-functional initial variants. In a recent tour de force, the evolution of a protein that binds a small molecule-protein complex, an elaborate pathway was needed to access the 5 mutations needed for minimal function and the 8 mutations eventually reached for high function. This included repeated high mutagenesis drift periods, a 3-position randomized library, steppingstone states (a panel of 16 small molecules), and testing of a wide range of selection stringencies (PACE and PANCE across 26 different stringency selection plasmids). High diversity libraries, like those used here in PANCS-Binders, are prepared using *in vitro* diversification techniques capable of making tens of targeted mutations in the initial variant. PANCS is a powerful approach to jumpstart more difficult evolutionary campaigns by increasing the navigable distance between functional states.

PANCS-Binders can screen multiplexed libraries of 10^10^ phage-encoded variants across dozens of targets in 2-3 days, yielding high affinity, selective binders with sufficient fidelity such that hits can be directly used in secondary assays, such as mammalian cell experiments. In our 96-well based high-throughput PANCS-Binders with 10^8^ variant libraries, we achieved a 55% hit rate across a range of targets with a high correlation between endpoint titer and validated binding (79/92). Repeating a subset of failed selections with a 100-fold larger library produced hits for 38% of those initial failures, showcasing how simply scaling up library size can yield hits for otherwise challenging targets. Additionally, either with large 10^10+^ libraries or through rapid affinity maturation via PACE, high affinity binders (<10 nM) can be obtained quickly.

PANCS-Binders generated hits for disordered protein targets and proteins that are challenging or impossible to purify, showcasing the potential of this screening and selection platform to discover binders for proteins that lack structural data or are incompatible with *in vitro* selection strategies. PANCS-Binders did not require significant optimization of selection conditions, as is commonly the case for 2-hybrid selections; while target solubility does impact the target-RNAP_C_ expression level, the differences were small enough to abrogate the need for target dependent tuning of expression. In other words, with PANCS-Binders, one can “plug-and-play” targets into the system.

## Methods

### Cloning and Bacterial Strain Handling

All plasmids and phage (**Supplementary Table 1)** were cloned by Gibson Assembly (GA) of PCR fragments generated using Q5 DNA polymerase (NEB). All primers (**Supplementary Table 2**) were ordered from IDT. For plasmids, GA mixtures were transformed into chemically competent DH10β *E. coli* and after a 1 h outgrowth in 2xYT media, were plated on antibiotic selective agar plates to isolate individual clones. For phage, GA mixtures were transformed into chemically competent S1030-1059 *E. coli*, and after a 2 h outgrowth, a plaque assay was performed to isolate individual phage clones. All plasmids and phage were confirmed by Sanger sequencing. All plasmid maps with annotations of key features are available in **Supplementary Table 1**. For constructing selection (+AP/−AP) and *E. coli* luciferase (2-22/N-lux/C-lux) strains, S1030 *E. coli* was made chemically competent and then single or double transformations were used (and then repeated as needed until all plasmids were incorporated). *E. coli* strains was grown on agar plates static at 37 °C or in solution at 37 °C with 200 rpm shaking with Luria Broth (LB) supplemented with the appropriate antibiotic unless otherwise indicated. Antibiotics were used at standard concentrations: kanamycin (40 ug/mL), chloramphenicol (33 ug/mL), and carbenicillin (100 ug/mL).

### Plaque Assays

Activity independent plaque assays can be used to determine the phage titer via plaque counting. Activity dependent plaque assays can be used to check for robust phage replication on a given strain. For activity independent plaque assays, an S1030-1059 *E. coli* culture (1059 plasmid encodes *gIII* expressed from the phage shock promoter to produce *gIII* after phage infection), is grown to stationary phase in LB with carbenicillin, subcultured 1/10 in fresh LB with antibiotic to an OD600 of 0.4-0.6, and then used as the selection strain in the plaque assay. Similarly, for activity dependent strains, S1030 with a +AP (and −AP) were grown similarly for use in the plaque assay. For the plaque assay, an initial dilution of the stock can be added based on the expected titer, but generally, 2 µL of a phage stock (or diluted stock) is added to 100 µL of subculture, mixed, and then serially diluted (2 µL into 100 µL) to create 4 dilutions. 750 µL of 50 °C top agar (7 g/L agar, 25 g/L LB) was added to each dilution and then transferred in its entirety to one quadrant of a bottom agar plate (15 g/L agar, 15 g/L LB). After 10-16 h of incubation at 37 °C, plaques become visible and were counted in the quadrant with 10-200 plaque forming units (PFU).

### Phage Amplification Rates

To determine phage amplification rate, the titer of a phage stock is determined using an activity independent plaque assay. Based on this titer, a diluted stock is made that should be 500 PFU/μL (the titer of this diluted stock is confirmed using an activity independent plaque assay). The activity dependent strain (+AP/−AP) is grown to stationary phase in LB with carbenicillin and kanamycin, subcultured 1/10 in fresh LB with antibiotic to an OD600 of 0.4-0.6. 2 µL, 1000 PFU, are added to 1 mL of this subculture and then incubated at 37 °C with shaking for 12 h. The cells are then pelleted and the cell-free supernatant collected for use in an activity independent plaque assay to determine the titer at the end of the amplification assay. The endpoint titer is divided by the starting titer (1000 PFU/mL) to determine the amplification rate.

### PACS/PACE

#### General procedures for continuous flow experiments

PACS^21^ and PACE^19^ were performed as previously described. All tubing, chemostat bottles, and lagoon flasks were bleached thoroughly, rinsed with DI water, and then autoclaved to ensure sterility. 10 L carboys of Davis Rich media were prepared as described previously^19^. Inlet lines consist of short needles unable to reach the culture and outlet lines consist of long needles able to reach the culture. Each chemostat had an inlet line for fresh media, an inlet line with a sterile filter for airflow, an outlet line for waste, and an outlet line for each lagoon. Each lagoon had an inlet line from the chemostat, an inlet line with a sterile filter for airflow, and an outlet line for waste. For PACE lagoons, each lagoon also has an inlet line for arabinose. Each chemostat and lagoon has a magnetic stir bar. Colonies of the selection strain were used to inoculate a 5 mL culture in the relevant media (see below) and grown to stationary phase. This culture was then used to inoculate a 200 mL chemostat (250 mL bottle). This culture was stirred in a 37 °C cabinet until an OD600 of ~0.5 and then fresh Davis Rich media was flowed into the chemostat at ~1 vol/h. The chemostat was monitored for 4 h to ensure that the flow rate maintained a stable OD600 of ~0.5. Phage was then added to each lagoon and then culture was flowed into the lagoon to a volume of 20-25 mL and let incubate for 1 h prior to beginning flow of 1 vol/h. Samples from the lagoons were collected from the waste lines at various timepoints.

#### PACE with libraries and with PANCS output

PACE was performed as describe previously^19^ in line with standard PACE protocols^40^. For PACE with RAF and IFNG with the affibody library (Figure S2), two selection strength strains were used for 36 h each with a 12 h mixing step (60 h total). +APs (ori, gIII/RNAP_C_ RBS strength): p15 SD8/SD8 to pSC101 SD8/SD8 for RAF and from pSC101 SD8/SD8 to p15 SD8/sd5 for IFNG. In addition to the +AP, each strain had a ZB_neg_ −AP (20-1) and MP6 (see Table S1). The initial selection strain supported activity dependent plaques of the affibody binders isolated from PANCS of the affibody library, confirming that binders capable of propagating on the initial selection strain exist in the library. One lagoon was used for each chemostat, initially seeded with 10^10^ PFU of library phage and arabinose began flowing during the 1 h of incubation of phage with culture in the lagoon prior to beginning flow at 1 vol/h. Samples were collected prior to beginning the mixed strain phase (24 h) and after completion of the second strain (60 h) and titers were assessed by activity independent plaque assay. For the PACE with Mdm2 (Figure 5A) starting from the final passage of PANCS with the Affibody library (Figure 3A), the same protocol was followed with the selection strengths of the +AP being pSC101 SD8/SD8 to p15 SD8/sd5 (both strains produced small activity dependent plaques using the passage 4 phage and large activity dependent plaques at the 60 h timepoint of PACE).

#### PACS with mock libraries

PACS was performed as describe previously^21^. The selection strain (S1030/31-69/20-6) was prepared by double transformation into chemically competent *E. coli*. Mock libraries were composed of 10^10^ PFU inactive phage (Affitin (SasA)) and varying amounts (0 (negative control), 10, 10^2^, 10^3^, 10^4^, or 10^5^ PFU) of active phage (RAF WT). In addition, a lagoon was seeded with only 10^3^ active phage as a positive control. Each of these was done in duplicate lagoons/chemostats. Samples were taken at 12, 24, and 48 h and the titer was determined by activity independent plaque assay (Figure S3).

### PANCS

#### General protocol for PANCS

For each passage, a selection strain (+AP/−AP) is grown to stationary phase in LB with carbenicillin and kanamycin, subcultured 1/10 in fresh LB with carbenicillin and kanamycin to an OD600 of 0.4-0.6 prior to adding phage. For passage 1, stock phage are added to the subculture and incubated at 37 °C with shaking (200 rpm) for 12 h, then centrifuged to pellet the cells and collect the cell-free supernatant (referred to as passage 1 phage). For subsequent passages, some fraction of the cell-free supernatant from the prior passage is added to the subculture and incubated at 37 °C with shaking (200 rpm) for 12 h, then centrifuged to pellet the cells and collect the cell-free supernatant (referred to as passage # phage). Titers of each passage or just the final passage were then determined using activity independent plaque assays, or for the 96-target panel, using qPCR (see below).

#### Details for specific PANCS

For PANCS development, a variety of culture volumes, transfer rates, and number of passages were used. For the Mock PANCS (**Fig. 2e**) and the 6-target panel PANCS (**Fig. 3a, Supplementary Table 4**), we performed 4-passage PANCS with 5 mL cultures, initially seeded with 10^10^ PFU for passage 1, and seeded with 250 µL of prior passage for passages 2-4 (5% transfer). For the 96-target panel PANCS (**Fig. 4a, Supplementary Table 5**), we performed 4-passage PANCS with 1 mL cultures (2 mL deep 96-well plates), initially seeded with 2*10^9^ PFU for passage 1, and seeded with 100 µL of prior passage for passages 2-4 (10% transfer). For additional passaging of this PANCS, we did 2% transfer for two passages (**Supplementary Fig. 26**). For the 10^10^ library PANCS (**Fig. 5a**), 6 passage PANCS was performed using either a 2% (RAF, IFNG, KRAS G12D, and HRAS) or 5% (PCNA, ALKBH5, PINK, PIK3CA, TRIM21, VHL, NIX, and BAX) transfer rate between passages and passage 1 was seeded with 5*10^10^ PFU; however, unlike previous PANCS, the volume of each passage was changed as well: 500 mL for passage 1, 125 mL for passage 2, 25 mL for passage 3, and 5 mL for passage 4-6.

### Split-RNAP *E. coli* Luciferase Assays

We followed a slightly modified version of our previously reported assay^19^. For each target, a two-plasmid strain (S1030/2-22/C-lux) was made chemically competent and each binder and non-binder N-Lux plasmid was transformed to make the three-plasmid luciferase strain. Colonies were picked for each binder and non-binder for each strain to inoculate 1 mL of LB with kanamycin, chloramphenicol, and carbenicillin and grown at 37 °C with shaking (200 rpm) for 12-16 h. Strains were then subcultured 7.5 µL into 143 µL of LB with kanamycin, chloramphenicol, carbenicillin, and L-arabinose (2 mg/mL final concentration) in white side, clear bottom 96-well assay plates (Corning 3610) and incubated at 37 °C with shaking (200 rpm) for 3.5 h prior to reading the OD600 and luminescence signal on a BioTek Synergy Neo2 plate reader. Luminescence signal is first divided by the OD600 to normalize luminescence to cell growth. Then the luminescence/OD600 is normalized for all strains with the same C-Lux plasmid (target-RNAP_C_ expression plasmid) are divided by the non-binder signal (non-binders set equal to 1). Due to differences in expression levels of each target and differences in how binding impacts expression level of a target, we do not believe direct comparisons in fold change over non-binder can be made across different targets, and therefore, we plot all binders for an RNAP_C_ expression plasmid separately from other RNAP_C_ expression plasmids.

### NGS of PANCS Hits

We used the Amplicon EZ service provided by Genewiz (Azenta) for Illumina sequencing of each of our hits (<u>GENEWIZ from Azenta | Amplicon-EZ</u>) which provides 50,000+ paired end reads per sample. We used primers to install Illumina partial adaptors (red/purple) and barcodes (blue) to PCR products extending from the linker to after the stop codon of our scaffold in phage (primed with green regions) – see **Supplementary Table 2**. PCR was performed with Q5 DNAP polymerase (NEB) directly from phage (1 µL) in a 25 µL PCR reaction; the initial denaturation step was 10 min at 98 °C to release the ssDNA from the phage particle, a 63 °C Ta, and a 40 second extension time were used with 30 cycles. PCR products are confirmed by gel (5 µL), and the remaining 20 µL of PCR product is pooled with other barcoded PCR products and column purified (Zymo DCC5). Qubit dsDNA High Sensitivity kit is used to determine an accurate concentration of the sample prior to dilution and submission for sequencing. For the NGS data in **Fig. 3, Supplementary Figs. 1, 11** and **17**, we used 7 barcodes/sequencing sample yielding >15,000 reads per condition (library or PANCS passage). For the NGS data in **Supplementary Figs. 22, 25**, and **28**, we used 24 barcodes/sequencing sample yielding >1000 reads for nearly all PANCS samples (reads listed in tables for each library-target pairing in each figure). BB Merge was used to merge each paired end reads (https://jgi.doe.gov/data-and-tools/software-tools/bbtools/bb-tools-user-guide/bbmerge-guide/) and then MatLab was used to separate reads by barcode and to translate reads using modified scripts as described previously (MatLab scripts provided as a supplement)^41^.

### Alpha fold predictions

As a preliminary estimate of how our binders interact with their target, we used AlphaFold2 multimer collab (<u>AlphaFold2.ipynb - Colab</u> (google.com))^42,43^ for predicting the interaction between binder and target for the top 4 variants above 1% of reads (**Supplementary Fig. 22**). AlphaFold3 was released after this analysis, and we subsequently used AlphaFold3 (https://golgi.sandbox.google.com/)^37^ to predict binding interactions for each top variants from our 6-target panel PANCS (**Supplementary Fig. 12**), our 10^10^ library PANCS (**Supplementary Fig. 28**), and for all of the binder-target pairs examined in **Fig. 4c** (**Supplementary Fig. 27**). We implore readers to utilize these predictions only for hypothesis generation rather than as data indicative of an actual interaction.

### qPCR to estimate phage titers

qPCR was tested across several primers that prime to M13 phage genes for linear response of a phage serial dilution. Primers VC-525 and VC-526 were chosen (**Supplementary Table 2**). Power Up SYBR mix was used with the following PCR protocol: 10 minutes at 95 °C (to denature phage particle and release ssDNA); 40 cycles of 20 seconds at 95 °C, 20 seconds at 60 °C, and 20 seconds at 72 °C; then 10 seconds at 95 °C and 60 seconds at 65 °C. qPCR was run on a QuantStudio6Pro. Each run includes a standard curve for which the titer is assessed using activity independent plaque assay.

### Library Construction

#### General Protocol

Libraries were designed based on previously published randomizations^28,44^. Randomization was installed into a template phage using primers with degenerate codons (IDT; see **Supplementary Table 2**). We optimized each step of this protocol to maximize the number of clones obtained. PCR conditions were optimized for each library to produce robust PCR product at 25 cycles and then tested for production at lower cycles to reduce amplification bias (18 or fewer cycles were used for each library reported here). PCR was then scaled up to produce 20-100 μg of PCR product. PCR products were concentrated using the Wizard Kit (Promega) and then digested using DpnI and NheI-HF (NEB) using a multidose cycle: for ~20-50 μg of PCR product in 300-400 μL, and then digested with DpnI and NheI, purified using a Zymo Gel Extraction kit, and then ligated with T4 DNA Ligase (NEB). Ligated products were then electroporated into 1059 *E. coli* cells. The cells were then recovered in 50 mL of 37 °C SOC media and incubated for 2 h at 37 °C with shaking – samples were collected throughout this time for determining the titer by plaque assay. At 2 h, the cells were pelleted and the cell-free supernatant was collected. 1030-1059 (activity independent replication strain) was grown overnight and then subcultured 1:10 to and OD600 of 0.6 at 37 °C with shaking (200 rpm). The phage (cell-free supernatant) was then amplified by adding the phage to this subculture for 8-10 h. At the conclusion of this outgrowth, cells were pelleted and the cell-free supernatant was sterile filtered to create the final library stock (titer determined by activity independent plaque assay).

### Affibody Library

The affibody library (**Supplementary Fig. 1**) was cloned from the Affibody (PDL1) phage (**Supplementary Table 1**) using MS-783 and MS-618 (**Supplementary Table 2**) by Q5 DNAP (NEB) with a T_a_ of 68 °C. For generating the 10^8^ size library, the *E. coli* strain used for the electroporation was SS320 (a highly electrocompetent strain that is capable of phage replication) rather than 10β. The 10^8^ library size was generated with a single transformation of 3 μg ligation product. For the 10^10^ library size, eight transformations of 10 μg ligation product were performed (**Supplementary Table 6**).

### Affitin Library

The affitin library (**Supplementary Fig. 19**) was cloned from either Affitin (SasA), 10^10^ library, or a version of the Affitin (SasA) phage with three stop codons inserted into a region randomized by the primers (**Supplementary Table 1**), 10^8^ library, using MS-624 and MS-799 primers (**Supplementary Table 2**) by Q5 DNAP with GC enhancer with a T_a_ of 68 °C. Both the 10^8^ and 10^10^ libraries were generated following the general protocol with a single 4 μg and eight 8 μg transformations, respectively, (**Supplementary Table 6**).

### Protein Purification

#### General Protocol for Target Proteins

Each target protein (KRAS G12D (1-169), RAF, IFNG, and Mdm2 was cloned into a pET28 vector with a C-terminal 6xHis tag and transformed into BL21 *E. coli* (**Supplementary Table 1**). Cells were grown to an OD600 of 0.8 (37 °C with shaking), chilled on ice, induced with 1 mM IPTG, and then incubated with shaking at 16 °C overnight. Cells were pelleted and resuspended in a lysis buffer (25 mM Tris (pH 7.8), 10% glycerol, 200 mM NaCl). Prior to lysing by sonication, cells were treated with PMSF. The soluble fraction of the lysate was incubated with Ni^2+^ resin, washed with lysis buffer containing 50 mM imidazole, then eluted in lysis buffer containing 250 mM imidazole, and finally buffer exchanged into lysis buffer and concentrated.

#### General Protocol for Binder Proteins

Each binder variant was cloned into a pET30 vector with an N-terminal 3xFLAG and GST tag and transformed into BL21 *E. coli* (**Supplementary Table 1**). Cells were grown to an OD600 of 0.8 (37 °C with shaking), chilled on ice, induced with 1 mM IPTG, and then incubated with shaking at 16 °C overnight. Cells were pelleted and resuspended in a lysis buffer (25 mM Tris (pH 7.8), 10% glycerol, 100 mM NaCl). The soluble fraction of the lysate was incubated with GST resin, washed with lysis buffer, then eluted in lysis buffer containing 10 mM L-glutathione, and finally buffer exchanged into lysis buffer and concentrated. Purified binders shown in **Supplementary Fig. 14**.

### Surface Plasmon Resonance

Surface Plasmon Resonance was performed on a Biacore 8000 using a NTA chip for immobilizing the His-tagged target proteins. Target concentrations were optimized to elicit a response of ~50-100 RU (180 s of 5 uL/s) and then a range of binder concentrations were tested to identify concentrations that produced robust binding (90 s of 30 uL/s). All SPR conducted at 10 °C to maintain slow dissociation of the His-tagged immobilized protein. All dose-responses were fit to a kinetic model for 1:1 binding using the Biacore evaluation software – all fits passed the quality checks in this software (**Supplementary Table 8**).

### Split Nano-Luciferase Assay

62.5 ng of the N-terminus of Nano-Luciferase-binder fusion plasmid and 62.5 ng of the KRas(G12D)-C-terminus of Nano-Luciferase fusion plasmid were co-transfected into HEK293T cells using 500 ng of PEI in 96-well glass bottom plate (Cellvis, P96-1-N). Transfection was performed in triplicate. After 36 hours, the Nano-luciferase activity was measured using Nano-Glo® Live Cell Assay System (Promega, N2011).

### Endogenous KRAS Degradation Assay

1000 ng of binder-LIR fusion plasmids were transfected into U2OS cells by 0.3 uL of Lipofectamin 3000 in a 24-well plate. After 4 h, the media was replaced. After 48 h, the cells were collected and subjected to western blot analysis with the appropriate antibodies.

### Mdm2 binder-Mdm2 Co-Localization Assay

125 ng of the GFP-binder fusion plasmid and 125 ng of the mCherry-Mdm2 fusion plasmid were co-transfected into HEK293T cells using 0.075 µL of Xfect™ Transfection Reagent (Takara Bio, 631317) in 96-well glass bottom plate (Cellvis, P96-1-N). After 4 h, the media was replaced. After 36 h, cells were imaged with a Leica fluorescence microscope.

### Mdm2-p53 Inhibition Assay

1000 ng of Mdm2 binder plasmids were transfected into U2OS cells by 0.3 uL of Xfect™ Transfection Reagent (Takara Bio, 631317) in a 24-well plate. After 4 h, the media was replaced. After 48 h, the cells were collected and subjected to western blot analysis with the appropriate antibodies.

### Reporting summary

Further information on research design is available in the Nature Portfolio Reporting Summary linked to this article.

## Data Availability

Links to electronic vector maps are included in Supplementary Information. Key vectors will be deposited with Addgene and all physical vectors will be made available upon reasonable request. Source data are provided with paper.

## Acknowledgements

This work was supported by the National Institute of General Medical Sciences (GM119840 to B.C.D and F32GM147968 to M.J.S) and then National Cancer Institute (P30CA014599) of the National Institutes of Health, and by the Camille and Henry Dreyfus Foundation Teacher Scholar Award (B.C.D.). We thank S. Ahmadiantehrani for assistance with preparing this paper.

## Contributions

Conceptualization: M.J.S. and B.C.D. Methodology: M.J.S. Investigation: M.S., J.A.P.; T.W.; C.B.; and S.L. Writing – original draft: M.J.S., J.A.P., and B.C.D. Writing – reviewing and editing: T.W. Supervision: B.C.D.

## SUPPORTING INFORMATION

Details of bacterial strains, plasmids, primers, additional data figures and tables.

## References

1 Bandrowski, A., Pairish, M., Eckmann, P., Grethe, J. & Martone, M. E. The Antibody Registry: ten years of registering antibodies. Nucleic Acids Res 51, D358–D367 (2023). 10.1093/nar/gkac927

2 Stanton, B. Z., Chory, E. J. & Crabtree, G. R. Chemically induced proximity in biology and medicine. Science 359 (2018). 10.1126/science.aao5902

3 Park, M. Surface Display Technology for Biosensor Applications: A Review. Sensors (Basel) 20 (2020). 10.3390/s20102775

4 Carter, P. J. & Lazar, G. A. Next generation antibody drugs: pursuit of the ‘high-hanging fruit’. Nat Rev Drug Discov 17, 197–223 (2018). 10.1038/nrd.2017.227

5 Ayoubi, R. et al. Scaling of an antibody validation procedure enables quantification of antibody performance in major research applications. Elife 12 (2023). 10.7554/eLife.91645

6 Laustsen, A. H., Greiff, V., Karatt-Vellatt, A., Muyldermans, S. & Jenkins, T. P. Animal Immunization, in Vitro Display Technologies, and Machine Learning for Antibody Discovery. Trends Biotechnol 39, 1263–1273 (2021). 10.1016/j.tibtech.2021.03.003

7 Sidhu, S. S., Lowman, H. B., Cunningham, B. C. & Wells, J. A. Phage display for selection of novel binding peptides. Methods Enzymol 328, 333–363 (2000). 10.1016/s0076-6879(00)28406-1

8 Xie, V. C., Styles, M. J. & Dickinson, B. C. Methods for the directed evolution of biomolecular interactions. Trends Biochem Sci 47, 403–416 (2022). 10.1016/j.tibs.2022.01.001

9 Wellner, A. et al. Rapid generation of potent antibodies by autonomous hypermutation in yeast. Nat Chem Biol 17, 1057–1064 (2021). 10.1038/s41589-021-00832-4

10 Philpott, D. N. et al. Rapid On-Cell Selection of High-Performance Human Antibodies. ACS Cent Sci 8, 102–109 (2022). 10.1021/acscentsci.1c01205

11 Lopez-Morales, J. et al. Protein Engineering and High-Throughput Screening by Yeast Surface Display: Survey of Current Methods. Small Science 3 (2023). 10.1002/smsc.202300095

12 Porebski, B. T. et al. Rapid discovery of high-affinity antibodies via massively parallel sequencing, ribosome display and affinity screening. Nat Biomed Eng 8, 214–232 (2024). 10.1038/s41551-023-01093-3

13 McConnell, A., Batten, S. L. & Hackel, B. J. Determinants of Developability and Evolvability of Synthetic Miniproteins as Ligand Scaffolds. J Mol Biol 435, 168339 (2023). 10.1016/j.jmb.2023.168339

14 Kordon, S. P. et al. Isoform- and ligand-specific modulation of the adhesion GPCR ADGRL3/Latrophilin3 by a synthetic binder. Nat Commun 14, 635 (2023). 10.1038/s41467-023-36312-7

15 Cao, L. et al. Design of protein-binding proteins from the target structure alone. Nature 605, 551–560 (2022). 10.1038/s41586-022-04654-9

16 Bennett, N. R. et al. Atomically accurate de novo design of single-domain antibodies. bioRxiv (2024). 10.1101/2024.03.14.585103

17 Sappington, I. et al. Improved protein binder design using beta-pairing targeted RFdiffusion. bioRxiv (2024). 10.1101/2024.10.11.617496

18 Huang, B. et al. Designed endocytosis-inducing proteins degrade targets and amplify signals. Nature (2024). 10.1038/s41586-024-07948-2

19 Pu, J., Zinkus-Boltz, J. & Dickinson, B. C. Evolution of a split RNA polymerase as a versatile biosensor platform. Nat Chem Biol 13, 432–438 (2017). 10.1038/nchembio.2299

20 Esvelt, K. M., Carlson, J. C. & Liu, D. R. A system for the continuous directed evolution of biomolecules. Nature 472, 499–503 (2011). 10.1038/nature09929

21 Zinkus-Boltz, J., DeValk, C. & Dickinson, B. C. A Phage-Assisted Continuous Selection Approach for Deep Mutational Scanning of Protein-Protein Interactions. ACS Chem Biol 14, 2757–2767 (2019). 10.1021/acschembio.9b00669

22 He, H., Zhou, C. & Chen, X. ATNC: Versatile Nanobody Chimeras for Autophagic Degradation of Intracellular Unligandable and Undruggable Proteins. J Am Chem Soc 145, 24785–24795 (2023). 10.1021/jacs.3c08843

23 Xie, V. C., Pu, J., Metzger, B. P., Thornton, J. W. & Dickinson, B. C. Contingency and chance erase necessity in the experimental evolution of ancestral proteins. Elife 10 (2021). 10.7554/eLife.67336

24 Miller, S. M. et al. Continuous evolution of SpCas9 variants compatible with non-G PAMs. Nat Biotechnol 38, 471–481 (2020). 10.1038/s41587-020-0412-8

25 Hubbard, B. P. et al. Continuous directed evolution of DNA-binding proteins to improve TALEN specificity. Nat Methods 12, 939–942 (2015). 10.1038/nmeth.3515

26 Packer, M. S., Rees, H. A. & Liu, D. R. Phage-assisted continuous evolution of proteases with altered substrate specificity. Nat Commun 8, 956 (2017). 10.1038/s41467-017-01055-9

27. Thuronyi, B. W., et al. Continuous evolution of base editors with expanded target compatibility and improved activity. Nat Biotechnol 37, 1070–1079 (2019). 10.1038/s41587-019-0193-0

28 Woldring, D. R., Holec, P. V., Stern, L. A., Du, Y. & Hackel, B. J. A Gradient of Sitewise Diversity Promotes Evolutionary Fitness for Binder Discovery in a Three-Helix Bundle Protein Scaffold. Biochemistry 56, 1656–1671 (2017). 10.1021/acs.biochem.6b01142

29 Behar, G. et al. Whole-bacterium ribosome display selection for isolation of highly specific anti-Staphyloccocus aureus Affitins for detection- and capture-based biomedical applications. Biotechnol Bioeng 116, 1844–1855 (2019). 10.1002/bit.26989

30 Hu, J. H. et al. Evolved Cas9 variants with broad PAM compatibility and high DNA specificity. Nature 556, 57–63 (2018). 10.1038/nature26155

31 Dewey, J. A., Azizi, S. A., Lu, V. & Dickinson, B. C. A System for the Evolution of Protein-Protein Interaction Inducers. ACS Synth Biol 10, 2096–2110 (2021). 10.1021/acssynbio.1c00276

32 Block, C., Janknecht, R., Herrmann, C., Nassar, N. & Wittinghofer, A. Quantitative structure-activity analysis correlating Ras/Raf interaction in vitro to Raf activation in vivo. Nat Struct Biol 3, 244–251 (1996). 10.1038/nsb0396-244

33 Spencer-Smith, R. et al. Inhibition of RAS function through targeting an allosteric regulatory site. Nat Chem Biol 13, 62–68 (2017). 10.1038/nchembio.2231

34 Grimm, S., Salahshour, S. & Nygren, P. A. Monitored whole gene in vitro evolution of an anti-hRaf-1 affibody molecule towards increased binding affinity. N Biotechnol 29, 534–542 (2012). 10.1016/j.nbt.2011.10.008

35 Karlsson, G. B. et al. Activation of p53 by scaffold-stabilised expression of Mdm2-binding peptides: visualisation of reporter gene induction at the single-cell level. Br J Cancer 91, 1488–1494 (2004). 10.1038/sj.bjc.6602143

36 Jing, L. et al. Screening and production of an affibody inhibiting the interaction of the PD-1/PD-L1 immune checkpoint. Protein Expr Purif 166, 105520 (2020). 10.1016/j.pep.2019.105520

37 Abramson, J. et al. Accurate structure prediction of biomolecular interactions with AlphaFold 3. Nature 630, 493–500 (2024). 10.1038/s41586-024-07487-w

38 He, H., Zhou, C. & Chen, X. ATNC: Versatile Nanobody Chimeras for Autophagic Degradation of Intracellular Unligandable and Undruggable Proteins. Journal of the American Chemical Society 145, 24785–24795 (2023). 10.1021/jacs.3c08843

39 Dixon, A. S. et al. NanoLuc Complementation Reporter Optimized for Accurate Measurement of Protein Interactions in Cells. ACS Chem Biol 11, 400–408 (2016). 10.1021/acschembio.5b00753

40 Miller, S. M., Wang, T. & Liu, D. R. Phage-assisted continuous and non-continuous evolution. Nat Protoc 15, 4101–4127 (2020). 10.1038/s41596-020-00410-3

41 Rentero Rebollo, I., Sabisz, M., Baeriswyl, V. & Heinis, C. Identification of target-binding peptide motifs by high-throughput sequencing of phage-selected peptides. Nucleic Acids Res 42, e169 (2014). 10.1093/nar/gku940

42 Mirdita, M. et al. ColabFold: making protein folding accessible to all. Nat Methods 19, 679–682 (2022). 10.1038/s41592-022-01488-1

43 Evans, R. et al. (2022). 10.1101/2021.10.04.463034

44 Mouratou, B. et al. Remodeling a DNA-binding protein as a specific in vivo inhibitor of bacterial secretin PulD. Proc Natl Acad Sci U S A 104, 17983–17988 (2007). 10.1073/pnas.0702963104

