## Supplementary Information for "High-throughput protein binder discovery by rapid in vivo selection" (from another version of this preprint: version 1, 2025-01-06)

**TABLE OF CONTENTS**

**Supplementary Table 1: Plasmids**

**Supplementary Table 2: Primers**

**Supplementary Table 3: Information on each target protein of interest**

**Supplementary Fig. 1: Affibody library construction**

**Supplementary Fig. 2: PACE of RAF and IFNG using affibody library**

**Supplementary Fig. 3: PACS with mock library**

**Supplementary Fig. 4: Rate of phage infection**

**Supplementary Fig. 5: Prediction of PANCS Titers based on amplification rates**

**Supplementary Fig. 6: PANCS with mock library and varied +APs**

**Supplementary Fig. 7: PANCS with mock library and varied -APs**

**Supplementary Fig. 8: PANCS with mock library and varied transfer rates**

**Supplementary Fig. 9: PANCS with mock library and varied initial cell-to-phage ratio**

**Supplementary Fig. 10: Mock library selections with additional binder-target pairs**

**Supplementary Table 4: Titers and plaque assays for *de novo* PANCS on 6-target panel**

**Supplementary Fig. 11: Monitoring enrichment across passages by NGS**

**Supplementary Fig. 12: AlphaFold Multimer predictions for binders from 6-target panel PANCS**

**Supplementary Fig. 13: Specificity of binders from 6-target panel PANCS**

**Supplementary Fig. 14: Purification of binders**

**Supplementary Fig. 15: SPR for binders from 6-target panel PANCS**

**Supplementary Fig. 16: Replicate PANCS for 6-target panel**

**Supplementary Fig. 17: Comparison of parallel replicate PANCS by NGS**

**Supplementary Fig. 18: Extending the linker length connecting the target and $RNAP_C$**

**Supplementary Fig. 19: Affitin library construction**

**Supplementary Fig. 20: Proximity independent, $N_{WT}$ phage amplification rates on 96-target panel**

**Supplementary Fig. 21: 96-target panel PANCS titer heatmap**

**Supplementary Table 5: Titers for *de novo* selection on 96-target panel PANCS**

**Supplementary Note S1: Setting a threshold endpoint titer for selections**

**Supplementary Fig. 22: Analysis of NGS for hits from 96-target panel PANCS**

**Supplementary Fig. 23: Luciferase assay results for all preliminary hits**

**Supplementary Fig. 24: Summary of NGS, AlphaFold, and luciferase assay results**

**Supplementary Fig. 25: NGS data for PANCS wells that were just under the threshold**

**Supplementary Fig. 26: Additional passaging 96-target panel PANCS to further separate hits from failures**

**Supplementary Fig. 27: AlphaFold3 predictions of all binder-target pairs testing Figure 4C**

**Supplementary Table 6: Constructing the $10^{10}$ affibody and affitin libraries**

**Supplementary Table 7: Endpoint titers for $10^{10}$ library PANCS**

**Supplementary Fig. 28: NGS and AlphaFold3 predictions of hits from$10^{10}$ library PANCS**

**Supplementary Fig. 29: Luciferase assay for hits from$10^{10}$ library PANCS**

**Supplementary Fig. 30: SPR for KRAS G12D and RAF affibody hits from$10^{10}$ library PANCS**

**Supplementary Fig. 31: SPR for PACE evolved Mdm2 binders**

**Supplementary Table 8: Compiled SPR model fits (for all SPR data)**

**Supplementary Fig. 32: AlphaFold predictions compared to *in vitro* binding affinities**

**Supplementary Fig. 33: KRAS degradation**

**Supplementary Fig. 34: Mdm2 binder co-localization**

**Supplementary Fig. 35: Mdm2-p53 inhibition**

**Supplementary Table 1: Plasmids**

**Phage**

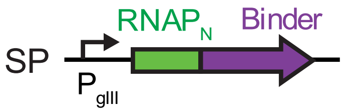


| ID | Description | Binder AA Sequence | Plasmid map |
|---|---|---|---|
| 70-70 | Gen 2 RNAP$_N$-RAF (WT) | SKTSNTIRVFLPNKQRTVVNVRNGMSLHDCLMKALKVRGLQPECCAVFRLLHEHKGKKARLDWNTDAASLIGEELQVDFL | https://benchling.com/s/seq-JJq1S2iNyzsJmd6xiEdv?m=slm-aRoQFIVy6IMYBSH2fwfJ |
| 70-71 | Gen 2 RNAP$_N$-RAF (Q66A) | SKTSNTIRVFLPNKARTVVNVRNGMSLHDCLMKALKVRGLQPECCAVFRLLHEHKGKKARLDWNTDAASLIGEELQVDFL | https://benchling.com/s/seq-rY6quiRgFHXfckenpJEf?m=slm-gCc7foLcfXRHKLDA5uAc |
| 70-72 | Gen 2 RNAP$_N$-RAF (K84A) | SKTSNTIRVFLPNKQRTVVNVRNGMSLHDCLMAALKVRGLQPECCAVFRLLHEHKGKKARLDWNTDAASLIGEELQVDFL | https://benchling.com/s/seq-nSr0g01RqNMCJJ2a4g5L?m=slm-xLrFr8p8h2BD9f9gzg4V |
| 70-73 | Gen 2 RNAP$_N$-RAF (R89L) | SKTSNTIRVFLPNKQRTVVNVRNGMSLHDCLMKALKVLGLQPECCAVFRLLHEHKGKKARLDWNTDAASLIGEELQVDFL | https://benchling.com/s/seq-iIbNxc5abNZFszpmc3xa?m=slm-d6rIiU34PHGHWuq48pRh |
| 70-78 | Gen 2 RNAP$_N$-Affitin (SasA) | GSVKVKFVVRGEEKEVDTSKIRSVWRGGKWVDFSYDDNGKQGYGYVLEKDAPKELLDMLARAEREKKL | https://benchling.com/s/seq-kGbU1x3dZtABiwGPTOvT?m=slm-ktQkaNLZJpvR0y70HFcJ |
| 70-74 | Gen 2 RNAP$_N$-Monobody (KRAS) | GSVSSVPTKLEVVAATPTSLLISWDAPAVTVDYYVITYGETGGNSPVQKFEVPGSKSTATISGLKPGVDYTITVYAWGWHGQVYYYMGSPISINYRT | https://benchling.com/s/seq-0OmrO7iWfuJUJs4VUj3m?m=slm-ZzNRWIOe8dXJESbPQ4yf |
| 70-80 | Gen 2 RNAP$_N$-Monobody (hSUMO1) | VSSVPTKLEVVAATPTSLLISWDAPAVTVDHYVITYGETGGNSPVQEFTVPGSKSTATISGLSPGVDYTITVYAWVSDEYTHGYYSYSPISINYRT | https://benchling.com/s/seq-BMAv02BGSfvUsKloWoMq?m=slm-k4Z8qtY23ikBzE9EvwAc |
| 70-81 | Gen 2 RNAP$_N$-Affibody (RAF) | VDNKFNKEVNLAADEIWLLPNLNNQQAWAFITSLKDDPSQSANLLAEAKKLNDAQAPK | https://benchling.com/s/seq-jlkztBe79lrkgW3fBhwO?m=slm-HWw47rRT6Gkt6WjA4nRo |
| 75-103 | Gen 2 RNAP$_N$-Affibody (PDL1) | VDNKFNKEPRAARLEITVLPNLNREQGGAFIVSLWDDPSQSANLLAEAKKLNDAQAPK | https://benchling.com/s/seq-w0kNAOKhSETKaNFpG3r2?m=slm-ZXJq3YteiKU5l3VNQVs7 |
| 73-9 | Gen 2 RNAP$_N$-Affitin (SasA) 3x STOP | GSVKVKF***GEEKEVDTSKIRSVWRGGKWVDFSYDDNGKQGYGYVLEKDAPKELLDMLARAEREKKL | https://benchling.com/s/seq-tdkTtlPlgVCtBPQclQDf?m=slm-lg2FWZ0yPhL7lZC5Uuyn |
| Lib 5 | Gen 2 RNAP$_N$-Affibody Library | VDNKF?KE???A???I??LPNLN??Q??AF??SL??DPSQSANLLAEAKKLNDAQAPK | https://benchling.com/s/seq-02e90z0Dmx8QenFnZSty?m=slm-OkP6P9eDWs34LeLC4wRx |
| Lib 8 | Gen 2 RNAP$_N$-Affiitn Library | GSVKVKF???GEEKEVDTSKI??V?R?GK?V?F?YDDNGK?G?G?V?EKDAPKELLDMLARAEREKKL | https://benchling.com/s/seq-fTpmkq9ednWVnGyH7r59?m=slm-nwE4qG2wmIsMLJcMUDpJ |
| 70-63 | Gen1 RNAP$_N$ (WT)- IFNG (not PPI dependent) | MQDPYVKEAENLKKYFNAGHSDVADNGTLFLGILKNWKEESDRKIMQSQIVSFYFKLFKNFKDDQSIQKSVETIKEDMNVKFFNSNKKKRDDFEKLTNYSVTDLNVQRKAIHELIQVMAELSPAAKTGKRKRSQ | https://benchling.com/s/seq-SU7Qimp1V6bYvoc7zZex?m=slm-JcqTYIMgGMwxBlz1DvSX |

**+APs**

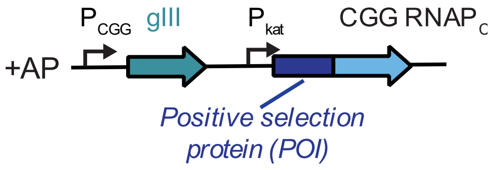


| ID | Target | ORI, RBS ($P_{CGG}$)/($P_{kat}$) | Plasmid Map |
|---|---|---|---|
| 6 AA Linker Sequence: TSGGSG | | | |
| 31-69 | KRAS (WT) | p15a, SD8/SD8; 6 AA linker | https://benchling.com/s/seq-QorElY6vvP0bDTiVgCUN?m=slm-kgoT1gDlDSgp3nNcP5H5 |
| 31-70 | KRAS (WT) | pSC101, SD8/SD8; 6 AA linker | https://benchling.com/s/seq-uoeGETX8o6z6qTEIVTPn?m=slm-p6KSgwAx4P8nB3U49dHb |
| 67-130 | KRAS (G12D) | p15a, SD8/SD8; 6 AA linker | https://benchling.com/s/seq-poKVGTKsSekoJzf0µLBD?m=slm-aeOcCRLcCJpb1zrrXo7l |
| 66-5 | Mdm2 (1-188) | p15a, SD8/SD8; 6 AA linker | https://benchling.com/s/seq-SRsejdlxiEms9jFC0ibj?m=slm-1aB34M8HQwvzADSw7hCs |
| 66-6 | IFNG | p15a, SD8/SD8; 6 AA linker | https://benchling.com/s/seq-cE41F90kMq2nnML3QDax?m=slm-YQGlywLAKMJQDZGARKrH |
| 67-49 | Sos1 | p15a, SD8/SD8; 6 AA linker | https://benchling.com/s/seq-33luLfC8RYnwpA6Vxqyd?m=slm-KviINCCSUz1RAfU78r3A |
| 8-37 | Myc DBD | p15a, SD8/SD8; 6 AA linker | https://benchling.com/s/seq-GCrOfOj1C7ES7b84OlZ6?m=slm-kgth2LUXb87ZnDX7f8Yt |
| 73-40 | RAF RBD | p15a, SD8/SD8; 6 AA linker | https://benchling.com/s/seq-LhbbW3RWsk8tXV9MpSbw?m=slm-gOUPJK3YD23OUQX50WoG |
| 73-66 | RAF RBD | pSC101, SD8/SD8; 6 AA linker | https://benchling.com/s/seq-zPDFzGI2710ZcxBfVXmC?m=slm-3KED3hx4BloSKvG0qJ6s |
| 73-67 | Mdm2 (1-188) | pSC101, SD8/SD8; 6 AA linker | https://benchling.com/s/seq-11PJ9yjPBiCrT3E7ogok?m=slm-GPlAbKKKKlgUjsNy6lwK |
| 73-65 | Mdm2 (1-188) | p15a, SD8/sd5; 6 AA linker | https://benchling.com/s/seq-BFCFEgkEqQQSgZvvdTKn?m=slm-ujGzZ2gWiXd2OUxBREli |
| 79-128 | IFNG | pSC101, SD8/SD8; 6 AA linker | https://benchling.com/s/seq-HAHkYMDpp4nMn9BYp7vr?m=slm-TLI4ukXqGrZkvPFhwpEI |
| 73-63 | IFNG | p15a, SD8/sd5; 6 AA linker | https://benchling.com/s/seq-jjYgcBHgpvQ3j5IxrSXU?m=slm-CDjEcS1ZXe44iDHPe6jN |
| 20 AA Linker Sequence: TSGGSGGSGSSGSGGTSGSG | | | |
| 73-31 | KRAS (G12D) | p15a, SD8/SD8; 20 AA linker | https://benchling.com/s/seq-5wpJUNYgrOkT5qjh8mhV?m=slm-fgsFkeIA99Ur5wTWEFob |
| 73-41 | RAF RBD | p15a, SD8/SD8; 20 AA linker | https://benchling.com/s/seq-QrfAd2dknEc0GXwKmIHj?m=slm-cbzOGy0XpZlaQM0NllU3 |
| 73-34 | ZB | p15a, SD8/SD8; 20 AA linker | https://benchling.com/s/seq-M0fP6w4gtYx8laJwX4D3?m=slm-MW3QPL7xWaJz78SXsTz6 |
| 40 AA Linker Sequence: TSGGSGGSGSSGSGGTSGSGGSGSGTGSGSSGSGGTSGSG | | | |
| 73-32 | KRAS (G12D) | p15a, SD8/SD8; 40 AA linker | https://benchling.com/s/seq-XbAD0KtbE3nhoaX664OO?m=slm-uvktq1pE8pMiLJJ5FwcN |
| 73-42 | RAF RBD | p15a, SD8/SD8; 40 AA linker | https://benchling.com/s/seq-6wAbVdVuhWk9PosRBmHH?m=slm-L9XAD432ChIUuwL6q35O |
| 73-35 | ZB | p15a, SD8/SD8; 40 AA linker | https://benchling.com/s/seq-vFjmVRA3rWP6JOKIWC0s?m=slm-INqCQEzSnM4x7w84aAR9 |
| 60 AA Linker Sequence: TSGGSGGSGSSGSGGTSGSGGSGTGTGTGSSGSGGTSGSGGSGSGTGSGSSGSGGTSGSG | | | |

| | | | |
|---|---|---|---|
| 75-154 | No Fusion | p15a, SD8/SD8; 60 AA linker | https://benchling.com/s/seq-OAkkUSRN3TJhdoLilhRD?m=slm-bEfoe3bp41FrsbQWjGA9 |
| 73-21 | KRAS (G12D) | p15a, SD8/SD8; 60 AA linker | https://benchling.com/s/seq-Zr3IVLYCsltuBwWaWxJr?m=slm-wRhiEYUzN1wIXpbqdbih |
| 73-36 | ZB | p15a, SD8/SD8; 60 AA linker | https://benchling.com/s/seq-B3VpJwTHBJNtDB96TRU3?m=slm-iQ706YAxBZMMwGKFHJVF |
| 73-39 | GFP | p15a, SD8/SD8; 60 AA linker | https://benchling.com/s/seq-v5EQQGCh486ebYJKJ51r?m=slm-ypcPwaGL9gxg6AgQIkhn |
| 73-150 | p65 (17-293) | p15a, SD8/SD8; 60 AA linker | https://benchling.com/s/seq-cdSliWqWg0tdHYgBTdOP?m=slm-pEDkDzXE0Z4F8vYUSiit |
| 73-155 | FGFR1c domain D2 | p15a, SD8/SD8; 60 AA linker | https://benchling.com/s/seq-Uzm5jeJGfrrqaKG4tbn6?m=slm-OanxydYCDzyfpqszed47 |
| 73-156 | TCIM | p15a, SD8/SD8; 60 AA linker | https://benchling.com/s/seq-6IddRZ1HDqTN3MCPSaK8?m=slm-Ls84NSQmf0GHZLtK9WM8 |
| 73-157 | TRIM21 (277-475) | p15a, SD8/SD8; 60 AA linker | https://benchling.com/s/seq-oEa6R5xeroyTLQdBFxcy?m=slm-qT4xnij6laqu2uwpCMdM |
| 73-158 | PDE6D | p15a, SD8/SD8; 60 AA linker | https://benchling.com/s/seq-DNJTqsoqchb6Ifc8LmkH?m=slm-S7qdc6giMN9TxTm03COK |
| 75-122 | RAF | p15a, SD8/SD8; 60 AA linker | https://benchling.com/s/seq-653ffOYStxIyWy0crwUG?m=slm-popOBq01ASaK3UElPBLr |
| 75-124 | NRAS | p15a, SD8/SD8; 60 AA linker | https://benchling.com/s/seq-rBJD0ntLGgEuAuq1kSED?m=slm-yshqst61vCjixpRdgNbY |
| 74-29 | Beclin (175-267) | p15a, SD8/SD8; 60 AA linker | https://benchling.com/s/seq-bKebTeEQKLuRZBaobE1r?m=slm-2EpZm0zHwRtbAbAu6Dx8 |
| 75-128 | Mdm2 (1-188) | p15a, SD8/SD8; 60 AA linker | https://benchling.com/s/seq-PxdMGjVyvDqAJDmdvnLQ?m=slm-QGrbcwpi2958fCky1PHM |
| 75-129 | Calpro het 12aa | p15a, SD8/SD8; 60 AA linker | https://benchling.com/s/seq-hjEXy4EpozczGXmnCBaw?m=slm-kaXmkr7hxqA6SVFKn2FS |
| 75-130 | HlgA toxin | p15a, SD8/SD8; 60 AA linker | https://benchling.com/s/seq-Et7y9HSeppcxczv2YEmK?m=slm-cIebjjpTK98RJnKI19EK |
| 75-131 | Bcl2 | p15a, SD8/SD8; 60 AA linker | https://benchling.com/s/seq-RMXbBBMr4cQa8yuBErNu?m=slm-phe0A6NggSSQHM8pYvx9 |
| 75-132 | Calpro mon s100a9 | p15a, SD8/SD8; 60 AA linker | https://benchling.com/s/seq-ACHXVm1JqRuo6vu5yxVR?m=slm-qEU7Hbl2g9DTVirxt3e9 |
| 75-133 | KIX | p15a, SD8/SD8; 60 AA linker | https://benchling.com/s/seq-75hSB7CK7Ss5mdiXwroT?m=slm-dFHu3SzeYs6wCnenE2hQ |
| 75-135 | HRAS | p15a, SD8/SD8; 60 AA linker | https://benchling.com/s/seq-GOe4zjTl1Mdf4PRb31OS?m=slm-aVc4rYGqhQtzmXMU7JX5 |
| 75-136 | KRAS wt | p15a, SD8/SD8; 60 AA linker | https://benchling.com/s/seq-ZeZ4aYcPMa4jt6kE0JA8?m=slm-urQbtWwArEqCnNkcetG1 |
| 75-137 | G3BP | p15a, SD8/SD8; 60 AA linker | https://benchling.com/s/seq-RmHYPArmkMpqTnNpr0OP?m=slm-2kBpsjPaVT6hPhPNxFxx |
| 75-143 | NIX | p15a, SD8/SD8; 60 AA linker | https://benchling.com/s/seq-r3wjG4NZ4yLcCRmEG7Yd?m=slm-P4Nzro5JL8BBBd0ORv5r |
| 75-140 | PP2A-B | p15a, SD8/SD8; 60 AA linker | https://benchling.com/s/seq-18IkmH1oviCvDSJqO8dh?m=slm-I67s5YoyoAAaUOnnArDm |
| 75-141 | Calpro mon s100a8 | p15a, SD8/SD8; 60 AA linker | https://benchling.com/s/seq-1dbycwm28J9gHToTdejh?m=slm-STtWIVscpGItdu8VEBy3 |
| 75-142 | FOXP3 | p15a, SD8/SD8; 60 AA linker | https://benchling.com/s/seq-5RUKlkWIsNYOKRFxp9H8?m=slm-5TvJ2jmfkNogUtIie2Rr |
| 75-138 | KRAS (G12V) | p15a, SD8/SD8; 60 AA linker | https://benchling.com/s/seq-GFtPQ2Xf6XG5izoUTCsA?m=slm-CxBk737fChjA24RXaPyO |
| 75-146 | Sos1 | p15a, SD8/SD8; 60 AA linker | https://benchling.com/s/seq-w80QMJY61wsyX3exlMz4?m=slm-kRREHX73IefmCI7iWAO6 |

| | | | |
|---|---|---|---|
| 75-147 | PP2A-B' | p15a, SD8/SD8; 60 AA linker | https://benchling.com/s/seq-TwI5xEyskYV7Sw9GsjRh?m=slm-TzAhfO0Rng5ajuOAwFgi |
| 75-148 | mCherry | p15a, SD8/SD8; 60 AA linker | https://benchling.com/s/seq-So0ZUDzTf8WRPuUMf9t0?m=slm-bnWoNtGNBoNtU3YjwZPc |
| 75-149 | Myc (short) | p15a, SD8/SD8; 60 AA linker | https://benchling.com/s/seq-EnqwDf6RRDLp9d1xJV4T?m=slm-V2L8jbdYZITnjwyPReYe |
| 75-152 | PP2A-B'' | p15a, SD8/SD8; 60 AA linker | https://benchling.com/s/seq-4gOERZifGohryh49vugB?m=slm-zclfZd87C5VyT8h1OZ68 |
| 75-153 | MBP | p15a, SD8/SD8; 60 AA linker | https://benchling.com/s/seq-CTYBnSJpKhNMPCGe49xt?m=slm-Yy5Ydwaik5tUJ7PRdDW7 |
| 75-155 | Fibronectin | p15a, SD8/SD8; 60 AA linker | https://benchling.com/s/seq-zdsltF3MW4Ugw2tCggjl?m=slm-pY7ugHj1aXcbiLaCPTIR |
| 75-156 | VHL | p15a, SD8/SD8; 60 AA linker | https://benchling.com/s/seq-qb7tgAd3oKFG9Z6EqSdo?m=slm-jGDEhoC2D3a3kO81tTL6 |
| 75-157 | BTK (PH) | p15a, SD8/SD8; 60 AA linker | https://benchling.com/s/seq-5ALCwW2eLTMX3flVJet3?m=slm-sBY39ws0zDeS7oLoG5SH |
| 75-160 | Calpro het 6aa | p15a, SD8/SD8; 60 AA linker | https://benchling.com/s/seq-CBtjqRnxx4ayrkRAR6XS?m=slm-yZoA13Q4kBtyBDHr8w9R |
| 75-161 | GABARAP | p15a, SD8/SD8; 60 AA linker | https://benchling.com/s/seq-CU3BT33LpDQcqNz52AYW?m=slm-pM8ur5mMQ5m8gMYjfW4x |
| 75-164 | GTF2I | p15a, SD8/SD8; 60 AA linker | https://benchling.com/s/seq-D6ZmbBc9m2P7cdeZfjkI?m=slm-wbxKJ1T7nHnx8u0dFvP3 |
| 75-165 | OmpF (ec) | p15a, SD8/SD8; 60 AA linker | https://benchling.com/s/seq-bpykRa7MihJDhXglLuqN?m=slm-pgYxnvoDyuSMfeGKks8P |
| 75-166 | 14-3-3z Monomeric (L12Q, R18E and S58E) | p15a, SD8/SD8; 60 AA linker | https://benchling.com/s/seq-IWbQqjRmQzcjUYtigywe?m=slm-yt4r5Yag09nYxefCzHFJ |
| 75-167 | PCNA-AF | p15a, SD8/SD8; 60 AA linker | https://benchling.com/s/seq-XJUWLK9eWZX7HAWauHcu?m=slm-oDX0s1RnucKQJcBeHREu |
| 75-169 | TEV | p15a, SD8/SD8; 60 AA linker | https://benchling.com/s/seq-81ffIJaX0kRAI6VLInsY?m=slm-galMvGSadT65JKoFySYK |
| 75-170 | FnbpA | p15a, SD8/SD8; 60 AA linker | https://benchling.com/s/seq-GjkFFNf2siUBskbdkmDc?m=slm-ASm8m8Enw4i3jjTBtBmf |
| 75-171 | IBTK | p15a, SD8/SD8; 60 AA linker | https://benchling.com/s/seq-Fy1aT64LESd9RoHJb4yy?m=slm-nfQUjcyqVNACboxbxfRP |
| 75-172 | PAL | p15a, SD8/SD8; 60 AA linker | https://benchling.com/s/seq-ecoHsMaIhmdTYwSoQV5i?m=slm-c8zDVquNiy8OOq62bqlL |
| 75-173 | PIK3CB (3-115) | p15a, SD8/SD8; 60 AA linker | https://benchling.com/s/seq-7A7cpLNdlIJM7KkPgZaS?m=slm-Y9jqMOLRNBhwlpGµLJOH |
| 75-174 | SortaseA (25-206) | p15a, SD8/SD8; 60 AA linker | https://benchling.com/s/seq-jGkTemzmKXVsPiaVzTmO?m=slm-jMbvmE2g0pp16TVTSZ0c |
| 75-176 | PilQ (468-549) | p15a, SD8/SD8; 60 AA linker | https://benchling.com/s/seq-khnmHmrdshMd7UFhA7fe?m=slm-vsPUDoswrA1dUAEs8dIZ |
| 75-177 | DltD (ef) | p15a, SD8/SD8; 60 AA linker | https://benchling.com/s/seq-pr9kZE87jy877ZeV2lwM?m=slm-01vwtAKq8spWRM1IWVcC |
| 75-178 | TNFR2 (33-205) | p15a, SD8/SD8; 60 AA linker | https://benchling.com/s/seq-mAyNSHYPq2EBro3MaKM7?m=slm-x8FVA5aJGLiVHCDET5Li |
| 75-179 | CDKN1A | p15a, SD8/SD8; 60 AA linker | https://benchling.com/s/seq-PQ5irZBkcDjh3R4UHBKN?m=slm-cFdFMaHd2Gl1r6awo41C |
| 75-180 | SH3BP5 | p15a, SD8/SD8; 60 AA linker | https://benchling.com/s/seq-Avc55NRDP6F4mjVqeGlV?m=slm-o4BfkBdFLzii1YkDeJOP |

| | | | |
|---|---|---|---|
| 75-181 | OTUBI (1-102) | p15a, SD8/SD8; 60 AA linker | https://benchling.com/s/seq-O82hpMpccs8FoUNuFrfY?m=slm-DGRFyYlG6QHybTY31P3b |
| 75-182 | DltD (strep) | p15a, SD8/SD8; 60 AA linker | https://benchling.com/s/seq-CECchAZI4HL7qEmLC3np?m=slm-tgKN7aPE8K1Kr0fPaaS7 |
| 75-150 | PINK | p15a, SD8/SD8; 60 AA linker | https://benchling.com/s/seq-qPq6x39UehzUtZRCla0I?m=slm-yHzeA88dr7KzF6ILd18b |
| 75-185 | RB1 | p15a, SD8/SD8; 60 AA linker | https://benchling.com/s/seq-22kbw64bDo5f2vL4T6J3?m=slm-IGZoUdtzriYriAAKfviD |
| 75-186 | SasG (1060-1140) | p15a, SD8/SD8; 60 AA linker | https://benchling.com/s/seq-vlqOK4ok3Nnz2RAIsmeV?m=slm-TGfDQbkCbhmIYbt8k1cU |
| 75-187 | SasG (930-1060) | p15a, SD8/SD8; 60 AA linker | https://benchling.com/s/seq-iLLWrGfjClMGYArCPXsI?m=slm-Fhe4Uyj9NfoC8CywBOKA |
| 75-188 | FKBP | p15a, SD8/SD8; 60 AA linker | https://benchling.com/s/seq-x40cA8lpEv7xirBM9uxs?m=slm-asEJgRJgltfptOrbcjjj |
| 75-190 | BRD4 | p15a, SD8/SD8; 60 AA linker | https://benchling.com/s/seq-uWLE77Emo9YGUtV2nx5v?m=slm-WmFAuRn2lQPoZsF4hFHX |
| 75-191 | IsdB (125-269) | p15a, SD8/SD8; 60 AA linker | https://benchling.com/s/seq-SzEr4lZJNcBJrnUB8QO2?m=slm-seJsEbcdhbgolFAfQwpN |
| 76-50 | HEWL | p15a, SD8/SD8; 60 AA linker | https://benchling.com/s/seq-CxlSVSbklEqF3Syw042I?m=slm-H3KNC82Pha4FHZB37MbH |
| 75-193 | HSPB1 (90-171) | p15a, SD8/SD8; 60 AA linker | https://benchling.com/s/seq-PJZGJ4mun3IKcOJErxe0?m=slm-Xs5wlOr4SfvbdgRUM6Ed |
| 75-194 | KRAS G12C | p15a, SD8/SD8; 60 AA linker | https://benchling.com/s/seq-2tvdweDEqlYO5sPUNAVY?m=slm-csLi8d4weLPbhqzzlYOU |
| 76-94 | Mst2 linker | p15a, SD8/SD8; 60 AA linker | https://benchling.com/s/seq-BemPhamuT5sywqWLa5h5?m=slm-Ymel1P0qdFuIUAm0Sf6u |
| 76-51 | LMO2 (28-150) | p15a, SD8/SD8; 60 AA linker | https://benchling.com/s/seq-NXc2ysSollG8dC1PXNgL?m=slm-YfpX1I0eijcrk5tYWpZ8 |
| 76-52 | HIF1a | p15a, SD8/SD8; 60 AA linker | https://benchling.com/s/seq-LrW4a6u72I2krof0lNiK?m=slm-Fo74FDHKJbx8z2PrQe0z |
| 76-23 | MXD1 | p15a, SD8/SD8; 60 AA linker | https://benchling.com/s/seq-bBeNxjomjgPQRSFiWGOn?m=slm-QyDn8Obf66PvZrYt6iKI |
| 75-195 | OmpA (ec, 84-224) | p15a, SD8/SD8; 60 AA linker | https://benchling.com/s/seq-qGyYe5Xj4xBsdfY7kAAg?m=slm-3Pw3eZnnYOG1s1rfDqRN |
| 76-40 | CD3z | p15a, SD8/SD8; 60 AA linker | https://benchling.com/s/seq-fF2AIC8XFYµLI5dEVeJk?m=slm-qUnT9qGpKKdfn3OOS1p1 |
| 76-41 | PIK3CA (1-105) | p15a, SD8/SD8; 60 AA linker | https://benchling.com/s/seq-Aned8XSfCRP1MoDFpYwR?m=slm-9Q9yuEB1ZIsZuzYopY4X |
| 76-44 | UEV | p15a, SD8/SD8; 60 AA linker | https://benchling.com/s/seq-WTDkd5RV746qVTqvZFsw?m=slm-l1U9FTb2LkTO29rWI5Wp |
| 76-46 | FRB | p15a, SD8/SD8; 60 AA linker | https://benchling.com/s/seq-mtFkW0QpGYU17MfoCaHn?m=slm-P9kNCCZeZbWASii3O9Bx |
| 76-47 | PD-1 | p15a, SD8/SD8; 60 AA linker | https://benchling.com/s/seq-bKsWCHYUYtEDZhayfEFE?m=slm-kkprEYggLqVT9u5bZgwT |
| 76-49 | EIF4A | p15a, SD8/SD8; 60 AA linker | https://benchling.com/s/seq-8mGcOba7WoGgvwySy429?m=slm-CzuPCOp2kSy8ZxM6VD7V |
| 75-123 | SasA | p15a, SD8/SD8; 60 AA linker | https://benchling.com/s/seq-6BAA9bssJHU2rrzvFfnQ?m=slm-BeLryRJBSo2jGsp6ZaQ2 |
| 75-126 | LC3B | p15a, SD8/SD8; 60 AA linker | https://benchling.com/s/seq-BVnxoGkEIF8WqzyciCqM?m=slm-zsHlH0xb2vn5ws0dScqE |
| 75-127 | ALKBH5 | p15a, SD8/SD8; 60 AA linker | https://benchling.com/s/seq-4J5UYICjPsrfCcBQexOw?m=slm-62da3X8otCUcHupJT2yw |
| 75-134 | HIF1b (1-474) | p15a, SD8/SD8; 60 AA linker | https://benchling.com/s/seq-ao4jpxAORmJ8nMHYWkdF?m=slm-TRnAWFqxP1nEEQ8iCra6 |

| | | | |
|---|---|---|---|
| 75-158 | IFNG (24-156) | p15a, SD8/SD8; 60 AA linker | https://benchling.com/s/seq-KsZlMXJQ7LRRHC7GSqti?m=slm-yIqgGLjFLyxIbwwJ2he8 |
| 75-144 | MST2 (313-437) | p15a, SD8/SD8; 60 AA linker | https://benchling.com/s/seq-jtOStbjKLwrI2hWHuPye?m=slm-z79ntiBq1r4xrktdn7Qj |
| 75-145 | p53 (1-293) | p15a, SD8/SD8; 60 AA linker | https://benchling.com/s/seq-xaOF9j7eTve5wPY7OSzA?m=slm-X1JsS1Y215Zor6alp6ry |
| 76-48 | PD-L1 | p15a, SD8/SD8; 60 AA linker | https://benchling.com/s/seq-B74sRi7ocrSyHvdQiQVZ?m=slm-xLrtaQpvUHFhWyCJ84mv |
| 76-1 | Bax peptide | p15a, SD8/SD8; 60 AA linker | https://benchling.com/s/seq-8J7TAMAqN4ZLESQkEQ4G?m=slm-SFiX8Vx8PpbEJ9s39bMJ |
| 75-151 | p53 AD (1-61) | p15a, SD8/SD8; 60 AA linker | https://benchling.com/s/seq-2OLFYp087OjPugBjKupY?m=slm-YlQqlGdSkzx0rHiTQfkZ |
| 76-93 | CD3d | p15a, SD8/SD8; 60 AA linker | https://benchling.com/s/seq-4hR2jNVµLOXJPDLGbX8B?m=slm-P33HEy8NnH2bAUbX7wbM |
| 75-159 | Myc (long) | p15a, SD8/SD8; 60 AA linker | https://benchling.com/s/seq-hendvxfIhlwtdR4OwfQv?m=slm-w8EH9I6dGWq16qgNJvYL |
| 75-162 | CRBN | p15a, SD8/SD8; 60 AA linker | https://benchling.com/s/seq-7mvwe76RtMPWca9NQqpC?m=slm-oLSx4DVQMj0Mh58fhIgZ |
| 75-163 | MAX | p15a, SD8/SD8; 60 AA linker | https://benchling.com/s/seq-rtp66xstwB8HFXFEQVje?m=slm-duRSyOvzl1gABrDn4rus |
| 75-168 | PPP1R11 | p15a, SD8/SD8; 60 AA linker | https://benchling.com/s/seq-MbPMAihvvKbWdKD6Jtg0?m=slm-D92oVzXfQµLUeLExAAve |
| 75-175 | Alpha amylase | p15a, SD8/SD8; 60 AA linker | https://benchling.com/s/seq-5emdqxm8ALcLLo7uNPJ6?m=slm-hi6YfuFD8Uc84mYewk3Y |
| 75-183 | PMP22 | p15a, SD8/SD8; 60 AA linker | https://benchling.com/s/seq-sPUgwnqHHCrprRrHXZk7?m=slm-JomYsxi1wgt1rgWU8Fai |
| 75-189 | BAX | p15a, SD8/SD8; 60 AA linker | https://benchling.com/s/seq-dv8svMLWo7jRgBNANgao?m=slm-0XX5aejmRwEcRVHgMrIl |
| 76-42 | Keap1 | p15a, SD8/SD8; 60 AA linker | https://benchling.com/s/seq-EdPHe3rCN1vOvTwxeQCZ?m=slm-KCWjIy09bMTDnKUY92o0 |
| 76-43 | HSPB1 | p15a, SD8/SD8; 60 AA linker | https://benchling.com/s/seq-IfKi5eLDKox1xmAmAeah?m=slm-CwP7gFgtQY7tnrVkHMt5 |
| 76-45 | HMGB1 | p15a, SD8/SD8; 60 AA linker | https://benchling.com/s/seq-5hGuCN4loBaLzdNTzHz2?m=slm-ZdlLMfeR7jQGkp5RVitC |

**-APs**

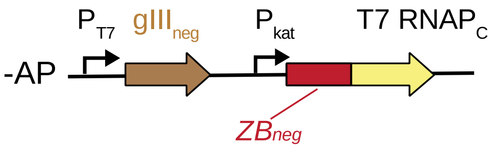


| ID | Description | Plasmid map |
|---|---|---|
| $ZB_{neg}$ | RBS for $P_{T7}$/ RBS for $P_{kat}$ | AA Sequence: MASEQLEKELQALEKELAQLKWKNQALEKKLAQ |
| 20-06 | $ZB_{neg}$, 6 AA linker, sd5/sd2 | https://benchling.com/s/seq-nHRnXEU6UGqfIjfZ9C0L?m=slm-GVJXRV43bBgZm75lqBm7 |
| 20-05 | $ZB_{neg}$, 6 AA linker, sd5/sd5 | https://benchling.com/s/seq-sR9zYNKCh7TIIPkbpZAf?m=slm-EMUTvNHfcWsCxBdhwxRZ |
| 20-01 | $ZB_{neg}$, 6 AA linker, sd8/sd8 | https://benchling.com/s/seq-VAE5JvugbkmYx1AYTWDf?m=slm-sI2hSzJ94ZnivxzSP675 |
| 73-79 | $ZB_{neg}$, 60 AA linker, sd8/sd8 | https://benchling.com/s/seq-qptnks4V9vCAkgQsGtmf?m=slm-0SHHTsCgDhaxvivYq6yY |

**Plasmids for luciferase assay**

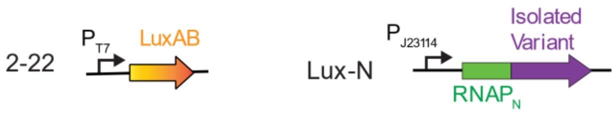


| ID | Description | AA Sequence | Plasmid map |
|---|---|---|---|
| 2-22 | $P_{T7}$ driven LuxAB on pSC101 | | https://benchling.com/s/seq-Udyz6dlESk9lOHBzqfsO?m=slm-pPVnaTxvld7HcIZ8InVO |
| 77-153 | Lux-N Affibody (PDL1) | VDNKFNKEPRAARLEITVLPNLNREQGGAFIVSLWDDPSQSANLLAEAKKLNDAQAPK | https://benchling.com/s/seq-4Hat7A1iLPOjVB5SfuyL?m=slm-kG73luQP4lmO9k1o5Xk0 |
| 77-144 | Lux-N Affitin (SasA) | GSVKVKFVVRGEEKEVDTSKIRSVWRGGKWVDFSYDDNGKQGYGYVLEKDAPKELLDMLARAEREKKL | https://benchling.com/s/seq-MzeKf9F5WI4KL6VcGZvu?m=slm-EWzvy4QEH0p4MplFX396 |
| 77-147 | Lux-N Monobody (hSUMO1) | VSSVPTKLEVVAATPTSLLISWDAPAVTVDHYVITYGETGGNSPVQEFTVPGSKSTATISGLSPGVDYTITVYAWVSDEYTHGYYSYSPISINYRT | https://benchling.com/s/seq-DIRqB3iHFxxrwaJ30gE7?m=slm-ZXarjzck1DUgI2snLzHR |
| 77-154 | Lux-N Affibody (RAF) | VDNKFNKEVNLAADEIWLLPNLNNQQAWAFITSLKDDPSQSANLLAEAKKLNDAQAPK | https://benchling.com/s/seq-BbfsAlyrXYrH5iGF0TXP?m=slm-8PKvi1so0qMW70ijQh4W |
| 77-134 | Lux-N Monobody (KRAS; NS1) | GSVSSVPTKLEVVAATPTSLLISWDAPAVTVDYYVITYGETGGNSPVQKFEVPGSKSTATISGLKPGVDYTITVYAWGWHGQVYYYMGSPISINYRT | https://benchling.com/s/seq-wbjyt230LVnMaXfV9pmI?m=slm-YRG6N9pCEBxRhtvmJQ4P |
| 77-155 | Lux-N Cl2 (Mdm2) | SSVEKKPEGVNTGAGDRHNLKTEWPELVGKSVEEAKKVILQDKPEAQIIVLPVGTMPRFMDYWEGLNRIDRVRLFVDKLDNIAEVPRVG | https://benchling.com/s/seq-LneOyfLblsQ1SqHsoraL?m=slm-TN599lHPKseJUorEDh3L |
| Lux-N Lib 5 | Lux-N Affiobdy Library Variant | VDNKF?KE???A???I??LPNLN??Q??AF??SL??DPSQSANLLAEAKKLNDAQAPK | https://benchling.com/s/seq-mVSKERuVdRGIeQ7j4OKo?m=slm-Hk4Rjrf5ZYCffF4uTrbZ |
| Lux-N Lib 8 | Lux-N Affitin Library Variant | GSVKVKF???GEEKEVDTSKI??V?R?GK?V?F?YDDNGK?G?G?V?EKDAPKELLDMLARAEREKKL | https://benchling.com/s/seq-nSI83aPslWFrsSEeCrpe?m=slm-Uzf6HPcmoEH5NHELKy7K |

| | | | |
|---|---|---|---|
| 65-50 | Lux-N $RNAP_{N(WT)}$-IFNG (not PPI dependent | MQDPYVKEAENLKKYFNAGHSDVADNGTLFLGIL KNWKEESDRKIMQSQIVSFYFKLFKNFKDDQSIQ KSVETIKEDMNVKFFNSNKKKRDDFEKLTNYSVT DLNVQRKAIHELIQVMAELSPAAKTGKRKRSQ | https://benchling.com/s/seq-z6k7WcWzlkehV2RIfEsz?m=slm-nF8asOivpVyPfh7mAXFB |

**Lux-C**

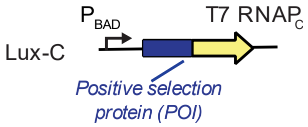


| Lux-C | Target | Lux-C Map |
|---|---|---|
| 6 AA Linker Sequence: TSGGSG | | |
| 12-48 | KRAS G12D | https://benchling.com/s/seq-uiSpEZdsBlTbxgExQKzA?m=slm-tjVz7COoGbkxxZsU8hwQ |
| 12-64 | RAF RBD | https://benchling.com/s/seq-QdDkS0EM8CzsfT20bokO?m=slm-syqroCCl8jVvk119MLUK |
| 72-83 | Mdm2 (1-188) | https://benchling.com/s/seq-qN74BV6NcLVA40q7sWOG?m=slm-tYP1L76qLlDmjmQSlgnI |
| 72-76 | IFNG | https://benchling.com/s/seq-9BsSkfIQe1RzlIl7Lynm?m=slm-buZIVDFvFMxriNzCASlF |
| 60 AA Linker Sequence: TSGGSGGSGSSGSGGTSGSGGSGTGTGTGSSGSGGTSGSGGSGSGTGSGSSGSGGTSGSG | | |
| 77-59 | No Fusion | https://benchling.com/s/seq-yExBSN9d57SyhGQvdX8k?m=slm-4cQuASRmUdkXWzuNG9ol |
| 73-21 | KRAS (G12D) | https://benchling.com/s/seq-bHUdLMQyJLyYaNnCzWYo?m=slm-kqR0h1614DqdUkezQdiP |
| 73-22 | ZB | https://benchling.com/s/seq-cUHUKYXlIfvrpiwQffeK?m=slm-f8l0TBya1R6o360d4148 |
| 73-30 | GFP | https://benchling.com/s/seq-jW7Zau5g2UgK1so86pGd?m=slm-b8yxDRGFVA9FnKMtqaMv |
| 76-152 | p65 (17-293) | https://benchling.com/s/seq-MFerrPNUPyioan1e5CPY?m=slm-7aRHx4Zp6Q72hCtJL1od |
| 76-105 | FGFR1c domain D2 | https://benchling.com/s/seq-9Ex1W5lAuuNbWmXbwFtM?m=slm-EVPbFTSOCZimmw58YBLK |
| 73-167 | TCIM | https://benchling.com/s/seq-nTnoMw1XCU0jQnn1ZXIE?m=slm-BYn2HKhFztBOk5msXYea |
| 73-173 | TRIM21 (277-475) | https://benchling.com/s/seq-qCMpyJNRnolctgFACU7q?m=slm-pOqLvGfB2rU0lwy0zrm3 |
| 76-88 | PDE6D | https://benchling.com/s/seq-8pMSWqjiCrQuri2DX3Kl?m=slm-3knkHTbWEwJnUEY4iboI |
| 73-23 | RAF | https://benchling.com/s/seq-O0XHCpWbTjdtbr6hzREO?m=slm-6HSgWO74Jc1t0JNGMmUi |
| 76-97 | NRAS | https://benchling.com/s/seq-MNEcNbqEA5bJjd5OxWp4?m=slm-kItXGR5rmeQg46qZz5G8 |
| 74-35 | Beclin (175-267) | https://benchling.com/s/seq-5yo7IVzcSnXLhI6MlISI?m=slm-lRI2HFD37sHjuktcnQJv |
| 76-74 | Mdm2 (1-188) | https://benchling.com/s/seq-lTuETCqeoOd8g8H8z2Xm?m=slm-dYuzgB8XONA4AEyrfbsV |
| 76-149 | Calpro het 12aa | https://benchling.com/s/seq-tkeIuocLYFD41AhUPhFY?m=slm-rfu4gttDjm6WRHHhjYi0 |
| 76-145 | HlgA toxin | https://benchling.com/s/seq-4rYld71VjIo3hrfWFvjX?m=slm-DNEtvtTLscs8BId0O0JV |
| 76-99 | Bcl2 | https://benchling.com/s/seq-XXBGkJPjCBIubOQuCogn?m=slm-DRPjkgVNKU0Dowo7PDjX |
| 76-75 | Calpro mon s100a9 | https://benchling.com/s/seq-Pr7YIp6rzdeTHHdAgiKE?m=slm-kCzo82h7fupB5DXLgs47 |
| 76-80 | KIX | https://benchling.com/s/seq-mkPkHaYFfPfnW81wRv1f?m=slm-FalsloRZmTpa36t0T2Ly |
| 76-98 | HRAS | https://benchling.com/s/seq-pgTuUgg65DUpR1fn20uo?m=slm-zeyRxnr9T8TrgjagcSCj |
| 76-96 | KRAS wt | https://benchling.com/s/seq-YNF06s3yMNWhoWHllQZm?m=slm-pOpYJTkPYLfYd25kMJqP |
| 76-106 | G3BP | https://benchling.com/s/seq-p6tyxpHvpRKNiw5c2wfR?m=slm-YTOOFgotSluGRC3VgMAY |
| 76-154 | NIX | https://benchling.com/s/seq-xefyGFfkOtoGCYsSLXtH?m=slm-pKPjhLMHVIswd1rsarhV |
| 76-175 | PP2A-B | https://benchling.com/s/seq-NUTSz9eO3JOJlfiwyrE5?m=slm-yAu6yfORB4nt6Pzc8qrU |
| 77-58 | Calpro mon s100a8 | https://benchling.com/s/seq-41mAWyETIGyDdPQXoXmY?m=slm-RqgYH6D5qrjU2ZN5jgBx |
| | FOXP3 | |
| 76-95 | KRAS (G12V) | https://benchling.com/s/seq-2mdc5SUWhQmRJX7ImMwk?m=slm-3YNe0gdVMwQA84eFeMZJ |
| 76-169 | Sos1 | https://benchling.com/s/seq-BetRKoyZuNBd1nUuMrp6?m=slm-dYRwQhHl0u5AfxcmmTU9 |
| 76-176 | PP2A-B' | https://benchling.com/s/seq-vMWqBFiKweSC3lbfvTet?m=slm-C3zToe5IZgJanvOwg5Jb |
| 76-150 | mCherry | https://benchling.com/s/seq-kYMKyHRGdy2oTZvt2IU3?m=slm-xRUOuQLeTy6oTcayxgBM |

| | | |
|---|---|---|
| 76-73 | Myc (short) | https://benchling.com/s/seq-s86vJay1ws2nFE8d2svR?m=slm-SlShPTCEI2HTtX0nl5If |
| 76-166 | PP2A-B'' | https://benchling.com/s/seq-BY4emeWMOUzOfwxXcfvt?m=slm-tcINDmZcOgsImkBKTvfx |
| 76-151 | MBP | https://benchling.com/s/seq-RP3yWG9oCdfyKJMPJdi7?m=slm-bdiXkPhY9MVs3uDPfc3c |
| 76-146 | Fibronectin | https://benchling.com/s/seq-ZzKvhno6Q626q81EjK1d?m=slm-qlkowg5flGJeA9Vk1DI0 |
| 76-78 | VHL | https://benchling.com/s/seq-e5ttFNgV64jbsyNZQ3PY?m=slm-jnoBlNbfrsKSaq2Ep8E9 |
| 76-159 | BTK (PH) | https://benchling.com/s/seq-cZ9gtCN49KI6B2RBmoyP?m=slm-qIKp2sQWKCw2RvVRqM1X |
| 76-148 | Calpro het 6aa | https://benchling.com/s/seq-Vrfr5UGSgFAO7gMuIx6F?m=slm-UXGody3pDClwTz535g2a |
| 76-77 | GABARAP | https://benchling.com/s/seq-QWrUgcltA0dgKyKnI5fN?m=slm-TRc3OoEsRBgRJn0jMb2z |
| 76-160 | GTF2I | https://benchling.com/s/seq-wbAjXeo3iMuzOgsGCymv?m=slm-HZwpAi3kcEaOrU708mZN |
| 76-85 | OmpF (ec) | https://benchling.com/s/seq-PCc30ARPOrkMJ7TZhvKw?m=slm-pWneGyvjBIBOYNQrZtnA |
| 76-101 | 14-3-3z Monomeric (L12Q, R18E and S58E) | https://benchling.com/s/seq-rYnWfjvOKjhc9ak4KfD7?m=slm-P6aLDJAlg1nAvJWgnMIF |
| 73-165 | PCNA-AF | https://benchling.com/s/seq-NyXUSZ8TBJmIT3RRKy2Q?m=slm-QIapA5oxPlJqO1kuZv6V |
| 73-175 | TEV | https://benchling.com/s/seq-gtv7gSu5b2FGnMJJbnlD?m=slm-4fv9YIXApwoWPXu8mhkd |
| 76-172 | FnbpA | https://benchling.com/s/seq-IheXG0iYJlZxW30gL3Y0?m=slm-XvsmEjdyVf3S8µLrjM0E |
| 76-161 | IBTK | https://benchling.com/s/seq-duQlBjHihWV9fAG3GRnc?m=slm-ELq4y4tZ8zUtm72XAPUG |
| 76-162 | PAL | https://benchling.com/s/seq-DARw5POmwzvIJ9uWmeGO?m=slm-sDNz3BY9RLpDtN6tyjXy |
| 73-171 | PIK3CB (3-115) | https://benchling.com/s/seq-M7YPl5R5qVffKYHqggRo?m=slm-kcbPs3f0DZT63IINpyc4 |
| 76-173 | SortaseA (25-206) | https://benchling.com/s/seq-Ru7jLju6GyoLCTdaSbsn?m=slm-XLo9CRmzkKxn4yGdNwQk |
| 76-163 | PilQ (468-549) | https://benchling.com/s/seq-Ru7jLju6GyoLCTdaSbsn?m=slm-XLo9CRmzkKxn4yGdNwQk |
| 76-164 | DltD (ef) | https://benchling.com/s/seq-i5tvCTZlzMwFyoDZtVIu?m=slm-MdE8CotxvlvH2bozLaZ2 |
| 76-104 | TNFR2 (33-205) | https://benchling.com/s/seq-IvN8nTYDCjqopYD1VWkE?m=slm-r3CzTPlvJvIKRsO8WfZb |
| 73-166 | CDKN1A | https://benchling.com/s/seq-istCEzGSqnP9JetiQCuN?m=slm-RISOjKarN6Ff2X2KVkaH |
| 76-81 | SH3BP5 | https://benchling.com/s/seq-I1mxq7TOjVqvnYaXbq69?m=slm-S9kmHpbmaAehQ1Cif04r |
| 76-82 | OTUBI (1-102) | https://benchling.com/s/seq-yBlNWiwMPwIP08fY6rTL?m=slm-Ek7d1P6R9a8umhclHnGY |
| 76-165 | DltD (strep) | https://benchling.com/s/seq-LbBpShA9x0Ode6p8PHAA?m=slm-KY2U9M0jhl6wfoGTp8OG |
| 76-155 | PINK | https://benchling.com/s/seq-A2zauOogFx5uYT7qmcT0?m=slm-1lbMTuobO1B89f2kHsSu |
| 76-83 | RB1 | https://benchling.com/s/seq-Xpy5OSXeQ51g6hi8WiGl?m=slm-28UkfqgISqw1C5vZBdBU |
| 76-86 | SasG (1060-1140) | https://benchling.com/s/seq-w5s9LZ1NQkLdhYruHFzm?m=slm-13LcEXHVKY7lNMyzHFw8 |
| 76-171 | SasG (930-1060) | https://benchling.com/s/seq-eAt4uQ54H1lpbx756DmO?m=slm-OMUDgLmYp8pQsQsHP17a |
| 73-179 | FKBP | https://benchling.com/s/seq-EjiHC9vGIVQLxz16BYzG?m=slm-FlerSN3uC2FpJhXgIewN |
| 73-180 | BRD4 | https://benchling.com/s/seq-bHjj1lDx3IogSnCjPck3?m=slm-qsp748T6DRU2YsMWrx3g |
| 76-87 | IsdB (125-269) | https://benchling.com/s/seq-BX4NIEpvXhR1IIAns94D?m=slm-4mQ9Te3U6StSI7HIN3Ny |
| 76-102 | HEWL | https://benchling.com/s/seq-vfQnr94eWLSHUc62TMCm?m=slm-xUA1Kr8fRnMd1cavpdlO |
| 73-164 | HSPB1 (90-171) | https://benchling.com/s/seq-GUv6WsRBpj4gz6bKvn3W?m=slm-iXkRj7y0gZgfL1PG29pI |
| 73-181 | KRAS G12C | https://benchling.com/s/seq-fSgcR78NvYN0yJjJ3DmN?m=slm-HB1w62dQtsObif5QH6NI |
| | Mst2 linker | |
| 76-103 | LMO2 (28-150) | https://benchling.com/s/seq-isKCMFIWJoigsZl8JsEJ?m=slm-5kBZFRPsk6xqvvslyub2 |
| 76-178 | HIF1a | https://benchling.com/s/seq-yEjxiXRm7O8Cvwc7ygFu?m=slm-jY0kKsGMQjCrXn1AumaJ |
| 76-158 | MXD1 | https://benchling.com/s/seq-SXFtHNpq5CNf3Y1GqBzz?m=slm-yAz9909ZIqdM8yxl3RbS |
| 76-84 | OmpA (ec, 84-224) | https://benchling.com/s/seq-gEac4hfWY7YSGwDZ2t1K?m=slm-FHCEGTHZ6TtM3BR38qLT |
| 73-168 | CD3z | https://benchling.com/s/seq-zhEuJgddfbTMl3LhVoeE?m=slm-xkjuBuJDo1ZVinKxxInh |
| 73-172 | PIK3CA (1-105) | https://benchling.com/s/seq-uarUD9PQsIuP4MamVl5m?m=slm-N3NhFTD6CqskaShokbqy |
| 73-176 | UEV | https://benchling.com/s/seq-1x3VYe3Rn6Cl2xHsf2My?m=slm-uBwK3p8e3BRgYpSLxbYu |
| 73-178 | FRB | https://benchling.com/s/seq-P2jkD6cGA69xaoxjgGiV?m=slm-cpQ3pvLLLgDsEorNynV0 |
| 73-182 | PD-1 | https://benchling.com/s/seq-yQzKo1Jh4I6HZq5rVG2V?m=slm-14baz2qEaANS3WMAKP5E |
| 73-187 | EIF4A | https://benchling.com/s/seq-zvEJ48QR8qUPsd40SF8M?m=slm-FVmn8NH5S01MEdwUVnYh |
| 76-71 | SasA | https://benchling.com/s/seq-Gx9NNJ91hTJzPumE9pZ7?m=slm-L1RumENXQNADsFgxn9T5 |
| 76-79 | LC3B | https://benchling.com/s/seq-rlt5IzSBDshWhl9Rb2fu?m=slm-SµL56BpmJVP1Nz5nKQcW |
| 76-156 | ALKBH5 | https://benchling.com/s/seq-s6c48MGHowq9jMME85ff?m=slm-23tUDuPOmB1iHJnMrSFO |
| 76-177 | HIF1b (1-474) | https://benchling.com/s/seq-4qbTb9Y8HldXECOCqQxH?m=slm-bH3t5YRzVtbYIlKm7XGF |
| 76-69 | IFNG (24-156) | https://benchling.com/s/seq-Rxb7OuSxhwC1KxJykaPR?m=slm-qgjWG0Jvg76X7yQpLdCb |
| 77-54 | MST2 (313-437) | https://benchling.com/s/seq-3QXRIY0pRUN97KNELCgH?m=slm-eJW6pYZYT851VaVwyHhC |

| | | |
|---|---|---|
| 76-144 | p53 (1-293) | https://benchling.com/s/seq-Fu83Td8HuPIQwC2nRt64?m=slm-SWkrs4aEcLewx5C6B1oC |
| 73-183 | PD-L1 | https://benchling.com/s/seq-RvhD3N2PU1hj77T4fwoo?m=slm-FWNH2UNTgw5H7MTcHQFl |
| 77-60 | Bax peptide | https://benchling.com/s/seq-aINRsklXSYSJcJv8aURp?m=slm-zKz2j49bk5r1v2mVWmMc |
| 76-68 | p53 AD (1-61) | https://benchling.com/s/seq-oboHu3wQGGrOxVZXR8mT?m=slm-CL2SXiwFRno7endZDLq8 |
| 73-169 | CD3d | https://benchling.com/s/seq-WcQwZtijlEoLB1KU2yBt?m=slm-XAGmL1pF0bAJlqi2L7qA |
| 76-147 | Myc (long) | https://benchling.com/s/seq-KWoSoJbUvTLOHeE1TIut?m=slm-2LNV6oSuS8u6YcLQCfKx |
| 76-153 | CRBN | https://benchling.com/s/seq-BAPh6kwfucKjYDmmPXFp?m=slm-i01HyK2nieJmwBXY5n6z |
| 76-100 | MAX | https://benchling.com/s/seq-GKeMoRGv4qd77TtfORFX?m=slm-i9MBG0ae66SnGxHPZgvF |
| 73-170 | PPP1R11 | https://benchling.com/s/seq-XHGhEcm7nvFEbuDNURBH?m=slm-SRI1vBi0GmF6ybMdXEbZ |
| 76-76 | Alpha amylase | https://benchling.com/s/seq-SBXDewfrSaxr7UHdZPKi?m=slm-xPkWa4IgjbTC38snLtEc |
| 73-174 | PMP22 | https://benchling.com/s/seq-AFHDnBiEoYvLILOGoita?m=slm-Bi5fpdTxyr84R5wT0uow |
| 77-53 | BAX | https://benchling.com/s/seq-SxhwkmdaNisoVdvFR6IP?m=slm-pbu3gAKdt55C3yvTXplM |
| 73-186 | Keap1 | https://benchling.com/s/seq-c4pJiEFPzHNjGtroj130?m=slm-VCVLym1r9ip2BtvQwbWY |
| 73-163 | HSPB1 | https://benchling.com/s/seq-sOzF6JEREPkrjU54uVFv?m=slm-vhYiNPEDQpHkYFGytVSr |
| 73-177 | HMGB1 | https://benchling.com/s/seq-Hme2Y2BCw59yBZQevApH?m=slm-00Z5hzIuKqBGFduBmlC0 |

**pET for protein purification**

| Description | Plasmid map |
|---|---|
| KRAS G12D 1-169 pET (HIS tag) | https://benchling.com/s/seq-1SWQgC0RLPELFTGwTnCV?m=slm-bIm9FfhwnFWabV0jgDcx |
| RAF (RBD) pET (His tag) | https://benchling.com/s/seq-D3VuadJQQRyylcSsMwAN?m=slm-DpvKaocf8aFnOcZS3uWv |
| Mdm2 (1-188) pET (His tag) | https://benchling.com/s/seq-oAWBkEqTG3gFhdht0XgG?m=slm-d7xeIGRU5PZIm4ooKwTr |
| IFNG pET (His tag) | https://benchling.com/s/seq-bJuJ9eGchlO8XJPvCn7f?m=slm-sF68NxIg9TzwSyTlADkm |
| 3xFLAG-GST-Affibody binder | https://benchling.com/s/seq-2h4RsRWrou5PCfYx4A9S?m=slm-PbduezfrnBQcWVeCQ025 |

**CMV Plasmids**

| Description | Plasmid Map |
|---|---|
| Target split NanoLuc | https://benchling.com/s/seq-irhIGNg6FDqyVXUYSpRO?m=slm-tRQlv6Im796pYvLSLXDB |
| Binder split NanoLuc | https://benchling.com/s/seq-OoyonZHpjwnrXIcd9qDI?m=slm-ewQylhZqd9FbB6Axnbs6 |
| Binder-LIR fusion | https://benchling.com/s/seq-oEP0ngFLZMXyyvU9NUIh?m=slm-yHYpECMuMIUp7G2Y5EIP |
| Binder only | https://benchling.com/s/seq-a42TroNswKf69IzCp9oA?m=slm-KgcTScaWOEOcWOHPsAfh |
| Mdm2-mCherry | https://benchling.com/s/seq-apmlnageLFs9kyBLSZqi?m=slm-FSLivJ1lYDoy6UhLVwMU |
| Binder-GFP | https://benchling.com/s/seq-jsSL7OyGOMNORbAGnD7U?m=slm-gwZEdWQQKi2Z8uw1Bjdn |

**Supplementary Table 2: Primers**

| ID | Description | **Sequence** ($X_{12}$ indicates sequence specific to target) |
|---|---|---|
| VC-525 | Forward primer for qPCR | ggttcagcaaggtgatgctt |
| VC-526 | Reverse primer for qPCR | accgcctcacctctgtttta |
| MS-783 | Forward primer for Affibody Library | CATCATCATGCTAGCGTAGATAATAAATTTWMYAAAGAAKKGNNYDMYGCADHYNNYSARATTNNYNNYCTGCCGAACTTAAACSNSNNYCAGVDRNNYGCCTTCVTHNNYAGCCTGNNYRAYGACCCGTCCCAAAGCGCTAATTTGCTGGCGGAG |
| MS-618 | Reverse primer for Affibody Library | CATCATgctagcGCCAGAGCCAGAGCCACCCTTGTAGAC |
| MS-624 | Forward primer for Affitin Library | CATCATgctagcTCGTGCGGAACGTGAGAAGAAACTGTAATGGAG |
| MS-799 | Reverse primer for Affitin Library | CATCATGCTAGCATATCTAATAATTCTTTCGGTGCATCTTTTTCRNNAACRNNACCRNNACCRNNTTTACCATTATCATCATARNNAAARNNAACRNNTTTACCRNNACGRNNAACRNNRNNAATTTTAGAAGTATCAACTTCTTTTTCTTCACCRNNRNNRNNAAACTTGACTTTAACTGATCCGCTCGAGCCAGAG |
| | General Forward for insert for cloning scaffold into phage or Lux-N | tggctctggctctggctcgagcXXXXXXXXXXXX |
| | General Reverse for insert for cloning scaffold into phage | cgattgagggagcatgttgaaaatctccattaXXXXXXXXXXXX |
| | General Reverse for insert for cloning scaffold into Lux-N | gctgaggagtgccgttaattaagtttaXXXXXXXXXXXX |
| BR-305 | Forward for vector for cloning scaffold into phage | tggagattttcaacatgctccctcaatcg |
| MS-40 | Forward for vector for cloning scaffold into Lux-N | taaacttaattaacggcactcctcagcaaatataatgacc |
| KJ-14 | Reverse for vector for cloning scaffold into phage or Lux-N | CTCTGGCTCTGGCTCGAGC |
| | General Forward for insert for cloning target into +AP | ggatccctcgaaaggaggaaaaaaaaATGXXXXXXXXXXXX |
| | General Forward for insert for cloning target into Lux-C | gtctgacataaatgaccgctATGXXXXXXXXXXXX |
| | General Reverse for insert for cloning target into +AP/Lux-C | ctttaccgcttccacccgacgtXXXXXXXXXXXX |
| KJ-19 | Forward for vector for cloning target into +AP or Lux-C | ACGTCGGGTGGAAGCGGT |
| MS-46 | Reverse for vector for cloning target into +AP | ttttttttcctcctttcgagGGATCCtaggtag |
| BR-116 | Reverse for vector for cloning scaffold into Lux-C | ACAACTCAAGTCTGACATAAATGACCGCT |
| MS-659 | Forward for insert for subcloning library variants into Lux-N | cacgtctacaagGGTGGCTCTG |
| JD-856 | Reverse for insert for subcloning library variants into Lux-N | tcaatcggttgaatgtcgccctt |
| MS-660 | Forward for vector for subcloning library variants into Lux-N | aatcggttgaatgtcgcccttacttaattaacggcactcctcagc |
| MS-661 | Reverse for vector for subcloning library variants into Lux-N | cacgtctacaagGGTGGCTCT |

| | | |
|---|---|---|
| MS-829 | Forward primer for extending 6 aa linker to 20 aa | cgggtggaagcggtGGTAGCGGGTCGTCGGGAAGTGGTGGTACGTCCGGGTCCGGTaaagcatttatgcaagttgtcgaggc |
| MS-298 | Reverse primer for extending 6 aa linker to 20 aa or 40 aa | CGTCGGGTGGAAGCGGT |
| MS-830 | Forward primer for extending 6 aa linker to 40 aa | cgggtggaagcggtGGCTCGGGCTCATCAGGTTCTGGTGGGACTAGTGGGTCCGGAGGATCAGGATCGGGGACCGGAAGTGGCTCCTCTGGTTCTGGCGGAACTTCCGGTTCAGGAaaagcatttatgcaagttgtcgaggc |
| MS-831 | Forward primer for extending 6 aa linker to 60 aa | CAGGCACAGGGACCGGAAGTAGCGGATCGGGTGGCACAAGTGGGTCTGGCGGCTCTGGATCAGGCACCGGCAGTGGGTCTTCGGGGAGTGGAGGTACCTCAGGCTCAGGAaaagcatttatgcaagttgtcgaggc |
| MS-832 | Reverse primer for extending 6 aa linker to 60 aa | CGTCGGGTGGAAGCGGTGGTTCAGGGTCCTCAGGTTCGGGTGGAACTAGTGGCTCTGGTGGGTCGGGGACAGGCACAGGGACC |
| MS-1156 | Reverse primer for Amplicon-EZ NGS for all barcodes (purple is adaptor, green is priming region) | gactggagttcagacgtgtgctcttccgatctgggcgacattcaaccgattgaggg |
| MS-1144 | Forward primer for Amplicon-EZ NGS Barcode 1 (red is adaptor, blue is barcode, green is priming region) | acactctttccctacacgacgctcttccgatctAAGGTTgcgcgtagggcacgtctacaag |
| MS-1145 | Forward primer for Amplicon-EZ NGS Barcode 2 | acactctttccctacacgacgctcttccgatctTAGGATgcgcgtagggcacgtctacaag |
| MS-1146 | Forward primer for Amplicon-EZ NGS Barcode 3 | acactctttccctacacgacgctcttccgatctTGAGATgcgcgtagggcacgtctacaag |
| MS-1147 | Forward primer for Amplicon-EZ NGS Barcode 4 | acactctttccctacacgacgctcttccgatctAGTGATgcgcgtagggcacgtctacaag |
| MS-1148 | Forward primer for Amplicon-EZ NGS Barcode 5 | acactctttccctacacgacgctcttccgatctAAGTGTgcgcgtagggcacgtctacaag |
| MS-1149 | Forward primer for Amplicon-EZ NGS Barcode 6 | acactctttccctacacgacgctcttccgatctAAGTTGgcgcgtagggcacgtctacaag |
| MS-1150 | Forward primer for Amplicon-EZ NGS Barcode 7 | acactctttccctacacgacgctcttccgatctAGATGTgcgcgtagggcacgtctacaag |
| MS-1151 | Forward primer for Amplicon-EZ NGS Barcode 8 | acactctttccctacacgacgctcttccgatctAGTAGTgcgcgtagggcacgtctacaag |
| MS-1152 | Forward primer for Amplicon-EZ NGS Barcode 9 | acactctttccctacacgacgctcttccgatctGAATTGgcgcgtagggcacgtctacaag |
| MS-1153 | Forward primer for Amplicon-EZ NGS Barcode 10 | acactctttccctacacgacgctcttccgatctGATATGgcgcgtagggcacgtctacaag |
| MS-1154 | Forward primer for Amplicon-EZ NGS Barcode 11 | acactctttccctacacgacgctcttccgatctGTAATGgcgcgtagggcacgtctacaag |
| MS-1155 | Forward primer for Amplicon-EZ NGS Barcode 12 | acactctttccctacacgacgctcttccgatctGTATAGgcgcgtagggcacgtctacaag |
| MS-1346 | Forward primer for Amplicon-EZ NGS Barcode 13 | acactctttccctacacgacgctcttccgatctAACCTTgcgcgtagggcacgtctacaag |
| MS-1347 | Forward primer for Amplicon-EZ NGS Barcode 14 | acactctttccctacacgacgctcttccgatctTACCATgcgcgtagggcacgtctacaag |

| | | |
|---|---|---|
| MS-1348 | Forward primer for Amplicon-EZ NGS Barcode 15 | acactctttccctacacgacgctcttccgatctTCACATgcgcgtagggcacgtctacaag |
| MS-1349 | Forward primer for Amplicon-EZ NGS Barcode 16 | acactctttccctacacgacgctcttccgatctACTCATgcgcgtagggcacgtctacaag |
| MS-1350 | Forward primer for Amplicon-EZ NGS Barcode 17 | acactctttccctacacgacgctcttccgatctAACTCTgcgcgtagggcacgtctacaag |
| MS-1351 | Forward primer for Amplicon-EZ NGS Barcode 18 | acactctttccctacacgacgctcttccgatctAACTTCgcgcgtagggcacgtctacaag |
| MS-1352 | Forward primer for Amplicon-EZ NGS Barcode 19 | acactctttccctacacgacgctcttccgatctACATCTgcgcgtagggcacgtctacaag |
| MS-1353 | Forward primer for Amplicon-EZ NGS Barcode 20 | acactctttccctacacgacgctcttccgatctACTACTgcgcgtagggcacgtctacaag |
| MS-1354 | Forward primer for Amplicon-EZ NGS Barcode 21 | acactctttccctacacgacgctcttccgatctCAATTCgcgcgtagggcacgtctacaag |
| MS-1355 | Forward primer for Amplicon-EZ NGS Barcode 22 | acactctttccctacacgacgctcttccgatctCATATCgcgcgtagggcacgtctacaag |
| MS-1356 | Forward primer for Amplicon-EZ NGS Barcode 23 | acactctttccctacacgacgctcttccgatctCTAATCgcgcgtagggcacgtctacaag |
| MS-1357 | Forward primer for Amplicon-EZ NGS Barcode 24 | acactctttccctacacgacgctcttccgatctCTATACgcgcgtagggcacgtctacaag |

**Supplementary Table 3: Sequence and details of target proteins in this study.**

| Target | AA Sequence | AlphaFold3 Structure | Other Notes |
|---|---|---|---|
| Very high (pLDDT > 90) | Confident (90 > pLDDT > 70) | Low (70 > pLDDT > 50) | Very low (pLDDT < 50) |
| No Fusion | M | | |
| KRAS (G12D) | MTEYKLVVVGADGVGKSALTIQLIQNHFVDEYDPTIEDSYRKQVVIDGETCLLDILDTAGQEEYSAMRDQYMRTGEGFLCVFAINNTKSFEDIHHYREQIKRVKDSEDVPMVLVGNKCDLPSRTVDTKQAQDLARSYGIPFIETSAKTRQGVDDAFYTLVREIRKHKEKMSKDGKKKKKKSKTKCVIM | 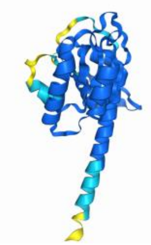 | Uniprot: P01116. Human, cytoplasm, GTPase, oncogene, full length isoform 4b G12D mutant. |
| ZB | MASEQLEKKLQALEKKLAQLEWKNQALEKKLAQ | 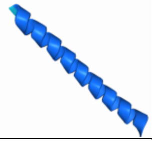 | Synthetic construct; PMID: 15631464 |
| HPV-pE7 | MHGDTPTLHEYMLDLQPETTDLYCYEQLNDSSEEEDEIDGPAGQAEPDRAHYNIVTFCCKCDSTLRLCVQSTHVDIRTLEDLLMGTLGIVCPICSQKP | 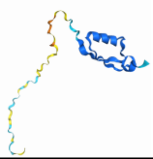 | Uniprot: P03129. Human Papillomavirus (HPV), involved in controlling cell cycle and viral genome replication, partially structured, full length. |

| | | | |
|---|---|---|---|
| p65 (17-293) | MSGPYVEIIEQPKQRGMRFRYKCEG<br>RSAGSIPGERSTDTTKTHPTIKINGYT<br>GPGTVRISLVTKDPPHRPHPHELVGK<br>DCRDGFYEAELCPDRCIHSFQNLGIQ<br>CVKKRDLEQAISQRIQTNNNPFQVPI<br>EEQRGDYDLNAVRLCFQVTVRDPSG<br>RPLRLPPVLSHPIFDNRAPNTAELKIC<br>RVNRNSGSCLGGDEIFLLCDKVQKE<br>DIEVYFTGPGWEARGSFSQADVHRQ<br>VAIVFRTPPYADPSLQAPVRVSMQLR<br>RPSDRELSEPMEFQYLPDTD | 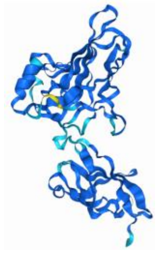 | Uniprot: Q04206. Human, aka NF-κβ, nucleus, transcription factor, structured region |
| FGFR1c domain D2 (153-251) | MAPYWTNTEKMEKRLHAVPAANTVK<br>FRCPAGGNPMPTMRWLKNGKEFKQ<br>EHRIGGYKVRNQHWSLIMESVVPSD<br>KGNYTCVVENEYGSINHTYHLDVVER | 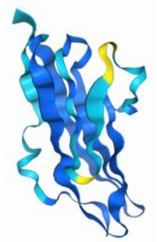 | Uniprot: P21802. Human, single-pass type 1 membrane protein, extracellular domain |
| TCIM | MKAKRSHQAVIMSTSLRVSPSIHGYH<br>FDTASRKKAVGNIFENTDQESLERLF<br>RNSGDKKAEERAKIIFAIDQDVEEKTR<br>ALMALKKRTKDKLFQFLKLRKYSIKVH | 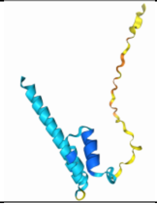 | Uniprot: Q9NR00. Human, nucleus/speckle/nucleolus, full length, disordered |
| TRIM21 (277-475) | MGLKKMLRTCAVHITLDPDTANPWLI<br>LSEDRRQVRLGDTQQSIPGNEERFD<br>SYPMVLGAQHFHSGKHYWEVDVTG<br>KEAWDLGVCRDSVRRKGHFLLSSKS<br>GFWTIWLWNKQKYEAGTYPQTPLHL<br>QVPPCQVGIFLDYEAGMVSFYNITDH<br>GSLIYSFSECAFTGPLRPFFSPGFND<br>GGKNTAPLTLCPLNIGSQGSTDY | 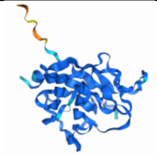 | Uniprot: P19474. Human, cytoplasm/nucleus, E3 ligase, structured domain, Fc binding region |
| PDE6D (1-150) | MSAKDERAREILRGFKLNWMNLRDA<br>ETGKILWQGTEDLSVPGVEHEARVP<br>KKILKCKAVSRELNFSSTEQMEKFRL<br>EQKVYFKGQCLEEWFFEFGFVIPNST<br>NTWQSLIEAAPESQMMPASVLTGNVI<br>IETKFFDDDLLVSTSRVRLFYV | 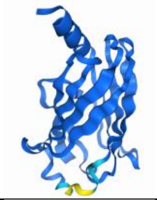 | Uniprot: O43924. Human, cytoplasm/membrane, binds/shuttles prenylated proteins including RAS family proteins; full length |
| RAF (52-131) | MSKTSNTIRVFLPNKQRTVVNVRNG<br>MSLHDCLMKALKVRGLQPECCAVFR<br>LLHEHKGKKARLDWNTDAASLIGEEL<br>QVDFL | 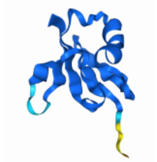 | Uniprot: P04049. Human, cytoplasm, proto oncogene, RAS binding domain |
| NRAS | MTEYKLVVVGAGGVGKSALTIQLIQN<br>HFVDEYDPTIEDSYRKQVVIDGETCL<br>LDILDTAGQEEYSAMRDQYMRTGEG<br>FLCVFAINNSKSFADINLYREQIKRVK<br>DSDDVPMVLVGNKCDLPTRTVDTKQ<br>AHELAKSYGIPFIETSAKTRQGVEDAF<br>YTLVREIRQYRMKKLNSSDDGTQGC<br>MGLPCVVM | 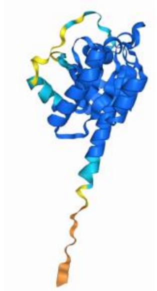 | Uniprot: P01111. Human, cytoplasm/cell membrane/Golgi apparatus membrane, proto oncogene, full lenth |

| | | | |
|---|---|---|---|
| Beclin (125-215) | MIMSGQTDVDHPLCEECTDTLLDQLDTQLNVTENECQNYKRCLEILEQMNEDDSEQLQMELKELALEEERLIQELEDVEKNRKIVAENLEKV | 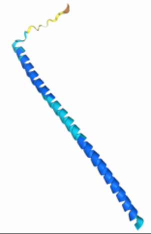 | Uniprot: Q14457. Human, cytoplasm, central role in autophagy and the PI3K complex, region that interacts with BCL2 family proteins. |
| Mdm2 (1-188) | MCNTNMSVPTDGAVTTSQIPASEQETLVRPKPLLLKLLKSVGAQKDTYTMKEVLFYLGQYIMTKRLYDEKQQHIVYCSNDLLGDLFGVPSFSVKEHRKIYTMIYRNLVVVNQQESSDSGTSVSENRCHLEGGSDQKDLVQELQEEKPSSSHLVSRPSTSSRRRAISETEENSDELSGERQRKRHKSDS | 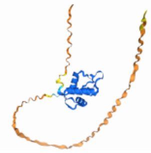 | Uniprot: Q00987. Human, cytoplasm/nucleus, E3 ligase and proto oncogene, region that binds p53. |
| Calprotectin heterodimer with 12aa linker | MTCKMSQLERNIETIINTFHQYSVKLGHPDTLNQGEFKELVRKDLQNFLKKENKNEKVIEHIMEDLDTNADKQLSFEEFIMLMARLTWASHEKMHEGDEGPGHHHKPGLGEGTPGGSGGSGGSGGSMLTELEKALNSIIDVYHKYSLIKGNFHAVYRDDLKKLLETECPQYIRKKGADVWFKELDINTDGAVNFQEFLILVIKMGVAAHKKSHEESHKE | 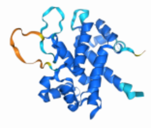 | Uniprot: P05109 and P06702. Human, secreted, calcium and zinc binder involved in inflammation and immune response, full length of each isoform fused with a 12 amino acid linker. |
| HlgA (30-309) | MGHHHHHHAMENKIEDIGQGAEIIKRTQDITSKRLAITQNIQFDFVKDKKYNKDALVVKMQGFISSRTTYSDLKKYPYIKRMIWPFQYNISLKTKDSNVDLINYLPKNKIDSADVSQKLGYNIGGNFQSAPSIGGSGSFNYSKTISYNQKNYVTEVESQNSKGVKWGVKANSFVTPNGQVSAYDQYLFAQDPTGPAARDYFVPDNQLPPLIQSGFNPSFITTLSHERGKGDKSEFEITYGRNMDATYAYVTRHRLAVDRKHDAFKNRNVTVKYEVNWKTHEVKIKSITPK | 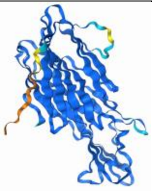 | Uniprot: P0A074. Staphylococcus aureus, secreted hemolytic toxin, folded domain (accidentally included His-tag). |
| Bcl2 | MAHAGRTGYDNREIVMKYIHYKLSQRGYEWDAGDVGAAPPGAAPAPGIFSSQPGHTPHPAASRDPVARTSPLQTPAAPGAAAGPALSPVPPVVHLTLRQAGDDFSRRYRRDFAEMSSQLHLTPFTARGRFATVVEELFRDGVNWGRIVAFFEFGGVMCVESVNREMSPLVDNIALWMTEYLNRHLHTWIQDNGGWDAFVELYGPSMRPLFDFSWLSLK | 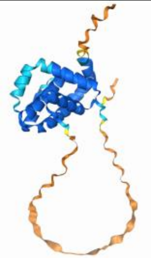 | Uniprot: P10415. Human, cytoplasm/mitochondrion outer membrane/nucleus membrane, suppresses apoptosis, full length. |
| Calpro mon s100a9 | MTCKMSQLERNIETIINTFHQYSVKLGHPDTLNQGEFKELVRKDLQNFLKKENKNEKVIEHIMEDLDTNADKQLSFEEFIMLMARLTWASHEKMHEGDEGPGHHHKPGLGEGTP | 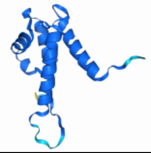 | Uniprot: P06702. Human, secreted, calcium and zinc binder involved in inflammation and immune response, full length. |

| | | | |
|---|---|---|---|
| KIX | MGNIGSLSTIPTAAPPSSTGVRKGWH<br>EHVTQDLRSHLVHKLVQAIFPTPDPA<br>ALKDRRMENLVAYAKKVEGDMYESA<br>NSRDEYYHLLAEKIYKIQKELEEKRRS<br>RL | 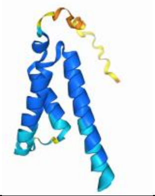 | Uniprot: Q92793. Human, nucleus, transcription factor, CREB Binding Protein (CBP) KIX domain (568-673) |
| HRAS | MTEYKLVVVGAGGVGKSALTIQLIQN<br>HFVDEYDPTIEDSYRKQVVIDGETCL<br>LDILDTAGQEEYSAMRDQYMRTGEG<br>FLCVFAINNTKSFEDIHQYREQIKRVK<br>DSDDVPMVLVGNKCDLAARTVESRQ<br>AQDLARSYGIPYIETSAKTRQGVEDA<br>FYTLVREIRQHKLRKLNPPDESGPGC<br>MSCKCVLS | 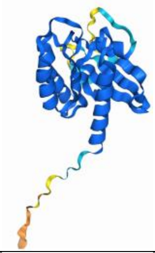 | Uniprot: P01112. Human, cytoplasm, GTPase, oncogene, full length. |
| KRAS wt | MTEYKLVVVGAGGVGKSALTIQLIQN<br>HFVDEYDPTIEDSYRKQVVIDGETCL<br>LDILDTAGQEEYSAMRDQYMRTGEG<br>FLCVFAINNTKSFEDIHHYREQIKRVK<br>DSEDVPMVLVGNKCDLPSRTVDTKQ<br>AQDLARSYGIPFIETSAKTRQGVDDA<br>FYTLVREIRKHKEKMSKDGKKKKKKS<br>KTKCVIM | 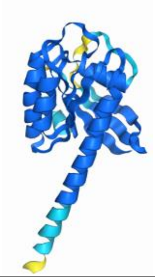 | Uniprot: P01116. Human, cytoplasm, GTPase, oncogene, full length isoform 4b. |
| G3BP (11-139) | MVGREFVRQYYTLLNQAPDMLHRFY<br>GKNSSYVHGGLDSNGKPADAVYGQ<br>KEIHRKVMSQNFTNCHTKIRHVDAHA<br>TLNDGVVVQVMGLLSNNNQALRRFM<br>QTFVLAPEGSVANKFYVHNDIFRYQD<br>EVFG | 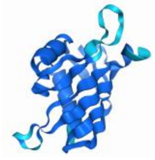 | Uniprot: Q13283. Human, cytoplasm, RAS GTPase activating protein, |
| NIX | MSSHLVEPPPPLHNNNNNCEENEQS<br>LPPPAGLNSSWVELPMNSSNGNDN<br>GNGKNGGLEHVPSSSSIHNGDMEKIL<br>LDAQHESGQSSSRGSSHCDSPSPQ<br>EDGQIMFDVEMHTSRDHSSQSEEEV<br>VEGEKEVEALKKSADWVSDWSSRPE<br>NIPPKEFHFRHPKRSVSLSMRKSGA<br>MKKGGIFSAEFLKVFIPSLFLSHVLAL<br>GLGIYIGKRLSTPSASTY | 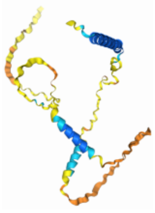 | Uniprot: O60238. Human, nucleus envelope/ER/mitochondrial outer membrane, single-pass membrane protein involved in apoptosis by suppressing BCl2, full length. |
| PP2A-B | MAGAGGGNDIQWCFSQVKGAVDDD<br>VAEADIISTVEFNHSGELLATGDKGG<br>RVVIFQQEQENKIQSHSRGEYNVYST<br>FQSHEPEFDYLKSLEIEEKINKIRWLP<br>QKNAAQFLLSTNDKTIKLWKISERDK<br>RPEGYNLKEEDGRYRDPTTVTTLRV<br>PVFRPMDLMVEASPRRIFANAHTYHI<br>NSISINSDYETYLSADDLRINLWHLEIT<br>DRSFNIVDIKPANMEELTEVITAAEFH<br>PNSCNTFVYSSSKGTIRLCDMRASAL<br>CDRHSKLFEEPEDPSNRSFFSEIISSI<br>SDVKFSHSGRYMMTRDYLSVKIWDL<br>NMENRPVETYQVHEYLRSKLCSLYE<br>NDCIFDKFECCWNGSDSVVMTGSYN<br>NFFRMFDRNTKRDITLEASRENNKPR<br>TVLKPRKVCASGKRKKDEISVDSLDF | 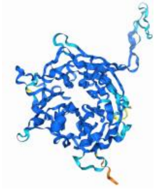 | Uniprot: P63151. Human, cytoplasm, regulatory subunit of a phosphatase, full length. |

| | | | |
|---|---|---|---|
| | NKKILHTAWHPKENIIAVATTNNLYIFQ<br>DKVN | | |
| Calpro<br>mon<br>s100a8 | MLTELEKALNSIIDVYHKYSLIKGNFH<br>AVYRDDLKKLLETECPQYIRKKGADV<br>WFKELDINTDGAVNFQEFLILVIKMGV<br>AAHKKSHEESHKE | 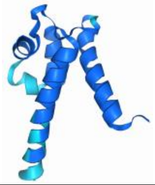 | Uniprot: P05109. Human, secreted, calcium and zinc binder involved in inflammation and immune response, full length. |
| Parkin | MIVFVRFNSSHGFPVEVDSDTSIFQL<br>KEVVAKRQGVPADQLRVIFAGKELRN<br>DWTVQNCDLDQQSIVHIVQRPWRKG<br>QEMNATGGDDPRNAAGGCEREPQS<br>LTRVDLSSSVLPGDSVGLAVILHTDS<br>RKDSPPAGSPAGRSIYNSFYVYCKG<br>PCQRVQPGKLRVQCSTCRQATLTLT<br>QGPSCWDDVLIPNRMSGECQSPHC<br>PGTSAEFFFKCGAHPTSDKETSVALH<br>LIATNSRNITCITCTDVRSPVLVFQCN<br>SRHVICLDCFHLYCVTRLNDRQFVHD<br>PQLGYSLPCVAGCPNSLIKELHHFRIL<br>GEEQYNRYQQYGAEECVLQMGGVL<br>CPRPGCGAGLLPEPDQRKVTCEGG<br>NGLGCGFAFCRECKEAYHEGECSAV<br>FEASGTTTQAYRVDERAAEQARWEA<br>ASKETIKKTTKPCPRCHVPVEKNGGC<br>MHMKCPQPQCRLEWCWNCGCEWN<br>RVCMGDHWFDV | 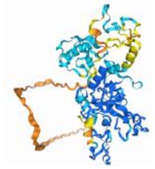 | Uniprot: O60260. Human, cytoplasm, E3 ligase, full length. |
| KRAS<br>(G12V) | MTEYKLVVVGAVGVGKSALTIQLIQN<br>HFVDEYDPTIEDSYRKQVVIDGETCL<br>LDILDTAGQEEYSAMRDQYMRTGEG<br>FLCVFAINNTKSFEDIHHYREQIKRVK<br>DSEDVPMVLVGNKCDLPSRTVDTKQ<br>AQDLARSYGIPFIETSAKTRQGVDDA<br>FYTLVREIRKHKEKMSKDGKKKKKKS<br>KTKCVIM | 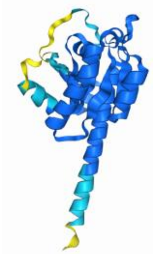 | Uniprot: P01116. Human, cytoplasm, GTPase, oncogene, full length isoform 4b G12V mutant. |
| Sos1<br>(1136-<br>1333) | MSSISLTKGTDEVPVPPPVPPRRRPE<br>SAPAESSPSKIMSKHLDSPPAIPPRQ<br>PTSKAYSPRYSISDRTSISDPPESPPL<br>LPPREPVRTPDVFSSSPLHLQPPPLG<br>KKSDHGNAFFPNSPSPFTPPPPQTP<br>SPHGTRRHLPSPPLTQEVDLHSIAGP<br>PVPPRQSTSQHIPKLPPKTYKREHTH<br>PSMHRDGPPLLENAHSS | 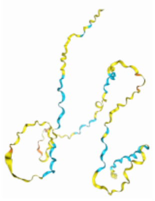 | Uniprot: Q07889. Human, cytoplasm, promotes RAS GTPase nucleotide exchange, disordered domain only. |
| PP2A-B' | MSSSSPPAGAASAAISASEKVDGFTR<br>KSVRKAQRQKRSQGSSQFRSQGSQ<br>AELHPLPQLKDATSNEQQELFCQKL<br>QQCCILFDFMDSVSDLKSKEIKRATL<br>NELVEYVSTNRGVIVESAYSDIVKMIS<br>ANIFRTLPPSDNPDFDPEEDEPTLEA<br>SWPHIQLVYEFFLRFLESPDFQPSIAK<br>RYIDQKFVQQLLELFDSEDPRERDFL<br>KTVLHRIYGKFLGLRAFIRKQINNIFLR<br>FIYETEHFNGVAELLEILGSIINGFALP<br>LKAEHKQFLMKVLIPMHTAKGLALFH<br>AQLAYCVVQFLEKDTTLTEPVIRGLLK | 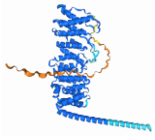 | Uniprot: Q15172. Human, cytoplasm, regulatory subunit of a phosphatase, full length. |

| | | | |
|---|---|---|---|
| | FWPKTCSQKEVMFLGEIEEILDVIEPT<br>QFKKIEEPLFKQISKCVSSSHFQVAE<br>RALYFWNNEYILSLIEENIDKILPIMFA<br>SLYKISKEHWNPTIVALVYNVLKTLME<br>MNGKLFDDLTSSYKAERQREKKKEL<br>EREELWKKLEELKLKKALEKQNSAYN<br>MHSILSNTSAE | | |
| mCherry | MVSKGEEDNMAIIKEFMRFKVHMEG<br>SVNGHEFEIEGEGEGRPYEGTQTSK<br>LKVTKGGPLPFAWDILSPQFMYGSKA<br>YVKHPADIPDYLKLSFPEGFKWERVM<br>NFEDGGVVTVTQDSSLQDGEFIYKVK<br>LRGTNFPSDGPVMQKKTMGWEASS<br>ERMYPEDGALKGEIKQRLKLKDGGH<br>YDAEVKTTYKAKKPVQLPGAYNVNIK<br>LDITSHNEDYTIVEQYERAEGRHSTG<br>GMDELYK | 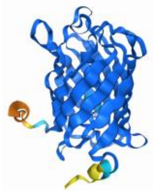 | Synthetic evolved protein from DsRed of Discosoma sea anemones. Red fluorescent protein full length. |
| Myc (DBD) | MNVKRRTHNVLERQRRNELKRSFFA<br>LRDQIPELENNEKAPKVVILKKATAYIL<br>SVQAEEQKLISEEDLLRKRREQLKHK<br>LEQL | 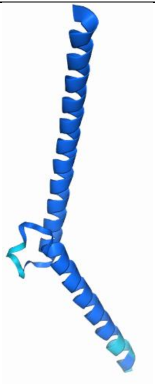 | Uniprot: P01106. Human, nucleus, proto oncogene transcription factor, DNA binding domain (368-451). |
| PP2A-B'' (622-1150) | MQILQETLTTSSQANLSVCRSPVGDK<br>AKDTTSAVLIQQTPEVIKIQNKPEKKP<br>GTPLPPPATSPSSPRPLSPVPHVNNV<br>VNAPLSINIPRFYFPEGLPDTCSNHE<br>QTLSRIETAFMDIEEQKADIYEMGKIA<br>KVCGCPLYWKAPMFRAAGGEKTGF<br>VTAQSFIAMWRKLLNNHHDDASKFIC<br>LLAKPNCSSLEQEDFIPLLQDVVDTH<br>PGLTFLKDAPEFHSRYITTVIQRIFYTV<br>NRSWSGKITSTEIRKSNFLQTLALLEE<br>EEDINQITDYFSYEHFYVIYCKFWELD<br>TDHDLYISQADLSRYNDQASSSRIIER<br>IFSGAVTRGKTIQKEGRMSYADFVWF<br>LISEEDKRNPTSIEYWFRCMDVDGD<br>GVLSMYELEYFYEEQCERMEAMGIE<br>PLPFHDLLCQMLDLVKPAVDGKITLR<br>DLKRCRMAHIFYDTFFNLEKYLDHEQ<br>RDPFAVQKDVENDGPEPSDWDRFA<br>AEEYETLVAEESAQAQFQEGFEDYE<br>TDEPASPSEFGNKSNKILSASLPEKC<br>GKLQSVDEE | 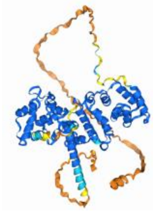 | Uniprot: Q06190. Human, cytoplasm, regulatory subunit of a phosphatase, primarily the structured domain. |

| | | | |
|---|---|---|---|
| MBP (27-396) | MKIEEGKLVIWINGDKGYNGLAEVGKKFEKDTGIKVTVEHPDKLEEKFPQVAATGDGPDIIFWAHDRFGGYAQSGLLAEITPDKAFQDKLYPFTWDAVRYNGKLIAYPIAVEALSLIYNKDLLPNPPKTWEEIPALDKELKAKGKSALMFNLQEPYFTWPLIAADGGYAFKYENGKYDIKDVGVDNAGAKAGLTFLVDLIKNKHMNADTDYSIAEAAFNKGETAMTINGPWAWSNIDTSKVNYGVTVLPTFKGQPSKPFVGVLSAGINAASPNKELAKEFLENYLLTDEGLEAVNKDKPLGAVALKSYEEELAKDPRIAATMENAQKGEIMPNIPQMSAFWYAVRTAVINAASGRQTVDEALKDAQTRITK | 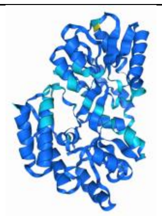 | Uniprot: P0AEX9. Escherichia coli, periplasm, maltose binding protein structured region. |
| Fibronectin (1080-1541) | MGVFTTLQPGSSIPPYNTEVTETTIVITWTPAPRIGFKLGVRPSQGGEAPREVTSDSGSIVVSGLTPGVEYVYTIQVLRDGQERDAPIVNKVVTPLSPPTNLHLEANPDTGVLTVSWERSTTPDITGYRITTTPTNGQQGNSLEEVVHADQSSCTFDNLSPGLEYNVSVYTVKDDKESVPISDTIIPEVPQLTDLSFVDITDSSIGLRWTPLNSSTIIGYRITVVAAGEGIPIFEDFVDSSVGYYTVTGLEPGIDYDISVITLINGGESAPTTLTQQTAVPPPTDLRFTNIGPDTMRVTWAPPPSIDLTNFLVRYSPVKNEEDVAELSISPSDNAVVLTNLLPGTEYVVSVSSVYEQHESTPLRGRQKTGLDSPTGIDFSDITANSFTVHWIAPRATITGYRIRHHPEHFSGRPREDRVPHSRNSITLTNLTPGTEYVVSIVALNGREESPLLIGQQSTVSDV | 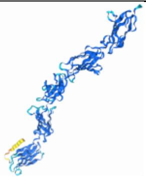 | Uniprot: P02751. Human, secreted, extracellular matrix, binds cell surfaces, various fibronectin type domains. |
| VHL (61-209) | MPVLRSVNSREPSQVIFCNRSPRVVLPVWLNFDGEPQPYPTLPPGTGRRIHSYRGHLWLFRDAGTHDGLLVNQTELFVPSLNVDGQPIFANITLPVYTLKERCLQVVRSLVKPENYRRLDIVRSLYEDLEDHPNVQKDLERLTQERIAHQ | 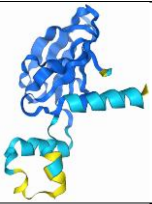 | Uniprot: P40337. Human, cytoplasm/ nucleus, E3 ligase, structured domain. |
| BTK (PH) | MAAVILESIFLKRSQQKKKTSPLNFKKRLFLLTVHKLSYYEYDFERGRRGSKKGSIDVEKITCVETVVPEKNPPPERQIPRRGEESSEMEQISIIERFPYPFQVVYDEGPLYVFSPTEELRKRWIHQLKNVIR | 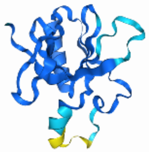 | Uniprot: Q06187. Human, cytoplasm, kinase, PH domain (1-133). |
| Calprotectin het 6aa | MTCKMSQLERNIETIINTFHQYSVKLGHPDTLNQGEFKELVRKDLQNFLKKENKNEKVIEHIMEDLDTNADKQLSFEEFIMLMARLTWASHEKMHEGDEGPGHHHKPGLGEGTPGGSGGSMLTELEKALNSIIDVYHKYSLIKGNFHAVYRDDLKKLLETECPQYIRKKGADVWFKELDINTDGAVNFQEFLILVIKMGVAAHKKSHEESHKE | 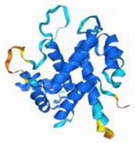 | Uniprot: P05109 and P06702. Human, secreted, calcium and zinc binder involved in inflammation and immune response, full length of each isoform fused with a 6 amino acid linker. |

| | | | |
|---|---|---|---|
| GABARAP | MKFVYKEEHPFEKRRSEGEKIRKKYPDRVPVIVEKAPKARIGDLDKKKYLVPSDLTVGQFYFLIRKRIHLRAEDALFFFVNNVIPPTSATMGQLYQEHHEEDFFLYIAYSDESVYGL | 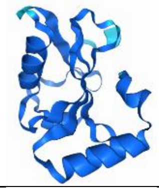 | Uniprot: O95166. Human, cytoplasm, involved in autophagy, full length. |
| GTF2I (1-106) | MAQVAMSTLPVEDEESSESRMVVTFLMSALESMCKELAKSKAEVACIAVYETDVFVVGTERGRAFVNTRKDFQKDFVKYCVEEEEKAAEMHKMKSTTQANRMSVDA | 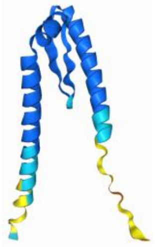 | Uniprot: P78347. Human, nucleus, transcription factor, DNA binding domain. |
| OmpF (233-350) | MAEVVRVQLDVKFDFDKSKVKENSYADIKNLADFMKQYPSTSTTVEGHTDSVGTDAYNQKLSERRANAVRDVLVNEYGVEGGRVNAVGYGESRPVADNATAEGRAINRRVEAEVEAEAK | 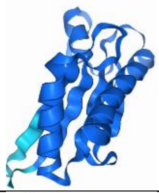 | Uniprot: A6V748. *Psudomonas aeruginosa*, outer membrane protein, extracellular domain. |
| 14-3-3z Monomeric | MDKNELVQKAKQAEQAEDYDDMAACMKSVTEQGAELSNEERNLLSVAYKNVVGARRSEWRVVSSIEQKTEGAEKKQQMAREYREKIETELRDICNDVLSLLEKFLIPNASQAESKVFYLKMKGDYYRYLAEVAAGDDKKGIVDQSQQAYQEAFEISKKEMQPTHPIRLGLALNFSVFYYEILNSPEKACSLAKTAFDEAIAELDTLSEESYKDSTLIMQLLRDNLTLWTSDTQGDEAEAGEGGEN | 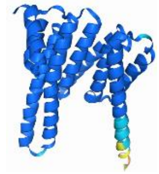 | Uniprot: P63104. Human, cytoplasm, interacts with phosphorylated proteins, full length. Modified to prevent dimerization (L12Q, R18E and S58E). |
| PCNA-AF | MVRTKADSVPGTYRKVVAARAPRKVLGSSTSATNSTSVSSRKAENKYAGGNPVCVRPTPKWQKGIGEFFRLSPKDSEKENQIPEEAGSSGLGKAKRKACPLQPDHTNDEKE | 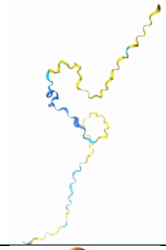 | Uniprot: Q15004. Human, nuclear, regulator of DNA replication, intrinsically disordered, full length. |
| TEV (2038-2274) | MGESLFKGPRDYNPISSTICHLTNESDGHTTSLYGIGFGPFIITNKHLFRRNNGTLLVQSLHGVFKVKNTTTLQQHLIDGRDMIIIRMPKDFPPFPQKLKFREPQREERICLVTTNFQTKSMSSMVSDTSCTFPSSDGIFWKHWIQTKDGQCGSPLVSTRDGFIVGIHSASNFTNTNNYFTSVPKNFMELLTNQEAQQWVSGWRLNADSVLWGGHKVFMVKPEEPFQPVKEATQLMN | 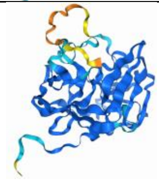 | Uniprot: Q88507. Tobacco etch virus, protease, proteolytic domain. |
| FnbpA (307-461) | MKYYTNLNGSIETFNKADNKFTHVAYVKPINGNKSESVSITGSLTQGSNVSGDSPIVKVYEYQGKETDLPKSVSVNLTDNSKFKDVTSDMQNKLTVQENGNYQLNLEKLDKTYVIHYTGEYSKETDEVNFRTQVSAYPENSYRYYSYYNNHYTLTW | 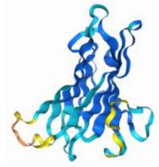 | Uniprot: P14738. *Staphylococcus aureus*, cell surface/secreted, fibronectin binding domain. |

| | | | |
|---|---|---|---|
| IBTK<br>(1159-<br>1339) | MLIDIISSKMIALTTKENNSGMNSMET<br>VLFTPSKVPKPVNAWASSLHSVSSK<br>SFRDFLLEEKKSVTSHSSGDHVKKVS<br>FKGIENSQAPKIVRCSTHGTPGPEGN<br>HISDLPLLDSPNPWLSSSVTAPSMVA<br>PVTFASIVEEELQQEAALIRSREKPLA<br>LIQIEEHAIQDLLVFYEAFGNPEEFVIV<br>ERTPQGPLAVPMWNKHGC | 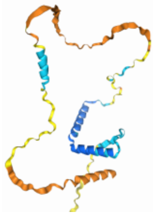 | Uniprot: Q9P2D0. Human, cytoplasm/nucleus, involved in B-cell development through BTK and NF-kb, disordered domain. |
| PAL | MLQMQQLQQNNIVYFDLDKYDIRSDF<br>AQMLDAHANFLRSNPSYKVTVEGHA<br>DERGTPEYNISLGERRANAVKMYLQ<br>GKGVSADQISIVSYGKEKPAVLGHDE<br>AAYSKNRRAVLVY | 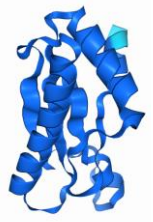 | Uniprot: P0A912. *Escherichia coli*, outer membrane protein involved in cell division, extracellular domain. |
| PIK3CB<br>(3-115) | MFSFIMPPAMADILDIWAVDSQIASD<br>GSIPVDFLLPTGIYIQLEVPREATISYIK<br>QMLWKQVHNYPMFNLLMDIDSYMFA<br>CVNQTAVYEELEDETRRLCDVRPFLP<br>VLKLVTRS | 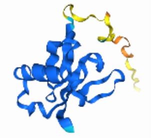 | Uniprot: P42338. Human, cytoplasm/nucleus, kinase, adaptor binding domain (ABD). |
| SortaseA<br>(25-206) | MAKPHIDNYLHDKDKDEKIEQYDKNV<br>KEQASKDKKQQAKPQIPKDKSKVAG<br>YIEIPDADIKEPVYPGPATPEQLNRGV<br>SFAEENESLDDQNISIAGHTFIDRPNY<br>QFTNLKAAKKGSMVYFKVGNETRKY<br>KMTSIRDVKPTDVGVLDEQKGKDKQ<br>LTLITCDDYNEKTGVWEKRKIF | 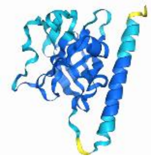 | Uniprot: A0A0H2XJ22. *Staphylococcus aureus*, cell surface/secreted, extracellular domain. |
| PilQ<br>(468-<br>549) | MGVRWGGAYHKGNWSGYGKDGNI<br>GIKDEDGMNCGPIAGSCTFPTTGTSK<br>SPSPFVDLGAKDATSGIGIGFITDNIIL<br>DLQLSA | 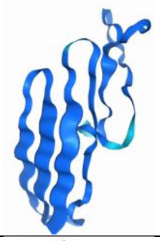 | Uniprot: P34750. *Psudomonas aeruginosa*, outer membrane protein, involved in formation of pili, part of the extracellular domain. |
| DltD (ef)<br>(41-422) | MDISNPETIRRASSSMSVNVLKGDAI<br>KNYALSEKQYIPFFGSSELSRISPFHP<br>SVLAEKYQRNYRPFLLGAPGTQSLS<br>QYMMMRSAGDAMKNKKVVFIISPQW<br>FVKNGVKTDYFNTYYSELQTYDWLF<br>SMKKVTPADRYLARRLLTFSKVKEND<br>TLTAILQTIKKGKLPLPESLNQLRSQW<br>NMLKREDEVFSNIGLKDRQQKIDHES<br>KRLPKQYQETELSILANQIGERETTN<br>NPFGLKNDFYTHRIRAHEPELKQSQK<br>NWDYRFSPEFSDFQLVLDQLAKNHN<br>EVLFIIPPVNEKWSDYTGLSQEMLQG<br>FAKKIKFQLNSQGFNRIADFVNQAGT<br>NYFMEDTIHLGWKGWLAADQQIRPF<br>LEENHITASKYHLDDAFFSKSWQHQI<br>PDKLQLK | 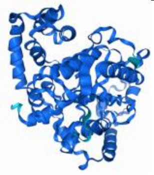 | Uniprot: Q3XZT4. Enterococcus faecium, outer membrane protein involved in cell wall synthesis, extracellular domain. |

| | | | |
|---|---|---|---|
| TNFR2 (33-205) | MAPEPGSTCRLREYYDQTAQMCCS<br>KCSPGQHAKVFCTKTSDTVCDSCED<br>STYTQLWNWVPECLSCGSRCSSDQ<br>VETQACTREQNRICTCRPGWYCALS<br>KQEGCRLCAPLRKCRPGFGVARPGT<br>ETSDVVCKPCAPGTFSNTTSSTDICR<br>PHQICNVVAIPGNASMDAVCTSTSP | 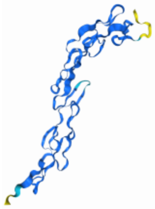 | Uniprot: P20333. Human, cell membrane, involved in regulating apoptosis, extracellular domain. |
| CDKN1A | MSEPAGDVRQNPCGSKACRRLFGP<br>VDSEQLSRDCDALMAGCIQEARERW<br>NFDFVTETPLEGDFAWERVRGLGLP<br>KLYLPTGPRRGRDELGGGRRPGTSP<br>ALLQGTAEEDHVDLSLSCTLVPRSGE<br>QAEGSPGGPGDSQGRKRRQTSMTD<br>FYHSKRRLIFSKRKP | 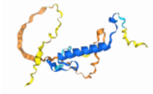 | Uniprot: P38936. Human, cytoplasm/nucleus, kinase inhibitor involved in cell cycle, intrinsically disordered, full length. |
| SH3BP5 | MLNHATQRVMEAEQTKTRSELVHKE<br>TAARYNAAMGRMRQLEKKLKRAINK<br>SKPYFELKAKYYVQLEQLKKTVDDLQ<br>AKLTLAKGEYKMALKNLEMISDEIHER<br>RRSSAMGPRGCGVGAEGSSTSVED<br>L | 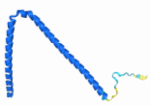 | Uniprot: C9JK30. Human, cytoplasm, guanine nucleotide exchange factor (GEF), full length. |
| OTUB1 (1-102) | MKPSWLSRTEFSKRLLCRTLWCQSG<br>WSSRSYTRSMLKMTTSINRRSRTST<br>KSTRTSARPGLTATVSIGLSDSPTWR<br>HCWMTARSCSGGEADLCRRPAGLL<br>Q | 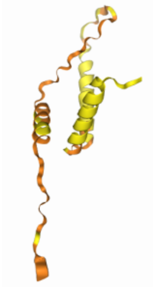 | Uniprot: Q96FW1. Human, cytoplasm, de-ubiquitinase, intrinsically disordered, full length. |
| DltD (strep) (31-390) | MDYYKVKDNSIRYSTSYEKYKSRDIL<br>TSNITPNTLVLMGSSELVATINEDYHP<br>NKIFNYNDFNIMQIGTSYSQNIIQATTL<br>GSIEGSMTKRKVAIVESVQWFEKDGT<br>HQDAFLNKASQEHIFHMLDNGKISKE<br>TKEKLINRIIEITKGNKQQNDIYKKYKS<br>YFIDGKGTIVDKKLLEFENAMYSFKLK<br>QTFYQKHAKSDYPSLGDKTPDYDWQ<br>KMTNQFVEEVKQKTDNNDYAVDNNY<br>YNTYLKDRYASLKDSNKDLSYLESPE<br>YSDMELFLTVAKELGIEVEVIIFPVNG<br>KWNDYTGVSREMREKTYKKIEDVAK<br>SHGATVLNYGNREYDDYFLFDVMHV<br>GVKGWMEVEKELYKFANQAN | 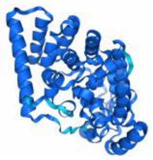 | Uniprot: X8KHP8. Steptococcus sp. CM6, outer membrane protein involved in cell wall synthesis, extracellular domain. |
| PINK (107-353) | MGIPEFEEKQAESRRAVSACQEIQAI<br>FTQKSKPGPDPLDTRRLQGFRLEEYL<br>IGQSIGKGCSAAVYEATMPTLPQNLE<br>VTKSTGLLPGRGPGTSAPGEGQERA<br>PGAPAFPLAIKMMWNISAGSSSEAIL<br>NTMSQELVPASRVALAGEYGAVTYR<br>KSKRGPKQLAPHPNIIRVLRAFTSSVP<br>LLPGALVDYPDVLPSRLHPEGLGHG<br>RTLFLVMKNYPCTLRQYLCVNTPSPR<br>LAAMMLLQLLEGVDHL | 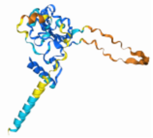 | Uniprot: Q9BXM7. Human, cytoplasm/mitochondrion membrane, kinase, kinase domain, partially disordered. |

| | | | |
|---|---|---|---|
| RB1 (1-103) | MPPKTPRKTAATAAAAAAEPPAPPPPPPPEEDPEQDSGPEDLPLVRLEFEETEEPDFTALCQKLKIPDHVRERAWLTWEKVSSVDGVLVLRSPVFKPWLTMGN | 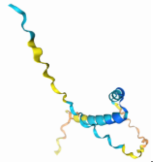 | Uniprot: P06400. Human, nucleus, regulates cell cycle, disordered domain (isoform 4). |
| SasG (1060-1140) | MPKTGTPETKTVEIPFETKREFNPKLQPGEERVKQEGQPGSKTITTPITVNPLTGEKVGEGQPTEEITKQPVDKIVEFGGE | 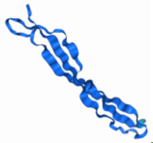 | Uniprot: A0A0H2WXE4. *Staphylococcus aureus*, cell surface, part of the extracellular domain. |
| SasG (930-1060) | MKYGPVKGDSIVEKEEIPFEKERKFNPDLAPGTEKVTREGQKGEKTITTPTLKNPLTGEIISKGESKEEITKDPVNELTEFGGEKIPQGHKDIFDPNLPTDQTEKVPGKPGIKNPDTGKVIEEPVDDVIKHG | 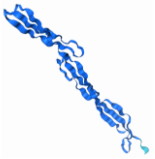 | Uniprot: A0A0H2WXE4. *Staphylococcus aureus*, cell surface, part of the extracellular domain. |
| FKBP (39-145) | MGVQVETISPGDGRTFPKRGQTCVVHYTGMLEDGKKFDSSRDRNKPFKFMLGKQEVIRGWEEGVAQMSVGQRAKLTISPDYAYGATGHPGIIPPHATLVFDVELLKLE | 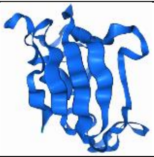 | Uniprot: Q0VDC6. Human, cytoplasm/sarcoplasmic reticulum membrane, cytoplasmic domain (excluding the membrane anchor). |
| BRD4 (349-459) | MKVSEQLKCCSGILKEMFAKKHAAYAWPFYKPVDVEALGLHDYCDIIKHPMDMSTIKSKLEAREYRDAQEFGADVRLMFSNCYKYNPPDHEVVAMARKLQDVFEMRFAKMPD | 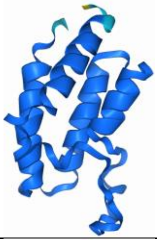 | Uniprot: O60885. Human, nucleus, chromatin reader, bromo 2 domain. |
| IsdB (125-269) | MNQELREAIKNPAIKDKDHSAPNSRPIDFEMKKENGEQQFYHYASSVKPARVIFTDSKPEIELGLQSGQFWRKFEVYEGDKKLPIKLVSYDTVKDYAYIRFSVSNGTKAVKIVSSTHFNNKEEKYDYTLMEFAQPIYNSADKFKT | 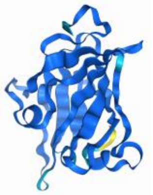 | Uniprot: Q99UX5. *Staphylococcus aureus*, cell surface/secreted, heme receptor, NEAT 1 domain. |
| HEWL (19-147) | MKVFGRCELAAAMKRHGLDNYRGYSLGNWVCAAKFESNFNTQATNRNTDGSTDYGILQINSRWWCNDGRTPGSRNLCNIPCSALLSSDITASVNCAKKIVSDGNGMNAWVAWRNRCKGTDVQAWIRGCRL | 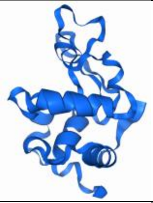 | Uniprot: P00698. *Gallus gallus* (chicken), secreted, lysozyme, structured domain (excluding signal domain). |
| HSPB1 (90-171) | MHTADRWRVSLDVNHFAPDELTVKTKDGVVEITGKHEERQDEHGYISRCFTRKYTLPPGVDPTQVSSSLSPEGTLTVEAPMPK | 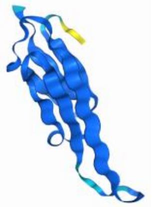 | Uniprot: P04792. Human, cytoplasm/nucleus, molecular chaperone, structured domain only. |

| | | | |
|---|---|---|---|
| KRAS G12C | MTEYKLVVVGACGVGKSALTIQLIQN<br>HFVDEYDPTIEDSYRKQVVIDGETCL<br>LDILDTAGQEEYSAMRDQYMRTGEG<br>FLCVFAINNTKSFEDIHHYREQIKRVK<br>DSEDVPMVLVGNKCDLPSRTVDTKQ<br>AQDLARSYGIPFIETSAKTRQGVDDA<br>FYTLVREIRKHKEKMSKDGKKKKKKS<br>KTKCVIM | 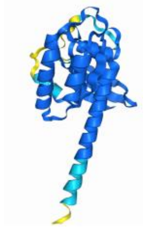 | Uniprot: P01116. Human, cytoplasm, GTPase, oncogene, full length isoform 4b G12C mutant. |
| Mst2 linker (350-380) | MIEHNSTMLESDLGTMVINSEDEEEE<br>DGTMK | 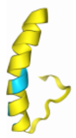 | Uniprot: Q13188. Human, cytoplasm/nucleus, pro-apoptotic kinase, disordered domain that interacts with TRIM21.[1] |
| LMO2 | MSSAIERKSLDPSEEPVDEVLQIPPSL<br>LTCGGCQQNIGDRYFLKAIDQYWHE<br>DCLSCDLCGCRLGEVGRRLYYKLGR<br>KLCRRDYLRLFGQDGLCASCDKRIRA<br>YEMTMRVKDKVYHLECFKCAACQKH<br>FCVGDRYLLINSDIVCEQDIYEWTKIN<br>GMI | 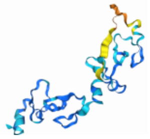 | Uniprot: P25791. Human, nucleus, transcription factor, mostly disordered, full length. |
| HIF-1a (1-350) | MEGAGGANDKKKISSERRKEKSRDA<br>ARSRRSKESEVFYELAHQLPLPHNV<br>SSHLDKASVMRLTISYLRVRKLLDAG<br>DLDIEDDMKAQMNCFYLKALDGFVM<br>VLTDDGDMIYISDNVNKYMGLTQFEL<br>TGHSVFDFTHPCDHEEMREMLTHRN<br>GLVKKGKEQNTQRSFFLRMKCTLTS<br>RGRTMNIKSATWKVLHCTGHIHVYDT<br>NSNQPQCGYKKPPMTCLVLICEPIPH<br>PSNIEIPLDSKTFLSRHSLDMKFSYCD<br>ERITELMGYEPEELLGRSIYEYYHALD<br>SDHLTKTHHDMFTKGQVTTGQYRML<br>AKRGGYVWVETQATVIYNTKNSQPQ<br>CIVCVNYVVSGIIQHDL | 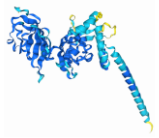 | Uniprot: Q16665. Human, cytoplasm/nucleus, transcriptional regulator, structured domain only. |
| MXD1 | MAAAVRMNIQMLLEAADYLERRERE<br>AEHGYASMLPYNNKDRDALKRRNKS<br>KKNNSSSRSTHNEMEKNRRAHLRLC<br>LEKLKGLVPLGPESSRHTTLSLLTKA<br>KLHIKKLEDCDRKAVHQIDQLQREQR<br>HLKRQLEKLGIERIRMDSIGSTVSSER<br>SDSDREEIDVDVESTDYLTGDLDWS<br>SSSVSDSDERGSMQSLGSDEGYSST<br>SIKRIKLQDSHKACLGL | 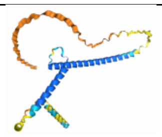 | Uniprot: Q05195. Human, nucleus, transcriptional repressor, intrinsically disordered, full length. |
| OmpA (ec, 84-224) | MADKQEAELRRQMEGTGVEVQRQG<br>DDIKLIMPGNITFATDSANIAPSFYAPL<br>NNLANSFKQYNQNTIEIVGYTDSTGS<br>RQHNMDLSQRRAQSVAGYLTAQGV<br>DGTRLSTRGMGPDQPIASNSTADGR<br>AQNRRVEVNLRPV | 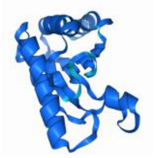 | Uniprot: Q9I5A7. *Psudomonas aeruginosa*, outer membrane protein, extracellular domain. |

| | | | |
|---|---|---|---|
| CD3z | MKWKALFTAAILQAQLPITEAQSFGLLDPKLCYLLDGILFIYGVILTALFLRVKFSRSADAPAYQQGQNQLYNELNLGRREEYDVLDKRRGRDPEMGGKPQRRKNPQEGLYNELQKDKMAEAYSEIGMKGERRRGKGHDGLYQGLSTATKDTYDALHMQALPPR | 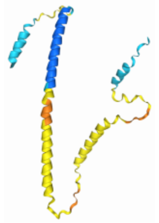 | Uniprot: P20963. Human, cell membrane, T-cell surface receptor, single pass-transmembrane protein, fully disordered, full length. |
| PIK3CA (1-105) | MPPRPSSGELWGIHLMPPRILVECLLPNGMIVTLECLREATLITIKHELFKEARKYPLHQLLQDESSYIFVSVTQEAEREEFFDETRRLCDLRLFQPFLKVIEPV | 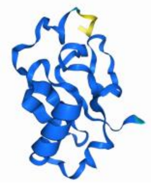 | Uniprot: P42336. Human, cytoplasm, kinase, adaptor binding domain (ABD). |
| UEV (1-145) | MAVSESQLKKMVSKYKYRDLTVRETVNVITLYKDLKPVLDSYVFNDGSSRELMNLTGTIPVPYRGNTYNIPICLWLLDTYPHNPPICFVKPTSSMTIKTGKHVDANGKIYLPYLHEWKHPQSDLLGLIQVMIVVFGDEPPVFSRP | 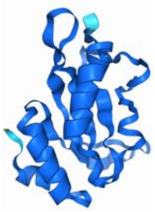 | Uniprot: Q99816. Human, cytoplasm/nucleus, binds ubiquitinated proteins, UEV domain. |
| FRB (2019-2114) | MVAILWHEMWHEGLEEASRLYFGERNVKGMFEVLEPLHAMMERGPQTLKETSFNQAYGRDLMEAQEWCRKYMKSGNVKDLTQAWDLYYHVFRRISKQ | 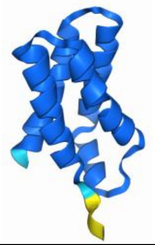 | Uniprot: P42345. Human, cytoplasm, kinase (mTOR), FKPB rapamycin associated domain (FRB. |
| PD-1 (25-160) | MLDSPDRPWNPPTFSPALLVVTEGDNATFTCSFSNTSESFVLNWYRMSPSNQTDKLAAFPEDRSQPGQDSRFRVTQLPNGRDFHMSVVRARRNDSGTYLCGAISLAPKAQIKESLRAELRVTERRAEVPTAHPSPSP | 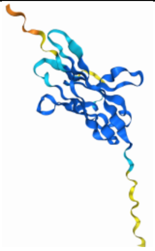 | Uniprot: Q15116. Human, cell membrane, inhibitor of T-cell activation, extracellular domain. |
| EIF4A | MSASQDSRSRDNGPDGMEPEGVIESNWNEIVDSFDDMNLSESLLRGIYAYGFEKPSAIQQRAILPCIKGYDVIAQAQSGTGKTATFAISILQQIELDLKATQALVLAPTRELAQQIQKVVMALGDYMGASCHACIGGTNVRAEVQKLQMEAPHIIVGTPGRVFDMLNRRYLSPKYIKMFVLDEADEMLSRGFKDQIYDIFQKLNSNTQVVLLSATMPSDVLEVTKKFMRDPIRILVKKEELTLEGIRQFYINVEREEWKLDTLCDLYETLTITQAVIFINTRRKVDWLTEKMHARDFTVSAMHGDMDQKERDVIMREFRSGSSRVLITTDLLARGIDVQQVSLVINYDLPTNRENYIHRIGRGGRFGRKGVAINMVTEEDKRTLRDIETFYNTSIEEMPLNVADLI | 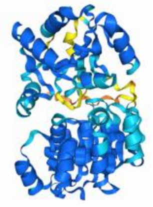 | Uniprot: P60842. Human, cytoplasm, translation initiation factor, full length. |

| | | | |
|---|---|---|---|
| SasA (154-211) | MADNNFNKEQQNAFYEILNMPNLNE EQRNGFIQSLKDDPSQSANLLSEAKK LNESQAPK | 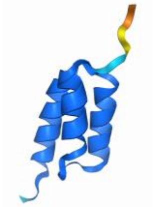 | Uniprot: P02976. *Staphylococcus aureus*, secreted, IgG binding protein, Ig binding domain A. |
| LC3B | MPSEKTFKQRRTFEQRVEDVRLIRE QHPTKIPVIIERYKGEKQLPVLDKTKF LVPDHVNMSELIKIIRRRLQLNANQAF FLLVNGHSMVSVSTPISEVYESEKDE DGFLYMVYASQETFGMKLSV | 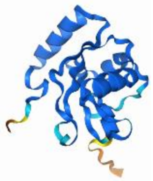 | Uniprot: A6NCE7. Human, cytoplasm/autophagosome membrane, autophagy cargo receptor, full length. |
| ALKBH5 | MAAASGYTDLREKLKSMTSRDNYKA GSREAAAAAAAAVAAAAAAAAAAEP YPVSGAKRKYQEDSDPERSDYEEQQ LQKEEEARKVKSGIRQMRLFSQDEC AKIEARIDEVVSRAEKGLYNEHTVDR APLRNKYFFGEGYTYGAQLQKRGPG QERLYPPGDVDEIPEWVHQLVIQKLV EHRVIPEGFVNSAVINDYQPGGCIVS HVDPIHIFERPIVSVSFFSDSALCFGC KFQFKPIRVSEPVLSLPVRRGSVTVL SGYAADEITHCIRPQDIKERRAVIILRK TRLDAPRLETKSLSSSVLPPSYASDR LSGNNRDPALKPKRSHRKADPDAAH RPRILEMDKEENRRSVLLPTHRRRGS FSSENYWRKSYESSEDCSEAAGSPA RKVKMRRHSRGSRA | 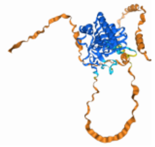 | Uniprot: Q6P6C2. Human, nucleus, RNA demethylase, partially disordered, full length. |
| HIF1b (1-474) | MAATTANPEMTSDVPSLGPAIASGNS GPGIQGGGAIVQRAIKRRPGLDFDDD GEGNSKFLRCDDDQMSNDKERFAR SDDEQSSADKERLARENHSEIERRR RNKMTAYITELSDMVPTCSALARKPD KLTILRMAVSHMKSLRGTGNTSTDGS YKPSFLTDQELKHLILEAADGFLFIVS CETGRVVYVSDSVTPVLNQPQSEWF GSTLYDQVHPDDVDKLREQLSTSEN ALTGRILDLKTGTVKKEGQQSSMRM CMGSRRSFICRMRCGSSSVDPVSVN RLSFVRNRCRNGLGSVKDGEPHFVV VHCTGYIKAWPPAGVSLPDDDPEAG QGSKFCLVAIGRLQVTSSPNCTDMS NVCQPTEFISRHNIEGIFTFVDHRCVA TVGYQPQELLGKNIVEFCHPEDQQLL RDSFQQVVKLKGQVLSVMFRFRSKN QEWLWMRTSSFTFQNPYSDEIEYIIC TNTNVKNSSQEPRPT | 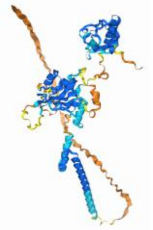 | Uniprot: P27540. Human, nucleus, heterodimerizes with ARNT to induce transcription, all structured regions, but only the N-terminal disordered domain. |
| IFNG (24-156) | MQDPYVKEAENLKKYFNAGHSDVAD NGTLFLGILKNWKEESDRKIMQSQIV SFYFKLFKNFKDDQSIQKSVETIKED MNVKFFNSNKKKRDDFEKLTNYSVT DLNVQRKAIHELIQVMAELSPAAKTG KRKRSQ | 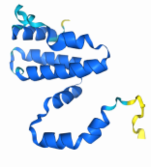 | Uniprot: P01579. Human, secreted, type II interferon, structured secreted domain only. |

| | | | |
|---|---|---|---|
| MST2 (313-437) | MENSDEDELDSHTMVKTSVESVGTMRATSTMSEGAQTMIEHNSTMLESDLGTMVINSEDEEEEDGTMKRNATSPQVQRPSFMDYFDKQDFKNKSHENCNQNMHEPFPMSKNVFPDNWKVPQDGDF | 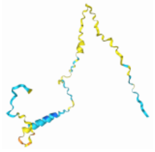 | Uniprot: Q13188. Human, cytoplasm/nucleus, pro-apoptotic kinase, disordered domain. |
| p53 (1-293) | MEEPQSDPSVEPPLSQETFSDLWKLLPENNVLSPLPSQAMDDLMLSPDDIEQWFTEDPGPDEAPRMPEAAPPVAPAPAAPTPAAPAPAPSWPLSSSVPSQKTYQGSYGFRLGFLHSGTAKSVTCTYSPALNKMFCQLAKTCPVQLWVDSTPPPGTRVRAMAIYKQSQHMTEVVRRCPHHERCSDSDGLAPPQHLIRVEGNLRVEYLDDRNTFRHSVVVPYEPPEVGSDCTTIHYNYMCNSSCMGGMNRRPILTIITLEDSSGNLLGRNSFEVRVCACPGRDRRTEEENLRKKG | 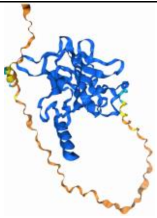 | Uniprot: P04637. Human, nucleus, transcription factor, N-terminal disordered domain and structured domain. |
| PD-L1 (18-239) | MAFTVTVPKDLYVVEYGSNMTIECKFPVEKQLDLAALIVYWEMEDKNIIQFVHGEEDLKVQHSSYRQRARLLKDQLSLGNAALQITDVKLQDAGVYRCMISYGGADYKRITVKVNAPYNKINQRILVVDPVTSEHELTCQAEGYPKAEVIWTSSDHQVLSGKTTTTNSKREEKLFNVTSTLRINTTTNEIFYCTFRRLDPEENHTAELVIPELPLAHPPNERT | 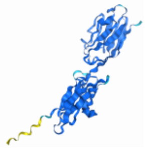 | Uniprot: Q9NZQ7. Human, cell membrane, important in T-cell activation, extracellular domain only. |
| Bax peptide (58-72) | MKKLSECLKRIGDELDS | 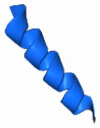 | Uniprot: Q07812. Human, cytoplasm, mitochondrion membrane, involved in mitochondrial apoptosis, BH3 domain. |
| p53 AD (1-61) | MEEPQSDPSVEPPLSQETFSDLWKLLPENNVLSPLPSQAMDDLMLSPDDIEQWFTEDPGPD | 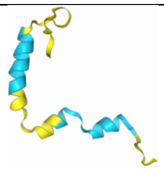 | Uniprot: P04637. Human, nucleus, transcription factor, N-terminal disordered activation domain. |
| CD3d | MEHSTFLSGLVLATLLSQVSPFKIPIEELEDRVFVNCNTSITWVEGTVGTLLSDITRLDLGKRILDPRGIYRCNGTDIYKDKESTVQVHYRMCQSCVELDPATVAGIIVTDVIATLLLALGVFCFAGHETGRLSGAADTQALLRNDQVYQPLRDRDDAQYSHLGGNWARNK | 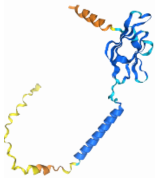 | Uniprot: P04234. Human, cell membrane, T-cell surface receptor, single pass-transmembrane protein, fully disordered, full length. |
| Myc (17-454) | MPLNVSFTNRNYDLDYDSVQPYFYCDEEENFYQQQQQSELQPPAPSEDIWKKFELLPTPPLSPSRRSGLCSPSYVAVTPFSLRGDNDGGGGSFSTADQLEMVTELLGGDMVNQSFICDPDDETFIKNIIIQDCMWSGFSAAAKLVSEKLASYQAARKDSGSPNPARGHSVCSTSSLYLQDLSAAASECIDPSVVFPYPLNDSSSPKSCASQDSSAFSPSSDSLLSSTESSPQGSPEPLVLHEETPPTTSSDSEEEQEDEEEIDVVSVEKRQAPGKRSESGSP | 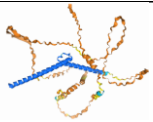 | Uniprot: P01106. Human, nucleus, proto oncogene transcription factor, intrinsically disordered, full length. |

| | | | |
|---|---|---|---|
| | SAGGHSKPPHSPLVLKRCHVSTHQHNYAAPPSTRKDYPAAKRVKLDSVRVLRQISNNRKCTSPRSSDTEENVKRRTHNVLERQRRNELKRSFFALRDQIPELENNEKAPKVVILKKATAYILSVQAEEQKLISEEDLLRKRREQLKHKLEQLRNSCA | | |
| CRBN (47-435) | MINFDTSLPTSHTYLGADMEEFHGRTLHDDDSCQVIPVLPQVMMILIPGQTLPLQLFHPQEVSMVRNLIQKDRTFAVLAYSNVQEREAQFGTTAEIYAYREEQDFGIEIVKVKAIGRQRFKVLELRTQSDGIQQAKVQILPECVLPSTMSAVQLESLNKCQIFPSKPVSREDQCSYKWWQKYQKRKFHCANLTSWPRWLYSLYDAETLMDRIKKQLREWDENLKDDSLPSNPIDFSYRVAACLPIDDVLRIQLLKIGSAIQRLRCELDIMNKCTSLCCKQCQETEITTKNEIFSLSLCGPMAAYVNPHGYVHETLTVYKACNLNLIGRPSTEHSWFPGYAWTVAQCKICASHIGWKFTATKKDMSPQKFWGLTRSALLPTIPDTEDEISP | 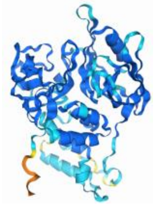 | Uniprot: Q96SW2. Human, cytoplasm/nucleus, E3 ligase, structured domain only. |
| MAX (23-87) | MDKRAHHNALERKRRDHIKDSFHSLRDSVPSLQGEKASRAQILDKATEYIQYMRRKNHTHQQDIDDLKRQNALLEQQGEHP | 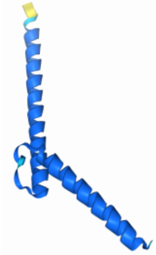 | Uniprot: P61244. Human, nucleus, transcription factor, intrinsically disordered, DNA binding domain only. |
| PPP1R11 | MAEAGAGLSETVTETTVTVTTEPENRSLTIKLRKRKPEKKVEWTSDTVDNEHMGRRSSKCCCIYEKPRAFGESSTESDEEEEEGCGHTHCVRGHRKGRRRATLGPTPTTPPQPPDPSQPPPGPMQH | 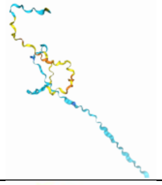 | Uniprot: O60927. Human, cytoplasm, nucleus, E3 ligase, intrinsically disordered, full-length. |
| Alpha amylase (1-226) | MKLFWLLFTIGFCWAQYSSNTQQGRTSIVHLFEWRWVDIALECERYLAPKGFGGVQVSPPNENVAIHNPFRPWWERYQPVSYKLCTRSGNEDEFRNMVTRCNNVGVRIYVDAVINHMCGNAVSAGTSSTCGSYFNPGSRDFPAVPYSGWDFNDGKCKTGSGDIENYNDATQVRDCRLSGLLDLALGKDYVRSKIAEYMNHLIDIGVAGFRIDASKHMWPGDIKAIL | 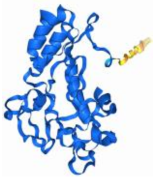 | Uniprot: P0DUB6. Human, secreted, glycosyl hydrolase, inactive fragment. |
| PMP22 | MLLLLLSIIVLHVAVLVLLFVSTIVSQWIVGNGHATDLWQNCSTSSSGNVHHCFSSSPNEWLQSVQATMILSIIFSILSLFLFFCQLFTLTKGGRFYITGIFQILAGLCVMSAAAIYTVRHPEWHLNSDYSYGFAYILAWVAFPLALLSGVIYVILRKRE | 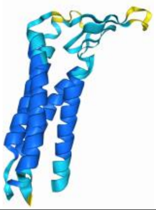 | Uniprot: Q01453. Human, cell membrane, involved in myelination, full length. |

| | | | |
|---|---|---|---|
| BAX (42-153) | MIAAVDTDSPREVFFRVAADMFSDGNFNWGRVVALFYFASKLVLKALCTKVPELIRTIMGWTLDFLRERLLGWIQDQGGWVRLLKPPHPHHRALTTAPAPPSLPPATPLGPW | 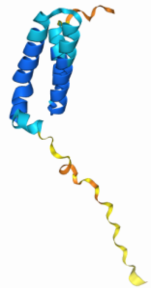 | Uniprot: Q07812. Human, cytoplasm, mitochondrion membrane, involved in mitochondrial apoptosis, isoform gamma. |
| Keap1 (321-611) | MAPKVGRLIYTAGGYFRQSLSYLEAYNPSDGTWLRLADLQVPRSGLAGCVVGGLLYAVGGRNNSPDGNTDSSALDCYNPMTNQWSPCAPMSVPRNRIGVGVIDGHIYAVGGSHGCIHHNSVERYEPERDEWHLVAPMLTRRIGVGVAVLNRLLYAVGGFDGTNRLNSAECYYPERNEWRMITAMNTIRSGAGVCVLHNCIYAAGGYDGQDQLNSVERYDVETETWTFVAPMKHRRSALGITVHQGRIYVLGGYDGHTFLDSVECYDPDTDTWSEVTRMTSGRSGVGVAVTME | 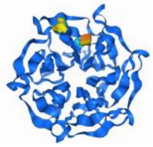 | Uniprot: Q14145. Human, cytoplasm/nucleus, E3 ligase, Kelch domains. |
| HSPB1 | MTERRVPFSLLRGPSWDPFRDWYPHSRLFDQAFGLPRLPEEWSQWLGGSSWPGYVRPLPPAAIESPAVAAPAYSRALSRQLSSGVSEIRHTADRWRVSLDVNHFAPDELTVKTKDGVVEITGKHEERQDEHGYISRCFTRKYTLPPGVDPTQVSSSLSPEGTLTVEAPMPKLATQSNEITIPVTFESRAQLGGPEAAKSDETAAK | 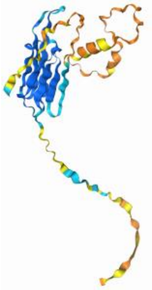 | Uniprot: P04792. Human, cytoplasm/nucleus, molecular chaperone, mostly disordered, full length. |
| HMGB1 | MGKGDPKKPRGKMSSYAFFVQTCREEHKKKHPDASVNFSEFSKKCSERWKTMSAKEKGKFEDMAKADKARYEREMKTYIPPKGETKKKFKDPNAPKRPPSAFFLFCSEYRPKIKGEHPGLSIGDVAKKLGEMWNNTAADDKQPYEKKAAKLKEKYEKDIAAYRAKGKPDAAKKGVVKAEKSKKKKEEEEDEEDEEDEEEEEDEEDEDEEEDDDDE | 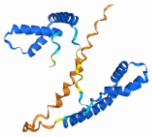 | Uniprot: P09429. Human, nucleus, transcription factor, partially disordered, full length. |

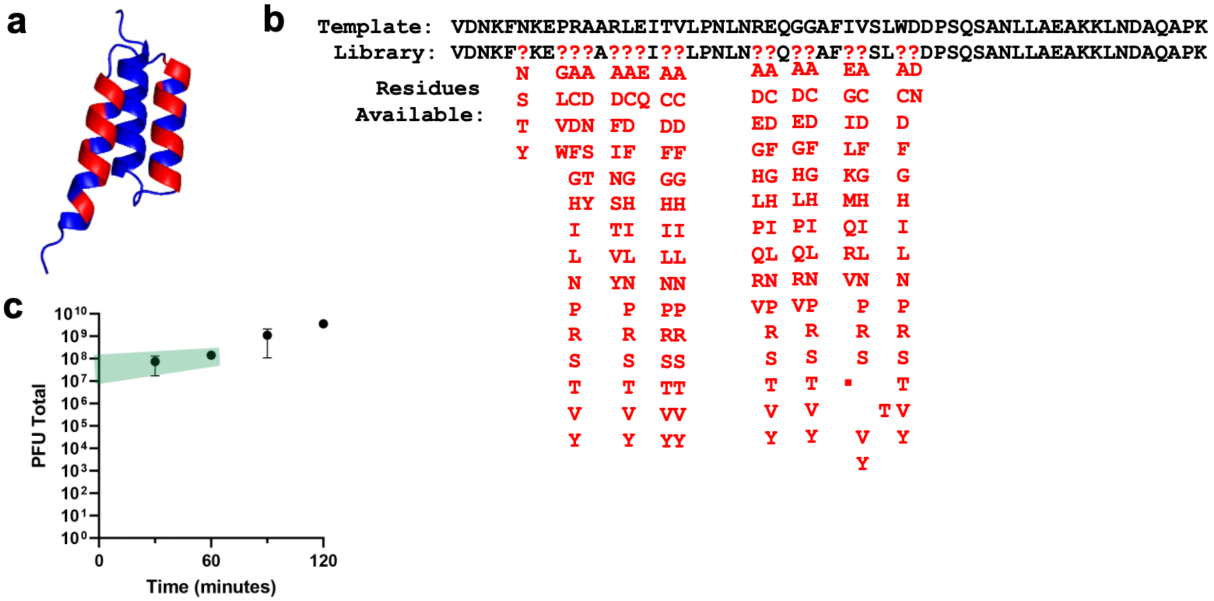


**Supplementary Fig. 1. Construction of the Affibody library.**

**a)** The affibody structure with constant residues in blue and randomized residues in red. **b)** The template phage used when cloning the library and the position and possible codons in the library variants. **c)** Monitoring of total phage after transformation. The early timepoints (highlighted in green) are a lower-bound on the number of variants in the library ($5*10^7$) and the 2 h timepoint is the upper-bound ($2*10^9$) as the phage were separated from the transformed cells at this timepoint. We refer to this as a $10^8$ variant library.

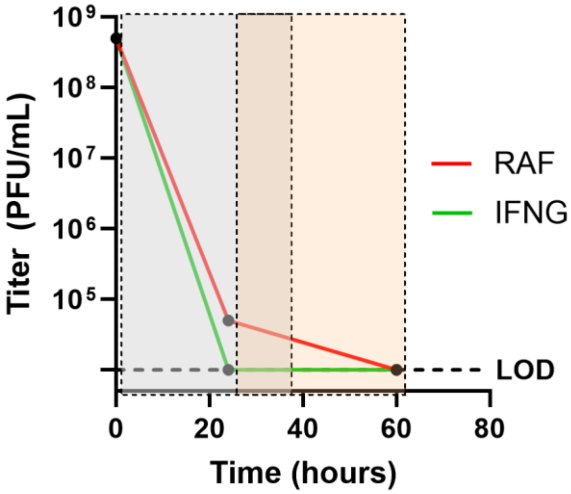


**Supplementary Fig. 2**. **PACE of the affibody library (Supplementary Fig. 1) on two targets – RAF and IFNG – with two different selection strength +APs (ori, gIII/RNAP$_C$ RBS strength).**

p15 SD8/SD8 to pSC101 SD8/SD8 for RAF and from pSC101 SD8/SD8 to p15 SD8/sd5 for IFNG. In addition to the +AP, each strain had a ZB$_{neg}$ -AP (20-1) and MP6 (see **Supplementary Table 1**). The first strain proceeded for 24 h (at which point a titer was collected) and then mixed 1:1 with the second strain for 12 h, then finally the second strain was used alone for 24 h at which point the titer was again collected. A single 20 mL lagoon was seeded with $10^{10}$ PFU library phage and filled with strain 1 and with arabinose to induce mutagenesis, after 1 h, the lagoon was flowed out to waste at a rate of 1 vol/h. During the mixing experiment, each selection strain chemostat flowed at half speed to maintain 1 vol/h.

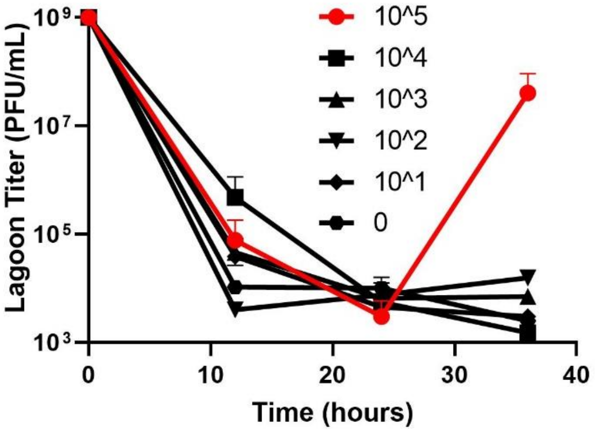


**Supplementary Fig. 3. PACS with mock library.**

The KRAS WT (+AP; 31-69)/$ZB_{neg}$ (-AP; 20-06) selection strain and active (RAF WT) and inactive (Affitin (SasA)) phage used in **Fig. 2** was used here in PACS (duplicate 20 mL lagoons were seeded with $10^{10}$ PFU library phage, and after 1 h, the lagoons were flowed out to waste at a rate of 1 vol/h. The mock library consisted of $10^{10}$ inactive phage mixed with varying amounts of active phage (legend). By 36 h, all lagoons had gone extinct except for the mock library spiked with $10^5$ active phage. Titers measured by plaque assay in duplicate (LOD of $10^3$ PFU/mL).

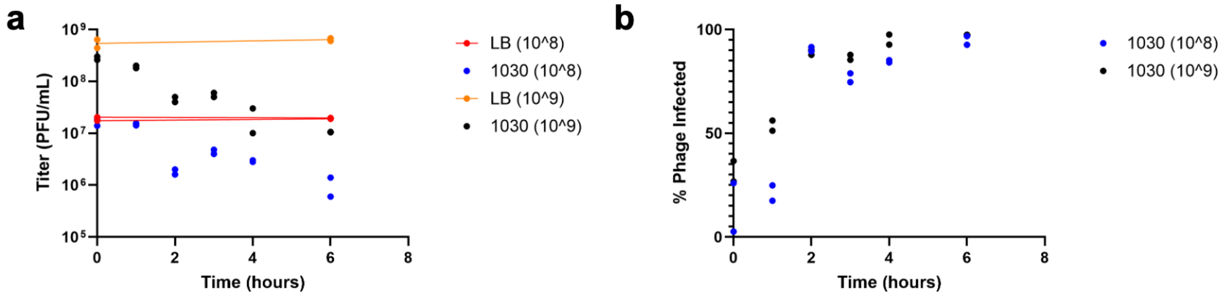


**Supplementary Fig. 4. Rate of phage infection.**

**a)** Phage (RNAP$_N$-RAF WT) were added to 1 mL LB or 1030 at two different concentrations (~ equal cell-to-phage ratio for 1030 and roughly 10x cell-phage ratio for 1030) and their titers were monitored over time. 1030 cells were grown to an OD600 of 0.3 prior to infection. 1030 cells are phage infectible, but do not support phage replication as they cannot supply *giii*. Duplicate tubes were performed for each combination and the titers were measured overtime. Phage and cells for the 0 hour timepoint were separated by centrifugation ~ 1 minute after combining and mixing. **b)** Percent of phage infected were determined by the difference between initial number of phage (determined as the average titer of phage in LB) and the remaining number of phage (as determined by the titer of the 1030 culture). Phage infections plateau at ~95% at 6 hours. These data indicate that the rate at which phage infect cells decreases logarithmically as the concentration of non-infected (and therefore infectable) cells decreases.

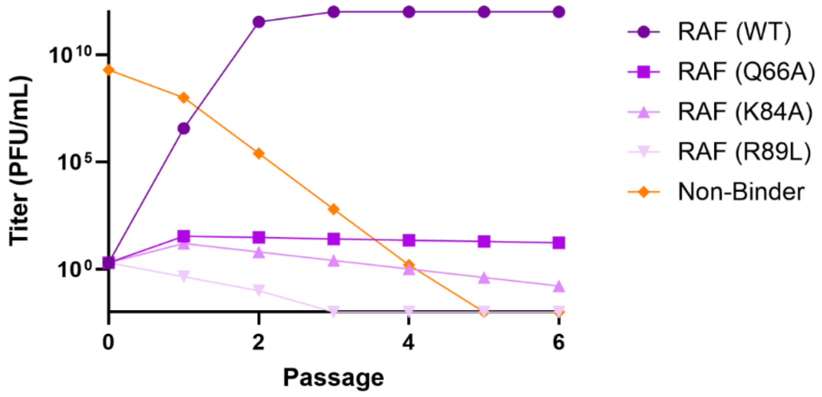


**Supplementary Fig. 5. Amplification rates of Raf phage using optimized conditions.**

Using the amplification rates in **Fig. 2c**, we predicted the expected enrichment of RAF mutant phage and de-enrichment of non-binder (Affitin (SasA)) phage based on an initial 5 mL volume, 10 or $10^{10}$ PFU respectively, and a 5% transfer between passages. We constrained the bottom to 0.01 and the top to $10^{12}$ PFU/mL respectively. The Inactive phage goes to extinction by passage 4 and the active variant enriches to saturation by passage 2. We expect that both rates are over-estimated by the amplification rate: RAF WT should amplify more slowly than in the amplification assay as most cells have been infected by an inactive phage and are therefore non-infectible; similarly, the non-binder should de-enrich more slowly as some fraction of phage will not infect a cell in the initial passage(s). Therefore, we settled on testing 4 passages in **Fig. 2e**.

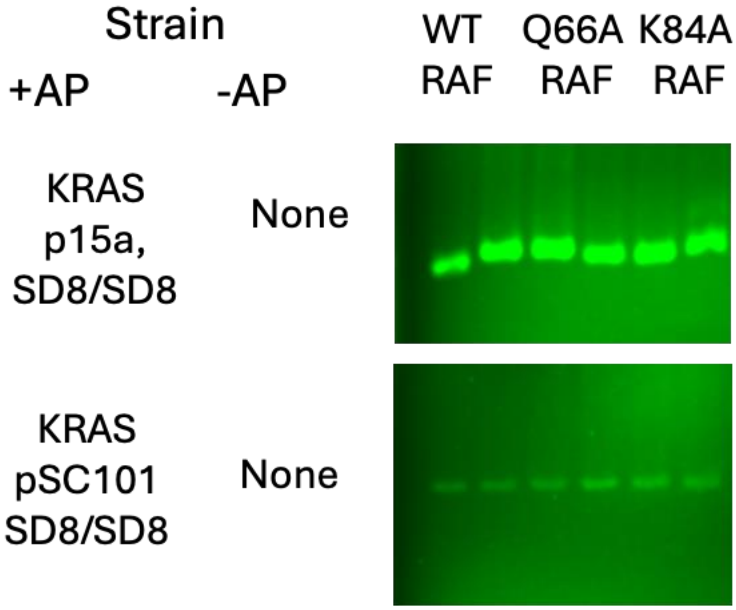


**Supplementary Fig. 6**. **PANCS with mock library and varied +APs.**

The active (RAF WT, Q66A K84A) and inactive (Affitin (SasA)) phage was used in mock library PANCS using different +AP strengths (31-69 (top) and 31-70 (bottom), see Supplementary Table 1). Each PANCS had a 1 mL passage volume with 5% transferred between passages (4 total) that was seeded with $10^9$ PFU inactive phage and 10 PFU active phage (duplicate PANCS). PCR was used to estimate the endpoint titer of each PANCS. The PCR cycle number, 25, was optimized so that $10^6$ PFU/mL produced a faint band (similar to that observed in the lower gel). The p15a, SD8/SD8 +AP strength was chosen for +AP designs moving forward.

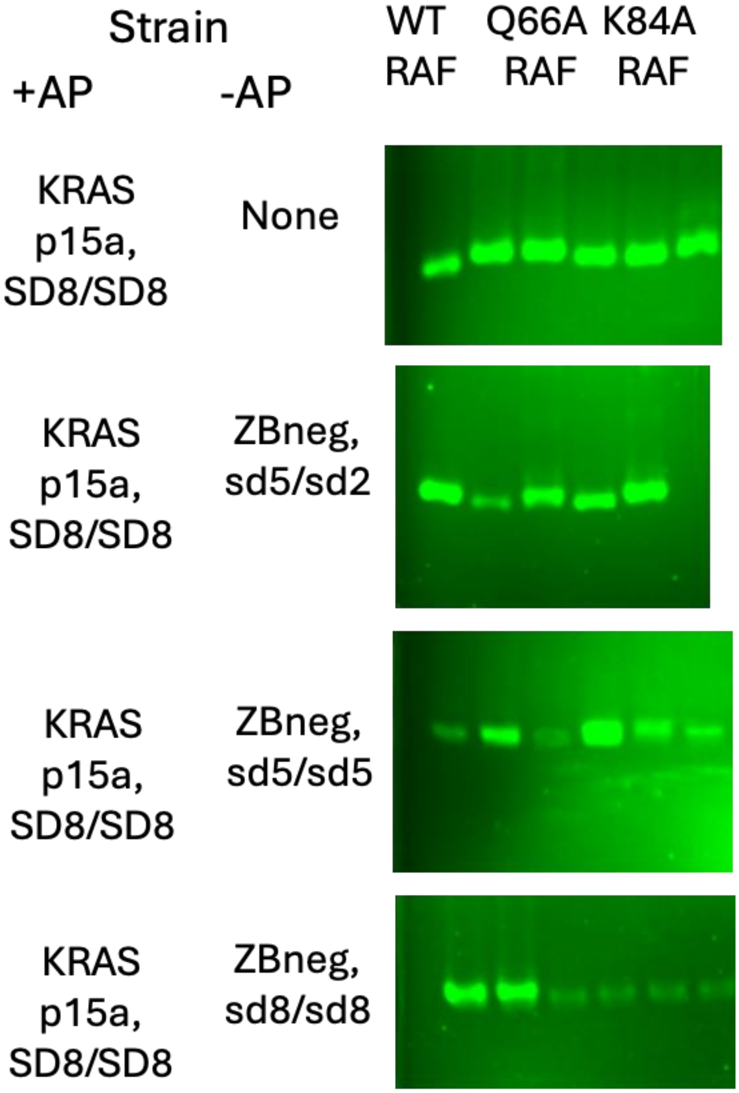


**Supplementary Fig. 7. PANCS with mock library and varied -APs.**

The active (RAF WT, Q66A K84A) and inactive (Affitin (SasA)) phage was used in mock library PANCS using different -AP strengths (top-bottom: none, 20-6, 20-5, and 20-1, see Supplementary Table 1). Each PANCS had a 1 mL passage volume with 5% transferred between passages (4 total) that was seeded with $10^9$ PFU inactive phage and 10 PFU active phage (duplicate PANCS). PCR was used to estimate the endpoint titer of each PANCS. The PCR cycle number, 25, was optimized so that $10^6$ PFU/mL produced a faint band (similar to that observed in the lowest gel on the right). The sd8/sd8 -AP strength was chosen for PANCS moving forward.

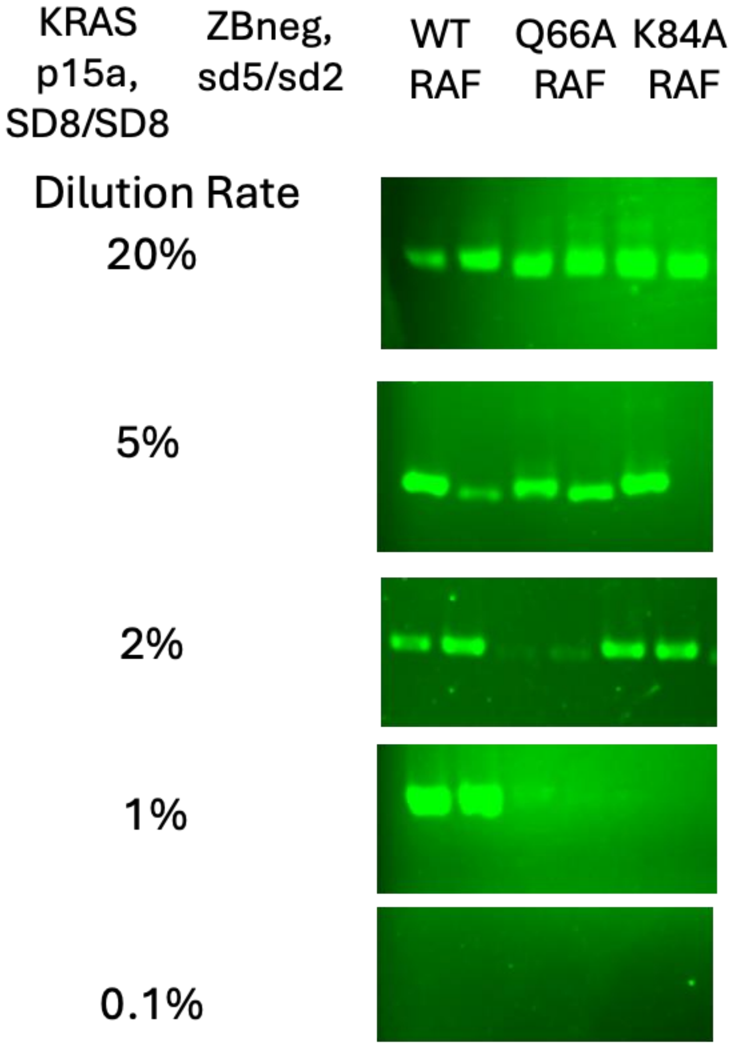


**Supplementary Fig. 8. PANCS with mock library and varied transfer rates.**

The active (RAF WT, Q66A K84A) and inactive (Affitin (SasA)) phage was used in mock library PANCS using different transfer rates. Each PANCS had a 1 mL passage volume with varying amounts transferred between passages (4 total) that was seeded with $10^9$ PFU inactive phage and 10 PFU active phage (duplicate PANCS). The same selection strain was used in each of these PANCS. PCR was used to estimate the endpoint titer of each PANCS. The PCR cycle number, 25, was optimized so that $10^6$ PFU/mL produced a faint band.

**a**

| Strain | Prior Infect | Active | Inactive | PFU/mL | Fold Change |
|---|---|---|---|---|---|
| 31-69 | --- | 10^6 | 0 | 245000000 | 490 |
| 31-69 | --- | 10^6 | 10^10 | 900000 | 1.8 |
| 31-69 | 10^10 | 10^6 | 0 | 1325000 | 2.65 |
| 31-69 | 10^10 | 10^6 | 10^10 | 600000 | 1.2 |
| 31-69/20-6 | --- | 10^6 | 0 | 223750000 | 447.5 |
| 31-69/20-6 | --- | 10^6 | 10^10 | 1000000 | 2 |
| 31-69/20-6 | 10^10 | 10^6 | 0 | 700000 | 1.4 |
| 31-69/20-6 | 10^10 | 10^6 | 10^10 | 875000 | 1.75 |

**b**

| Phage/Cell ratio | Volume | Result |
|---|---|---|
| 10x | 1 mL | Inactive only |
| 1x | 1 mL | Both active and inactive (~1:1) |
| 0.1x | 1 mL | Active only |
| 10x | 10 mL | Inactive only |
| 1x | 10 mL | Active only |
| 0.1x | 10 mL | Active only |
| 10x | 100 mL | Inactive only |
| 1x | 100 mL | Inactive only |
| 0.1x | 100 mL | Active only |

**Supplementary Fig. 9. PANCS with mock library and varied initial cell-to-phage ratio.**

**a)** Amplification rate of $5*10^5$ active (RAF WT) phage in 1 mL of KRAS +AP selection strain grown for 12 h with either no inactive phage, simultaneous addition of $10^{10}$ inactive phage (>10x the number of cells), or prior infection (1 h of $10^{10}$ inactive phage (>10x the number of cells). Active phage were monitored using activity dependent plaque assays (active phage shows robust plaques, inactive do not produce plaques). Both simultaneous and prior infection of excess phage prevented replication of the active phage when the phage-to-cell ratio was >10:1. **b)** The active (RAF WT) and inactive (Affitin (SasA)) phage was used in mock library PANCS on the KRAS (+AP)/$ZB_{neg}$ (-AP) using a 5% transfer rate. PCR band size was used to estimate the relative population of active and inactive phage after 4 passages. Each PANCS had a different passage volume and was seeded with 10 PFU active phage and an amount of inactive phage either 10x, 1x, or 0.1x the number of cells in the passage (assuming $10^9$ cells/mL). Red indicates failure to enrich the active phage during PANCS. These data demonstrate the necessity for having at most a 1:1 phage-to-cell ratio in the initial passage.

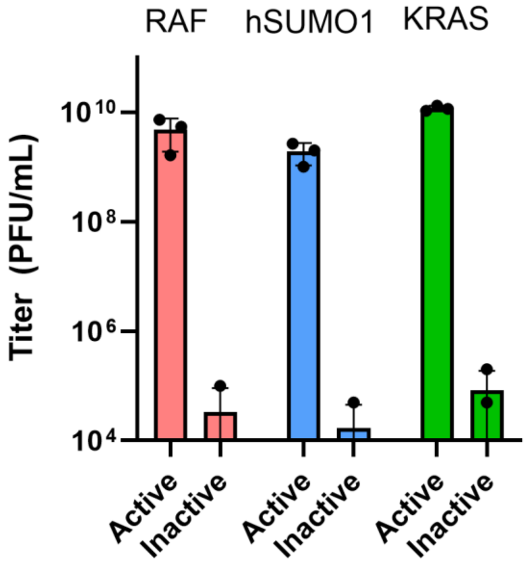


**Supplementary Fig. 10. Mock library selections with additional binder-target pairs.**

Previously published binders to RAF (Affibody (RAF)[2]), hSUMO1 (Monobody (hSUMO1)[3]), and KRAS WT (Monobody (KRAS), NS1[4]) were used in mock library (10 PFU active and $10^{10}$ PFU inactive (Affitin (SasA)[5]) using 4 passages, 5% transfer, 5 mL starting volume, with p15a/SD8/SD8 +APs and sd8/sd5 (20-01) $ZB_{neg}$ -AP. PANCS was performed in triplicate and endpoint (passage 4) titers were measured with an activity independent plaque assay. LOD was $5*10^4$ PFU/mL (0 PFU, data points not shown). Error bars indicate SD.

**Supplementary Table 4. Plaque Assay results for 6-target panel *de novo* library PANCS (Fig. 3a).**

| Target | Titer (PFU/mL)[a] | On-Target Activity Dependent Plaques[b] | Off-Target Activity Dependent Plaques[c] | 1030 Activity Dependent Plaques[d] |
|---|---|---|---|---|
| KRAS G12D | $1*10^9$ | Yes | No | No |
| RAF | $8*10^8$ | Yes | No | No |
| Mdm2 | $1*10^{10}$ | Yes | No | No |
| IFNG | $1*10^{10}$ | Yes | No | No |
| Myc DBD | < LOD[e] | No[f] | No[f] | No[f] |
| Sos1 | < LOD[e] | No[f] | No[f] | No[f] |

[a]Activity independent titer determined from counting plaques using 1030-1059.
[b]Presence of plaques when a plaque assay was performed using the strain used in PANCS.
[c]Presence of plaques when a plaque assay was performed using an off-target hSUMO1 selection strain +AP.
[d]Presence of plaques when a plaque assay was performed using 1030 (no *giii* present in the *E. coli*).
[e]Limit of detection (LOD) of 1 PFU was equal to $5*10^4$ PFU/mL.
[f]Because the titer was below the LOD, the observation of no activity dependent plaques is of limited information as no phage were likely added to the quadrant in the activity dependent plaque assay.

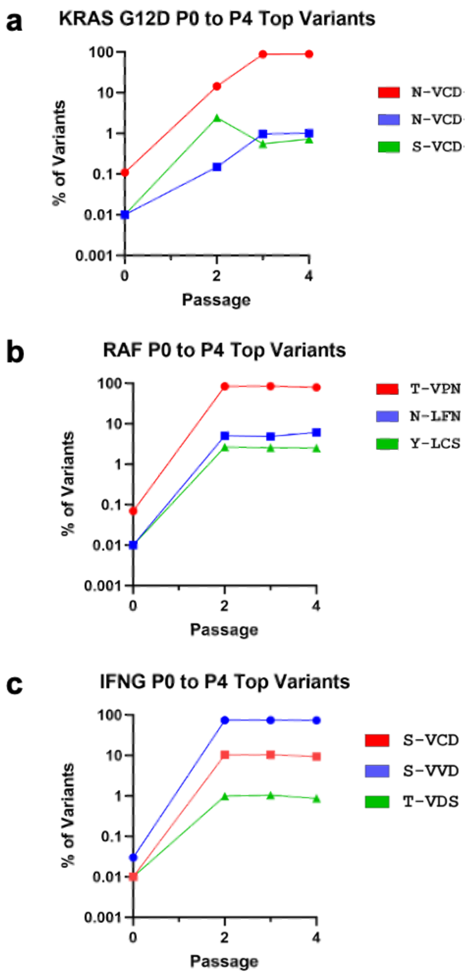


**Supplementary Fig. 11. Tracking enrichment of top variants.**

For the first 6-target panel PANCS, we collected NGS of the initial affibody library, passage 2, passage 3, and passage 4. This allowed us to monitoring enrichment of the active phage across passages as shown for the top three variants at passage 4 for **a)** KRAS G12D, **b)** RAF RBD, and **c)** IFNG. Generally, the relative ratio of variants stabilizes by passage 2 or 3.

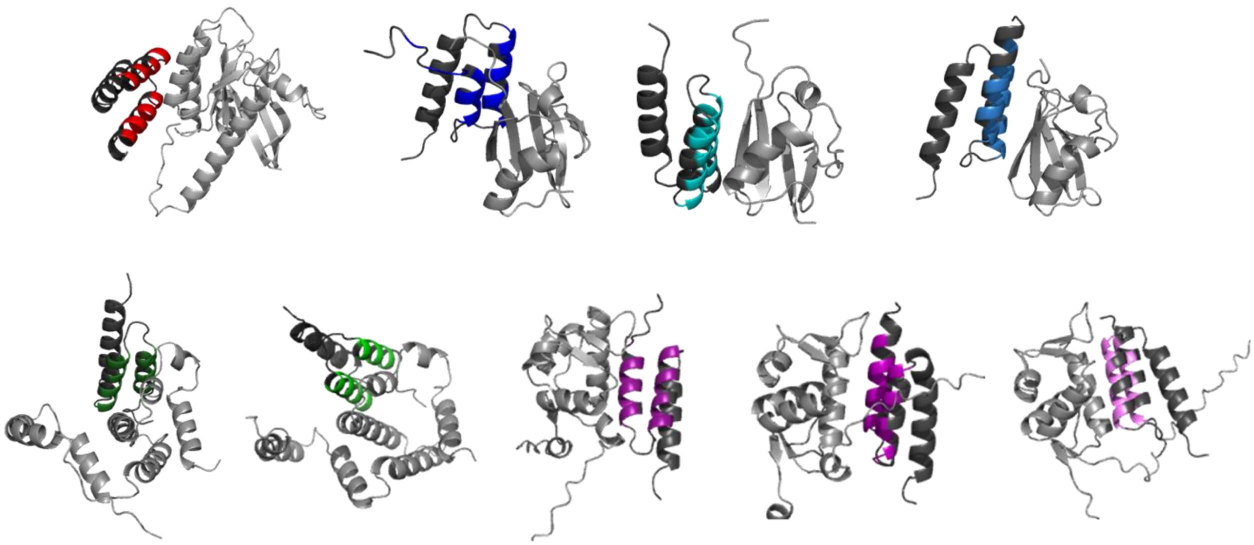

**Supplementary Fig. 12. AlphaFold3 predictions of binder-target interactions for binders from 6-target panel PANCS (Fig. 3b).**

KRAS G12D/N-VCD, RAF/T-VPN, RAF/N-LFN, RAF/Y-LCS, IFNG/S-VVD, IFNG/S-VCD, Mdm2/S-WFY, Mdm2/S-WYS, and Mdm2/T-WDN (top to bottom, left to right). The target is shown in light grey, the binder scaffold in dark grey, and the randomized residues in color (corresponding to colors in **Figure 3B**).

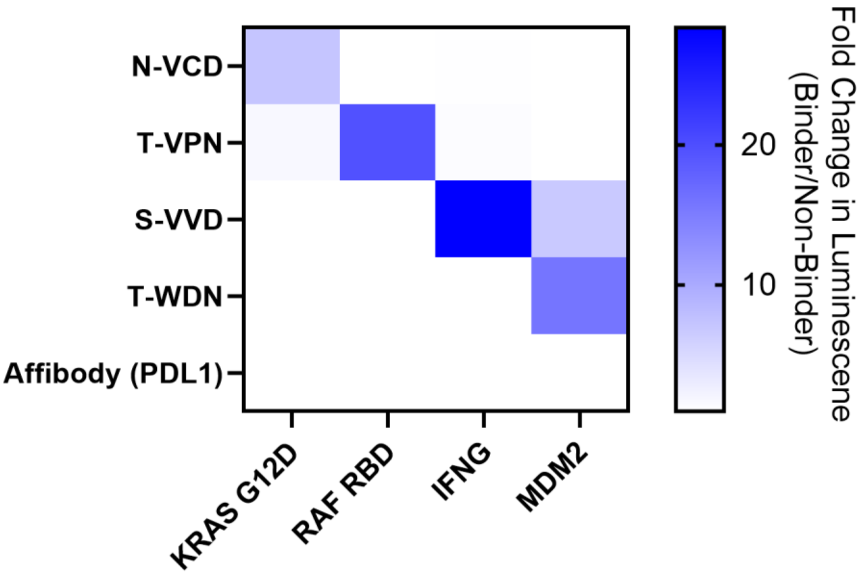


**Supplementary Fig. 13. Specificity of PANCS derived binders.**

Fold-change in luciferase signal for select variants from **Fig. 3b** over the template affibody (PDL1) used in cloning the library (a non-binder). Individual values in the heatmap are the average of triplicates. Only one off-target has a fold-change significantly (p = 0.05) greater than 1.0 (S-VVD on Mdm2).

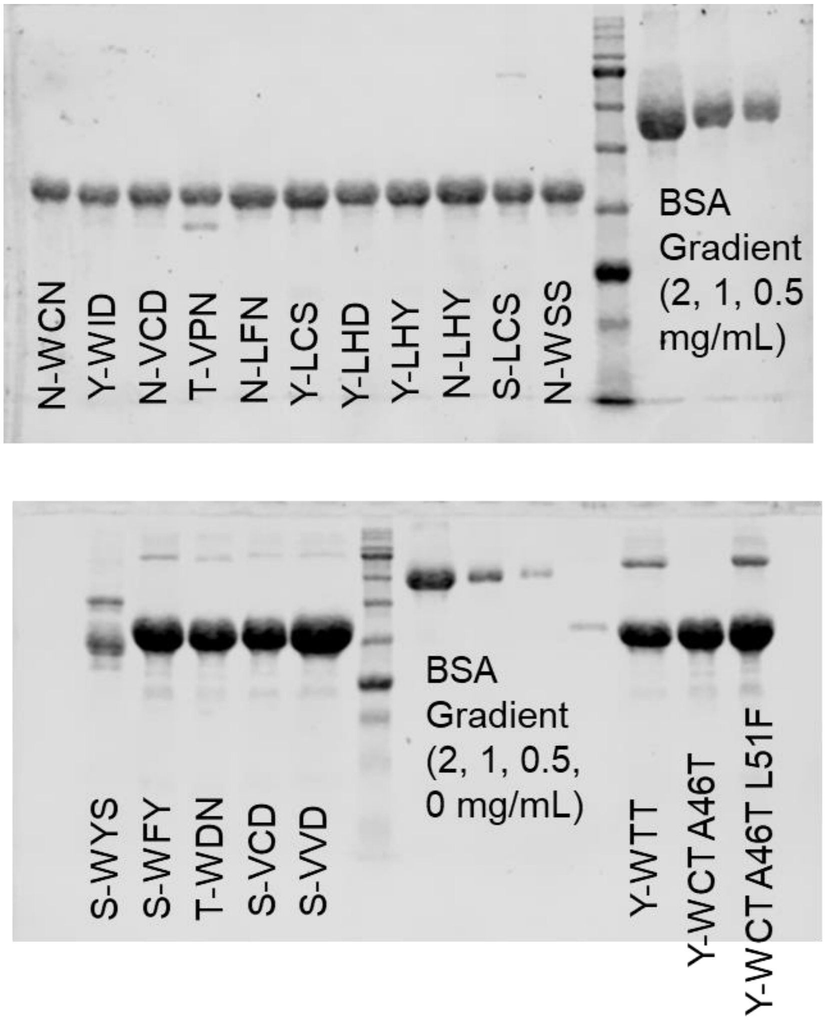


**Supplementary Fig. 14. Purification of GST-tagged binders.**

Each final purified binder was run on a 12% acrylamide SDS PAGE gel and run next to a gradient of BSA. Stained with Coumassie Blue and imaged using a LICOR. All *in vitro* binding affinities reported here (**Supplementary Table 8**) are from these purified proteins.

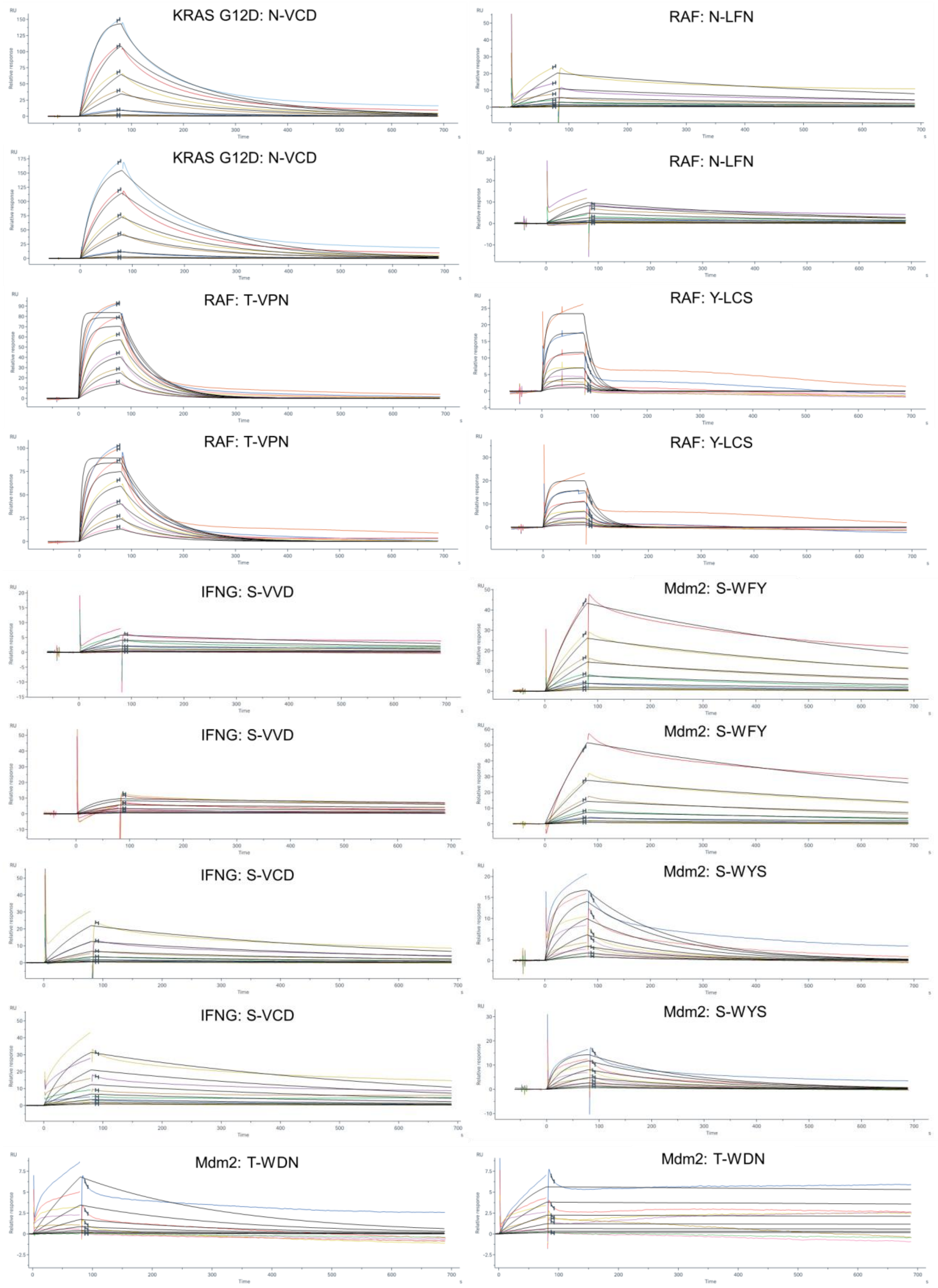
KRAS G12D: N-VCD
RAF: N-LFN
KRAS G12D: N-VCD
RAF: N-LFN
RAF: T-VPN
RAF: Y-LCS
RAF: T-VPN
RAF: Y-LCS
IFNG: S-VVD
Mdm2: S-WFY
IFNG: S-VVD
Mdm2: S-WFY
IFNG: S-VCD
Mdm2: S-WYS
IFNG: S-VCD
Mdm2: S-WYS
Mdm2: T-WDN
Mdm2: T-WDN
RU
Relative response
Time
s

**Supplementary Fig. 15**: **Surface Plasmon Resonance (SPR) dose-response and kinetic model fits for $K_d$s shown in Fig. 3b** (**Supplementary Table 8** for details of model fits).

Each target-binder pair required a different concentration range: KRAS G12D with N-VCD (2000, 1000, 500, 250, 62.5, 15.3, 4, 0 nM); RAF with T-VPN (4000, 2000, 1000, 500, 250, 125, 62.3, and 0 nM), N-LFN (24000, 12000, 6000, 3000, 1500, 750, 375, 0 nM), and Y-LCS (4000, 2000, 1000, 500, 250, 125, 62.3, and 0 nM); IFNG with S-VVD (15000, 10000, 5000, 2500, 1250, 625, 313, 0 nM) and S-VCD (24000, 12000, 6000, 3000, 1500, 750, 375, 0 nM); and Mdm2 with S-WFY (1000, 500, 250, 125, 62.5, 31.3, 15.6, 0 nM), S-WYS (2000, 1000, 500, 250, 62.5, 15.3, 4, 0 nM), and T-WDN (2000, 1000, 500, 250, 62.5, 15.3, 4, 0 nM). All interactions were measured in duplicate and all data was fit to a 1:1 binding model.

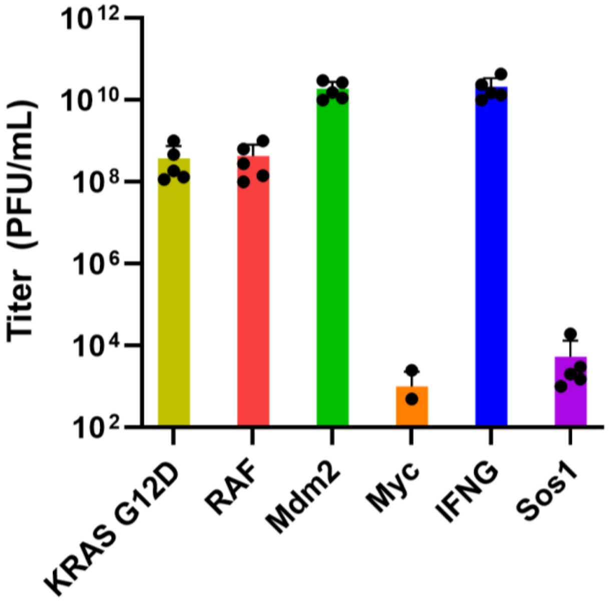


**Supplementary Fig. 16. Replicate 6-target panel PANCS.**

In addition to the initial replicate, technical quadruplicates were performed several months later (**Fig. 3a**, **Supplementary Table 4**). Each of the 5 total PANCS are shown as individual datapoints, error bar indicates SD. For Myc, all four of the parallel replicates had a titer below our LOD, and therefore, are shown as being equal to our LOD (500 PFU/mL).

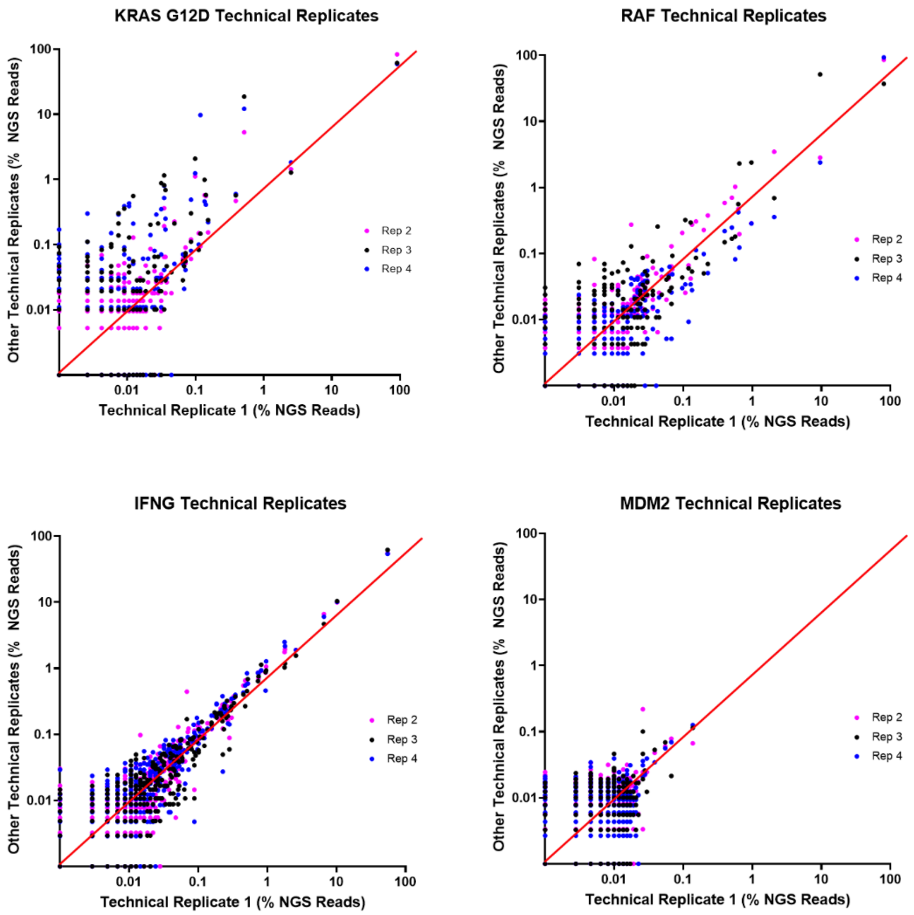


**Supplementary Fig. 17. Comparison of parallel replicates by NGS.**

Percentage of reads for each unique variant in one of the parallel replicate PANCS compared to the percentage of reads for each unique variant in each of the other parallel replicates (**Supplementary Fig. 12**). If a variant was present in one NGS sample but not in another, it was coded as 0.001%. of reads. For variants >0.1% of NGS reads, there was a high correlation between parallel replicates (r = 0.95; average of each pairwise Pearson's Correlation).

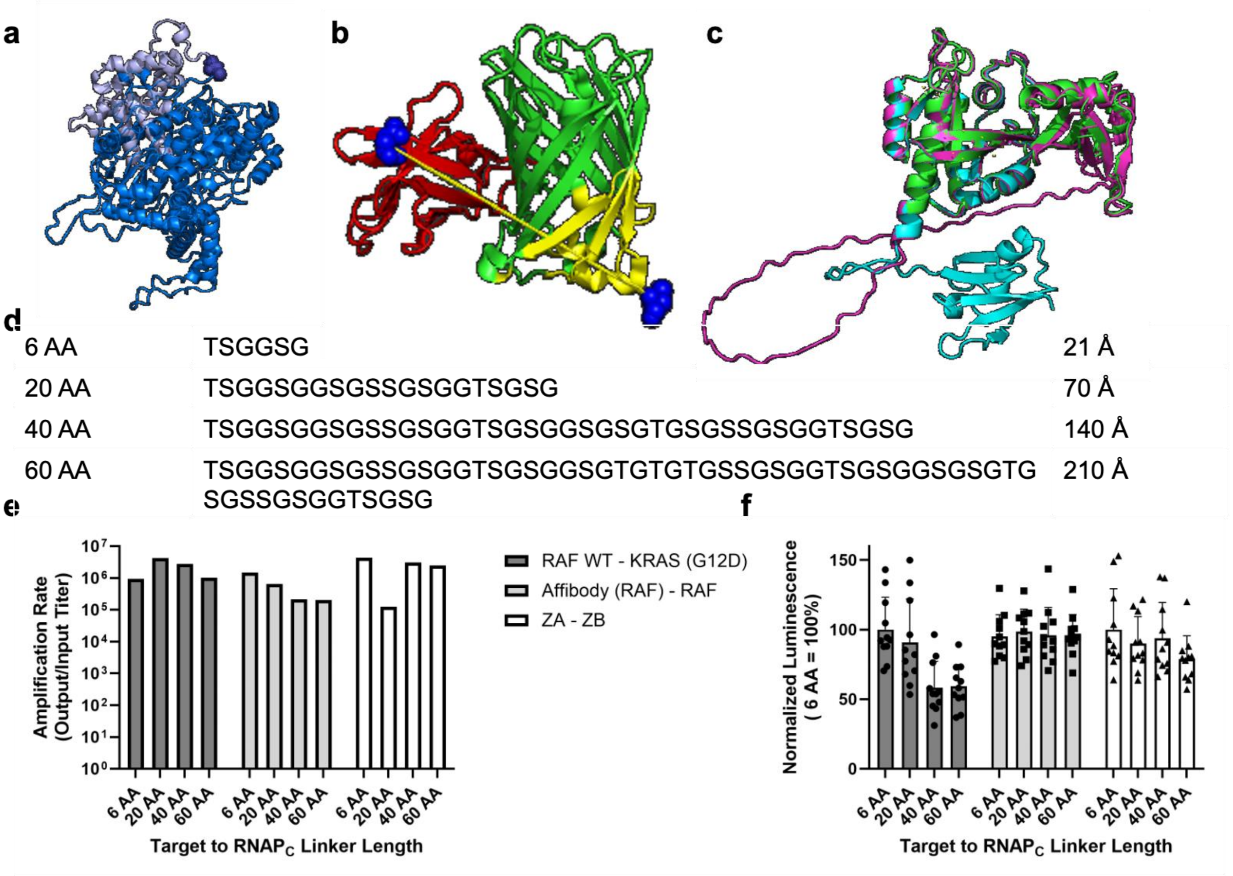


**Supplementary Fig. 18. Extending the linker length connecting the target and RNAP$_C$.**

**a)** The structure of T7 RNAP (1CEZ) with the split site shown in dark blue as spheres, the N-terminal in light blue, and C-terminal in blue. **b)** The structure (3OGO) of a nanobody (red) binding to GFP (green) with each terminus that would be fused to spRNAP in our system shown as blue spheres – the distance between them is >50 Å. On GFP, all atoms within 20 Å are shown in yellow indicating the likely radius reachable for a 6 AA linker. **c)** AlphaFold3 prediction of the interaction of HRAS (1-169) with the RAF (RBD) (green) compared to the predicted folding of HRAS-6aa-8aa-RAF (cyan) and HRAS-60aa-8aa-RAF (purple) aligned to HRAS where 6aa, 8aa, and 60aa refer to linker lengths used in our binder/target fusions with spRNAP. The 60 AA linker is predicted to allow HRAS and RAF to bind in the actual PPI conformation, but the 6 AA linker is not. **d)** Extended linker lengths, sequence, and predicted maximum extended length (assuming 3.5 Å /residue) cloned into several target-RNAP$_C$ fusion +AP and Lux-C plasmids that function well with a short linker. **e)** Phage amplification rate with varied linker lengths for several binder-target pairs (each measured as a single replicate). **f)** Change in luciferase signal when the linker length is extended for several binder-target pairs (same legend as in **e**). Each pair has three biological replicates of technical quadruplicates (12 total measurements). Each luminescence signal is normalized to the 6 AA linker for each target. Overall, E and F suggest that extending the linker length decreases replication and signal, but to a minimal extent (<10x lower amplification rate and <2x lower luciferase signal).

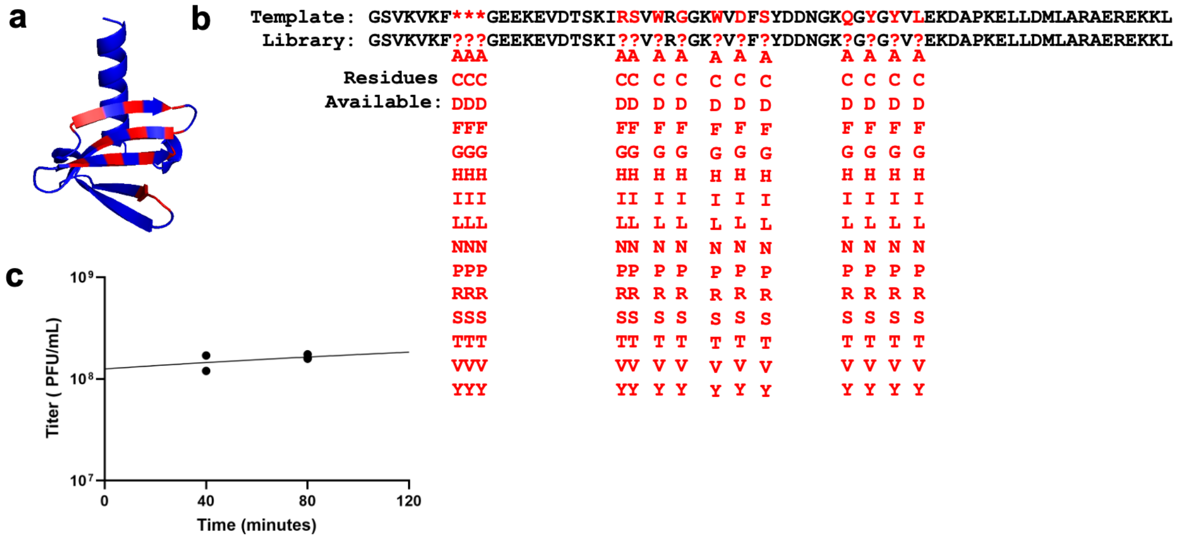


**Supplementary Fig. 19: Construction of the affitin library.**

**a)** The affitin structure with constant residues in blue and randomized residues in red. **b)** The template phage (Affitin (SasA) 3X STOP) used when cloning the library and the position and possible codons in the library variants (all are NNY at the DNA level). **c)** Monitoring of total phage after transformation. Because this transformation utilized the 10β-1059 (non-infectible *E. coli*), the titer does not change much between the timepoints (12% increase). We estimate that the likely total number of variants is similar to total titer measured at 40 minutes: $1.5*10^8$.

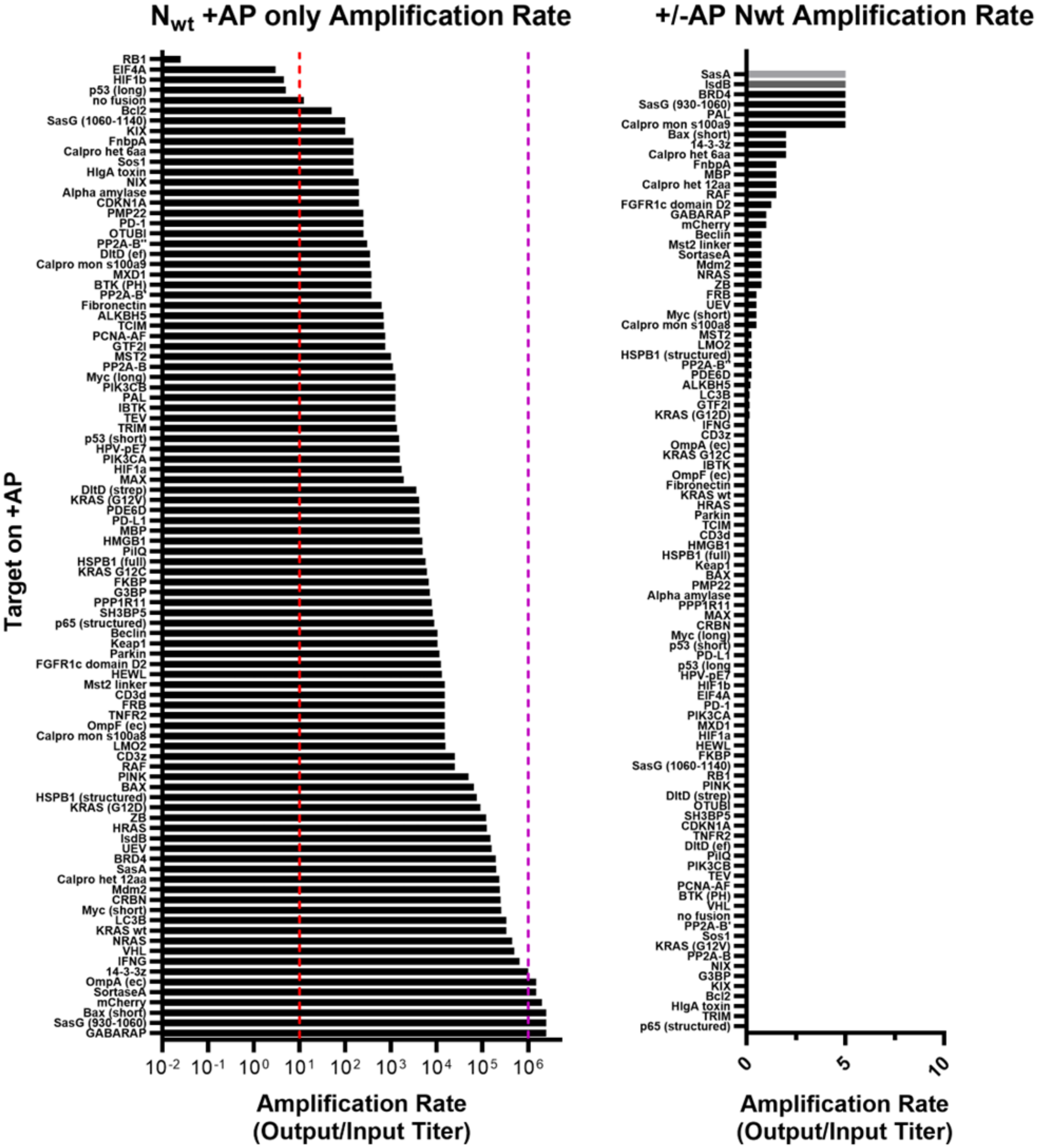


**Supplementary Fig. 20. Background replication rates of all targets.**

Proximity independent, $N_{WT}$ phage amplification rates on 96-target panel +AP only (left) and +AP with -AP (right). Amplification rates obtained using 40,000 PFU RNAP-$N_{WT}$ phage (70-63; **Supplementary Table 1**) as the input and the output was measured using activity independent plaque assays. (Left) Red line indicates where a transfer rate of 10% would prevent enrichment during PANCS; purple line indicates rate for RAF WT phage on KRAS used in optimizing PANCS (**Fig. 2c**).

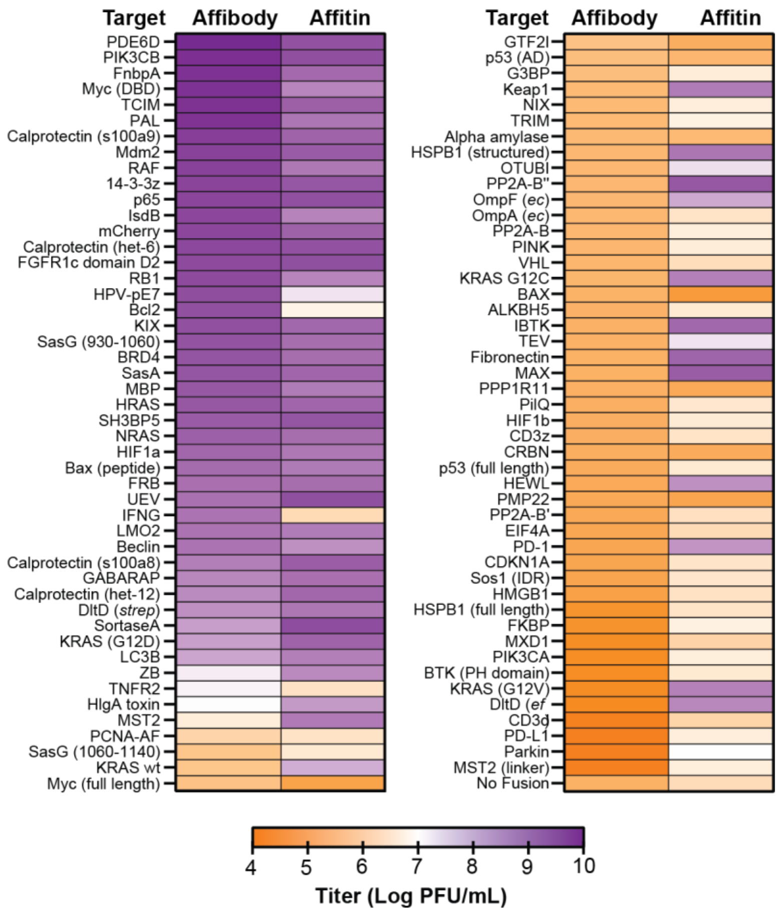


**Supplementary Fig. 21. 95-target panel PANCS endpoint titer heatmap.**

Endpoint titer heatmap for each well of the 192 selections (**Fig. 4a**): legend shows titer (Log PFU/mL) with purple indicating a high titer, white indicating $10^7$, and orange indicating low titers to extinction. Titers determined by single replicate qPCR and compiled in **Supplementary Table 5**.

**Supplementary Table 5. Titers from passage 4 of the 96-target panel PANCS with affibody and affitin libraries (Supplementary Fig. 21).** Titers measured using qPCR.

| Target | Affibody Titer (PFU/mL) | Affitin Titer (PFU/mL) | Target | Affibody Titer (PFU/mL) | Affitin Titer (PFU/mL) |
|---|---|---|---|---|---|
| No Fusion | 1.45E+05 | 1.55E+06 | Myc (fµl length) | 3.63E+05 | 6.61E+04 |
| MST2 (linker) | 2.88E+03 | 3.72E+06 | KRAS wt | 4.27E+05 | 9.55E+07 |
| Parkin | 8.91E+03 | 8.95E+06 | SasG (1060-1140) | 4.27E+05 | 3.02E+06 |
| PD-L1 | 9.55E+03 | 3.55E+06 | PCNA-AF | 1.02E+06 | 2.09E+06 |
| CD3d | 1.05E+04 | 9.55E+05 | MST2 | 3.39E+06 | 4.17E+08 |
| DltD (*ef*) | 1.66E+04 | 2.82E+08 | HlgA toxin | 1.10E+07 | 1.70E+08 |
| KRAS (G12V) | 1.95E+04 | 3.31E+08 | TNFR2 | 1.51E+07 | 2.04E+06 |
| BTK (PH domain) | 2.00E+04 | 2.75E+06 | ZB | 1.74E+07 | 2.57E+08 |
| PIK3CA | 2.04E+04 | 3.72E+06 | LC3B | 1.23E+08 | 3.55E+08 |
| MXD1 | 2.09E+04 | 9.12E+05 | KRAS (G12D) | 1.45E+08 | 8.13E+08 |
| FKBP | 2.75E+04 | 4.17E+06 | SortaseA | 1.48E+08 | 1.78E+09 |
| HSPB1 (fµLl length) | 3.02E+04 | 2.24E+06 | DltD (*strep*) | 2.19E+08 | 4.57E+08 |
| HMGB1 | 6.03E+04 | 2.00E+06 | Calprotectin (het-12) | 2.45E+08 | 7.08E+08 |
| Sos1 (IDR) | 7.41E+04 | 2.40E+06 | GABARAP | 2.82E+08 | 5.50E+08 |
| CDKN1A | 7.94E+04 | 2.34E+06 | Calprotectin (s100a8) | 3.63E+08 | 9.33E+08 |
| PD-1 | 8.32E+04 | 1.91E+08 | Beclin | 5.01E+08 | 2.14E+08 |
| EIF4A | 8.51E+04 | 1.41E+06 | LMO2 | 5.01E+08 | 3.98E+08 |
| PP2A-B' | 9.33E+04 | 1.86E+06 | IFNG | 5.01E+08 | 1.29E+06 |
| PMP22 | 9.55E+04 | 7.59E+04 | UEV | 5.37E+08 | 1.55E+09 |
| HEWL | 1.02E+05 | 2.19E+08 | FRB | 5.50E+08 | 5.75E+08 |
| p53 (fµLl length) | 1.15E+05 | 2.88E+06 | Bax (peptide) | 6.31E+08 | 4.07E+08 |
| CRBN | 1.26E+05 | 1.07E+05 | HIF1a | 7.08E+08 | 4.47E+08 |
| CD3z | 1.29E+05 | 2.19E+06 | NRAS | 9.12E+08 | 5.75E+08 |
| HIF1b | 1.29E+05 | 3.16E+06 | SH3BP5 | 1.05E+09 | 1.32E+09 |
| PilQ | 1.32E+05 | 2.63E+06 | HRAS | 1.17E+09 | 7.41E+08 |
| PPP1R11 | 1.41E+05 | 9.77E+04 | MBP | 1.20E+09 | 3.98E+08 |
| MAX | 1.45E+05 | 1.00E+09 | SasA | 1.32E+09 | 7.41E+08 |
| Fibronectin | 1.48E+05 | 7.76E+08 | BRD4 | 1.35E+09 | 5.75E+08 |
| TEV | 1.51E+05 | 2.14E+07 | SasG (930-1060) | 1.38E+09 | 5.89E+08 |
| IBTK | 1.55E+05 | 6.92E+08 | KIX | 1.48E+09 | 6.92E+08 |
| ALKBH5 | 1.58E+05 | 2.95E+06 | Bcl2 | 1.51E+09 | 4.79E+06 |
| BAX | 1.62E+05 | 4.37E+04 | HPV-pE7 | 1.70E+09 | 2.17E+07 |
| KRAS G12C | 1.82E+05 | 3.47E+08 | RB1 | 1.86E+09 | 2.95E+08 |
| VHL | 1.91E+05 | 1.58E+06 | FGFR1c domain D2 | 1.95E+09 | 1.55E+09 |
| PINK | 1.95E+05 | 3.31E+06 | Calprotectin (het-6) | 2.19E+09 | 1.45E+09 |
| PP2A-B | 2.00E+05 | 3.63E+06 | mCherry | 2.24E+09 | 8.32E+08 |
| OmpA (*ec*) | 2.00E+05 | 2.24E+06 | IsdB | 2.29E+09 | 3.09E+08 |
| OmpF (*ec*) | 2.04E+05 | 1.07E+08 | p65 | 2.34E+09 | 1.51E+09 |
| PP2A-B" | 2.09E+05 | 1.20E+09 | 14-3-3z | 2.51E+09 | 1.23E+09 |
| OTUBI | 2.09E+05 | 2.40E+07 | RAF | 2.82E+09 | 4.47E+08 |
| HSPB1 (structured) | 2.09E+05 | 4.68E+08 | Mdm2 | 2.88E+09 | 1.05E+09 |
| Alpha amylase | 2.14E+05 | 2.24E+05 | Calprotectin (s100a9) | 3.39E+09 | 8.13E+08 |
| TRIM | 2.29E+05 | 4.27E+06 | PAL | 5.13E+09 | 4.79E+08 |
| NIX | 2.29E+05 | 3.55E+06 | TCIM | 5.25E+09 | 8.71E+08 |
| Keap1 | 2.29E+05 | 3.98E+08 | Myc (DBD) | 5.37E+09 | 2.88E+08 |
| G3BP | 2.88E+05 | 3.24E+06 | FnbpA | 5.37E+09 | 6.76E+08 |
| p53 (AD) | 2.88E+05 | 1.95E+05 | PIK3CB | 6.03E+09 | 1.58E+09 |
| GTF2I | 3.39E+05 | 1.26E+05 | PDE6D | 8.91E+09 | 1.45E+09 |

**Note S1. Setting a threshold endpoint titer.**

**Setting the criteria for a hit based on titer, NGS, and AlphaFold predictions**

We sought to establish criteria based on titer, NGS, and AlphaFold predictions to identifying hits and reduce false positives. We used the no-fusion endpoint titers as a preliminary cutoff: any well with an endpoint titer >10x the no-fusion endpoint titer was considered a preliminary hit and sequenced (cutoffs of $1.5*10^6$ PFU/mL for affibody and $1.6*10^7$ PFU/mL for affitin), yielding 44 affibody and 56 affitin preliminary hits. We compiled the sequences that made up >1% of NGS reads and performed AlphaFold2 Multimer predictions for the top four hits (excluding truncations, **Supplementary Fig. 22**). We collected E. coli luciferase assay data for dominant variants from each of these preliminary hits that did not have a truncation dominate (>10% of NGS reads; **Supplementary Fig. 23**). The NGS generally matched what we observed in our initial 6-panel PPI-PANCS: an average of 4.5 variants with >1% of reads (range = 1-24), the most dominant sequence from the selection had on average 57% of reads (range = 3-96%), and convergence was a frequent occurrence with 58% of hits converging onto >50% on one variant; only 17% of hits had a dominant variant with < 20% of reads (**Supplementary Fig. 24**). We observed a background level (~0.1-1%) of truncation variants in the NGS results indicating incomplete de-enrichment and confirmed that a subset of selections below our threshold contained high levels of truncations (**Supplementary Fig. 25**). Additionally, we showed that with an additional day of passaging (2 rounds), these no-fusion and failed selections do continue to de-enrich to extinction (**Supplementary Fig. 26**). We conclude from this NGS data, that a lower cutoff would increase our false positive rate by including samples that had not de-enriched; however, with additional passaging, a much lower threshold could have been used as extinctions and successful enrichment become more separated.

Several hits had dominant variants that were truncations, low convergence, or variants that were enriched on multiple targets. We investigated each of these hits further by luciferase assay (**Supplementary Fig. 23**). Of the 143 variants that had >10% of reads within a hit, there were 12 truncations across 10 targets. Each of these targets had a hit in the other library; 8 out of 10 top variants on these targets did have binders (only Bax peptide and SasG did not, however, the truncated variant did bind SasG) within their hits suggesting that enrichment of a truncation as the top variant in one library was not a strong indication of false positives in the other library's hit. Of the 131 variants that had >10% of reads within a hit (excluding truncations), there were 126 unique variants – 5 variants enriched in two target selections. KRAS G12D and KRAS G12C enriched the same affitin variant, KRAS WT and KRAS G12V enriched the same affitin variant, KIX and NRAS enriched the same affibody variant, RB1 and UEV enriched the same affitin variant, and the dominant RAF affibody variant was also the 2nd most common affibody variant on BAX peptide. The common KRAS binding variants are unsurprising, but we subcloned the hits for these other 6 targets: UEV and KIX bound their top variants, RAF, NRAS, Bax peptide, and RB1 did not (**Supplementary Fig. 23**).

Lastly, we hoped to use AlphaFold2 or AlphaFold3 predictions to assist in identifying false positives. We saw a wide distribution of confidence in AlphaFold2 multimer prediction confidences for the top variant from each hit (ipTM of 0.11-0.92; **Supplementary Fig. 24**) Additionally, we observed with a panel of binders and targets, all tested in luciferase assay for binding (18x16), that AlphaFold3 was not able to identify the binding pairs (**Supplementary Fig. 27**). Lastly, we decided to compare the ipTM to in vitro Kd for all of our measured interactions and saw only a weak correlation (**Supplementary Fig. 32**). We conclude from these results that we could not use low ipTM values as a filter. Based on our examination of several features in the NGS data, ignoring the AlphaFold predictions, we conclude that a titer greater than our threshold (10x the No Fusion) and a dominant variant that is not a truncation are the criteria for identifying a hit; based on this we have 42 affibody hits and 46 affitin hits across 59 targets. Six target tested in luciferase (**Supplementary Fig. 23**) failed to have a binder (at least 2x lux signal) across both libraries – Bax peptide, NRAS, FGFR, TEV, PP2A-B'', and KRAS G12V.

**Supplementary Fig. 22 (below)**: **Analysis of NGS for hits from 96-target panel PANCS with affibody and affitin.** We performed NGS on each hit (see Methods) and show in the figures below each variant that had >1% of reads for a hit (organized as each affibody hit and then each affitin hit below). We then use AlphaFold2 multimer collab (AlphaFold2.ipynb - Colab (google.com)) for predicting the interaction between binder and target for the top 4 variants (above 1% of reads). Alphafold metrics: pLDDT (Average Predicted Local Distance Difference Test): Tests for local structural confidence - Very Low (<50), Low (60), OK (70), Confident (80), Very High (>90); pTM (Predicted Template Modeling Score): Measures structural congruency - pTM > 0.5 is generally recognized as strong enough to accept as inference; and ipTM (Interface Predicted Template Modeling Score): Similar to pTM but looking at multimer interactions - >0.8 indicates high confidence and less than <0.2 indicates no confidence in the predicted interaction interface. Predicted structure is omitted for predictions in which the pLDDT was <50 or the pTM was <0.5. Targets are shown in grey, binders are color coded based on the pie chart.

| Target | KRAS (G12D) |
|---|---|
| Scaffold | Affibody |
| Total HTPS Reads | 6198 |
| Unique HTPS Reads | 259 |
| Top Variant % of Total Reads | 86 |
| Final Phage Titer (PFU/mL) | 1.45E+08 |

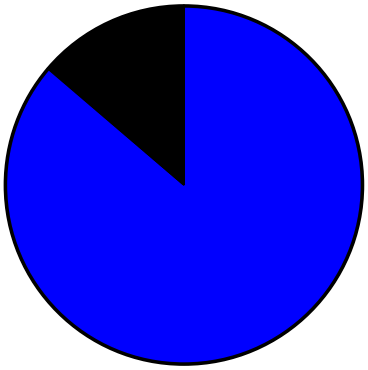

Library: VDNKF?KE???A???I??LPNLN??Q??AF??SL??DPSQSANLLAEAKKLNDAQAPK

86.24% VDNKFNKEVCDAYAQIGNLPNLNVYQIVAFIRSLDNDPSQSANLLAEAKKLNDAQAPK

13.76% Other Sequence Less than 1% Total Reads

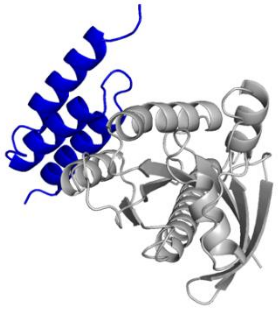

pLDDT = 88.2

pTM = 0.854

ipTM = 0.844

| Target | ZB |
|---|---|
| Scaffold | Affibody |
| Total HTPS Reads | 4677 |
| Unique HTPS Reads | 3668 |
| Top Variant % of Total Reads | 78 |
| Final Phage Titer (PFU/mL) | 1.74E+07 |

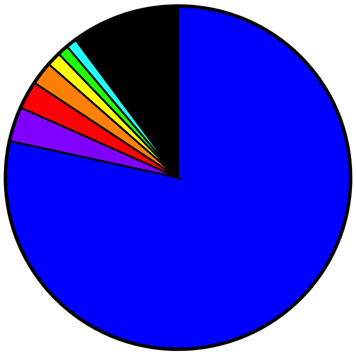

```
Library: VDNKF?KE???A???I??LPNLN??Q??AF??SL??DPSQSANLLAEAKKLNDAQAPK
78.43%   VDNKFTKEVRSAAHQIIYLPNLNLHQQCAFLDSLTNDPSQSANLLAEAKKLNDAQAPK
3.27%    VDNKFYKEVISAAREIAGLPNLNVGQRSAFIDSLIDDPSQSANLLAEAKKLNDAQAPK
2.63%    VDNKFTKEVATAVCQICCLPNLNVDQMNAFISSLFDDPSQSANLLAEAKKLNDAQAPK
2.18%    VDNKFTKEWCSAYVEIGVLPNLNDGQKNAFLYSLVDDPSQSANLLAEAKKLNDAQAPK
1.33%    VDNKFSKEVDNAINEIGDLPNLNVYQIIAFVRSLSDDPSQSANLLAEAKKLNDAQAPK
1.03%    VDNKFTKEVDDASFEILYLPNLNDFQENAFLVSLSDDPSQSANLLAEAKKLNDAQAPK
1.00%    VDNKFTKELVYATNEIYNLPNLNRLQRRAFLDSLSNDPSQSANLLAEAKKLNDAQAPK
10.13%   Other Sequence less than 1%
```

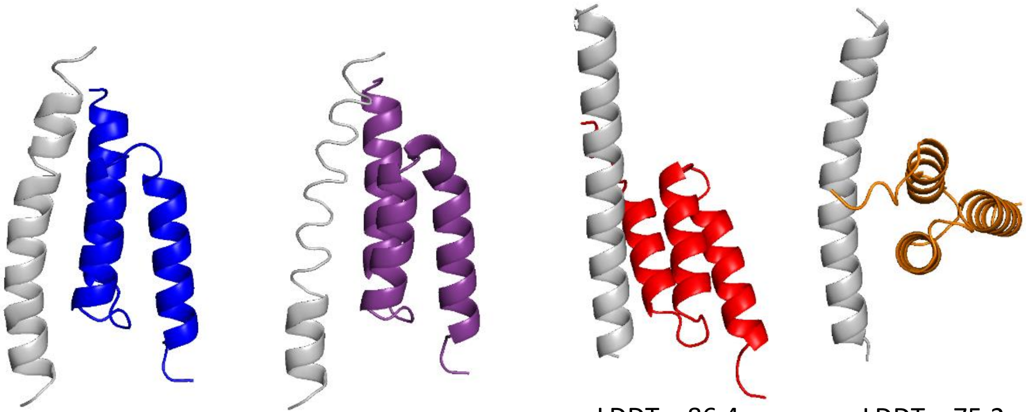

| | | | |
|---|---|---|---|
| pLDDT = 73.4 | pLDDT = 73.5 | pLDDT = 86.4 | pLDDT = 75.2 |
| pTM = 0.780 | pTM = 0.834 | pTM = 0.726 | pTM = 0.599 |
| ipTM = 0.691 | ipTM = 0.793 | ipTM = 0.523 | ipTM = 0.137 |

| Target | p65 (17-293) |
|---|---|
| Scaffold | Affibody |
| Total HTPS Reads | 7057 |
| Unique HTPS Reads | 226 |
| Top Variant % of Total Reads | 90 |
| Final Phage Titer (PFU/mL) | 2.34E+09 |

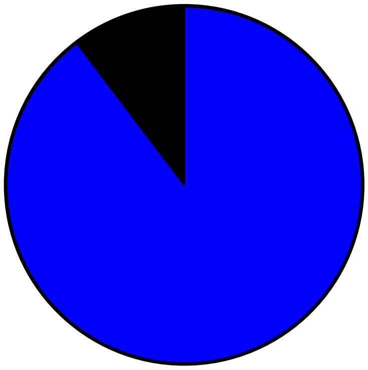

**Library: VDNKF?KE???A???I??LPNLN??Q??AF??SL??DPSQSANLLAEAKKLNDAQAPK**

**89.66% VDNKFTKESFI---EIVNLPNLNLRQIDAFVSSLFDDPSQSANLLAEAKKLNDAQAPK**

**10.34% Other Sequence Less than 1% Total Reads**

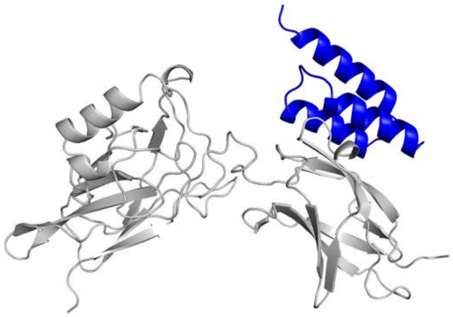

pLDDT = 88.2
pTM = 0.854
ipTM = 0.844

| Target | HPV-pE7 |
|---|---|
| Scaffold | Affibody |
| Total HTPS Reads | 22667 |
| Unique HTPS Reads | 687 |
| Top Variant % of Total Reads | 56 |
| Final Phage Titer (PFU/mL) | 1.70E+09 |

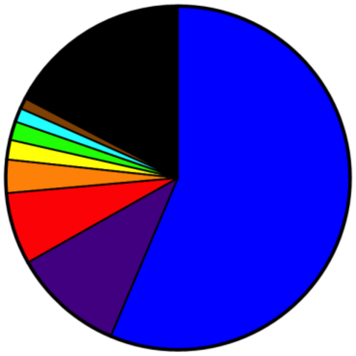

| | |
|---|---|
| Library: | VDNKF?KE???A???I??LPNLN??Q??AF??SL??DPSQSANLLAEAKKLNDAQAPK |
| 56.38% | VDNKFSKEWTYAAFQIPSLPNLNRNQRRAFIFSLGDDPSQSANLLAEAKKLNDAQAPK |
| 10.44% | VDNKFSKEVVDAVVQISALPNLNRDQRPAFVISLVDDPSQSANLLAEAKKLNDAQAPK |
| 6.76% | VDNKFNKEVRYAVVEIPRLPNLNPTQGRAFVFSLGDDPSQSANLLAEAKKLNDAQAPK |
| 3.14% | VDNKFSKELCYAVCEIDPLPNLNPSQGRAFLISLVDDPSQSANLLAEAKKLNDAQAPK |
| 1.85% | VDNKFYKEVDAAASEICSLPNLNRNQRSAFICSLYNDPSQSANLLAEAKKLNDAQAPK |
| 1.76% | VDNKFSKEWTYAAFQIPSLPNLNRDQRPAFVISLVDDPSQSANLLAEAKKLNDAQAPK |
| 1.31% | VDNKFSKEVVDAVVQISALPNLNRNQRRAFIFSLGDDPSQSANLLAEAKKLNDAQAPK |
| 1.02% | VDNKFSKEWTYAAFQIPSLPNLNRNQRRAFIFSLGDDPSQSANLLAEAKKLNDAQAPK |
| 17.34% | Other Sequence Less Than 1% Total Reads |

| | | | |
|---|---|---|---|
| **Insufficient Confidence to Predict Complex Structure (pTM)** | **Insufficient Confidence to Predict Complex Structure (pTM)** | **Insufficient Confidence to Predict Complex Structure (pTM)** | **Insufficient Confidence to Predict Complex Structure (pTM)** |
| pLDDT = 52.2<br>pTM = 0.411<br>ipTM = 0.256 | pLDDT = 57.1<br>pTM = 0.365<br>ipTM = 0.154 | pLDDT = 51.8<br>pTM = 0.366<br>ipTM = 0.209 | pLDDT = 50.2<br>pTM = 0.395<br>ipTM = 0.260 |

| | |
|---|---|
| Target | FGFR1c domain D2 |
| Scaffold | Affibody |
| Total HTPS Reads | 8101 |
| Unique HTPS Reads | 327 |
| Top Variant % of Total Reads | 86 |
| Final Phage Titer (PFU/mL) | 1.95E+09 |

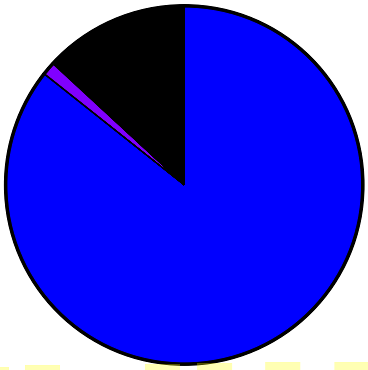

**Library:** VDNKF?KE???A???I??LPNLN??Q??AF??SL??DPSQSANLLAEAKKLNDAQAPK

**85.66%** VDNKFTKEVATAVCQICCLPNLNVDQMNAFISSLFDDPSQSANLLAEAKKLNDAQAPK

**1.21%** VDNKFSKEVVYAVYQITGLPNLNLVQKFAFVTSLCDDPSQSANLLAEAKKLNDAQAPK

**13.13%** Other Sequence Less than 1% Total Reads

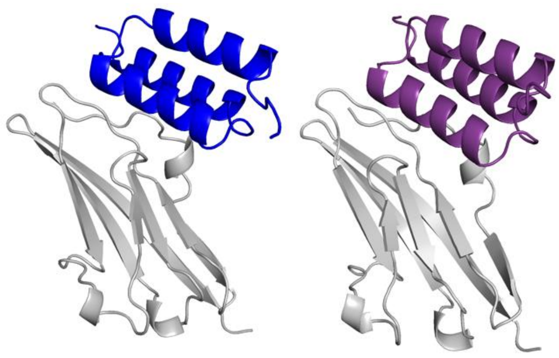

pLDDT = 80.1
pTM = 0.623
ipTM = 0.151

pLDDT = 80.7
pTM = 0.606
ipTM = 0.130

| Target | TCIM |
|---|---|
| Scaffold | Affibody |
| Total HTPS Reads | 9826 |
| Unique HTPS Reads | 8861 |
| Top Variant % of Total Reads | 90 |
| Final Phage Titer (PFU/mL) | 5.25E+09 |

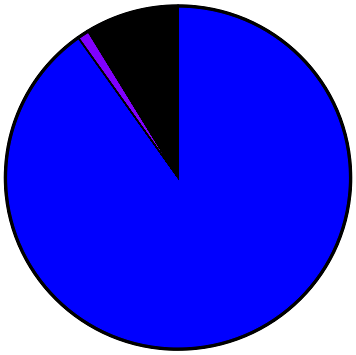

```
  Library: VDNKF?KE???A???I??LPNLN??Q??AF??SL??DPSQSANLLAEAKKLNDAQAPK
  90.18%   VDNKFTKEWCSAYVEIGVLPNLNDGQKNAFLYSLVDDPSQSANLLAEAKKLNDAQAPK
  1.09%    VDNKFTKEWTDAFCEISCLPNLNAHQKVAFLGSLLDDPSQSANLLAEAKKLNDAQAPK
  8.73%   Other Sequence less than 1%
```

| Insufficient Confidence to Predict Complex Structure (pTM) | Insufficient Confidence to Predict Complex Structure (pTM) |
|---|---|
| pLDDT = 53.0 | pLDDT = 52.4 |
| pTM = 0.387 | pTM = 0.430 |
| ipTM = 0.174 | ipTM = 0.256 |

| | |
|---|---|
| Target | PDE6D |
| Scaffold | Affibody |
| Total HTPS Reads | 7257 |
| Unique HTPS Reads | 4982 |
| Top Variant % of Total Reads | 69 |
| Final Phage Titer (PFU/mL) | 8.91E+09 |

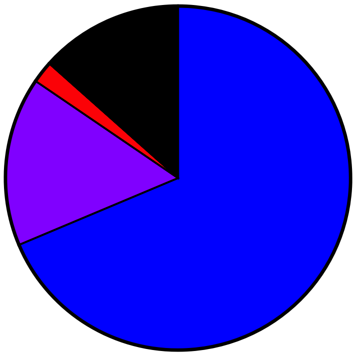

**Library:** VDNKF?KE???A???I??LPNLN??Q??AF??SL??DPSQSANLLAEAKKLNDAQAPK

68.65% VDNKFYKELLDAVRQIRSLPNLNVDQKAAFIFSLDNDPSQSANLLAEAKKLNDAQAPK

15.89% VDNKFYKELFYAVRQINGLPNLNRSQRYAFICSLDDDPSQSANLLAEAKKLNDAQAPK

2.05% VDNKFTKEWCSAYVEIGVLPNLNDGQKNAFLYSLVDDPSQSANLLAEAKKLNDAQAPK

13.41% Other Sequence less than 1%

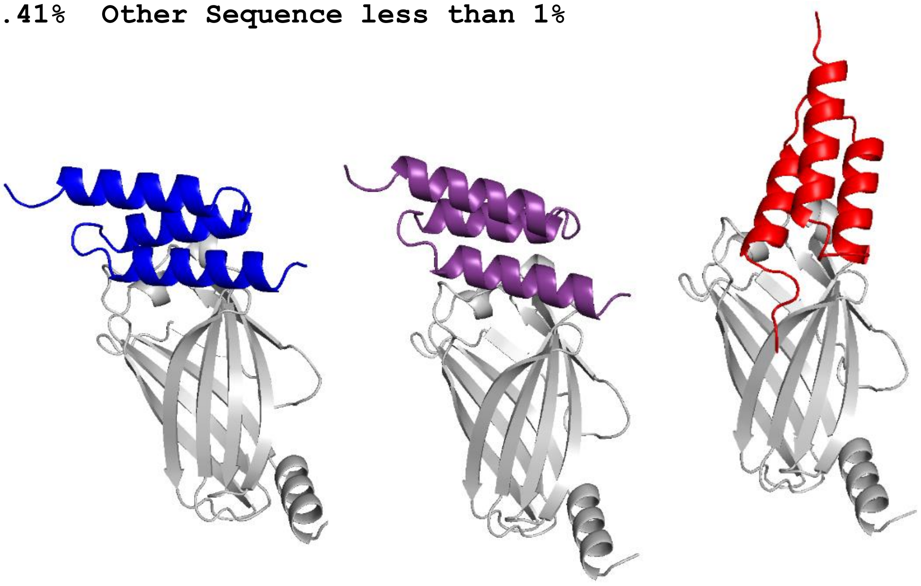

pLDDT = 84.9
pTM = 0.701
ipTM = 0.177

pLDDT = 86.1
pTM = 0.728
ipTM = 0.270

pLDDT = 87.8
pTM = 0.732
ipTM = 0.279

| Target | RAF RBD |
|---|---|
| Scaffold | Affibody |
| Total HTPS Reads | 1997 |
| Unique HTPS Reads | 354 |
| Top Variant % of Total Reads | 18 |
| Final Phage Titer (PFU/mL) | 2.82E+09 |

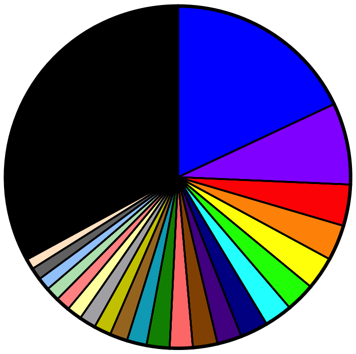

| | |
|---|---|
| Library: | VDNKF?KE???A???I??LPNLN??Q??AF??SL??DPSQSANLLAEAKKLNDAQAPK |
| 17.98% | VDNKFNKEGDYAVFQICGLPNLNEGQGAAFVGSLHNDPSQSANLLAEAKKLNDAQAPK |
| 7.71% | VDNKFYKEVAAASGEIRDLPNLNRDQGDAFIGSLVNDPSQSANLLAEAKKLNDAQAPK |
| 3.91% | VDNKFNKEGADAVDEIHVLPNLNRRQRVAFLGSLHDDPSQSANLLAEAKKLNDAQAPK |
| 3.36% | VDNKFYKEWVTAYNEICLLPNLNGLQICAFIGSLDDDPSQSANLLAEAKKLNDAQAPK |
| 3.10% | VDNKFTKEVATAVCQICCLPNLNVDQMNAFISSLFDDPSQSANLLAEAKKLNDAQAPK |
| 2.75% | VDNKFTKELRDANLQIGGLPNLNGTQVSAFLDSLRDDPSQSANLLAEAKKLNDAQAPK |
| 2.60% | VDNKFTKESFI-EIV--NLPNLNLRQIDAFVSSLFDDPSQSANLLAEAKKLNDAQAPK |
| 2.30% | VDNKFNKEWSYAFYEIVFLPNLNECQVDAFISSLGDDPSQSANLLAEAKKLNDAQAPK |
| 2.60% | VDNKFTKEWCSAFCEICLLPNLNLSQMHAFVSSLLNDPSQSANLLAEAKKLNDAQAPK |
| 2.30% | VDNKFSKEVVYAVYQITGLPNLNLVQKFAFVTSLCDDPSQSANLLAEAKKLNDAQAPK |
| 2.15% | VDNKFNKEGDTAVLEIGHLPNLNLHQQYAFLHSLYNDPSQSANLLAEAKKLNDAQAPK |
| 2.15% | VDNKFTKEWDNAVNQIANLPNLNPAQKIAFISSLSDDPSQSANLLAEAKKLNDAQAPK |
| 1.85% | VDNKFNKEVADANSEIRYLPNLNLGQRLAFLGSLYNDPSQSANLLAEAKKLNDAQAPK |
| 1.65% | VDNKFTKELGAADDEILRLPNLNGNQRCAFVVSLFDDPSQSANLLAEAKKLNDAQAPK |
| 1.60% | VDNKFYKEWSTAICEIVGLPNLNLSQQHAFIFSLLDDPSQSANLLAEAKKLNDAQAPK |
| 1.55% | VDNKFSKEGCSADREIGGLPNLNLRQQDAFIGSLYNDPSQSANLLAEAKKLNDAQAPK |
| 1.40% | VDNKFNKEVCDAYAQIGNLPNLNVYQIVAFIRSLDNDPSQSANLLAEAKKLNDAQAPK |
| 1.35% | VDNKFTKEGCNAVIQINSLPNLNEAQRRAFICSLANDPSQSANLLAEAKKLNDAQAPK |
| 1.30% | VDNKFSKEVPSAFDEIGALPNLNVRQQNAFIVSLPDDPSQSANLLAEAKKLNDAQAPK |
| 1.20% | VDNKFSKELRDATCEIGSLPNLNRGQGNAFVTSLRDDPSQSANLLAEAKKLNDAQAPK |
| 1.15% | VDNKFSKEVGAADGEIVGLPNLNGSQKIAFLRSLVNDPSQSANLLAEAKKLNDAQAPK |
| 1.00% | VDNKFSKEVFTAFDQIGGLPNLNLCQKDAFICSLVDDPSQSANLLAEAKKLNDAQAPK |
| 33.00% | Other Sequence less than 1% |

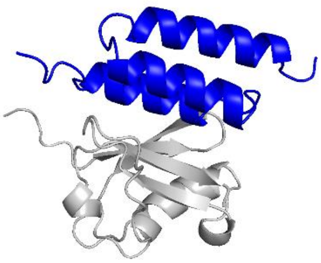

pLDDT = 75.9
pTM = 0.573
ipTM = 0.254

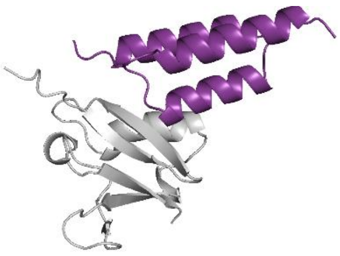

pLDDT = 75.9
pTM = 0.552
ipTM = 0.159

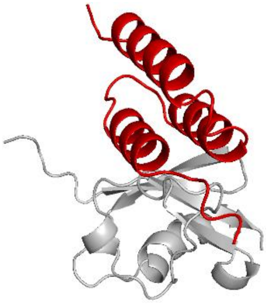

pLDDT = 80.2
pTM = 0.572
ipTM = 0.242

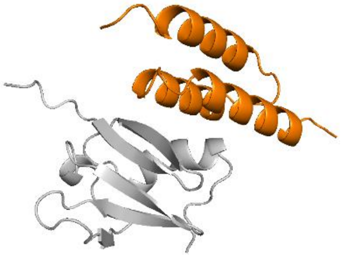

pLDDT = 83.0
pTM = 0.637
ipTM = 0.385

| Target | NRAS |
|---|---|
| Scaffold | Affibody |
| Total HTPS Reads | 834 |
| Unique HTPS Reads | 310 |
| Top Variant % of Total Reads | 19 |
| Final Phage Titer (PFU/mL) | 9.12E+08 |

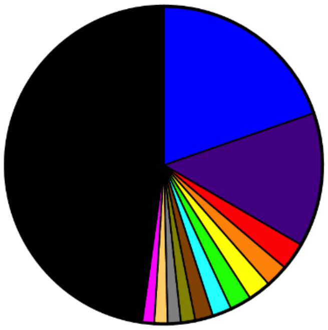

```
Library: VDNKF?KE???A???I??LPNLN??Q??AF??SL??DPSQSANLLAEAKKLNDAQAPK
19.68%   VDNKFTKEWCSAFCEICLLPNLNLSQMHAFVSSLLNDPSQSANLLAEAKKLNDAQAPK
13.69%   VDNKFYKEGTYASYEINNLPNLNLVQRSAFVFSLYNDPSQSANLLAEAKKLNDAQAPK
2.81%    VDNKFSKEWVDASCQIVSLPNLNLVQRSAFVFSLYNDPSQSANLLAEAKKLNDAQAPK
2.32%    VDNKFYKEWNNAADEILNLPNLNEVQGCAFVCSLNDDPSQSANLLAEAKKLNDAQAPK
2.32%    VDNKFSKEWTYAAFQIPSLPNLNRNQRRAFIFSLGDDPSQSANLLAEAKKLNDAQAPK
2.20%    VDNKFSKEVHNAVGEIVILPNLNASQVHAFIVSLGNDPSQSANLLAEAKKLNDAQAPK
1.96%    VDNKFSKEWFDATCEIFCLPNLNGVQKRAFLGSLDDDPSQSANLLAEAKKLNDAQAPK
1.83%    VDNKFSKEWHSAYAEIAHLPNLNPSQVCAFIGSLLDDPSQSANLLAEAKKLNDAQAPK
1.47%    VDNKFNKGRGFCKSRDSCPAELKPGPATRLRQQPGRDPSQSANLLAEAKKLNDAQAPK
1.34%    VDNKFNKEGGCCKL*DSPPAELKRVPGVCLHMQPD*DPSQSANLLAEAKKLNDAQAPK
1.34%    VDNKFYKEWFDANVQIAVLPNLNEYQVIAFISSLANDPSQSANLLAEAKKLNDAQAPK
1.22%    VDNKFYKEWCNANYEISCLPNLNDYQKNAFIVSLGNDPSQSANLLAEAKKLNDAQAPK
47.80%   Other Sequence Less Than 1% Total Reads
```

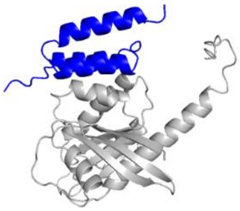

pLDDT = 88.5
pTM = 0.702
ipTM = 0.287

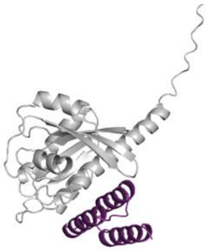

pLDDT = 77.9
pTM = 0.702
ipTM = 0.213

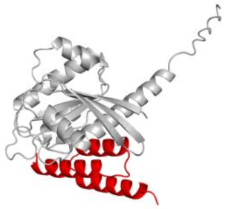

pLDDT = 75.8
pTM = 0.688
ipTM = 0.175

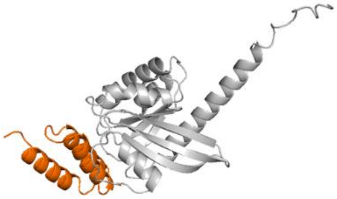

pLDDT = 78.8
pTM = 0.686
ipTM = 0.173

| Target | Beclin |
|---|---|
| Scaffold | Affibody |
| Total HTPS Reads | 11794 |
| Unique HTPS Reads | 5545 |
| Top Variant % of Total Reads | 47 |
| Final Phage Titer (PFU/mL) | 5.00E+08 |

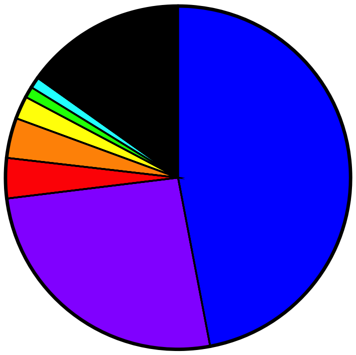

| | | |
|---|---|---|
| Library: | | VDNKF?KE???A???I??LPNLN??Q??AF??SL??DPSQSANLLAEAKKLNDAQAPK |
| ■ (blue) | 47.02% | VDNKFYKEVISAICQIFGLPNLNGTQRRAFLCSLGNDPSQSANLLAEAKKLNDAQAPK |
| ■ (purple) | 26.04% | VDNKFNKEWTAAFRQIVGLPNLNRSQRCAFLGSLLDDPSQSANLLAEAKKLNDAQAPK |
| ■ (red) | 3.79% | VDNKFYKELADAITEIVNLPNLNRAQRCAFLGSLADDPSQSANLLAEAKKLNDAQAPK |
| ■ (orange) | 3.78% | VDNKFSKEWANAFRQIHDLPNLNRCQRCAFLGSLLDDPSQSANLLAEAKKLNDAQAPK |
| ■ (yellow) | 2.16% | VDNKFNKEWTAAFRQIVGLPNLNGTQRRAFLCSLGNDPSQSANLLAEAKKLNDAQAPK |
| ■ (green) | 1.04% | VDNKFNKEWTAAFRQIVGLPNLNRSQRCAFLCSLGNDPSQSANLLAEAKKLNDAQAPK |
| ■ (cyan) | 1.04% | VDNKFTKEVDSAVFEIHILPNLNDYQRVAFIFSLFDDPSQSANLLAEAKKLNDAQAPK |
| ■ (black) | 15.13% | Other Sequence less than 1% |

**Insufficient Confidence to Predict Complex Structure (pTM)**

pLDDT = 66.8
pTM = 0.472
ipTM = 0.288

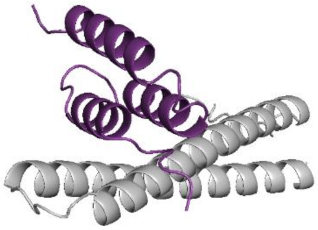

pLDDT = 78.8
pTM = 0.604
ipTM = 0.476

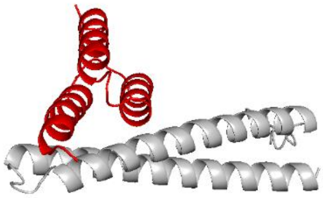

pLDDT = 79.2
pTM = 0.648
ipTM = 0.553

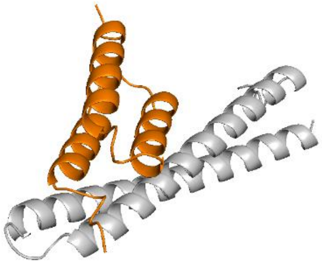

pLDDT = 82.6
pTM = 0.714
ipTM = 0.707

| Target | MDM2 |
|---|---|
| Scaffold | Affibody |
| Total HTPS Reads | 9656 |
| Unique HTPS Reads | 4130 |
| Top Variant % of Total Reads | 5 |
| Final Phage Titer (PFU/mL) | 2.88E+09 |

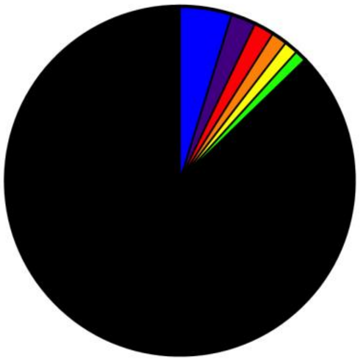

```
Library: VDNKF?KE???A???I??LPNLN??Q??AF??SL??DPSQSANLLAEAKKLNDAQAPK
4.82%    VDNKFYKEWNNAADEILNLPNLNEVQGCAFVCSLNDDPSQSANLLAEAKKLNDAQAPK
2.31%    VDNKFSKEWDSASREILLLPNLNGLQQYAFVYSLVDDPSQSANLLAEAKKLNDAQAPK
1.86%    VDNKFSKEWINASYEILSLPNLNESQEHAFLSSLGDDPSQSANLLAEAKKLNDAQAPK
1.45%    VDNKFSKEWLAATRQIIRLPNLNGIQECAFIRSLPDDPSQSANLLAEAKKLNDAQAPK
1.27%    VDNKFTKEWCSAFCEICLLPNLNLSQMHAFVSSLLNDPSQSANLLAEAKKLNDAQAPK
1.06%    VDNKFYKEGTYASYEINNLPNLNLVQRSAFVFSLYNDPSQSANLLAEAKKLNDAQAPK
87.23%   Other Sequence Less Than 1% Total Reads
```

**Insufficient Confidence to Predict Complex Structure (pTM)**

pLDDT = 60.0
pTM = 0.480
ipTM = 0.466

**Insufficient Confidence to Predict Complex Structure (pTM)**

pLDDT = 59.5
pTM = 0.478
ipTM = 0.459

**Insufficient Confidence to Predict Complex Structure (pTM)**

pLDDT = 61.4
pTM = 0.473
ipTM = 0.446

**Insufficient Confidence to Predict Complex Structure (pTM)**

pLDDT = 55.1
pTM = 0.462
ipTM = 0.371

| Target | Calpro het 12aa |
|---|---|
| Scaffold | Affibody |
| Total HTPS Reads | 3260 |
| Unique HTPS Reads | 382 |
| Top Variant % of Total Reads | 26 |
| Final Phage Titer (PFU/mL) | 2.45E+08 |

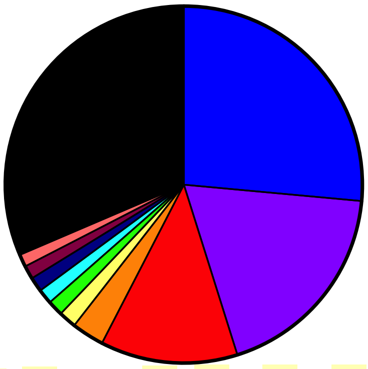

| | |
|---|---|
| **Library:** | **VDNKF?KE???A???I??LPNLN??Q??AF??SL??DPSQSANLLAEAKKLNDAQAPK** |
| **26.47%** | **VDNKFSKELIDADSEIYALPNLNELQEPAFVGSLGNDPSQSANLLAEAKKLNDAQAPK** |
| **18.68%** | **VDNKFNKELANAISEIASLPNLNRGQRDAFVGSLGNDPSQSANLLAEAKKLNDAQAPK** |
| **12.42%** | **VDNKFYKEVNAAIAQIYGLPNLNLDQEYAFVFSLLNDPSQSANLLAEAKKLNDAQAPK** |
| **2.94%** | **VDNKFYKEWDNAYSEIDVLPNLNLDQGCAFLVSLRDDPSQSANLLAEAKKLNDAQAPK** |
| **1.60%** | **VDNKFSKELTYAINEIHSLPNLNRTQGFAFVRSLIDDPSQSANLLAEAKKLNDAQAPK** |
| **1.41%** | **VDNKFSKEWCNADLQIGDLPNLNRCQQGAFICSLGDDPSQSANLLAEAKKLNDAQAPK** |
| **1.35%** | **VDNKFTKEVATAVCQICCLPNLNVDQMNAFISSLFDDPSQSANLLAEAKKLNDAQAPK** |
| **1.32%** | **VDNKFSKEVVYAVYQITGLPNLNLVQKFAFVTSLCDDPSQSANLLAEAKKLNDAQAPK** |
| **1.23%** | **VDNKFNKEVCDAYAQIGNLPNLNVYQIVAFIRSLDNDPSQSANLLAEAKKLNDAQAPK** |
| **1.13%** | **VDNKFNKEVADANSEIRYLPNLNLGQRLAFLGSLYNDPSQSANLLAEAKKLNDAQAPK** |
| **31.44%** | **Other Sequence Less than 1% Total Reads** |

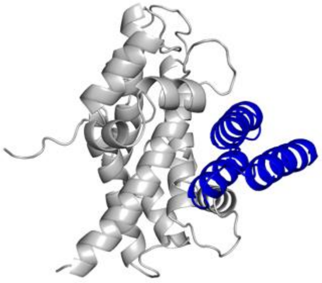

pLDDT = 71.6
pTM = 0.721
ipTM = 0.372

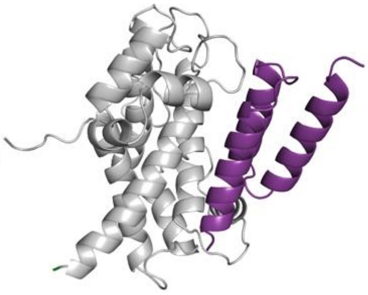

pLDDT = 76.6
pTM = 0.745
ipTM = 0.342

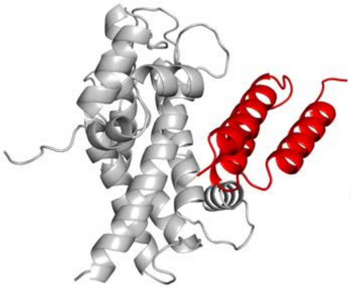

pLDDT = 79.8
pTM = 0.771
ipTM = 0.41

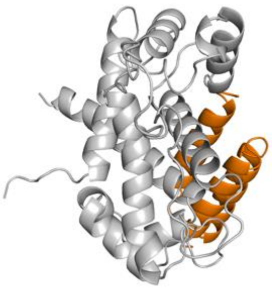

pLDDT = 74.2
pTM = 0.712
ipTM = 0.248

| Target | HlgA toxin |
|---|---|
| Scaffold | Affibody |
| Total HTPS Reads | 6627 |
| Unique HTPS Reads | 257 |
| Top Variant % of Total Reads | 86 |
| Final Phage Titer (PFU/mL) | 1.10E+07 |

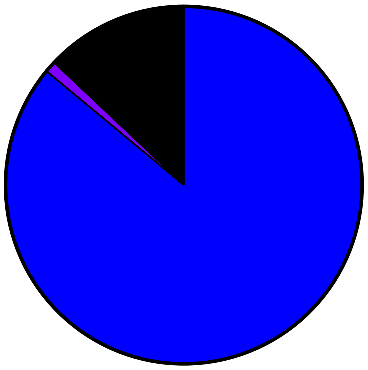

**Library: VDNKF?KE???A???I??LPNLN??Q??AF??SL??DPSQSANLLAEAKKLNDAQAPK**

**86.01% VDNKFYKEWSTAICEIVGLPNLNLSQQHAFIFSLLDDPSQSANLLAEAKKLNDAQAPK**

**1.01% VDNKFTKEWCSAFCEICLLPNLNLSQMHAFVSSLLNDPSQSANLLAEAKKLNDAQAPK**

**12.98% Other Sequence Less than 1% Total Reads**

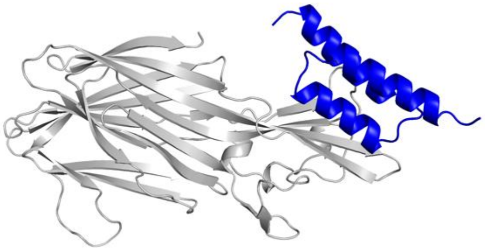

pLDDT = 87.4
pTM = 0.813
ipTM = 0.251

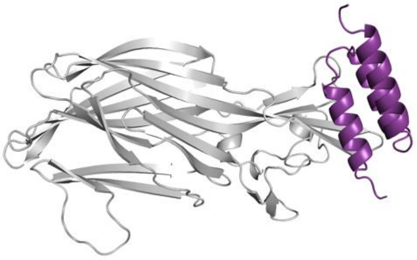

pLDDT = 88.0
pTM = 0.825
ipTM = 0.385

| | |
|---|---|
| Target | Bcl2 |
| Scaffold | Affibody |
| Total HTPS Reads | 6236 |
| Unique HTPS Reads | 271 |
| Top Variant % of Total Reads | 85 |
| Final Phage Titer (PFU/mL) | 1.51E+09 |

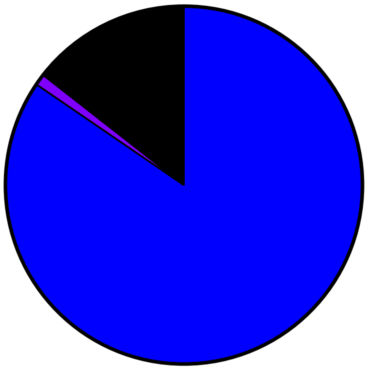

| | |
|---|---|
| Library: | VDNKF?KE???A???I??LPNLN??Q??AF??SL??DPSQSANLLAEAKKLNDAQAPK |
| 84.51% | VDNKFNKEVADANSEIRYLPNLNLGQRLAFLGSLYNDPSQSANLLAEAKKLNDAQAPK |
| 1.03% | VDNKFTKEWCSAFCEICLLPNLNLSQMHAFVSSLLNDPSQSANLLAEAKKLNDAQAPK |
| 14.46% | Other Sequence Less than 1% Total Reads |

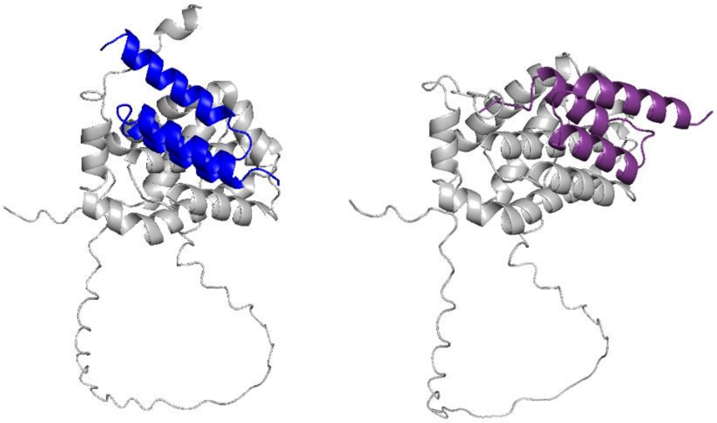

pLDDT = 77.7
pTM = 0.748
ipTM = 0.924

pLDDT = 64.9
pTM = 0.600
ipTM = 0.302

| Target | Calprotectin monomer s100a9 |
|---|---|
| Scaffold | Affibody |
| Total HTPS Reads | 5957 |
| Unique HTPS Reads | 778 |
| Top Variant % of Total Reads | 40 |
| Final Phage Titer (PFU/mL) | 3.39E+09 |

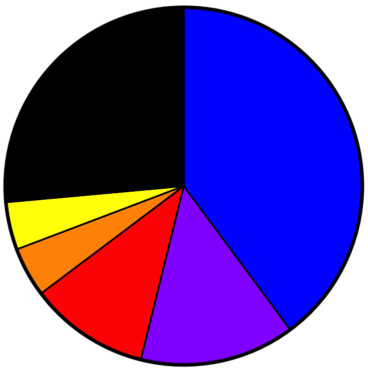

```
Library: VDNKF?KE???A???I??LPNLN??Q??AF??SL??DPSQSANLLAEAKKLNDAQAPK
39.84%   VDNKFYKELFNAAYEIYNLPNLNRVQVVAFIISLFDDPSQSANLLAEAKKLNDAQAPK
14.04%   VDNKFYKELVNASCQIYFLPNLNAIQVSAFICSLFDDPSQSANLLAEAKKLNDAQAPK
10.87%    VDNKFYKEVGYAAFQIYNLPNLNVCQRCAFVISLFDDPSQSANLLAEAKKLNDAQAPK
4.50%    VDNKFTKEWCDASVQIYILPNLNRAQQCAFLCSLSDDPSQSANLLAEAKKLNDAQAPK
4.33%    VDNKFTKEGVAAIREIYYLPNLNLYQQAAFVFSLNNDPSQSANLLAEAKKLNDAQAPK
26.43%   Other Sequence Less than 1% Total Reads
```

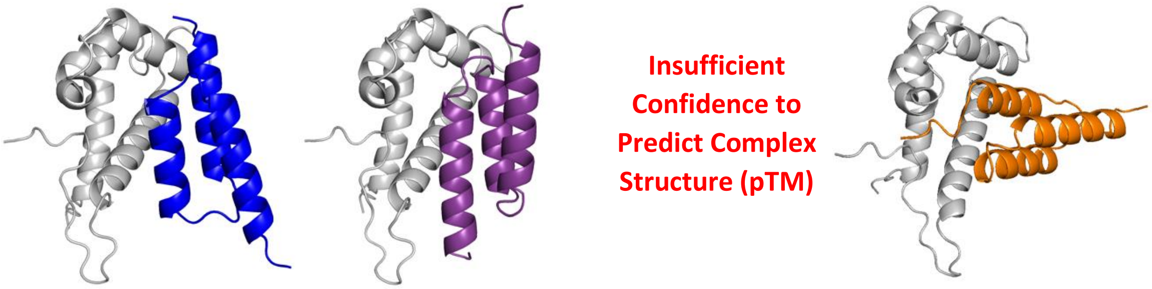


| | | | |
|---|---|---|---|
| pLDDT = 73.6 | pLDDT = 74.2 | pLDDT = 63.3 | pLDDT = 66.0 |
| pTM = 0.567 | pTM = 0.574 | pTM = 0.477 | pTM = 0.515 |
| ipTM = 0.161 | ipTM = 0.192 | ipTM = 0.197 | ipTM = 0.190 |

| Target | KIX Domain of CBP |
|---|---|
| Scaffold | Affibody |
| Total HTPS Reads | 8219 |
| Unique HTPS Reads | 495 |
| Top Variant % of Total Reads | 68 |
| Final Phage Titer (PFU/mL) | 1.48E+09 |

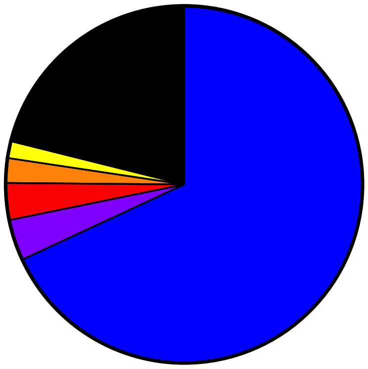

| | |
|---|---|
| Library: | VDNKF?KE???A???I??LPNLN??Q??AF??SL??DPSQSANLLAEAKKLNDAQAPK |
| 68.09% | VDNKFTKEWCSAFCEICLLPNLNLSQMHAFVSSLLNDPSQSANLLAEAKKLNDAQAPK |
| 3.71% | VDNKFSKEWIDAFVEINFLPNLNRNQRCAFVNSLLNDPSQSANLLAEAKKLNDAQAPK |
| 3.33% | VDNKFYKEWAAASSQIVDLPNLNVDQGLAFLTSLYDDPSQSANLLAEAKKLNDAQAPK |
| 2.24% | VDNKFSKEWFDATCEIFCLPNLNGVQKRAFLGSLDDDPSQSANLLAEAKKLNDAQAPK |
| 1.52% | VDNKFSKEWVTAFVEIASLPNLNQRQVRAFVGSLLNDPSQSANLLAEAKKLNDAQAPK |
| 21.11% | Other Sequence Less than 1% Total Reads |

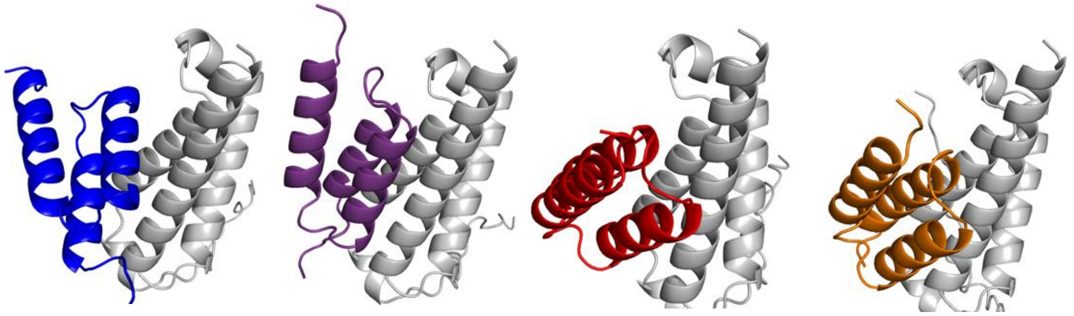

| | | | |
|---|---|---|---|
| pLDDT = 78.2 | pLDDT = 76.2 | pLDDT = 69.5 | pLDDT = 78.6 |
| pTM = 0.688 | pTM = 0.646 | pTM = 0.543 | pTM = 0.690 |
| ipTM = 0.708 | ipTM = 0.592 | ipTM = 0.318 | ipTM = 0.687 |

| Target | HRAS |
|---|---|
| Scaffold | Affibody |
| Total HTPS Reads | 9141 |
| Unique HTPS Reads | 8157 |
| Top Variant % of Total Reads | 89 |
| Final Phage Titer (PFU/mL) | 1.17E+09 |

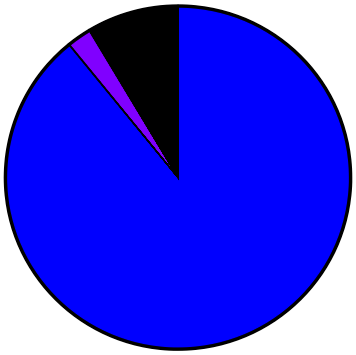

```
Library: VDNKF?KE???A???I??LPNLN??Q??AF??SL??DPSQSANLLAEAKKLNDAQAPK
89.13%   VDNKFSKEVDNAINEIGDLPNLNVYQIIAFVRSLSDDPSQSANLLAEAKKLNDAQAPK
2.29%    VDNKFTKEWCSAYVEIGVLPNLNDGQKNAFLYSLVDDPSQSANLLAEAKKLNDAQAPK
8.59%    Other Sequence less than 1%
```

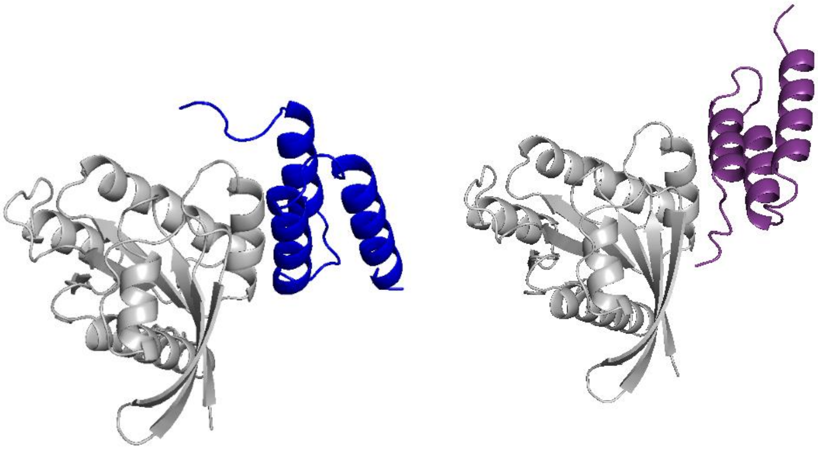

pLDDT = 90.1
pTM = 0.851
ipTM = 0.769

pLDDT = 86.1
pTM = 0.763
ipTM = 0.411

| Target | Calprotectin monomer s100a8 |
|---|---|
| Scaffold | Affibody |
| Total HTPS Reads | 8609 |
| Unique HTPS Reads | 305 |
| Top Variant % of Total Reads | 85 |
| Final Phage Titer (PFU/mL) | 3.63E+08 |

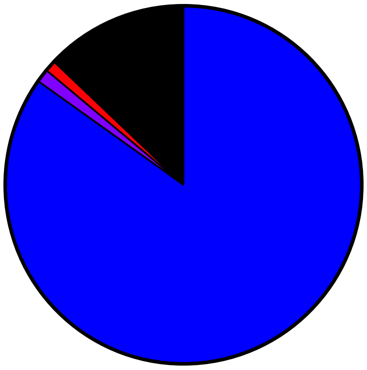

```
Library: VDNKF?KE???A???I??LPNLN??Q??AF??SL??DPSQSANLLAEAKKLNDAQAPK
84.83%   VDNKFSKEVVYAVYQITGLPNLNLVQKFAFVTSLCDDPSQSANLLAEAKKLNDAQAPK
1.21%    VDNKFTKEVATAVCQICCLPNLNVDQMNAFISSLFDDPSQSANLLAEAKKLNDAQAPK
1.02%    VDNKFTKELSAAVFQITSLPNLNVYQKSAFVFSLLDDPSQSANLLAEAKKLNDAQAPK
12.94%   Other Sequence Less than 1% Total Reads
```

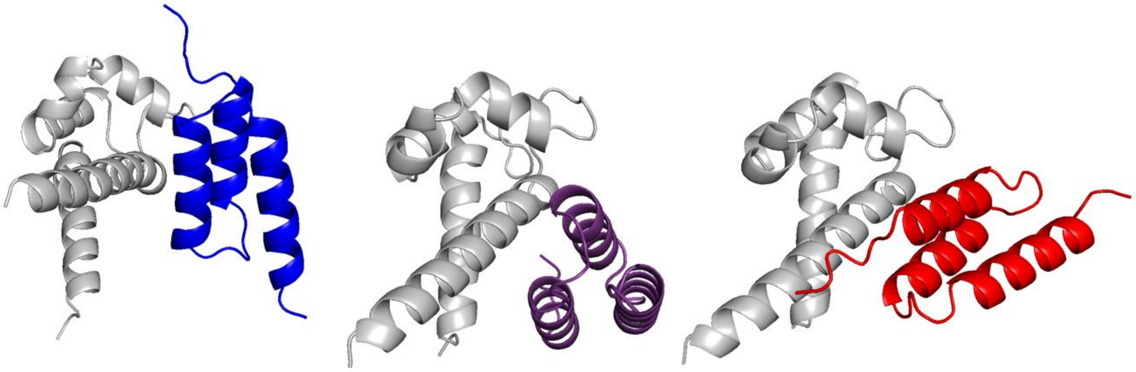

pLDDT = 78.6
pTM = 0.599
ipTM = 0.226

pLDDT = 77.9
pTM = 0.626
ipTM = 0.417

pLDDT = 76,1
pTM = 0.567
ipTM = 0.264

| | |
|---|---|
| Target | mCherry |
| Scaffold | Affibody |
| Total HTPS Reads | 6081 |
| Unique HTPS Reads | 414 |
| Top Variant % of Total Reads | 35 |
| Final Phage Titer (PFU/mL) | 2.24E+09 |

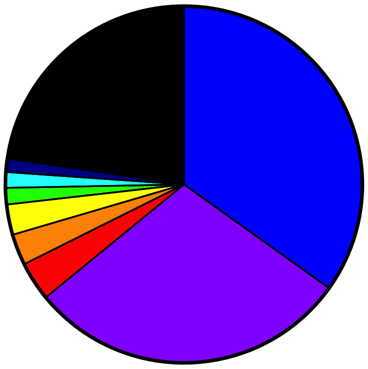

```
Library: VDNKF?KE???A???I??LPNLN??Q??AF??SL??DPSQSANLLAEAKKLNDAQAPK
34.88%   VDNKFSKEVGAADGEIVGLPNLNGSQKIAFLRSLVNDPSQSANLLAEAKKLNDAQAPK
29.21%   VDNKFNKEWTDADSQISRLPNLNHSQKIAFLRSLVNDPSQSANLLAEAKKLNDAQAPK
3.55%    VDNKFSKEVGAADGEIVGLPNLNHSQKIAFLRSLVNDPSQSANLLAEAKKLNDAQAPK
2.81%    VDNKFSKEVGAADGEIVGLPNLNGSQKIAFLRSLVNDPSQSANLLAEAKKLNDAQAPK
2.75%    VDNKFNKEWTDADSQISRLPNLNHSQKIAFLRSLVNDPSQSANLLAEAKKLNDAQAPK
1.51%    VDNKFSKEVVYAVYQITGLPNLNLVQKFAFVTSLCDDPSQSANLLAEAKKLNDAQAPK
1.40%    VDNKFNKEWGDADYQICGLPNLNQNQKLAFIRSLINDPSQSANLLAEAKKLNDAQAPK
1.22%    VDNKFTKEVATAVCQICCLPNLNVDQMNAFISSLFDDPSQSANLLAEAKKLNDAQAPK
22.68%   Other Sequence Less than 1% Total Reads
```

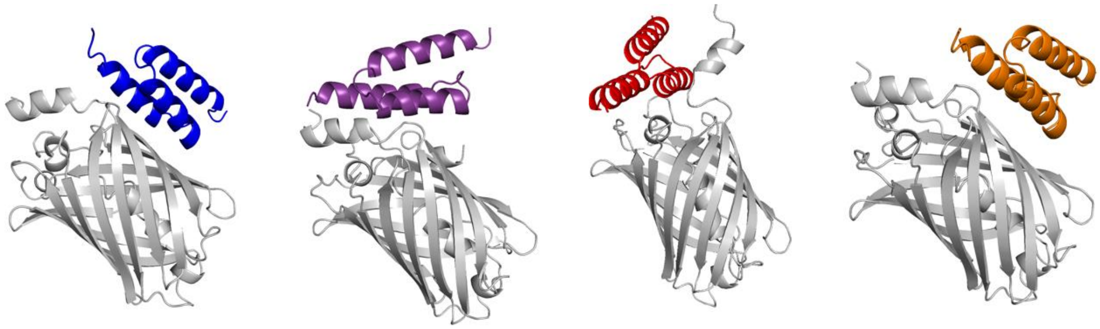

pLDDT = 81.0
pTM = 0.734
ipTM = 0.164

pLDDT = 82.8
pTM = 0.737
ipTM = 0.149

pLDDT = 83.0
pTM = 0.771
ipTM = 0.312

pLDDT = 81.0
pTM = 0.734
ipTM = 0.164

| Target | Myc (DBD) |
|---|---|
| Scaffold | Affibody |
| Total HTPS Reads | 7077 |
| Unique HTPS Reads | 468 |
| Top Variant % of Total Reads | 47 |
| Final Phage Titer (PFU/mL) | 5.37E+09 |

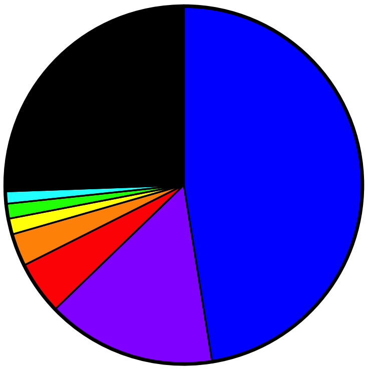

| | | |
|---|---|---|
| Library: | | VDNKF?KE???A???I??LPNLN??Q??AF??SL??DPSQSANLLAEAKKLNDAQAPK |
| (blue) | 47.46% | VDNKFSKEVFTAFDQIGGLPNLNLCQKDAFICSLVDDPSQSANLLAEAKKLNDAQAPK |
| (purple) | 15.33% | VDNKFYKEVCDAFDEICDLPNLNLLQLDAFLLSLVDDPSQSANLLAEAKKLNDAQAPK |
| (red) | 4.80% | VDNKFYKEVFSAFDEISGLPNLNGDQRNAFLDSLFNDPSQSANLLAEAKKLNDAQAPK |
| (orange) | 2.95% | VDNKFSKEVFTAFDQIGGLPNLNLLQLDAFLLSLVDDPSQSANLLAEAKKLNDAQAPK |
| (yellow) | 1.43% | VDNKFSKEVLNAFDEICILPNLNRSQKIAFIDSLFDDPSQSANLLAEAKKLNDAQAPK |
| (green) | 1.34% | VDNKFSKEVFTAFDQIGGLPNLNLCQKDAFICSLVDDPSQSANLLAEAKKLNDAQAPK |
| (cyan) | 1.12% | VDNKFSKEVVYAVYQITGLPNLNLVQKFAFVTSLCDDPSQSANLLAEAKKLNDAQAPK |
| (black) | 25.56% | Other Sequence Less than 1% Total Reads |

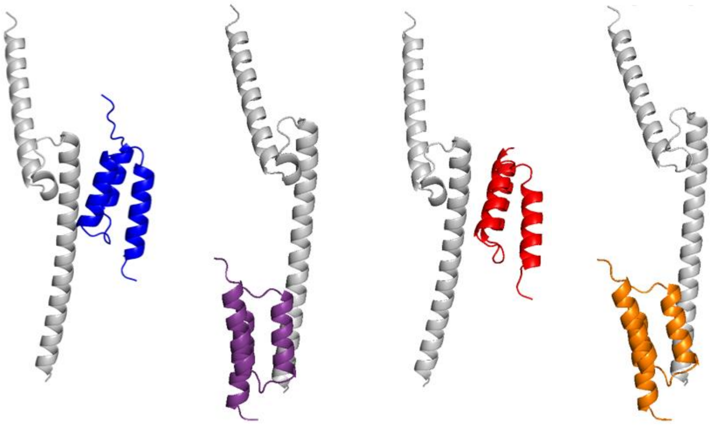

| | | | |
|---|---|---|---|
| pLDDT = 83.1 | pLDDT = 92.8 | pLDDT = 84.3 | pLDDT = 91.2 |
| pTM = 0.537 | pTM = 0.677 | pTM = 0.504 | pTM = 0.645 |
| ipTM = 0.220 | ipTM = 0.733 | ipTM = 0.130 | ipTM = 0.626 |

Dark Blue (47.46%) and Green (1.34%) have itendical AA sequences but differ in DNA coding at the V35 Residue (GTC vs GTT)

| Target | MBP |
|---|---|
| Scaffold | Affibody |
| Total HTPS Reads | 2771 |
| Unique HTPS Reads | 356 |
| Top Variant % of Total Reads | 23 |
| Final Phage Titer (PFU/mL) | 1.20E+09 |

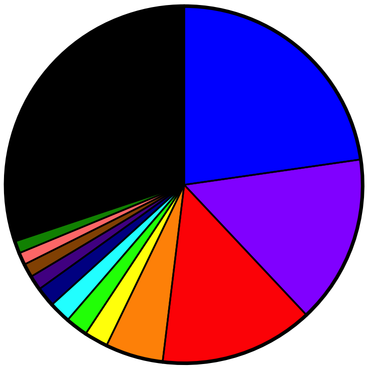

| | |
|---|---|
| Library: | VDNKF?KE???A???I??LPNLN??Q??AF??SL??DPSQSANLLAEAKKLNDAQAPK |
| 22.74% | VDNKFNKELFYAFHQIYALPNLNAAQVRAFVGSLDDDPSQSANLLAEAKKLNDAQAPK |
| 15.27% | VDNKFTKEVNYAASEIRYLPNLNHVQKLAFISSLGDDPSQSANLLAEAKKLNDAQAPK |
| 13.93% | VDNKFSKELGAANGQIIRLPNLNPCQQYAFILSLADDPSQSANLLAEAKKLNDAQAPK |
| 5.23% | VDNKFNKEVFYAYPQILCLPNLNASQRSAFVVSLLNDPSQSANLLAEAKKLNDAQAPK |
| 2.13% | VDNKFYKEGVDAVSQICTLPNLNLDQRSAFLGSLVDDPSQSANLLAEAKKLNDAQAPK |
| 2.02% | VDNKFSKEVVYAVYQITGLPNLNLVQKFAFVTSLCDDPSQSANLLAEAKKLNDAQAPK |
| 1.95% | VDNKFYKEGDNAISEIVLLPNLNLCQMLAFVNSLPDDPSQSANLLAEAKKLNDAQAPK |
| 1.34% | VDNKFTKEVATAVCQICCLPNLNVDQMNAFISSLFDDPSQSANLLAEAKKLNDAQAPK |
| 1.80% | VDNKFTKESFI---EIVNLPNLNLRQIDAFVSSLFDDPSQSANLLAEAKKLNDAQAPK |
| 1.23% | VDNKFYKEWSTAICEIVGLPNLNLSQQHAFIFSLLDDPSQSANLLAEAKKLNDAQAPK |
| 1.12% | VDNKFNKEVCDAYAQIGNLPNLNVYQIVAFIRSLDNDPSQSANLLAEAKKLNDAQAPK |
| 1.12% | VDNKFNKEGDTAVLEIGHLPNLNLHQQYAFLHSLYNDPSQSANLLAEAKKLNDAQAPK |
| 30.13% | Other Sequence Less than 1% Total Reads |

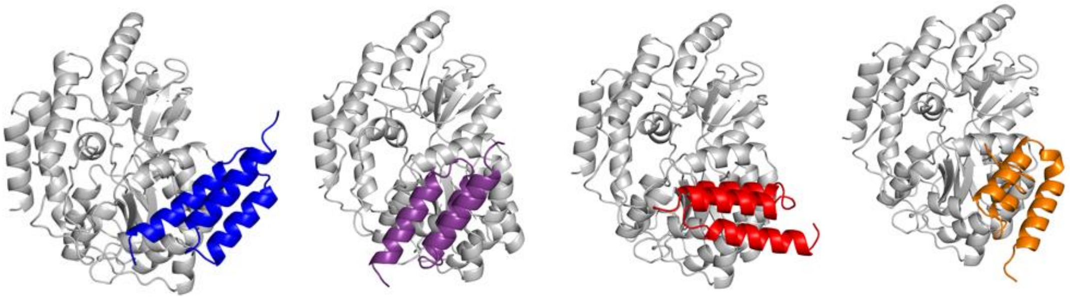

| | | | |
|---|---|---|---|
| pLDDT = 95.3 | pLDDT = 90.2 | pLDDT = 90.4 | pLDDT = 89.2 |
| pTM = 0.917 | pTM = 0.838 | pTM = 0.865 | pTM = 0.866 |
| ipTM = 0.817 | ipTM = 0.348 | ipTM = 0.438 | ipTM = 0.461 |

| Target | Calprotectin heterodimer 6aa linker |
|---|---|
| Scaffold | Affibody |
| Total HTPS Reads | 3822 |
| Unique HTPS Reads | 904 |
| Top Variant % of Total Reads | 24 |
| Final Phage Titer (PFU/mL) | 2.19E+09 |

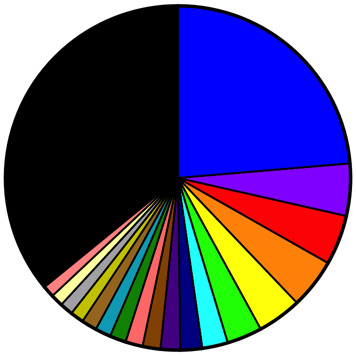

```
Library: VDNKF?KE???A???I??LPNLN??Q??AF??SL??DPSQSANLLAEAKKLNDAQAPK
23.65%   VDNKFYKEVDAATDEIHLLPNLNENQRHAFVTSLDNDPSQSANLLAEAKKLNDAQAPK
4.89%    VDNKFNKEGYYANYQIRILPNLNPNQKGAFLTSLNDDPSQSANLLAEAKKLNDAQAPK
4.68%    VDNKFNKEGNYAANQITGLPNLNVGQGIAFICSLTDDPSQSANLLAEAKKLNDAQAPK
4.68%    VDNKFNKEGRSADLQITILPNLNDGQRVAFIGSLADDPSQSANLLAEAKKLNDAQAPK
4.13%    VDNKFNKEGIYANSEITDLPNLNEGQLIAFVSSLGNDPSQSANLLAEAKKLNDAQAPK
3.30%    VDNKFTKEWCSAYVEIGVLPNLNDGQKNAFLYSLVDDPSQSANLLAEAKKLNDAQAPK
2.41%    VDNKFSKEGGSAAHQICNLPNLNGFQMAAFLGSLRNDPSQSANLLAEAKKLNDAQAPK
1.83%    VDNKFSKEVLNAVNQISFLPNLNASQECAFIDSLFNDPSQSANLLAEAKKLNDAQAPK
2.04%    VDNKFSKEVDNAINEIGDLPNLNVYQIIAFVRSLSDDPSQSANLLAEAKKLNDAQAPK
1.67%    VDNKFYKEWYDATSS------------------LFDDPSQSANLLAEAKKLNDAQAPK
1.62%    VDNKFSKELCAAIGQISSLPNLNGAQENAFIVSLADDPSQSANLLAEAKKLNDAQAPK
1.52%    VDNKFSKELFSAAFQIPDLPNLNGHQRTAFVGSLGNDPSQSANLLAEAKKLNDAQAPK
1.47%    VDNKFSKELNYANAQIHTLPNLNHLQLNAFVSSLCDDPSQSANLLAEAKKLNDAQAPK
1.44%    VDNKFTKELVYATNEIYNLPNLNRLQRRAFLDSLSNDPSQSANLLAEAKKLNDAQAPK
1.26%    VDNKFTKEWTDAFCEISCLPNLNAHQKVAFLGSLLDDPSQSANLLAEAKKLNDAQAPK
1.23%    VDNKFYKEVDAAASEICSLPNLNRNQRSAFICSLYNDPSQSANLLAEAKKLNDAQAPK
1.15%    VDNKFSKEGCAADFQIGFLPNLNAGQKIAFLHSLSDDPSQSANLLAEAKKLNDAQAPK
1.10%    VDNKFSKEVVSANSEIISLPNLNGVQRHAFVGSLHDDPSQSANLLAEAKKLNDAQAPK
35.92%   Other Sequence less than 1%
```

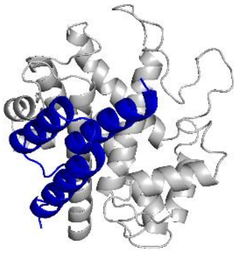

pLDDT = 73.5
pTM = 0.709
ipTM = 0.360

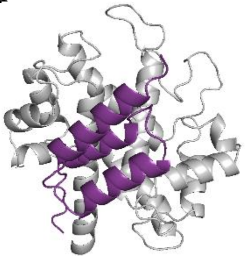

pLDDT = 78.2
pTM = 0.750
ipTM = 0.325

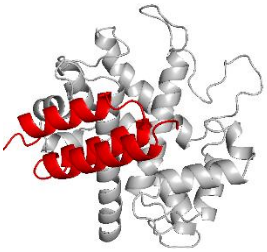

pLDDT = 83.2
pTM = 0.831
ipTM = 0.657

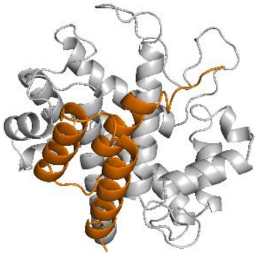

pLDDT = 86.4
pTM = 0.865
ipTM = 0.808

| | |
|---|---|
| Target | GABARAP |
| Scaffold | Affibody |
| Total HTPS Reads | 7244 |
| Unique HTPS Reads | 357 |
| Top Variant % of Total Reads | 57 |
| Final Phage Titer (PFU/mL) | 2.82E+08 |

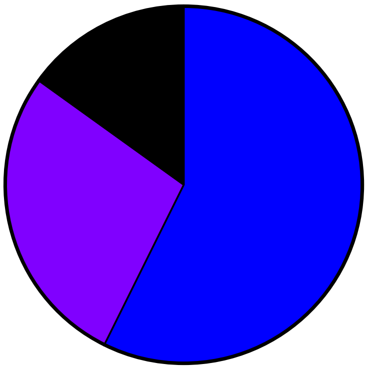

```
Library: VDNKF?KE???A???I??LPNLN??Q??AF??SL??DPSQSANLLAEAKKLNDAQAPK
57.34%   VDNKFNKEGDTAVLEIGHLPNLNLHQQYAFLHSLYNDPSQSANLLAEAKKLNDAQAPK
27.55%   VDNKFSKELVDADFEIIDLPNLNENQQVAFVGSLVNDPSQSANLLAEAKKLNDAQAPK
15.10%   Other Sequence Less than 1% Total Reads
```

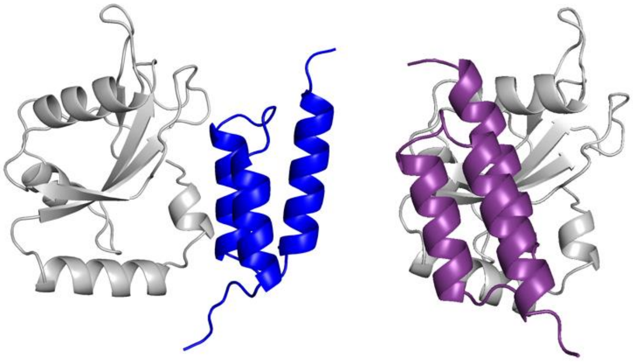

pLDDT = 84.8
pTM = 0.657
ipTM = 0.116

pLDDT = 81.4
pTM = 0.707
ipTM = 0.299

| Target | 14-3-3z Monomer |
|---|---|
| Scaffold | Affibody |
| Total HTPS Reads | 4556 |
| Unique HTPS Reads | 642 |
| Top Variant % of Total Reads | 38 |
| Final Phage Titer (PFU/mL) | 2.51E+09 |

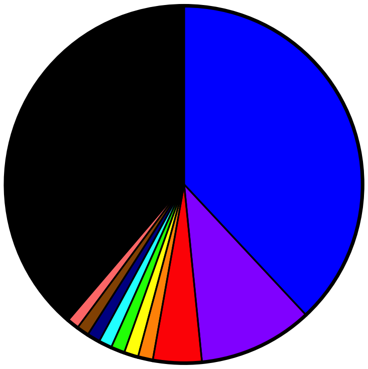

| | |
|---|---|
| Library: | VDNKF?KE???A???I??LPNLN??Q??AF??SL??DPSQSANLLAEAKKLNDAQAPK |
| 38.04% | VDNKFNKEWLAAARQICDLPNLNRFQKFAFIRSLLDDPSQSANLLAEAKKLNDAQAPK |
| 10.36% | VDNKFNKEWSAAIGEIGCLPNLNPDQGYAFVRSLLDDPSQSANLLAEAKKLNDAQAPK |
| 4.41% | VDNKFNKELSTAFLQIRYLPNLNDSQIYAFVYSLCDDPSQSANLLAEAKKLNDAQAPK |
| 1.34% | VDNKFSKEVVYAVYQITGLPNLNLVQKFAFVTSLCDDPSQSANLLAEAKKLNDAQAPK |
| 1.29% | VDNKFSKEWCSAIREICRLPNLNENQRCAFLFSLDNDPSQSANLLAEAKKLNDAQAPK |
| 1.25% | VDNKFSKEWNTAFVEIVDLPNLNESQMFAFLFSLDNDPSQSANLLAEAKKLNDAQAPK |
| 1.21% | VDNKFTKEVATAVCQICCLPNLNVDQMNAFISSLFDDPSQSANLLAEAKKLNDAQAPK |
| 1.19% | VDNKFNKEWLAAARQICDLPNLNPDQGYAFVRSLLDDPSQSANLLAEAKKLNDAQAPK |
| 1.10% | VDNKFYKEWSTAICEIVGLPNLNLSQQHAFIFSLLDDPSQSANLLAEAKKLNDAQAPK |
| 1.08% | VDNKFTKESFI---EIVNLPNLNLRQIDAFVSSLFDDPSQSANLLAEAKKLNDAQAPK |
| 38.74% | Other Sequence Less than 1% Total Reads |

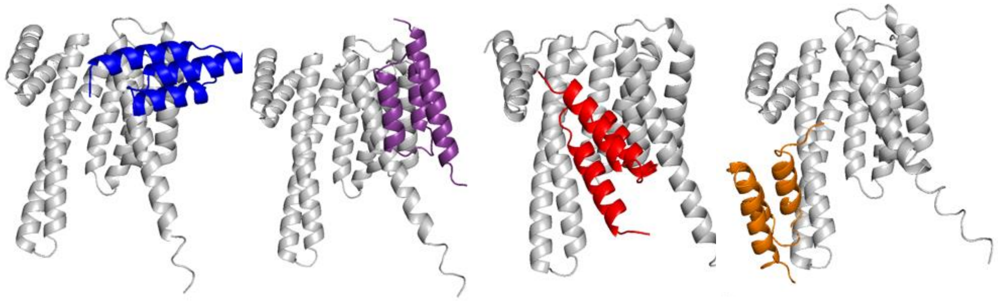

| | | | |
|---|---|---|---|
| pLDDT = 83.8 | pLDDT = 81.6 | pLDDT = 78.6 | pLDDT = 80.8 |
| pTM = 0.767 | pTM = 0.750 | pTM = 0.783 | pTM = 0.812 |
| ipTM = 0.316 | ipTM = 0.228 | ipTM = 0.309 | ipTM = 0.532 |

| Target | FnbpA |
|---|---|
| Scaffold | Affibody |
| Total HTPS Reads | 12225 |
| Unique HTPS Reads | 265 |
| Top Variant % of Total Reads | 90 |
| Final Phage Titer (PFU/mL) | 5.37E+09 |

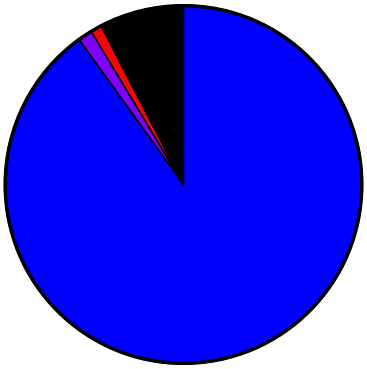

```
Library: VDNKF?KE???A???I??LPNLN??Q??AF??SL??DPSQSANLLAEAKKLNDAQAPK
90.04%   VDNKFSKEWSYAALQISRLPNLNQTQIVAFVNSLRNDPSQSANLLAEAKKLNDAQAPK
1.28%    VDNKFNKEWFTANYEIGTLPNLNDVQRYAFVCSLYNDPSQSANLLAEAKKLNDAQAPK
1.01%    VDNKFSKE*
7.66%  Other Sequence Less than 1% Total Reads
```

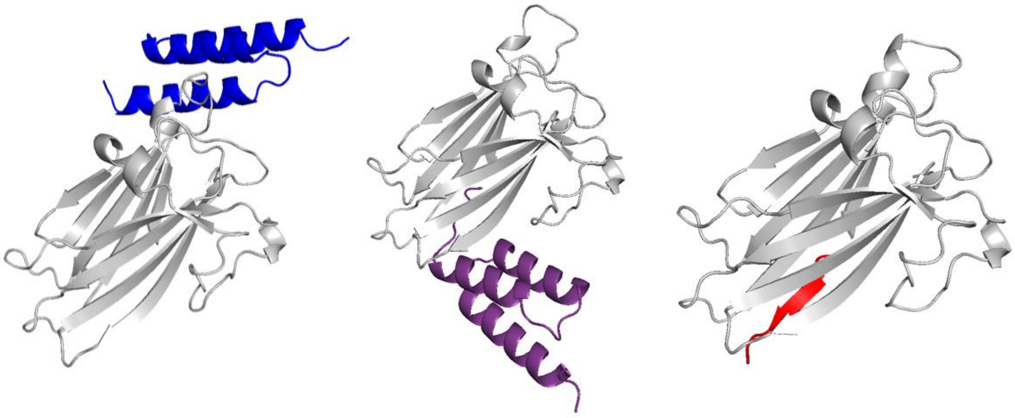

| pLDDT = 83.3 | pLDDT = 81.2 | pLDDT = 82.4 |
|---|---|---|
| pTM = 0.697 | pTM = 0.719 | pTM = 0.825 |
| ipTM = 0.170 | ipTM = 0.389 | ipTM = 0.517 |

| Target | PAL |
|---|---|
| Scaffold | Affibody |
| Total HTPS Reads | 3447 |
| Unique HTPS Reads | 637 |
| Top Variant % of Total Reads | 23 |
| Final Phage Titer (PFU/mL) | 5.13E+09 |

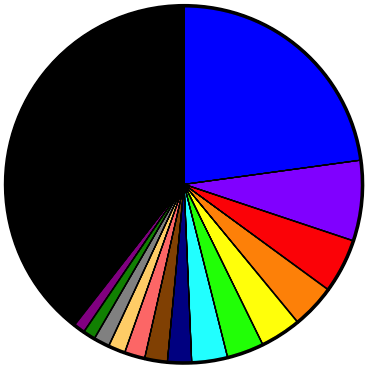

```
Library: VDNKF?KE???A???I??LPNLN??Q??AF??SL??DPSQSANLLAEAKKLNDAQAPK
22.83%   VDNKFSKELCSAICEISFLPNLNEDQVIAFLRSLGDDPSQSANLLAEAKKLNDAQAPK
7.28%    VDNKFYKELTTAAVQIRLLPNLNVAQKGAFIASLGDDPSQSANLLAEAKKLNDAQAPK
4.93%    VDNKFSKEVGDATLEINGLPNLNGSQEHAFLCSLRDDPSQSANLLAEAKKLNDAQAPK
3.97%    VDNKFSKEWSYAALQISRLPNLNQTQIVAFVNSLRNDPSQSANLLAEAKKLNDAQAPK
3.71%    VDNKFNKELSNAICEIHSLPNLNQDQKSAFLISLGDDPSQSANLLAEAKKLNDAQAPK
3.34%    VDNKFNKELYYAIVQIGILPNLNPGQVHAFLRSLGDDPSQSANLLAEAKKLNDAQAPK
3.25%    VDNKFYKEGTAAVNEIPLLPNLNHGQRTAFVGSLVDDPSQSANLLAEAKKLNDAQAPK
2.18%    VDNKFYKELPNAIIEICGLPNLNESQRTAFLRSLGNDPSQSANLLAEAKKLNDAQAPK
2.06%    VDNKFSKELSDAVIEINFLPNLNEYQRDAFLRSLGDDPSQSANLLAEAKKLNDAQAPK
1.86%    VDNKFSKE*
1.54%    VDNKFSKELNNAIVQINGLPNLNAGQLAAFLRSLGDDPSQSANLLAEAKKLNDAQAPK
1.45%    VDNKFSKEVSAASSEIVGLPNLNVGQRTAFLFSLGNDPSQSANLLAEAKKLNDAQAPK
1.10%    VDNKFNKEGIYAIFEISRLPNLNGCQRVAFLVSLYNDPSQSANLLAEAKKLNDAQAPK
1.04%    VDNKFNKELGNANSQIGGLPNLNGNQVPAFLISLYDDPSQSANLLAEAKKLNDAQAPK
Other Sequence Less than 1% Total Reads
```

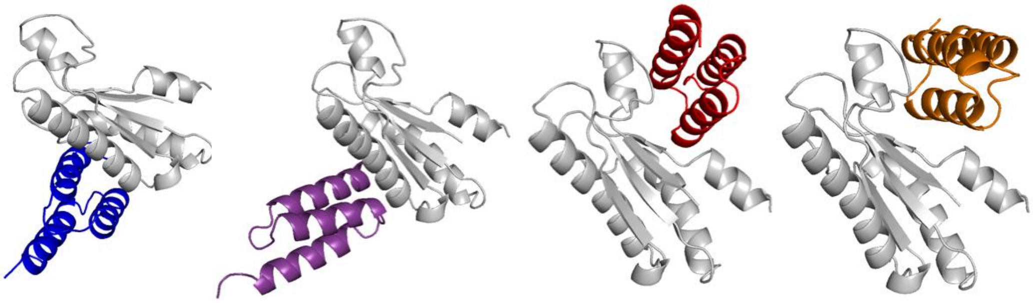

| pLDDT = 88.5 | pLDDT = 86.9 | pLDDT = 84.9 | pLDDT = 89.3 |
|---|---|---|---|
| pTM = 0.770 | pTM = 0.683 | pTM = 0.672 | pTM = 0.771 |
| ipTM = 0.466 | ipTM = 0.189 | ipTM = 0.160 | ipTM = 0.499 |

| Target | PIK3CB (3-115) |
|---|---|
| Scaffold | Affibody |
| Total HTPS Reads | 10189 |
| Unique HTPS Reads | 641 |
| Top Variant % of Total Reads | 45 |
| Final Phage Titer (PFU/mL) | 6.03E+09 |

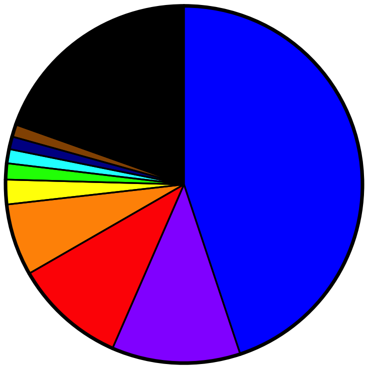

```
Library: VDNKF?KE???A???I??LPNLN??Q??AF??SL??DPSQSANLLAEAKKLNDAQAPK
 44.89%  VDNKFNKEGIYAIFEISRLPNLNGCQRVAFLVSLYNDPSQSANLLAEAKKLNDAQAPK
 11.66%  VDNKFNKEWCDAIFEISGLPNLNPNQKRASLLSLLDDPSQSANLLAEAKKLNDAQAPK
 10.14%  VDNKFYKELATAFIQIRNLPNLNVGQRVAFVSSLVDDPSQSANLLAEAKKLNDAQAPK
 6.52%   VDNKFYKELIDAYREIRILPNLNESQMCAFISSLIDDPSQSANLLAEAKKLNDAQAPK
 2.24%   VDNKFSKEWSYAALQISRLPNLNQTQIVAFVNSLRNDPSQSANLLAEAKKLNDAQAPK
 1.45%   VDNKFYKELRAAFSEIRALPNLNAFQLCAFVGSLVDDPSQSANLLAEAKKLNDAQAPK
 1.29%   VDNKFNKEGIYAIFEISRLPNLNVGQRVAFVSSLVDDPSQSANLLAEAKKLNDAQAPK
 1.14%   VDNKFTKEWTSAYFEIARLPNLNRSQRIAFILSLGDDPSQSANLLAEAKKLNDAQAPK
 1.13%   VDNKFSKE*
 19.55%  Other Sequence Less than 1% Total Reads
```

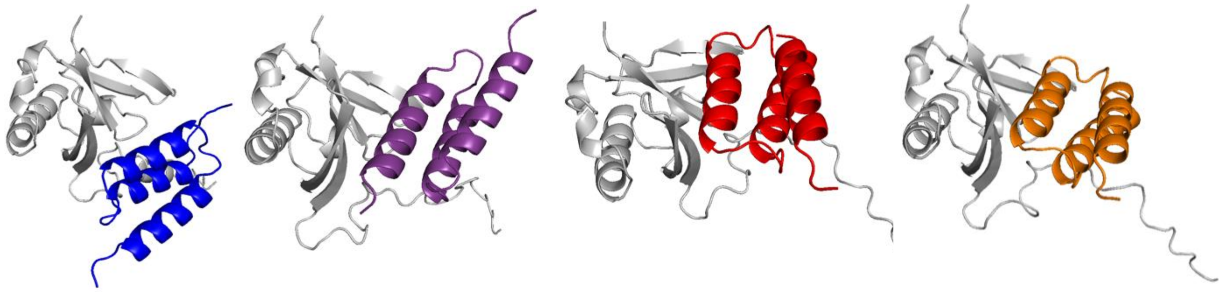

| | | | |
|---|---|---|---|
| pLDDT = 84.6 | pLDDT = 84.6 | pLDDT = 88.2 | pLDDT = 88.0 |
| pTM = 0.781 | pTM = 0.796 | pTM = 0.835 | pTM = 0.833 |
| ipTM = 0.686 | ipTM = 0.796 | ipTM = 0.878 | ipTM = 0.872 |

| Target | SortaseA (25-206) |
|---|---|
| Scaffold | Affibody |
| Total HTPS Reads | 6249 |
| Unique HTPS Reads | 182 |
| Top Variant % of Total Reads | 87 |
| Final Phage Titer (PFU/mL) | 1.48E+08 |

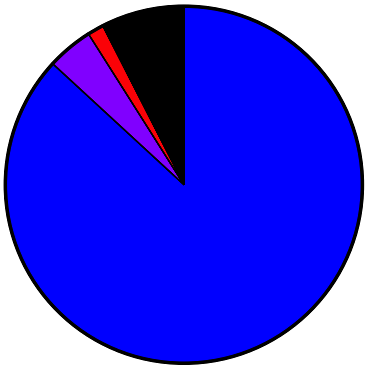

```
Library: VDNKF?KE???A???I??LPNLN??Q??AF??SL??DPSQSANLLAEAKKLNDAQAPK
86.85%   VDNKFSKE*
4.16%    VDNKFSKEWSYAALQISRLPNLNQTQIVAFVNSLRNDPSQSANLLAEAKKLNDAQAPK
1.47%    VDNKFNKEGIYAIFEISRLPNLNGCQRVAFLVSLYNDPSQSANLLAEAKKLNDAQAPK
7.52%  Other Sequence Less than 1% Total Reads
```

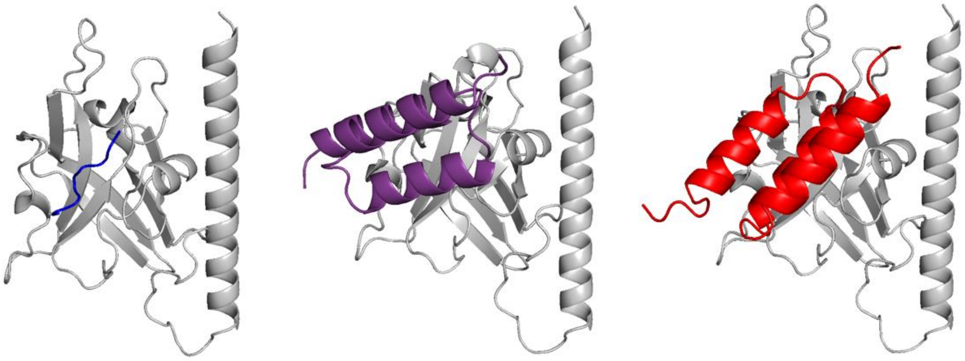

pLDDT = 86.0
pTM = 0.848
ipTM = 0.475

pLDDT = 65.2
pTM = 0.573
ipTM = 0.155

pLDDT = 73.1
pTM = 0.651
ipTM = 0.249

| | |
|---|---|
| Target | TNFR2 |
| Scaffold | Affibody |
| Total HTPS Reads | 6064 |
| Unique HTPS Reads | 3171 |
| Top Variant % of Total Reads | 52 |
| Final Phage Titer (PFU/mL) | 1.51E+07 |

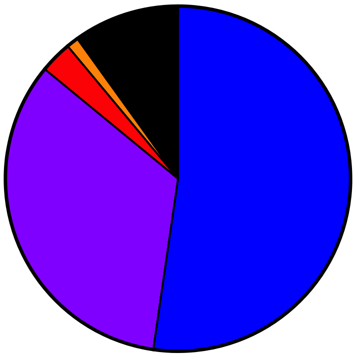

| | | |
|---|---|---|
| Library: | | VDNKF?KE???A???I??LPNLN??Q??AF??SL??DPSQSANLLAEAKKLNDAQAPK |
| (blue) | 52.29% | VDNKFTKEGNSAICEICCLPNLNPCQKCAFICSLSDDPSQSANLLAEAKKLNDAQAPK |
| (purple) | 33.66% | VDNKFTKEWCSAYVEIGVLPNLNDGQKNAFLYSLVDDPSQSANLLAEAKKLNDAQAPK |
| (red) | 3.00% | VDNKFSKEVDNAINEIGDLPNLNVYQIIAFVRSLSDDPSQSANLLAEAKKLNDAQAPK |
| (orange) | 1.12% | VDNKFYKEVDAAASEICSLPNLNRNQRSAFICSLYNDPSQSANLLAEAKKLNDAQAPK |
| (black) | 9.93% | Other Sequence less than 1% |

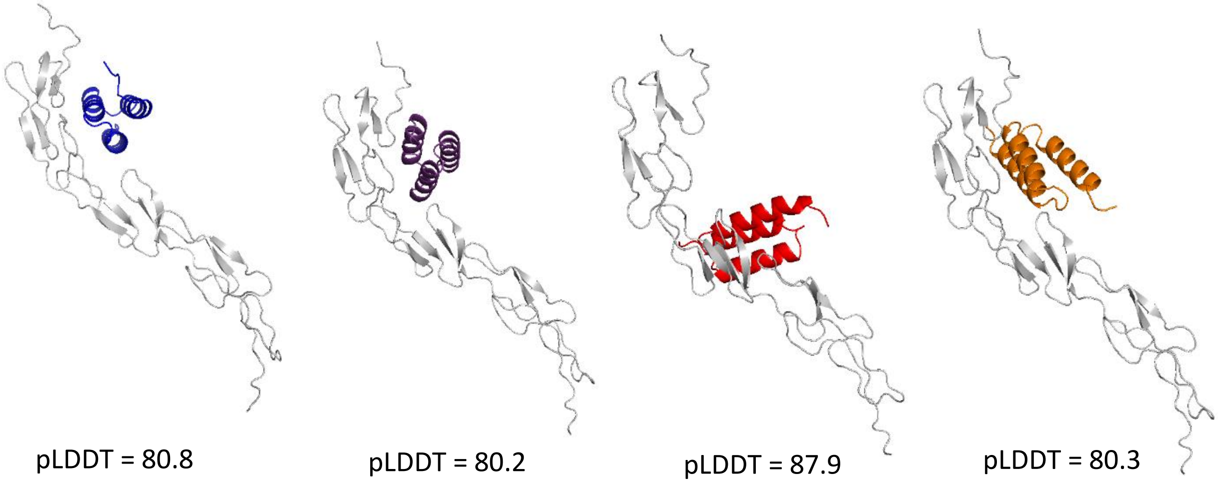

| | | | |
|---|---|---|---|
| pLDDT = 80.8 | pLDDT = 80.2 | pLDDT = 87.9 | pLDDT = 80.3 |
| pTM = 0.657 | pTM = 0.605 | pTM = 0.658 | pTM = 0.617 |
| ipTM = 0.395 | ipTM = 0.291 | ipTM = 0.623 | ipTM = 0.297 |

| Target | SH3BP5 |
|---|---|
| Scaffold | Affibody |
| Total HTPS Reads | 4845 |
| Unique HTPS Reads | 82 |
| Top Variant % of Total Reads | 96 |
| Final Phage Titer (PFU/mL) | 1.05E+09 |

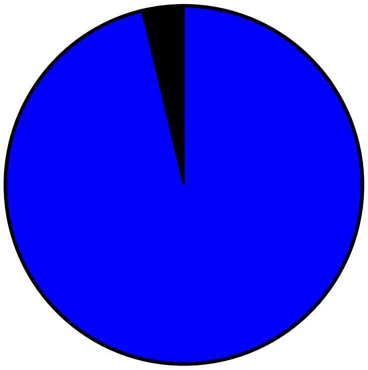

Library: VDNKF?KE???A???I??LPNLN??Q??AF??SL??DPSQSANLLAEAKKLNDAQAPK

96.24% VDNKFSKELAAAYCQIISLPNLNPHQRLAFLLSLPDDPSQSANLLAEAKKLNDAQAPK

3.76% Other Sequence Less than 1% Total Reads

**Insufficient Confidence to Predict Complex Structure (pTM)**

pLDDT = 72.6
pTM = 0.410
ipTM = 0.162

| Target | DltD (strep) |
|---|---|
| Scaffold | Affibody |
| Total HTPS Reads | 1342 |
| Unique HTPS Reads | 129 |
| Top Variant % of Total Reads | 79 |
| Final Phage Titer (PFU/mL) | 2.19E+08 |

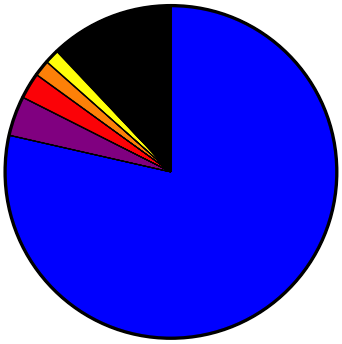

Library: VDNKF?KE???A???I??LPNLN??Q??AF??SL??DPSQSANLLAEAKKLNDAQAPK

78.54% VDNKFNKEGLDAFFQIASLPNLNRIQVFAFIASLRDDPSQSANLLAEAKKLNDAQAPK

3.87% VDNKFSKEVHSAYAEIGLLPNLNERQMAAFLCSLGDDPSQSANLLAEAKKLNDAQAPK

2.53% VDNKFSKELAAAYCQIISLPNLNPHQRLAFLLSLPDDPSQSANLLAEAKKLNDAQAPK

1.56% VDNKFNKEVVNASNQIVGLPNLNGSQGFAFVISLGDDPSQSANLLAEAKKLNDAQAPK

1.42% VDNKFYKEVIDAVGQIRRLPNLNPGQKGAFIRSLNDDPSQSANLLAEAKKLNDAQAPK

12.07% Other Sequence Less than 1% Total Reads

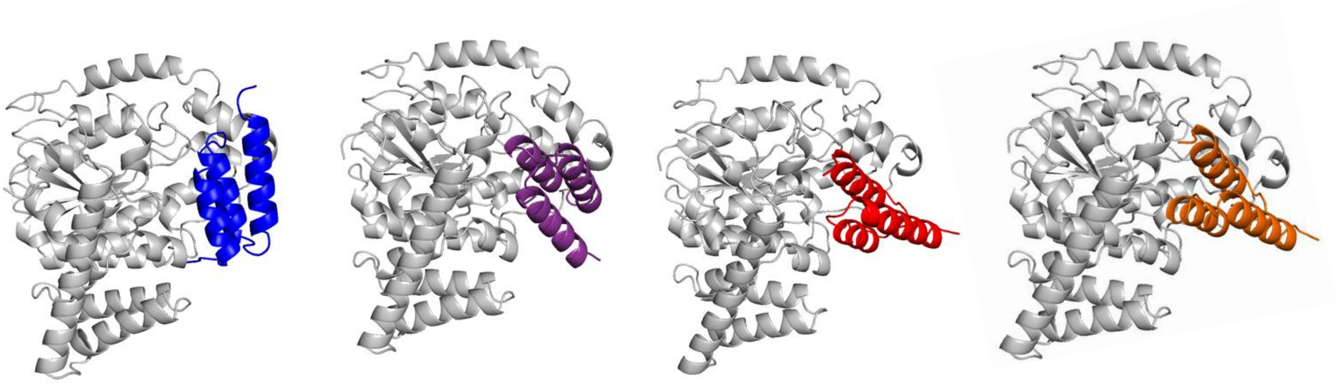

| | | | |
|---|---|---|---|
| pLDDT = 88.0<br>pTM = 0.869<br>ipTM = 0.389 | pLDDT = 87.4<br>pTM = 0.873<br>ipTM = 0.409 | pLDDT = 85.4<br>pTM = 0.844<br>ipTM = 0.252 | pLDDT = 87.6<br>pTM = 0.869<br>ipTM = 0.381 |

| Target | RB1 |
| --- | --- |
| Scaffold | Affibody |
| Total HTPS Reads | 6154 |
| Unique HTPS Reads | 5338 |
| Top Variant % of Total Reads | 87 |
| Final Phage Titer (PFU/mL) | 1.86E+09 |

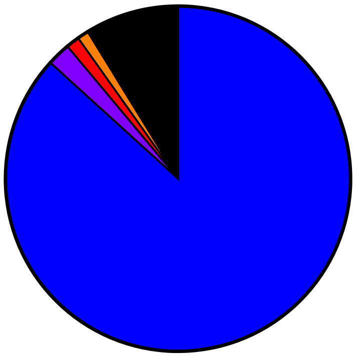

```
Library: VDNKF?KE???A???I??LPNLN??Q??AF??SL??DPSQSANLLAEAKKLNDAQAPK
86.74%   VDNKFTKELVYATNEIYNLPNLNRLQRRAFLDSLSNDPSQSANLLAEAKKLNDAQAPK
2.18%    VDNKFTKEWCSAYVEIGVLPNLNDGQKNAFLYSLVDDPSQSANLLAEAKKLNDAQAPK
1.25%    VDNKFSKEVDNAINEIGDLPNLNVYQIIAFVRSLSDDPSQSANLLAEAKKLNDAQAPK
1.06%    VDNKFTKEWTDAFCEISCLPNLNAHQKVAFLGSLLDDPSQSANLLAEAKKLNDAQAPK
8.77%    Other Sequence less than 1%
```

**Insufficient Confidence to Predict Complex Structure (pLDDT, pTM)**

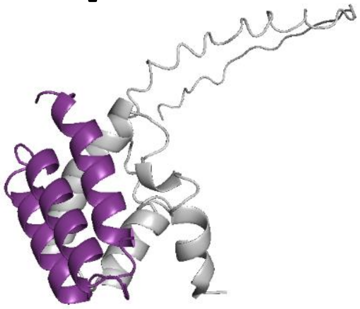

**Insufficient Confidence to Predict Complex Structure (pTM)**

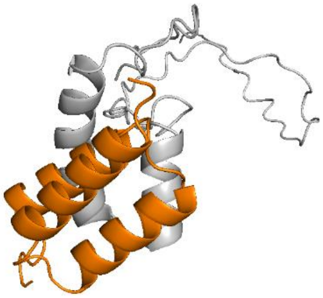

pLDDT = 48.9
pTM = 0.383
ipTM = 0.186

pLDDT = 61.2
pTM = 0.518
ipTM = 0.635

pLDDT = 54.2
pTM = 0.420
ipTM = 0.229

pLDDT = 65.1
pTM = 0.541
ipTM = 0.746

| Target | SasG (930-1060) |
|---|---|
| Scaffold | Affibody |
| Total HTPS Reads | 1166 |
| Unique HTPS Reads | 207 |
| Top Variant % of Total Reads | 58 |
| Final Phage Titer (PFU/mL) | 1.38E+09 |

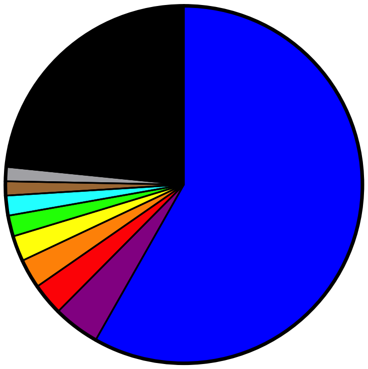

```
Library: VDNKF?KE???A???I??LPNLN??Q??AF??SL??DPSQSANLLAEAKKLNDAQAPK
58.23%   VDNKFSKELAAAYCQIISLPNLNPHQRLAFLLSLPDDPSQSANLLAEAKKLNDAQAPK
4.20%    VDNKFSKEVCDAIHEIHFLPNLNDAQRIAFIFSLFDDPSQSANLLAEAKKLNDAQAPK
2.92%    VDNKFYKELCSAADEIFVLPNLNDDQVLAFINSLRNDPSQSANLLAEAKKLNDAQAPK
2.66%    VDNKFSKELAAAYCQIISLPNLNPHQRLAFLLSLPDDPSQSANLLAEAKKLNDAQAPK
2.32%    VDNKFTKEVDSAVFEIHILPNLNDYQRVAFIFSLFDDPSQSANLLAEAKKLNDAQAPK
1.89%    VDNKFYKELSTAVYEIFALPNLNESQVVAFVVSLFDDPSQSANLLAEAKKLNDAQAPK
1.80%    VDNKFNKEVCDAYAQIGNLPNLNVYQIVAFIRSLDNDPSQSANLLAEAKKLNDAQAPK
1.29%    VDNKFTKEVPNASDQIFLLPNLNDYQVLAFLHSLRDDPSQSANLLAEAKKLNDAQAPK
1.29%    VDNKFYKELAYAIHEIFCLPNLNEPQVCAFVVSLCDDPSQSANLLAEAKKLNDAQAPK
23.41%   Other Sequence Less than 1% Total Reads
```

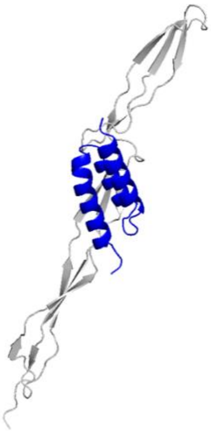

pLDDT = 77.6
pTM = 0.521
ipTM = 0.206

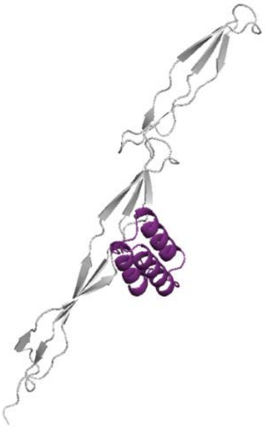

pLDDT = 81.8
pTM = 0.516
ipTM = 0.118

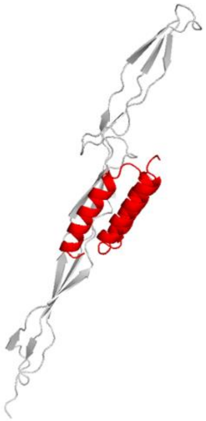

pLDDT = 79.8
pTM = 0.482
ipTM = 0.163

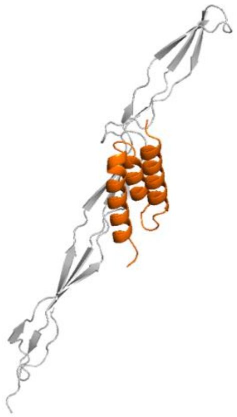

pLDDT = 77.6
pTM = 0.521
ipTM = 0.206

| | |
|---|---|
| Target | BRD4 |
| Scaffold | Affibody |
| Total HTPS Reads | 10988 |
| Unique HTPS Reads | 1657 |
| Top Variant % of Total Reads | 13 |
| Final Phage Titer (PFU/mL) | 1.35E+09 |

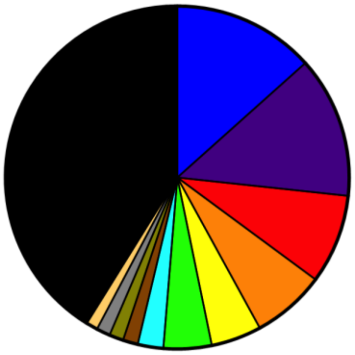

```
Library: VDNKF?KE???A???I??LPNLN??Q??AF??SL??DPSQSANLLAEAKKLNDAQAPK
13.37%   VDNKFSKEVHNAVGEIVILPNLNASQVHAFIVSLGNDPSQSANLLAEAKKLNDAQAPK
13.34%   VDNKFSKEWHSAYAEIAHLPNLNPSQVCAFIGSLLDDPSQSANLLAEAKKLNDAQAPK
 8.43%   VDNKFYKEWFDANVQIAVLPNLNEYQVIAFISSLANDPSQSANLLAEAKKLNDAQAPK
 6.90%   VDNKFYKEGPNAFREIGCLPNLNGTQKAAFLRSLCDDPSQSANLLAEAKKLNDAQAPK
 4.74%   VDNKFYKEVSDAITQISILPNLNDNQVRAFVYSLSDDPSQSANLLAEAKKLNDAQAPK
 4.52%   VDNKFYKEGGSAVVQIVALPNLNESQVHAFLCSLSNDPSQSANLLAEAKKLNDAQAPK
 2.40%   VDNKFTKEGYDANCQITSLPNLNLGQGSAFVRSLRNDPSQSANLLAEAKKLNDAQAPK
 1.41%   VDNKFYKEWFTAAGQISVLPNLNQSQRYAFLCSLSDDPSQSANLLAEAKKLNDAQAPK
 1.34%   VDNKFYKEWFSANVEISNLPNLNEAQMFAFIGSLYNDPSQSANLLAEAKKLNDAQAPK
 1.29%   VDNKFYKEWFSASSQICALPNLNEDQIFAFLCSLGDDPSQSANLLAEAKKLNDAQAPK
 1.08%   VDNKFYKEGTYASYEINNLPNLNLVQRSAFVFSLYNDPSQSANLLAEAKKLNDAQAPK
41.17%   Other Sequence Less Than 1% Total Reads
```

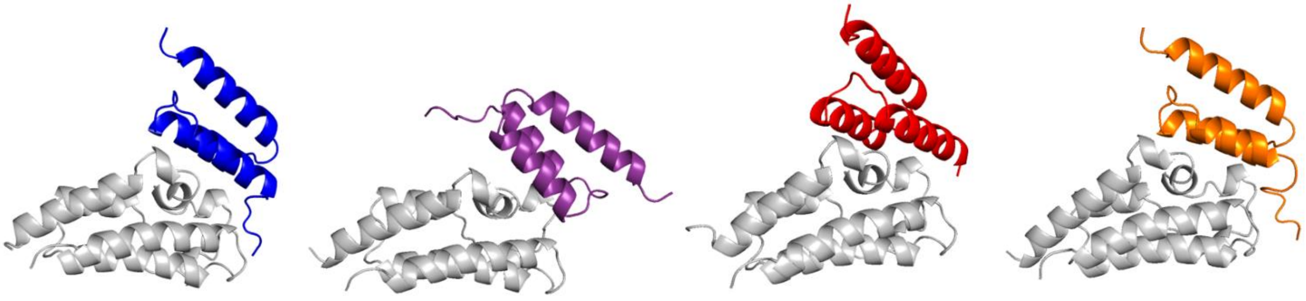

pLDDT = 88.5
pTM = 0.775
ipTM = 0.480

pLDDT = 86.8
pTM = 0.707
ipTM = 0.287

pLDDT = 95.2
pTM = 0.904
ipTM = 0.866

pLDDT = 83.9
pTM = 0.721
ipTM = 0.327

| Target | IsdB |
|---|---|
| Scaffold | Affibody |
| Total HTPS Reads | 5901 |
| Unique HTPS Reads | 199 |
| Top Variant % of Total Reads | 89 |
| Final Phage Titer (PFU/mL) | 2.29E+09 |

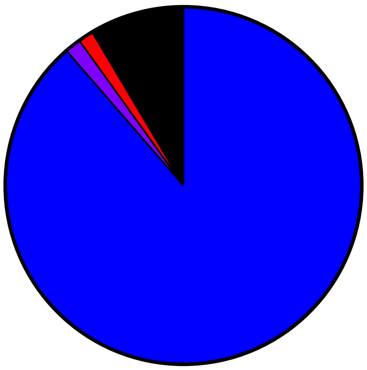

Library: VDNKF?KE???A???I??LPNLN??Q??AF??SL??DPSQSANLLAEAKKLNDAQAPK

88.60% VDNKFNKEVCSAFNEIGNLPNLNLGQKIAFVRSLYNDPSQSANLLAEAKKLNDAQAPK

1.46% VDNKFYKEVDAAASEICSLPNLNRNQRSAFICSLYNDPSQSANLLAEAKKLNDAQAPK

1.37% VDNKFYKEVIYAAAEIFGLPNLNVGQRSAFVCSLYDDPSQSANLLAEAKKLNDAQAPK

8.57% Other Sequence Less than 1% Total Reads

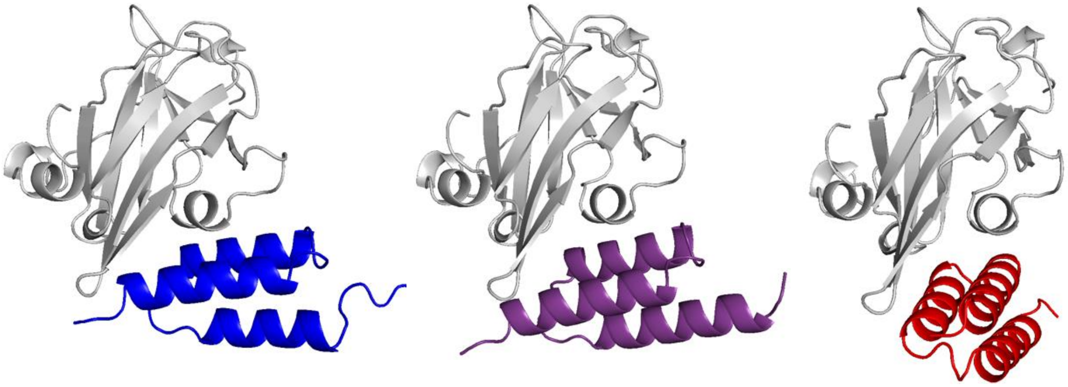

pLDDT = 89.0
pTM = 0.815
ipTM = 0.556

pLDDT = 84.2
pTM = 0.730
ipTM = 0.256

pLDDT = 85.4
pTM = 0.745
ipTM = 0.301

| | |
|---|---|
| Target | LMO2 (28-150) |
| Scaffold | Affibody |
| Total HTPS Reads | 6909 |
| Unique HTPS Reads | 291 |
| Top Variant % of Total Reads | 50 |
| Final Phage Titer (PFU/mL) | 5.01E+08 |

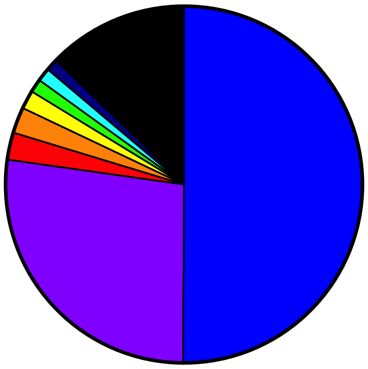

| | |
|---|---|
| Library: | VDNKF?KE???A???I??LPNLN??Q??AF??SL??DPSQSANLLAEAKKLNDAQAPK |
| 50.08% | VDNKFSKEGIYAISQIVRLPNLNGFQEVAFLSSLPNDPSQSANLLAEAKKLNDAQAPK |
| 27.15% | VDNKFTKEVCYAYIEIVYLPNLNPDQISAFISSLHDDPSQSANLLAEAKKLNDAQAPK |
| 2.46% | VDNKFSKEGIYAISQIVRLPNLNPDQISAFISSLHDDPSQSANLLAEAKKLNDAQAPK |
| 2.36% | VDNKFYKEGYSAYREICTLPNLNGCQRAAFICSLFDDPSQSANLLAEAKKLNDAQAPK |
| 1.68% | VDNKFYKEVDAAASEICSLPNLNRNQRSAFICSLYNDPSQSANLLAEAKKLNDAQAPK |
| 1.23% | VDNKFTKEGIYAISQIVRLPNLNGFQEVAFLSSLPNDPSQSANLLAEAKKLNDAQAPK |
| 1.20% | VDNKFYKEVIYAAAEIFGLPNLNVGQRSAFVCSLYDDPSQSANLLAEAKKLNDAQAPK |
| 1.10% | VDNKFNKEVCSAFNEIGNLPNLNLGQKIAFVRSLYNDPSQSANLLAEAKKLNDAQAPK |
| 12.74% | Other Sequence Less than 1% Total Reads |

**Insufficient Confidence to Predict Complex Structure (pTM)**

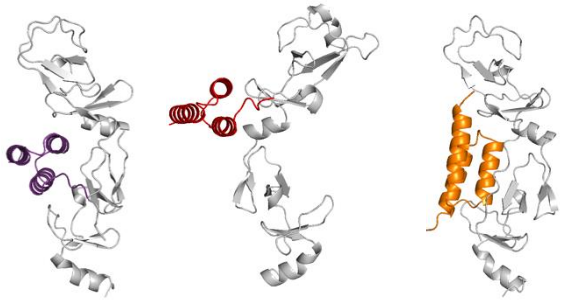

| | | | |
|---|---|---|---|
| pLDDT = 75.9 | pLDDT = 74.8 | pLDDT = 81.2 | pLDDT = 70.6 |
| pTM = 0.498 | pTM = 0.509 | pTM = 0.591 | pTM = 0.555 |
| ipTM = 0.251 | ipTM = 0.239 | ipTM = 0.631 | ipTM = 0.179 |

| Target | HIF1a |
|---|---|
| Scaffold | Affibody |
| Total HTPS Reads | 6360 |
| Unique HTPS Reads | 185 |
| Top Variant % of Total Reads | 87 |
| Final Phage Titer (PFU/mL) | 7.08E+08 |

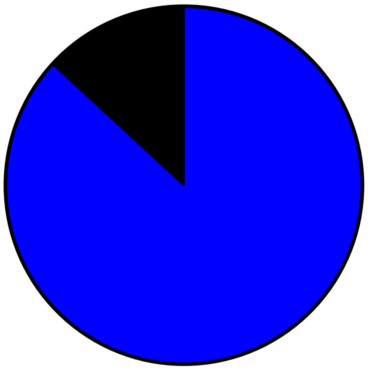

**Library: VDNKF?KE???A???I??LPNLN??Q??AF??SL??DPSQSANLLAEAKKLNDAQAPK**

**86.84% VDNKFYKEVDAAASEICSLPNLNRNQRSAFICSLYNDPSQSANLLAEAKKLNDAQAPK**

**13.16% Other Sequence Less than 1% Total Reads**

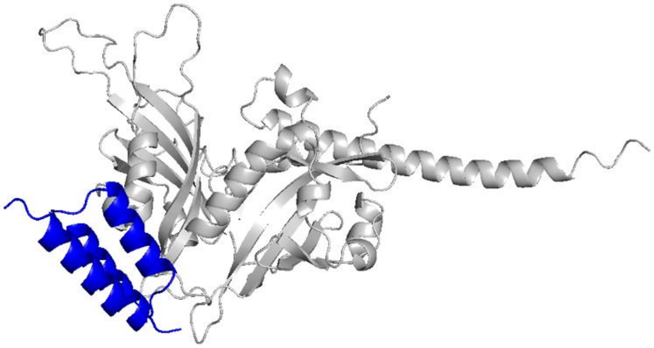

pLDDT = 72.2
pTM = 0.618
ipTM = 0.317

| Target | UEV |
|---|---|
| Scaffold | Affibody |
| Total HTPS Reads | 7537 |
| Unique HTPS Reads | 3899 |
| Top Variant % of Total Reads | 52 |
| Final Phage Titer (PFU/mL) | 5.37E+08 |

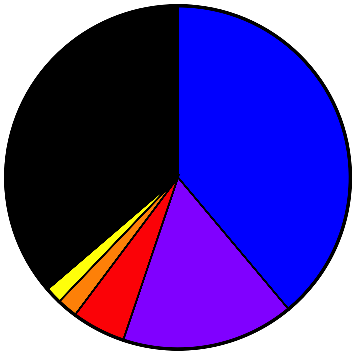

```
Library: VDNKF?KE???A???I??LPNLN??Q??AF??SL??DPSQSANLLAEAKKLNDAQAPK
38.92%   VDNKFTKEWTDAFCEISCLPNLNAHQKVAFLGSLLDDPSQSANLLAEAKKLNDAQAPK
16.23%   VDNKFNKEWIYAVNEIHRLPNLNEFQRVAFVRSLNNDPSQSANLLAEAKKLNDAQAPK
5.05%    VDNKFYKEWSDAFSEIVFLPNLNPCQRHAFLCSLLDDPSQSANLLAEAKKLNDAQAPK
1.92%    VDNKFTKEWCSAYVEIGVLPNLNDGQKNAFLYSLVDDPSQSANLLAEAKKLNDAQAPK
1.57%    VDNKFSKEVCAAVYQISSLPNLNVSQKLAFLPSLRDDPSQSANLLAEAKKLNDAQAPK
36.31%   Other Sequence less than 1%
```

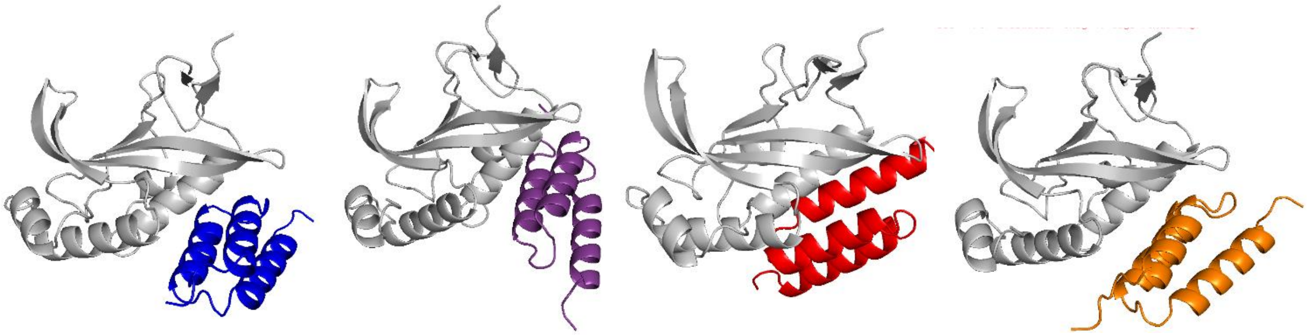

pLDDT = 85.4
pTM = 0.721
ipTM = 0.238

pLDDT = 91.1
pTM = 0.849
ipTM = 0.668

pLDDT = 86.5
pTM = 0.737
ipTM = 0.300

pLDDT = 85.8
pTM = 0.715
ipTM = 0.213

| Target | FRB |
|---|---|
| Scaffold | Affibody |
| Total HTPS Reads | 5623 |
| Unique HTPS Reads | 232 |
| Top Variant % of Total Reads | 83 |
| Final Phage Titer (PFU/mL) | 5.50E+08 |

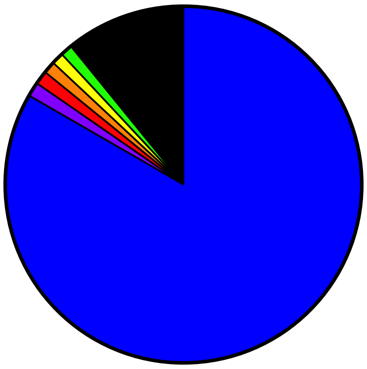

| | |
|---|---|
| Library: | VDNKF?KE???A???I??LPNLN??Q??AF??SL??DPSQSANLLAEAKKLNDAQAPK |
| 83.32% | VDNKFYKEVIYAAAEIFGLPNLNVGQRSAFVCSLYDDPSQSANLLAEAKKLNDAQAPK |
| 1.30% | VDNKFNKEVCYASSEIFYLPNLNDYQKHAFVVSLYDDPSQSANLLAEAKKLNDAQAPK |
| 1.24% | VDNKFNKEVCSAFNEIGNLPNLNLGQKIAFVRSLYNDPSQSANLLAEAKKLNDAQAPK |
| 1.08% | VDNKFYKEVIYAAAEIFGLPNLNVGQRSAFVVSLYDDPSQSANLLAEAKKLNDAQAPK |
| 1.08% | VDNKFYKEVDAAASEICSLPNLNRNQRSAFICSLYNDPSQSANLLAEAKKLNDAQAPK |
| 1.08% | VDNKFSKEGIYAISQIVRLPNLNGFQEVAFLSSLPNDPSQSANLLAEAKKLNDAQAPK |
| 10.88% | Other Sequence Less than 1% Total Reads |

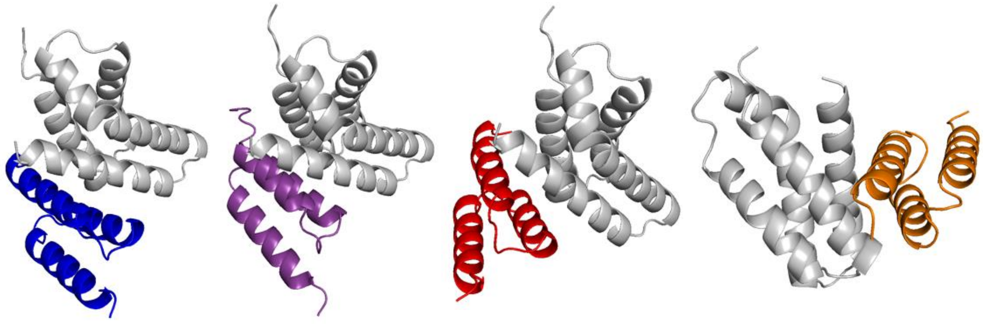

pLDDT = 92.1
pTM = 0.873
ipTM = 0.810

pLDDT = 82.9
pTM = 0.681
ipTM = 0.318

pLDDT = 85.1
pTM = 0.729
ipTM = 0.444

pLDDT = 91.1
pTM = 0.854
ipTM = 0.767

| Target | SasA |
|---|---|
| Scaffold | Affibody |
| Total HTPS Reads | 1359 |
| Unique HTPS Reads | 177 |
| Top Variant % of Total Reads | 54 |
| Final Phage Titer (PFU/mL) | 1.32E+09 |

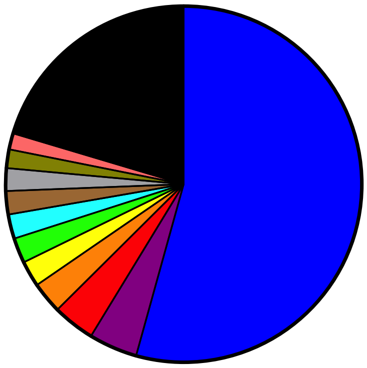

```
Library: VDNKF?KE???A???I??LPNLN??Q??AF??SL??DPSQSANLLAEAKKLNDAQAPK
54.30%   VDNKFYKEGLYAAGEIFRLPNLNGCQVVAFLSSLRDDPSQSANLLAEAKKLNDAQAPK
4.42%    VDNKFYKEVIYAAAEIFGLPNLNVGQRSAFVCSLYDDPSQSANLLAEAKKLNDAQAPK
3.83%    VDNKFNKEVCSAFNEIGNLPNLNLGQKIAFVRSLYNDPSQSANLLAEAKKLNDAQAPK
2.87%    VDNKFNKEVCSAFNEIGNLPNLNLGQKIAFVRSLYNDPSQSANLLAEAKKLNDAQAPK
2.35%    VDNKFYKEWFAAASQIFTLPNLNGVQVVAFINSLRDDPSQSANLLAEAKKLNDAQAPK
2.28%    VDNKFNKELDSAYNQICHLPNLNANQRSAFLSSLYDDPSQSANLLAEAKKLNDAQAPK
2.21%    VDNKFYKEVDAAASEICSLPNLNRNQRSAFICSLYNDPSQSANLLAEAKKLNDAQAPK
2.13%    VDNKFSKEGIYAISQIVRLPNLNGFQEVAFLSSLPNDPSQSANLLAEAKKLNDAQAPK
2.06%    VDNKFNKEWNNAAYEIFTLPNLNGCQVCAFVDSLGNDPSQSANLLAEAKKLNDAQAPK
1.69%    VDNKFTKEVCYAYIEIVYLPNLNPDQISAFISSLHDDPSQSANLLAEAKKLNDAQAPK
1.47%    VDNKFYKEGLYAAGEIFRLPNLNGCQVVAFLSSLRDDPSQSANLLAEAKKLNDAQAPK
20.38%   Other Sequence Less than 1% Total Reads
```

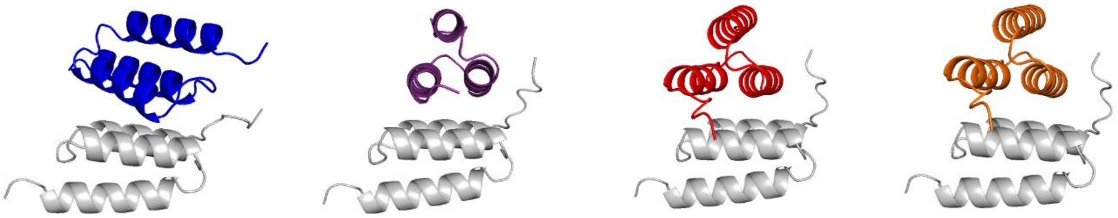

| | | | |
|---|---|---|---|
| pLDDT = 81.9<br>pTM = 0.718<br>ipTM = 0.579 | pLDDT = 80.6<br>pTM = 0.630<br>ipTM = 0.395 | pLDDT = 80.2<br>pTM = 0.619<br>ipTM = 0.372 | pLDDT = 80.2<br>pTM = 0.619<br>ipTM = 0.372 |

| Target | LC3B |
|---|---|
| Scaffold | Affibody |
| Total HTPS Reads | 7942 |
| Unique HTPS Reads | 3627 |
| Top Variant % of Total Reads | 46 |
| Final Phage Titer (PFU/mL) | 1.23E+08 |

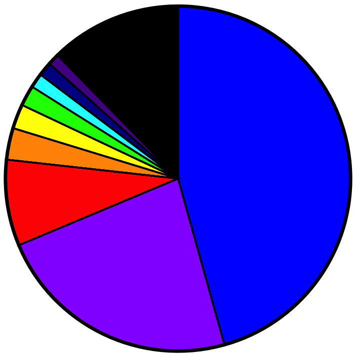

```
Library: VDNKF?KE???A???I??LPNLN??Q??AF??SL??DPSQSANLLAEAKKLNDAQAPK
45.67%   VDNKFTKEVDDASFEILYLPNLNDFQENAFLVSLSDDPSQSANLLAEAKKLNDAQAPK
23.02%   VDNKFTKEWTDAFCEISCLPNLNAHQKVAFLGSLLDDPSQSANLLAEAKKLNDAQAPK
8.07%    VDNKFNKEWIYAVNEIHRLPNLNEFQRVAFVRSLNNDPSQSANLLAEAKKLNDAQAPK
2.97%    VDNKFYKEWSDAFSEIVFLPNLNPCQRHAFLCSLLDDPSQSANLLAEAKKLNDAQAPK
2.29%    VDNKFSKEGGDADFEILSLPNLNENQDGCLHSQPG*DPSQSANLLAEAKKLNDAQAPK
1.93%    VDNKFTKEWCSAYVEIGVLPNLNDGQKNAFLYSLVDDPSQSANLLAEAKKLNDAQAPK
1.36%    VDNKFTKEWDDASFEILCLPNLNEDQGSAFISSLYNDPSQSANLLAEAKKLNDAQAPK
1.08%    VDNKFSKEVCAAVYQISSLPNLNVSQKLAFLPSLRDDPSQSANLLAEAKKLNDAQAPK
1.30%    VDNKFSKEVDNAINEIGDLPNLNVYQIIAFVRSLSDDPSQSANLLAEAKKLNDAQAPK
12.31%   Other Sequence less than 1%
```

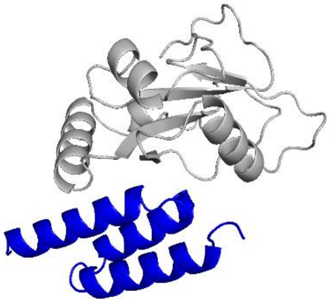

pLDDT = 85.4
pTM = 0.771
ipTM = 0.566

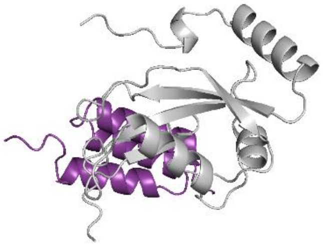

pLDDT = 79.4
pTM = 0.659
ipTM = 0.211

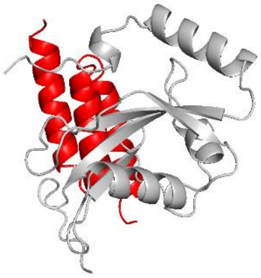

pLDDT = 75.1
pTM = 0.625
ipTM = 0.153

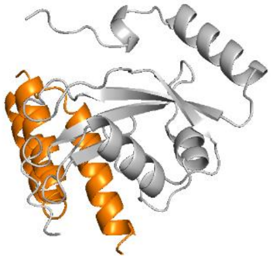

pLDDT = 76.9
pTM = 0.631
ipTM = 0.133

| | |
|---|---|
| Target | IFNG |
| Scaffold | Affibody |
| Total HTPS Reads | 9319 |
| Unique HTPS Reads | 5896 |
| Top Variant % of Total Reads | 63 |
| Final Phage Titer (PFU/mL) | 5.00E+08 |

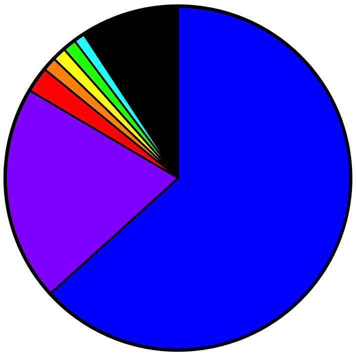

| | |
|---|---|
| Library: | VDNKF?KE???A???I??LPNLN??Q??AF??SL??DPSQSANLLAEAKKLNDAQAPK |
| 63.27% | VDNKFTKEVDSAVFEIHILPNLNDYQRVAFIFSLFDDPSQSANLLAEAKKLNDAQAPK |
| 20.25% | VDNKFYKELSTAVYEIFALPNLNESQVVAFVVSLFDDPSQSANLLAEAKKLNDAQAPK |
| 2.41% | VDNKFNKELFNAIYQIFVLPNLNEHQRVAFVISLFDDPSQSANLLAEAKKLNDAQAPK |
| 1.30% | VDNKFTKEVDSAVFEIHILPNLNESQVVAFVVSLFDDPSQSANLLAEAKKLNDAQAPK |
| 1.28% | VDNKFYKELFNAICQIFGLPNLNENQIVAFIISLFDDPSQSANLLAEAKKLNDAQAPK |
| 1.26% | VDNKFYKEVISAICQIFGLPNLNGTQRRAFLCSLGNDPSQSANLLAEAKKLNDAQAPK |
| 1.03% | VDNKFSKEVCDAIYEIHILPNLNVFQRLAFIFSLFDDPSQSANLLAEAKKLNDAQAPK |
| 9.21% | Other Sequence less than 1% |

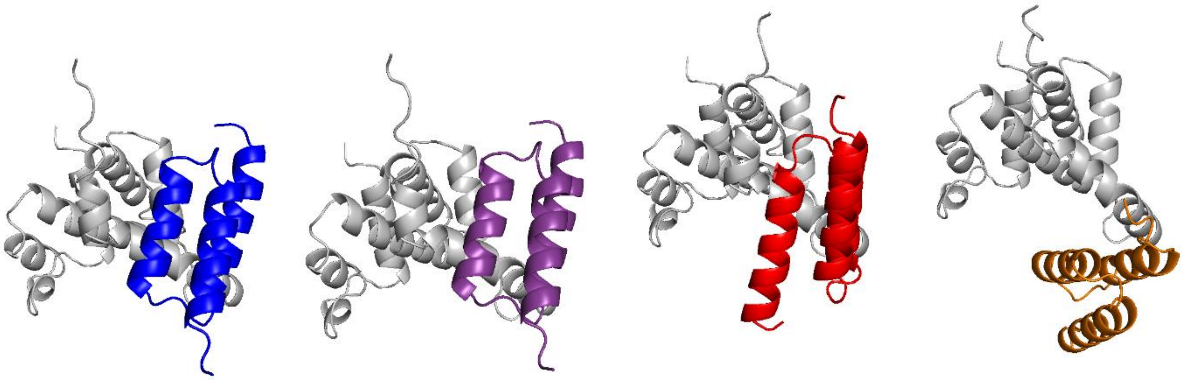

| | | | |
|---|---|---|---|
| pLDDT = 83.5 | pLDDT = 83.5 | pLDDT = 82.9 | pLDDT = 78.8 |
| pTM = 0.636 | pTM = 0.636 | pTM = 0.672 | pTM = 0.593 |
| ipTM = 0.247 | ipTM = 0.247 | ipTM = 0.378 | ipTM = 0.125 |

| Target | MST2 (313-437) |
|---|---|
| Scaffold | Affibody |
| Total HTPS Reads | 2628 |
| Unique HTPS Reads | 206 |
| Top Variant % of Total Reads | 34 |
| Final Phage Titer (PFU/mL) | 3.39E+06 |

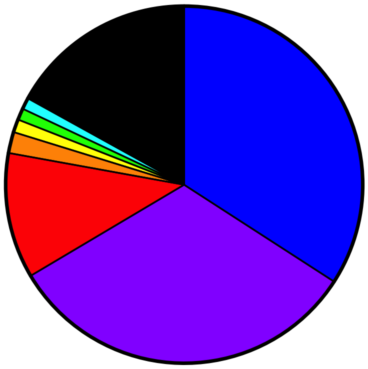

Library: VDNKF?KE???A???I??LPNLN??Q??AF??SL??DPSQSANLLAEAKKLNDAQAPK

34.13% VDNKFSKEVSAAACEIRILPNLNGSQRSAFIGSLVNDPSQSANLLAEAKKLNDAQAPK

32.34% VDNKFNKEGRSATSEIVVLPNLNGLQRDAFVVSLYDDPSQSANLLAEAKKLNDAQAPK

11.38% VDNKFTKELGYAASQISGLPNLNGFQQGAFLRSLINDPSQSANLLAEAKKLNDAQAPK

1.90% VDNKFYKEVDAAASEICSLPNLNRNQRSAFICSLYNDPSQSANLLAEAKKLNDAQAPK

1.18% VDNKFNKEVCSAFNEIGNLPNLNLGQKIAFVRSLYNDPSQSANLLAEAKKLNDAQAPK

1.07% VDNKFYKEVIYAAAEIFGLPNLNVGQRSAFVCSLYDDPSQSANLLAEAKKLNDAQAPK

1.03% VDNKFSKEGIYAISQIVRLPNLNGFQEVAFLSSLPNDPSQSANLLAEAKKLNDAQAPK

16.97% Other Sequence Less than 1% Total Reads

| | | | |
|---|---|---|---|
| **Insufficient Confidence to Predict Complex Structure (pLDDT, pTM)** | **Insufficient Confidence to Predict Complex Structure (pLDDT, pTM)** | **Insufficient Confidence to Predict Complex Structure (pLDDT, pTM)** | **Insufficient Confidence to Predict Complex Structure (pLDDT, pTM)** |
| pLDDT = 48.3<br>pTM = 0.335<br>ipTM = 0.205 | pLDDT = 45.8<br>pTM = 0.326<br>ipTM = 0.184 | pLDDT = 45.0<br>pTM = 0.323<br>ipTM = 0.146 | pLDDT = 47.4<br>pTM = 0.346<br>ipTM = 0.210 |

| Target | BAX Peptide |
|---|---|
| Scaffold | Affibody |
| Total HTPS Reads | 3576 |
| Unique HTPS Reads | 287 |
| Top Variant % of Total Reads | 55 |
| Final Phage Titer (PFU/mL) | 6.31E+08 |

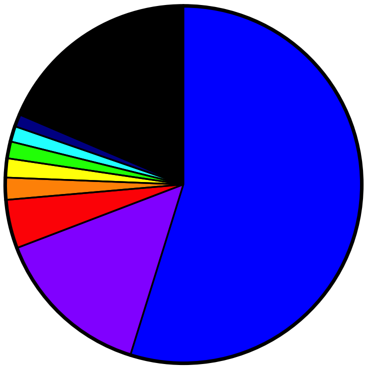

| | |
|---|---|
| Library: | VDNKF?KE???A???I??LPNLN??Q??AF??SL??DPSQSANLLAEAKKLNDAQAPK |
| 54.81% | VDNKFNKELDSAYNQICHLPNLNANQRSAFLSSLYDDPSQSANLLAEAKKLNDAQAPK |
| 14.40% | VDNKFNKEGDYAVFQICGLPNLNEGQGAAFVGSLHNDPSQSANLLAEAKKLNDAQAPK |
| 4.42% | VDNKFTKEGGYADLEIAFLPNLNDDQRFAFLPSLTNDPSQSANLLAEAKKLNDAQAPK |
| 1.99% | VDNKFNKEVCSAFNEIGNLPNLNLGQKIAFVRSLYNDPSQSANLLAEAKKLNDAQAPK |
| 1.73% | VDNKFYKEVDAAASEICSLPNLNRNQRSAFICSLYNDPSQSANLLAEAKKLNDAQAPK |
| 1.54% | VDNKFYKEVIYAAAEIFGLPNLNVGQRSAFVCSLYDDPSQSANLLAEAKKLNDAQAPK |
| 1.40% | VDNKFNKEVHYATNEIGCLPNLNRYQRSAFILSLHDDPSQSANLLAEAKKLNDAQAPK |
| 1.12% | VDNKFSKEGIYAISQIVRLPNLNGFQEVAFLSSLPNDPSQSANLLAEAKKLNDAQAPK |
| 18.60% | Other Sequence Less than 1% Total Reads |

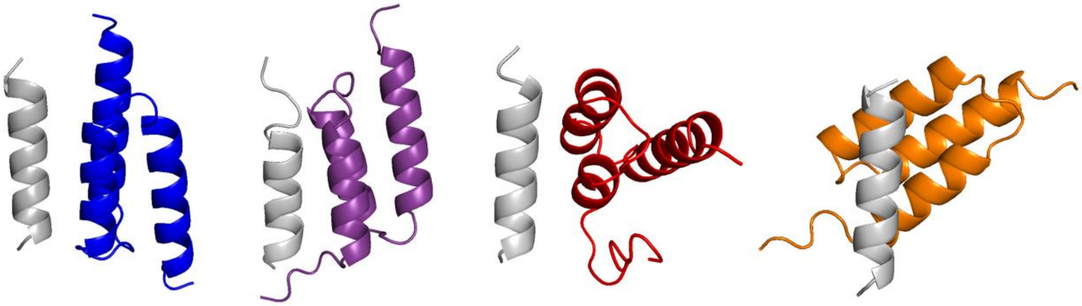

| | | | |
|---|---|---|---|
| pLDDT = 76.2 | pLDDT = 72.7 | pLDDT = 69.2 | pLDDT = 81.2 |
| pTM = 0.668 | pTM = 0.675 | pTM = 0.595 | pTM = 0.709 |
| ipTM = 0.249 | ipTM = 0.449 | ipTM = 0.404 | ipTM = 0.450 |

| Target | KRAS G12D |
|---|---|
| Scaffold | Affitin |
| Total HTPS Reads | 10736 |
| Unique HTPS Reads | 287 |
| Top Variant % of Total Reads | 84 |
| Final Phage Titer (PFU/mL) | 8.13E+08 |

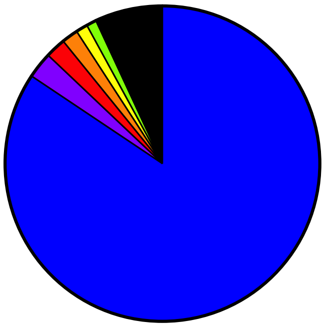

```
Library: GSVKVKF???GEEKEVDTSKI??V?R?GK?V?F?YDDNGK?G?G?V?EKDAPKELLDMLARAEREKKL
 84.36%  GSVKVKFIFSGEEKEVDTSKIHAVFRNGKYVTFFYDDNGKHGFGSVLEKDAPKELLDMLARAEREKKL
  2.69%  GSVKVKFATFGEEKEVDTSKIHDVDRYGKTVHFIYDDNGKCGLGYVNEKDAPKELLDMLARAEREKKL
  2.09%  GSVKVKFHYRGEEKEVDTSKIVHVHRDGKFVLFVYDDNGKNGVGGVYEKDAPKELLDMLARAEREKKL
  1.69%  GSVKVKFSYHGEEKEVDTSKINNVNRHGKHVLFDYDDNGKIGSGLVFEKDAPKELLDMLARAEREKKL
  1.20%  GSVKVKFVDFGEEKEVDTSKIRLVIRFGKSVHFIYDDNGKDGYGIVNEKDAPKELLDMLARAEREKKL
  1.03%  GSVKVKFRRPGEEKEVDTSKIGYVHRVGKFVAFFYDDNGKIGSGTVSEKDAPKELLDMLARAEREKKL
  6.94%  Other Sequence Less than 1% Tota
```

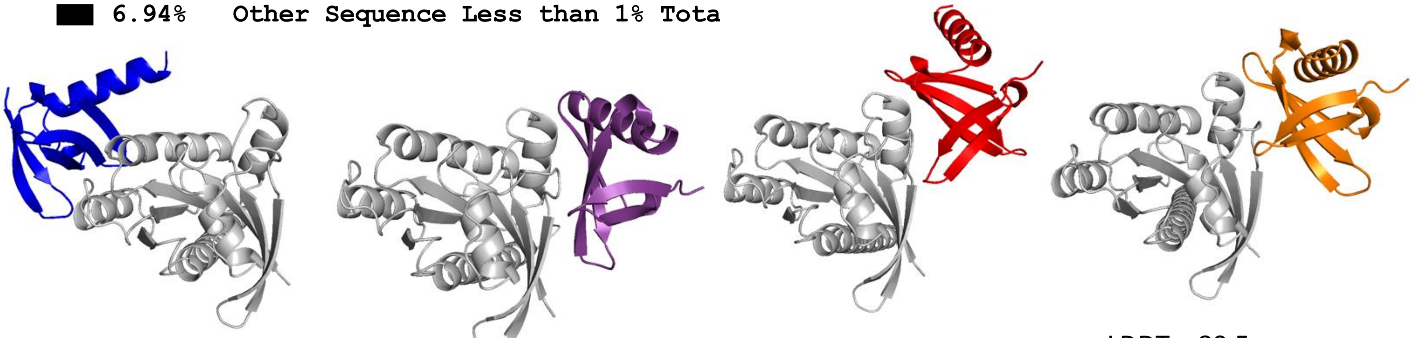

pLDDT = 87.4
pTM = 0.850
ipTM = 0.821

pLDDT = 78.8
pTM = 0.662
ipTM = 0.162

pLDDT = 82.1
pTM = 0.676
ipTM = 0.206

pLDDT = 80.5
pTM = 0.661
ipTM = 0.158

| Target | ZB |
| --- | --- |
| Scaffold | Affitin |
| Total HTPS Reads | 18028 |
| Unique HTPS Reads | 478 |
| Top Variant % of Total Reads | 85 |
| Final Phage Titer (PFU/mL) | 2.57E+08 |

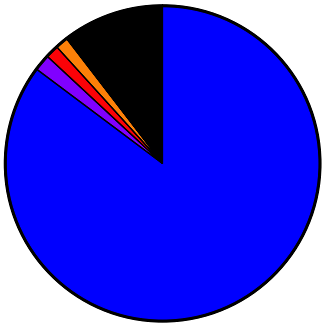

```
Library: GSVKVKF???GEEKEVDTSKI??V?R?GK?V?F?YDDNGK?G?G?V?EKDAPKELLDMLARAEREKKL
85.21%   GSVKVKFATFGEEKEVDTSKIHDVDRYGKTVHFIYDDNGKCGLGYVNEKDAPKELLDMLARAEREKKL
1.68%    GSVKVKFHYRGEEKEVDTSKIVHVHRDGKFVLFVYDDNGKNGVGGVYEKDAPKELLDMLARAEREKKL
1.35%    GSVKVKFIFSGEEKEVDTSKIHAVFRNGKYVTFFYDDNGKHGFGSVLEKDAPKELLDMLARAEREKKL
1.31%    GSVKVKFSYHGEEKEVDTSKINNVNRHGKHVLFDYDDNGKIGSGLVFEKDAPKELLDMLARAEREKKL
10.44%   Other Sequence Less than 1% Total Reads
```

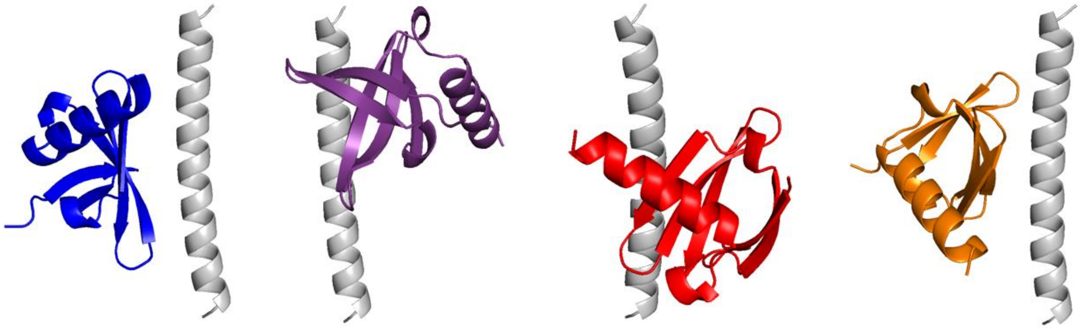

| | | | |
| --- | --- | --- | --- |
| pLDDT = 76.4 | pLDDT = 91.4 | pLDDT = 74.8 | pLDDT = 75.4 |
| pTM = 0.610 | pTM = 0.808 | pTM = 0.602 | pTM = 0.597 |
| ipTM = 0.193 | ipTM = 0.759 | ipTM = 0.197 | ipTM = 0.182 |

| | |
|---|---|
| Target | p65 (17-293) |
| Scaffold | Affitin |
| Total HTPS Reads | 10100 |
| Unique HTPS Reads | 309 |
| Top Variant % of Total Reads | 79 |
| Final Phage Titer (PFU/mL) | 1.51E+09 |

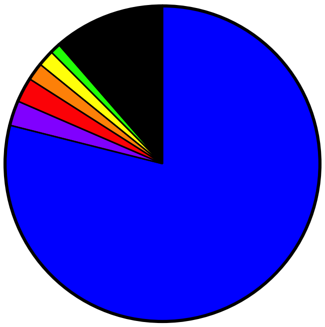

Library: GSVKVKF???GEEKEVDTSKI??V?R?GK?V?F?YDDNGK?G?G?V?EKDAPKELLDMLARAEREKKL

78.92% GSVKVKFVDFGEEKEVDTSKIRLVIRFGKSVHFIYDDNGKDGYGIVNEKDAPKELLDMLARAEREKKL

2.56% GSVKVKFHYRGEEKEVDTSKIVHVHRDGKFVLFVYDDNGKNGVGGVYEKDAPKELLDMLARAEREKKL

2.53% GSVKVKFATFGEEKEVDTSKIHDVDRYGKTVHFIYDDNGKCGLGYVNEKDAPKELLDMLARAEREKKL

1.83% GSVKVKFSYHGEEKEVDTSKINNVNRHGKHVLFDYDDNGKIGSGLVFEKDAPKELLDMLARAEREKKL

1.66% GSVKVKFIFSGEEKEVDTSKIHAVFRNGKYVTFFYDDNGKHGFGSVLEKDAPKELLDMLARAEREKKL

1.07% GSVKVKFFILGEEKEVDTSKIPHVLRYGKHVHFIYDDNGKHGYGTVPEKDAPKELLDMLARAEREKKL

11.42% Other Sequence Less than 1% Total Reads

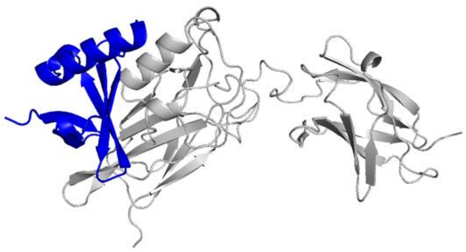

pLDDT = 83.9
pTM = 0.694
ipTM = 0.132

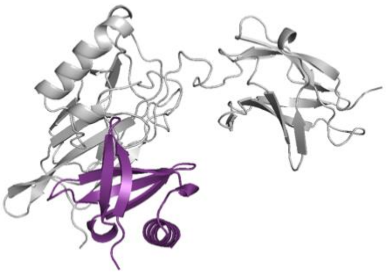

pLDDT = 84.4
pTM = 0.708
ipTM = 0.131

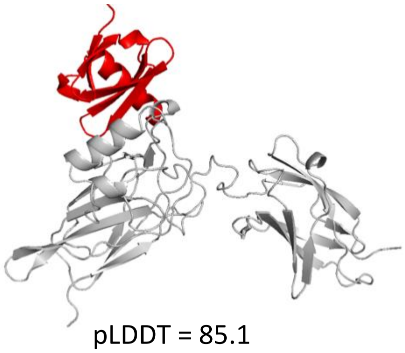

pLDDT = 85.1
pTM = 0.570
ipTM = 0.257

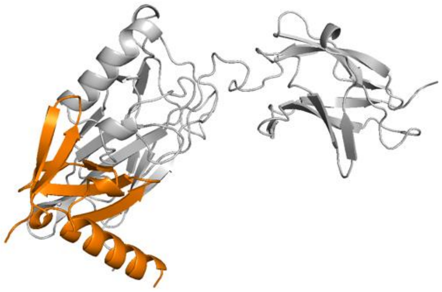

pLDDT = 83.8
pTM = 0.699
ipTM = 0.122

| Target | HPV-pE7 |
|---|---|
| Scaffold | Affitin |
| Total HTPS Reads | 19911 |
| Unique HTPS Reads | 1071 |
| Top Variant % of Total Reads | 82 |
| Final Phage Titer (PFU/mL) | 2.14E+07 |

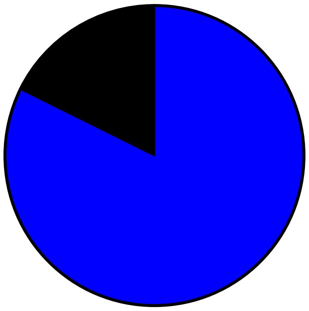

**Library:** GSVKVKF???GEEKEVDTSKI??V?R?GK?V?F?YDDNGK?G?G?V?EKDAPKELLDMLARAEREKKL

82.30% GSVKVKFHGTGEEKEVDTSKISVVIRAGKTVYFSYDDNGKRGTGRVSEKDAPKELLDMLARAEREKKL

17.70% Other Sequence Less than 1% Total Reads

**Insufficient Confidence to Predict Complex Structure (pTM)**

pLDDT = 57.9
pTM = 0.415
ipTM = 0.165

| Target | FGFR1c domain D2 |
|---|---|
| Scaffold | Affitin |
| Total HTPS Reads | 16959 |
| Unique HTPS Reads | 379 |
| Top Variant % of Total Reads | 87 |
| Final Phage Titer (PFU/mL) | 1.55E+09 |

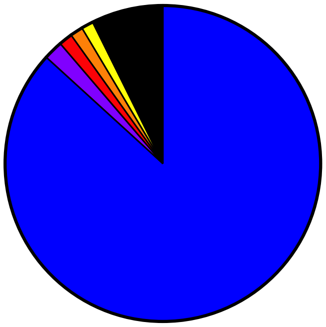

```
Library: GSVKVKF???GEEKEVDTSKI??V?R?GK?V?F?YDDNGK?G?G?V?EKDAPKELLDMLARAEREKKL
86.75%   GSVKVKFHYRGEEKEVDTSKIVHVHRDGKFVLFVYDDNGKNGVGGVYEKDAPKELLDMLARAEREKKL
1.98%    GSVKVKFATFGEEKEVDTSKIHDVDRYGKTVHFIYDDNGKCGLGYVNEKDAPKELLDMLARAEREKKL
1.40%    GSVKVKFSYHGEEKEVDTSKINNVNRHGKHVLFDYDDNGKIGSGLVFEKDAPKELLDMLARAEREKKL
1.31%    GSVKVKFIFSGEEKEVDTSKIHAVFRNGKYVTFFYDDNGKHGFGSVLEKDAPKELLDMLARAEREKKL
1.23%    GSVKVKFVDFGEEKEVDTSKIRLVIRFGKSVHFIYDDNGKDGYGIVNEKDAPKELLDMLARAEREKKL
7.34%    Other Sequence Less than 1% Total Reads
```

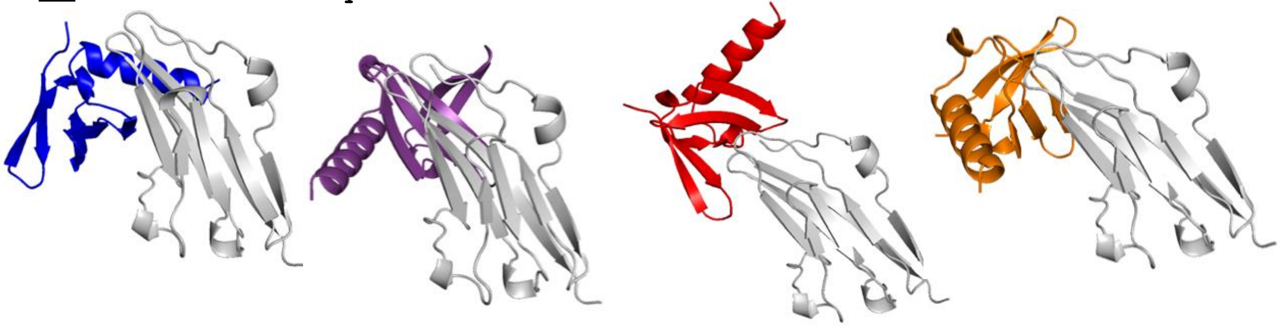

| pLDDT = 81.6<br>pTM = 0.586<br>ipTM = 0.141 | pLDDT = 81.6<br>pTM = 0.570<br>ipTM = 0.134 | pLDDT = 82.9<br>pTM = 0.567<br>ipTM = 0.117 | pLDDT = 80.8<br>pTM = 0.566<br>ipTM = 0.126 |
|---|---|---|---|

| | |
|---|---|
| Target | TCIM |
| Scaffold | Affitin |
| Total HTPS Reads | 15930 |
| Unique HTPS Reads | 996 |
| Top Variant % of Total Reads | 80 |
| Final Phage Titer (PFU/mL) | 8.71E+08 |

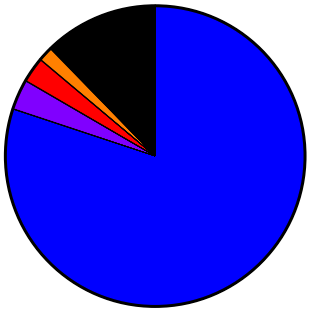

```
Library: GSVKVKF???GEEKEVDTSKI??V?R?GK?V?F?YDDNGK?G?G?V?EKDAPKELLDMLARAEREKKL
80.08%   GSVKVKFPSLGEEKEVDTSKILLVLRAGKDVLFFYDDNGKLGGGIVTEKDAPKELLDMLARAEREKKL
3.30%    GSVKVKF***GEEKEVDTSKIRSVWRGGKWVDFSYDDNGKQGYGYVEKDAPKELLDMLARAEREKKLL
2.73%    GSVKVKFLTHGEEKEVDTSKINAVHRTGKRVYFIYDDNGKPGFGFVFEKDAPKELLDMLARAEREKKL
1.58%    GSVKVKFSYAGEEKEVDTSKIYHVSRLGKDVIFCYDDNGKVGSGAVFEKDAPKELLDMLARAEREKKL
12.32%   Other Sequence Less than 1% Total Reads
```

| | | |
|---|---|---|
| **Insufficient Confidence to Predict Complex Structure (pTM)** | **Insufficient Confidence to Predict Complex Structure (pTM)** | **Insufficient Confidence to Predict Complex Structure (pTM)** |
| pLDDT = 51.1<br>pTM = 0.417<br>ipTM = 0.178 | pLDDT = 50.2<br>pTM = 0.417<br>ipTM = 0.202 | pLDDT = 50.4<br>pTM = 0.424<br>ipTM = 0.175 |

| Target | PDE6D |
|---|---|
| Scaffold | Affitin |
| Total HTPS Reads | 13640 |
| Unique HTPS Reads | 620 |
| Top Variant % of Total Reads | 81 |
| Final Phage Titer (PFU/mL) | 1.45E+09 |

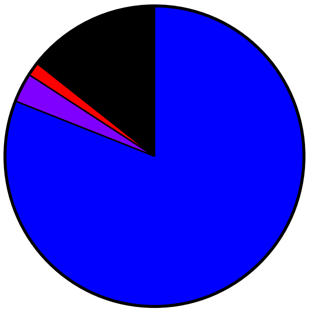

Library: GSVKVKF???GEEKEVDTSKI??V?R?GK?V?F?YDDNGK?G?G?V?EKDAPKELLDMLARAEREKKL

80.99% GSVKVKFLTHGEEKEVDTSKINAVHRTGKRVYFIYDDNGKPGFGFVFEKDAPKELLDMLARAEREKKL

3.15% GSVKVKFPSLGEEKEVDTSKILLVLRAGKDVLFFYDDNGKLGGGIVTEKDAPKELLDMLARAEREKKL

1.47% GSVKVKFSYAGEEKEVDTSKIYHVSRLGKDVIFCYDDNGKVGSGAVFEKDAPKELLDMLARAEREKKL

14.39% Other Sequence Less than 1% Total Reads

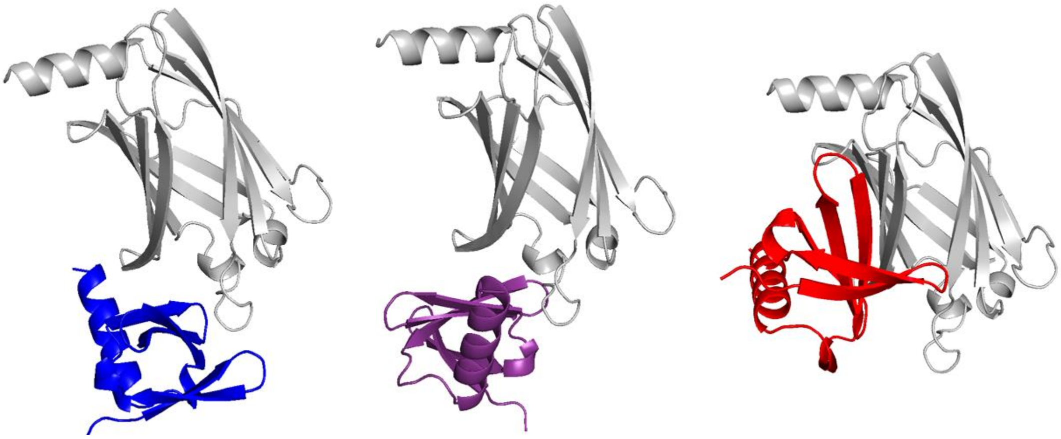

pLDDT = 91.5
pTM = 0.828
ipTM = 0.675

pLDDT = 85.3
pTM = 0.673
ipTM = 0.192

pLDDT = 82.1
pTM = 0.658
ipTM = 0.141

| | |
|---|---|
| Target | RAF RBD |
| Scaffold | Affitin |
| Total HTPS Reads | 11043 |
| Unique HTPS Reads | 804 |
| Top Variant % of Total Reads | 60 |
| Final Phage Titer (PFU/mL) | 4.47E+08 |

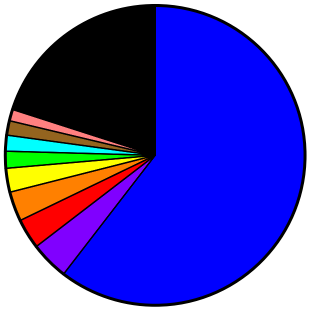

Library: GSVKVKF???GEEKEVDTSKI??V?R?GK?V?F?YDDNGK?G?G?V?EKDAPKELLDMLARAEREKKL

60.46% GSVKVKFSYAGEEKEVDTSKIYHVSRLGKDVIFCYDDNGKVGSGAVFEKDAPKELLDMLARAEREKKL

4.13% GSVKVKFSYHGEEKEVDTSKINGVSRCGKDVIFNYDDNGKSGLGAVYEKDAPKELLDMLARAEREKKL

3.26% GSVKVKFLTHGEEKEVDTSKINAVHRTGKRVYFIYDDNGKPGFGFVFEKDAPKELLDMLARAEREKKL

3.20% GSVKVKFPSLGEEKEVDTSKILLVLRAGKDVLFFYDDNGKLGGGIVTEKDAPKELLDMLARAEREKKL

2.57% GSVKVKF***GEEKEVDTSKIRSVWRGGKWVDFSYDDNGKQGYGYVLEKDAPKELLDMLARAEREKKL

1.82% GSVKVKFSYAGEEKEVDTSKIYHVSRLGKDVIFCYDDNGKSGLGAVYEKDAPKELLDMLARAEREKKL

1.72% GSVKVKFSYHGEEKEVDTSKIYHVSRLGKDVIFCYDDNGKVGSGAVFEKDAPKELLDMLARAEREKKL

1.55% GSVKVKFSYAGEEKEVDTSKINGVSRCGKDVIFNYDDNGKSGLGAVYEKDAPKELLDMLARAEREKKL

1.24% GSVKVKFSYAGEEKEVDTSKIYHVSRLGKDVIFCYDDNGKVGSGAVYEKDAPKELLDMLARAEREKKL

20.05% Other Sequence Less than 1% Total Reads

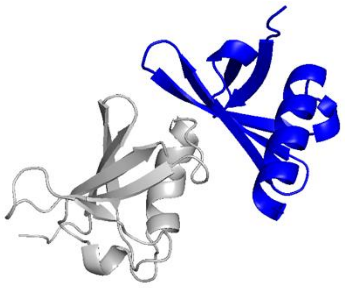

pLDDT = 89.6
pTM = 0.714
ipTM = 0.534

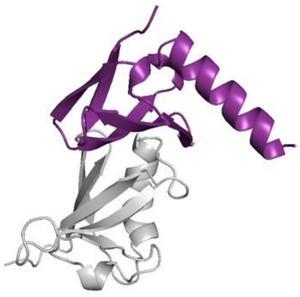

pLDDT = 87.6
pTM = 0.691
ipTM = 0.544

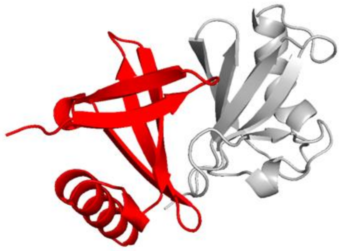

pLDDT = 79.6
pTM = 0.488
ipTM = 0.090

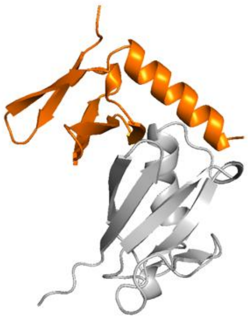

pLDDT = 81.1
pTM = 0.531
ipTM = 0.207

| Target | NRAS |
|---|---|
| Scaffold | Affitin |
| Total HTPS Reads | 12686 |
| Unique HTPS Reads | 371 |
| Top Variant % of Total Reads | 84 |
| Final Phage Titer (PFU/mL) | 5.75E+08 |

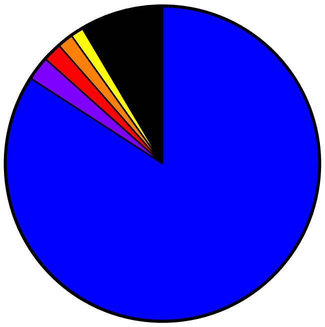

| | | |
|---|---|---|
| Library: | | GSVKVKF???GEEKEVDTSKI??V?R?GK?V?F?YDDNGK?G?G?V?EKDAPKELLDMLARAEREKKL |
| (blue) | 84.13% | GSVKVKFSYHGEEKEVDTSKINNVNRHGKHVLFDYDDNGKIGSGLVFEKDAPKELLDMLARAEREKKL |
| (purple) | 2.46% | GSVKVKFATFGEEKEVDTSKIHDVDRYGKTVHFIYDDNGKCGLGYVNEKDAPKELLDMLARAEREKKL |
| (red) | 1.93% | GSVKVKFHYRGEEKEVDTSKIVHVHRDGKFVLFVYDDNGKNGVGGVYEKDAPKELLDMLARAEREKKL |
| (orange) | 1.62% | GSVKVKFIFSGEEKEVDTSKIHAVFRNGKYVTFFYDDNGKHGFGSVLEKDAPKELLDMLARAEREKKL |
| (yellow) | 1.32% | GSVKVKFVDFGEEKEVDTSKIRLVIRFGKSVHFIYDDNGKDGYGIVNEKDAPKELLDMLARAEREKKL |
| (black) | 8.54% | Other Sequence Less than 1% Total Reads |

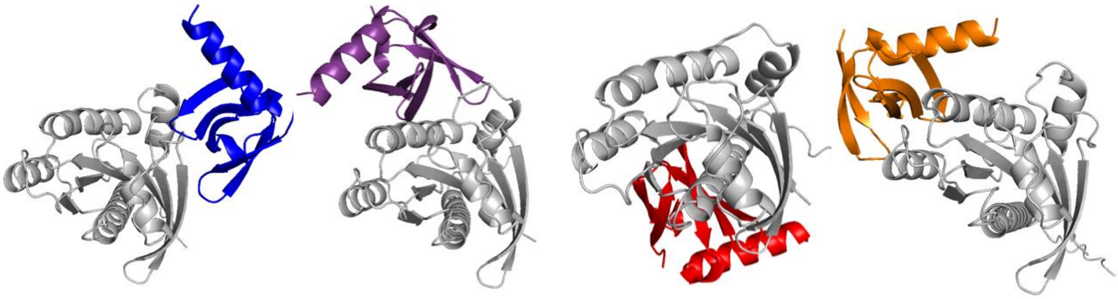

pLDDT = 80.9
pTM = 0.661
ipTM = 0.144

pLDDT = 78.8
pTM = 0.661
ipTM = 0.182

pLDDT = 85.4
pTM = 0.803
ipTM = 0.638

pLDDT = 78.1
pTM = 0.672
ipTM = 0.230

| Target | Beclin (175-267) |
|---|---|
| Scaffold | Affitin |
| Total HTPS Reads | 5073 |
| Unique HTPS Reads | 690 |
| Top Variant % of Total Reads | 14 |
| Final Phage Titer (PFU/mL) | 2.15E+08 |

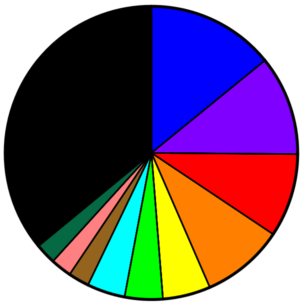

Library: GSVKVKF???GEEKEVDTSKI??V?R?GK?V?F?YDDNGK?G?G?V?EKDAPKELLDMLARAEREKKL

14.13% GSVKVKFIPFGEEKEVDTSKILIVNRLGKNVTFIYDDNGKRGYGFVSEKDAPKELLDMLARAEREKKL

11.00% GSVKVKFSSYGEEKEVDTSKILRVVRCGKHVGFLYDDNGKTGFGVVSEKDAPKELLDMLARAEREKKL

9.23% GSVKVKFCPLGEEKEVDTSKILNVNRCGKCVVFIYDDNGKSGTGHVSEKDAPKELLDMLARAEREKKL

9.11% GSVKVKFIPVGEEKEVDTSKIHAVHRAGKIVLFRYDDNGKNGSGFVLEKDAPKELLDMLARAEREKKL

5.28% GSVKVKFCNLGEEKEVDTSKIALVNRVGKCVFFRYDDNGKTGFGAVCEKDAPKELLDMLARAEREKKL

4.20% GSVKVKFTVHGEEKEVDTSKILRVTRVGKHVGFLYDDNGKIGTGIVTEKDAPKELLDMLARAEREKKL

4.20% GSVKVKFDFDGEEKEVDTSKITFVRRVGKLVFFVYDDNGKTGFGPVGEKDAPKELLDMLARAEREKKL

2.31% GSVKVKFAFRGEEKEVDTSKISHVSRRGKLVGFLYDDNGKTGIGIVAEKDAPKELLDMLARAEREKKL

2.31% GSVKVKFVLHGEEKEVDTSKIHHVVRSGKHVAFSYDDNGKPGYGVVYEKDAPKELLDMLARAEREKKL

2.29% GSVKVKFPVSGEEKEVDTSKIIFVRRIGKHVFFTYDDNGKYGAGAVLEKDAPKELLDMLARAEREKKL

35.96% Other Sequence less than 1%

| Insufficient Confidence to Predict Complex Structure (pTM) | Insufficient Confidence to Predict Complex Structure (pTM) | Insufficient Confidence to Predict Complex Structure (pTM) | Insufficient Confidence to Predict Complex Structure (pTM) |
|---|---|---|---|
| pLDDT = 68.6<br>pTM = 0.448<br>ipTM = 0.160 | pLDDT = 67.6<br>pTM = 0.490<br>ipTM = 0.229 | pLDDT = 63.9<br>pTM = 0.425<br>ipTM = 0.123 | pLDDT = 65.2<br>pTM = 0.449<br>ipTM = 0.232 |

| Target | MDM2 (1-188) |
|---|---|
| Scaffold | Affitin |
| Total HTPS Reads | 8067 |
| Unique HTPS Reads | 380 |
| Top Variant % of Total Reads | 61 |
| Final Phage Titer (PFU/mL) | 1.05E+09 |

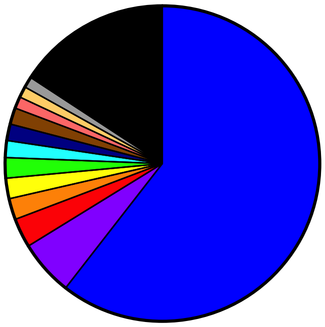

```
Library: GSVKVKF???GEEKEVDTSKI??V?R?GK?V?F?YDDNGK?G?G?V?EKDAPKELLDMLARAEREKKL
60.54%   GSVKVKFPCRGEEKEVDTSKIVHVSRYGKYVIFLYDDNGKHGFGHVAEKDAPKELLDMLARAEREKKL
5.70%    GSVKVKFTIAGEEKEVDTSKITPVARNGKSVCFLYDDNGKSGIGIVAEKDAPKELLDMLARAEREKKL
3.01%    GSVKVKFLFPGEEKEVDTSKIHSVDRAGKFVAFYYDDMVNAVLVMFTEKDAPKELLDMLARAEREKKL
2.16%    GSVKVKFYDNGEEKEVDTSKIICVIRFGKNVLFDYDDNGKPGDGIVLEKDAPKELLDMLARAEREKKL
2.11%    GSVKVKFHYRGEEKEVDTSKIVHVHRDGKFVLFVYDDNGKNGVGGVYEKDAPKELLDMLARAEREKKL
2.08%    GSVKVKFATFGEEKEVDTSKIHDVDRYGKTVHFIYDDNGKCGLGYVNEKDAPKELLDMLARAEREKKL
1.74%    GSVKVKFTIAGEEKEVDTSKIVHVSRYGKYVIFLYDDNGKHGFGHVAEKDAPKELLDMLARAEREKKL
1.70%    GSVKVKFPCRGEEKEVDTSKITPVARNGKSVCFLYDDNGKSGIGIVAEKDAPKELLDMLARAEREKKL
1.69%    GSVKVKFSYHGEEKEVDTSKINNVNRHGKHVLFDYDDNGKIGSGLVFEKDAPKELLDMLARAEREKKL
1.21%    GSVKVKFVDFGEEKEVDTSKIRLVIRFGKSVHFIYDDNGKDGYGIVNEKDAPKELLDMLARAEREKKL
1.13%    GSVKVKFIFSGEEKEVDTSKIHAVFRNGKYVTFFYDDNGKHGFGSVLEKDAPKELLDMLARAEREKKL
1.13%    GSVKVKFRRPGEEKEVDTSKIGYVHRVGKFVAFFYDDNGKIGSGTVSEKDAPKELLDMLARAEREKKL
15.81%   Other Sequence Less than 1% Total Reads
```

| | | | |
|---|---|---|---|
| **Insufficient Confidence to Predict Complex Structure (pTM)** | **Insufficient Confidence to Predict Complex Structure (pTM)** | **Insufficient Confidence to Predict Complex Structure (pTM)** | **Insufficient Confidence to Predict Complex Structure (pTM)** |
| pLDDT = 59.8<br>pTM = 0.448<br>ipTM = 0.330 | pLDDT = 60.7<br>pTM = 0.481<br>ipTM = 0.443 | pLDDT = 61.4<br>pTM = 0.443<br>ipTM = 0.277 | pLDDT = 61.0<br>pTM = 0.470<br>ipTM = 0.381 |

| | |
|---|---|
| Target | Calprotectin heterodimer 12aa linker |
| Scaffold | Affitin |
| Total HTPS Reads | 8206 |
| Unique HTPS Reads | 604 |
| Top Variant % of Total Reads | 68 |
| Final Phage Titer (PFU/mL) | 7.08E+08 |

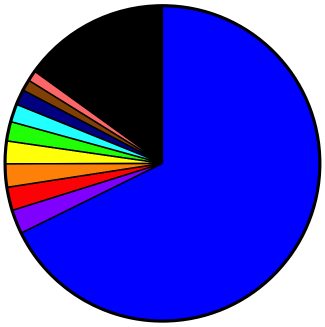

```
Library: GSVKVKF???GEEKEVDTSKI??V?R?GK?V?F?YDDNGK?G?G?V?EKDAPKELLDMLARAEREKKL
67.71%   GSVKVKFRRPGEEKEVDTSKIGYVHRVGKFVAFFYDDNGKIGSGTVSEKDAPKELLDMLARAEREKKL
2.47%    GSVKVKF***GEEKEVDTSKIRSVWRGGKWVDFSYDDNGKQGYGYVLEKDAPKELLDMLARAEREKKL
2.39%    GSVKVKFHYRGEEKEVDTSKIVHVHRDGKFVLFVYDDNGKNGVGGVYEKDAPKELLDMLARAEREKKL
2.38%    GSVKVKFVLSGEEKEVDTSKIGYVHRVGKFVAFFYDDNGKIGSGTVSEKDAPKELLDMLARAEREKKL
2.34%    GSVKVKFATFGEEKEVDTSKIHDVDRYGKTVHFIYDDNGKCGLGYVNEKDAPKELLDMLARAEREKKL
1.99%    GSVKVKFVLSGEEKEVDTSKIIRVNRFGKVVYFSYDDNGKSGFGYVSEKDAPKELLDMLARAEREKKL
1.84%    GSVKVKFSYHGEEKEVDTSKINNVNRHGKHVLFDYDDNGKIGSGLVFEKDAPKELLDMLARAEREKKL
1.58%    GSVKVKFIFSGEEKEVDTSKIHAVFRNGKYVTFFYDDNGKHGFGSVLEKDAPKELLDMLARAEREKKL
1.15%    GSVKVKFVDFGEEKEVDTSKIRLVIRFGKSVHFIYDDNGKDGYGIVNEKDAPKELLDMLARAEREKKL
1.08%    GSVKVKFPCRGEEKEVDTSKIVHVSRYGKYVIFLYDDNGKHGFGHVAEKDAPKELLDMLARAEREKKL
15.07%   Other Sequence Less than 1% Total Reads
```

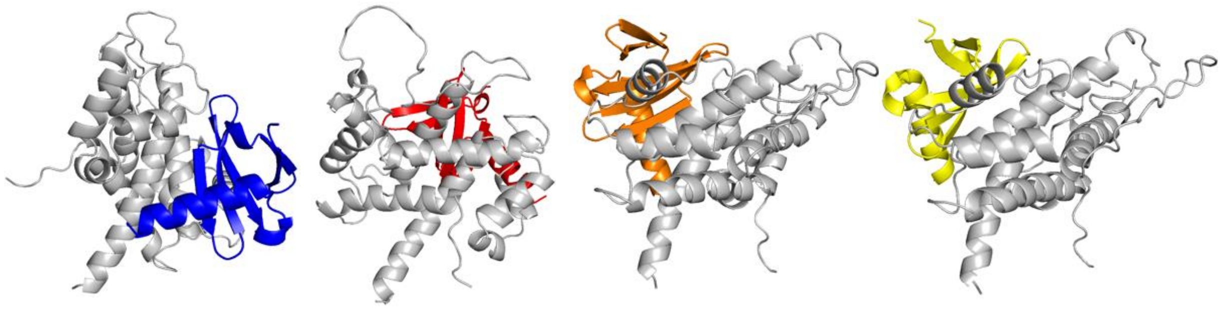

| | | | |
|---|---|---|---|
| pLDDT = 83.2 | pLDDT = 75.8 | pLDDT = 83.8 | pLDDT = 76.4 |
| pTM = 0.830 | pTM = 0.683 | pTM = 0.838 | pTM = 0.713 |
| ipTM = 0.785 | ipTM = 0.169 | ipTM = 0.792 | ipTM = 0.323 |

| Target | HlgA Toxin |
|---|---|
| Scaffold | Affitin |
| Total HTPS Reads | 10710 |
| Unique HTPS Reads | 798 |
| Top Variant % of Total Reads | 69 |
| Final Phage Titer (PFU/mL) | 1.70E+08 |

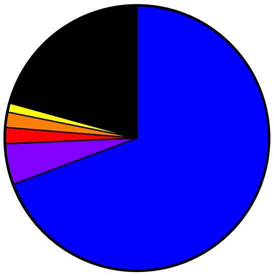

```
Library: GSVKVKF???GEEKEVDTSKI??V?R?GK?V?F?YDDNGK?G?G?V?EKDAPKELLDMLARAEREKKL
69.31%   GSVKVKFPHSGEEKEVDTSKITYVDRYGKVVVFTYDDNGKDGFGTVIEKDAPKELLDMLARAEREKKL
5.00%    GSVKVKFTSSGEEKEVDTSKITLVFRDGKTVFFAYDDNGKTGNGHVPEKDAPKELLDMLARAEREKKL
1.99%    GSVKVKFTSSGEEKEVDTSKITYVDRYGKVVVFTYDDNGKDGFGTVIEKDAPKELLDMLARAEREKKL
1.86%    GSVKVKFPHSGEEKEVDTSKITLVFRDGKTVFFAYDDNGKTGNGHVPEKDAPKELLDMLARAEREKKL
1.11%    GSVKVKFRIDGEEKEVDTSKICYVYRIGKAVIFIYDDNGKRGLGLVFEKDAPKELLDMLARAEREKKL
20.74%   Other Sequence Less than 1% Total Reads
```

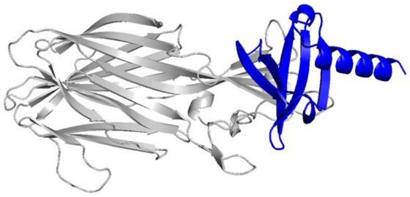

pLDDT = 90.6
pTM = 0.883
ipTM = 0.750

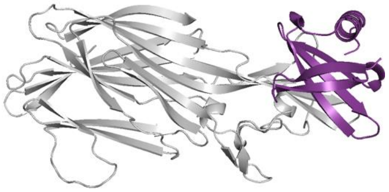

pLDDT = 90.3
pTM = 0.886
ipTM = 0.764

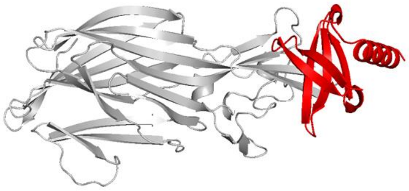

pLDDT = 88.4
pTM = 0.819
ipTM = 0.412

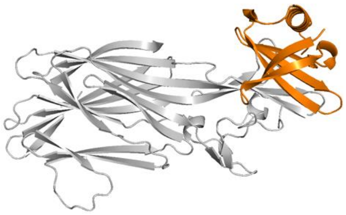

pLDDT = 89.4
pTM = 0.871
ipTM = 0.701

| | |
|---|---|
| Target | Calprotectin monomer s100a9 |
| Scaffold | Affitin |
| Total HTPS Reads | 7506 |
| Unique HTPS Reads | 841 |
| Top Variant % of Total Reads | 31 |
| Final Phage Titer (PFU/mL) | 8.13E+08 |

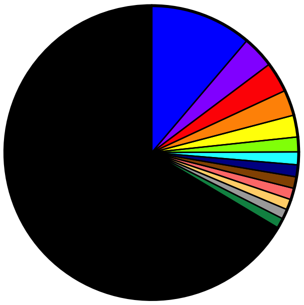

| | |
|---|---|
| Library: | GSVKVKF???GEEKEVDTSKI??V?R?GK?V?F?YDDNGK?G?G?V?EKDAPKELLDMLARAEREKKL |
| 11.20% | GSVKVKFATIGEEKEVDTSKIYRVARFGKTVFFLYDDNGKAGCGYVTEKDAPKELLDMLARAEREKKL |
| 3.54% | GSVKVKF***GEEKEVDTSKIRSVWRGGKWVDFSYDDNGKQGYGYVLEKDAPKELLDMLARAEREKKL |
| 3.32% | GSVKVKFDLSGEEKEVDTSKITHVLRIGKTVFFNYDDNGKRGTGFVVEKDAPKELLDMLARAEREKKL |
| 2.84% | GSVKVKFFYFGEEKEVDTSKIRSVGRSGKLVPFIYDDNGKPGAGIVPEKDAPKELLDMLARAEREKKL |
| 2.40% | GSVKVKFHYFGEEKEVDTSKIVHVLRIGKTVFFAYDDNGKNGFGFVIEKDAPKELLDMLARAEREKKL |
| 1.63% | GSVKVKFNFYGEEKEVDTSKITHVSRFGKTVFFTYDDNGKFGVGYVVEKDAPKELLDMLARAEREKKL |
| 1.41% | GSVKVKFLRFGEEKEVDTSKILSVFRRGKIVAFAYDDNGKCGYGFVREKDAPKELLDMLARAEREKKL |
| 1.32% | GSVKVKFISAGEEKEVDTSKIHIVRRVGKLVFFCYDDNGKHGYGFVYEKDAPKELLDMLARAEREKKL |
| 1.28% | GSVKVKFNGPGEEKEVDTSKILYVNRTGKVVHFLYDDNGKNGYGSVHEKDAPKELLDMLARAEREKKL |
| 1.27% | GSVKVKFHVAGEEKEVDTSKIRAVLRIGKTVFFAYDDNGKLGSGYVIEKDAPKELLDMLARAEREKKL |
| 1.15% | GSVKVKFFLGGEEKEVDTSKIICVSRFGKVVFFNYDDNGKSGTGYVFEKDAPKELLDMLARAEREKKL |
| 1.11% | GSVKVKFRIDGEEKEVDTSKICYVYRIGKAVIFIYDDNGKRGLGLVFEKDAPKELLDMLARAEREKKL |
| 1.11% | GSVKVKFCPCGEEKEVDTSKIRLVLRYGKIVTFCYDDNGKYGRGFVNEKDAPKELLDMLARAEREKKL |
| 66.44% | Other Sequence Less than 1% Total Reads |

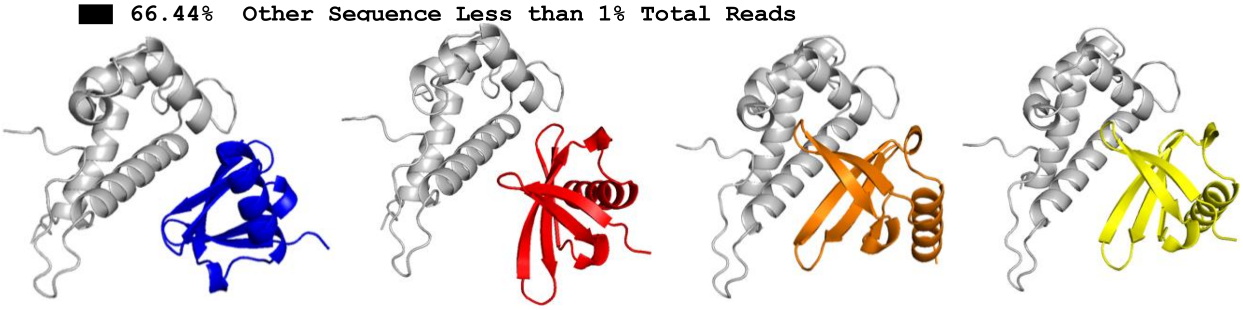

| | | | |
|---|---|---|---|
| pLDDT = 75.0 | pLDDT = 76.0 | pLDDT = 75.1 | pLDDT = 76.7 |
| pTM = 0.560 | pTM = 0.637 | pTM = 0.550 | pTM = 0.585 |
| ipTM = 0.195 | ipTM = 0.319 | ipTM = 0.183 | ipTM = 0.262 |

| Target | KIX |
|---|---|
| Scaffold | Affitin |
| Total HTPS Reads | 9333 |
| Unique HTPS Reads | 2655 |
| Top Variant % of Total Reads | 10 |
| Final Phage Titer (PFU/mL) | 6.92E+08 |

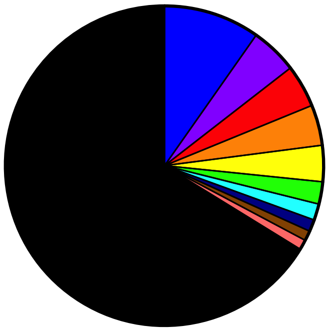

```
Library: GSVKVKF???GEEKEVDTSKI??V?R?GK?V?F?YDDNGK?G?G?V?EKDAPKELLDMLARAEREKKL
 9.77%   GSVKVKFVDRGEEKEVDTSKIVDVLRVGKFVSFLYDDNGKPGAGDVFEKDAPKELLDMLARAEREKKL
 4.76%   GSVKVKFYDAGEEKEVDTSKIDFVVRDGKGVVFFYDDNGKCGPGFVDEKDAPKELLDMLARAEREKKL
 4.32%   GSVKVKFSLSGEEKEVDTSKIRFVLRDGKYVHFDYDDNGKLGIGFVNEKDAPKELLDMLARAEREKKL
 4.10%   GSVKVKFVSSGEEKEVDTSKIDFVFRIGKAVGFIYDDNGKCGIGDVNEKDAPKELLDMLARAEREKKL
 3.66%   GSVKVKFDCCGEEKEVDTSKIVDVLRNGKNVCFLYDDNGKPGAGTVDEKDAPKELLDMLARAEREKKL
 2.26%   GSVKVKFSNSGEEKEVDTSKIDYVLRTGKFVHFDYDDNGKPGYGHVNEKDAPKELLDMLARAEREKKL
 1.64%   GSVKVKFFSGGEEKEVDTSKIYSVSRYGKHVSFCYDDNGKFGVGFVTEKDAPKELLDMLARAEREKKL
 1.21%   GSVKVKF***GEEKEVDTSKIRSVWRGGKWVDFSYDDNGKQGYGYVLEKDAPKELLDMLARAEREKKL
 1.02%   GSVKVKFRIDGEEKEVDTSKICYVYRIGKAVIFIYDDNGKRGLGLVFEKDAPKELLDMLARAEREKKL
 1.01%   GSVKVKFCPCGEEKEVDTSKIRLVLRYGKIVTFCYDDNGKYGRGFVNEKDAPKELLDMLARAEREKKL
66.25%   Other Sequence Less than 1% Total Reads
```

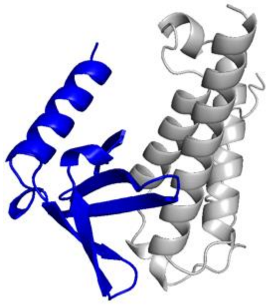

pLDDT = 74.9
pTM = 0.582
ipTM = 0.371

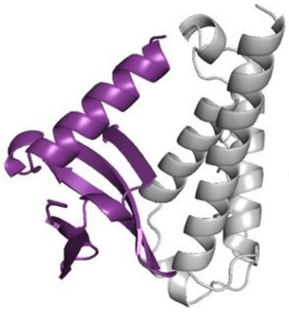

pLDDT = 70.1
pTM = 0.511
ipTM = 0.226

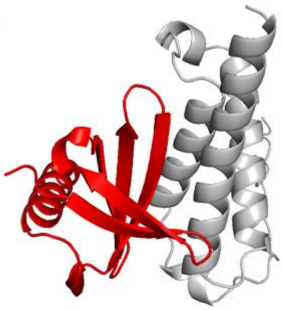

pLDDT = 80.1
pTM = 0.708
ipTM = 0.707

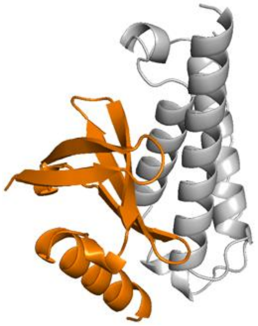

pLDDT = 69.2
pTM = 0.547
ipTM = 0.337

| Target | HRAS |
|---|---|
| Scaffold | Affitin |
| Total HTPS Reads | 7464 |
| Unique HTPS Reads | 947 |
| Top Variant % of Total Reads | 36 |
| Final Phage Titer (PFU/mL) | 7.41E+08 |

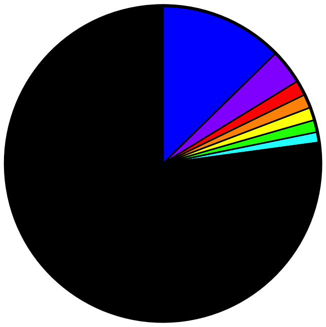

| | |
|---|---|
| Library: | GSVKVKF???GEEKEVDTSKI??V?R?GK?V?F?YDDNGK?G?G?V?EKDAPKELLDMLARAEREKKL |
| 12.69% | GSVKVKFPYPGEEKEVDTSKIVCVCRRGKIVNFRYDDNGKHGTGTVFEKDAPKELLDMLARAEREKKL |
| 3.58% | GSVKVKFSSIGEEKEVDTSKIICVCRYGKRVNFRYDDNGKFGAGNVFEKDAPKELLDMLARAEREKKL |
| 1.54% | GSVKVKFATPGEEKEVDTSKIICVCRTGKNVFFRYDDNGKSGAGSVNEKDAPKELLDMLARAEREKKL |
| 1.41% | GSVKVKFTHNGEEKEVDTSKIACVCRHGKSVNFRYDDNGKLGIGRVCEKDAPKELLDMLARAEREKKL |
| 1.34% | GSVKVKFHNNGEEKEVDTSKIACVCRTGKYVNFRYDDNGKIGSGCVLEKDAPKELLDMLARAEREKKL |
| 1.25% | GSVKVKFHSSGEEKEVDTSKIHCVCRTGKSVNFRYDDNGKVGSGVVLEKDAPKELLDMLARAEREKKL |
| 1.06% | GSVKVKFRIDGEEKEVDTSKICYVYRIGKAVIFIYDDNGKRGLGLVFEKDAPKELLDMLARAEREKKL |
| 77.14% | Other Sequence Less than 1% Total Reads |

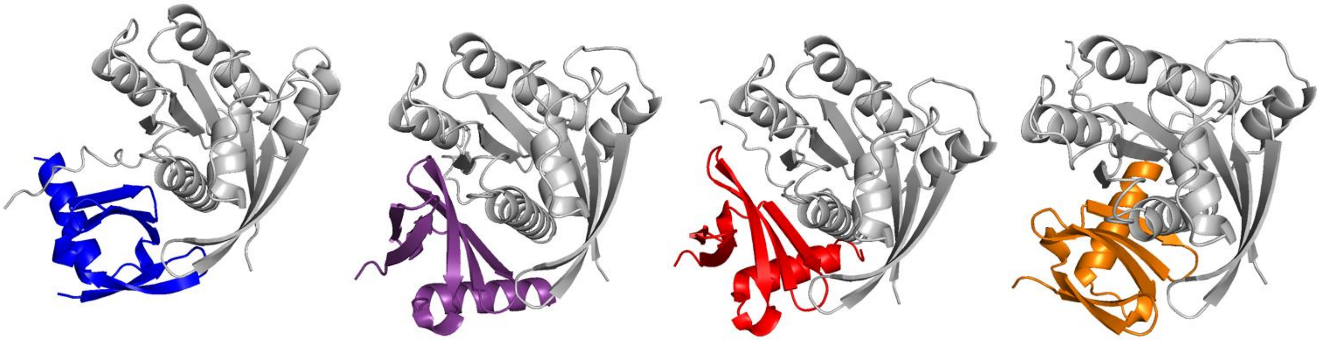

| | | | |
|---|---|---|---|
| pLDDT = 79.2 | pLDDT = 79.3 | pLDDT = 76.6 | pLDDT = 79.7 |
| pTM = 0.688 | pTM = 0.684 | pTM = 0.667 | pTM = 0.669 |
| ipTM = 0.258 | ipTM = 0.200 | ipTM = 0.169 | ipTM = 0.165 |

| Target | KRAS WT |
|---|---|
| Scaffold | Affitin |
| Total HTPS Reads | 7580 |
| Unique HTPS Reads | 818 |
| Top Variant % of Total Reads | 73 |
| Final Phage Titer (PFU/mL) | 9.55E+07 |

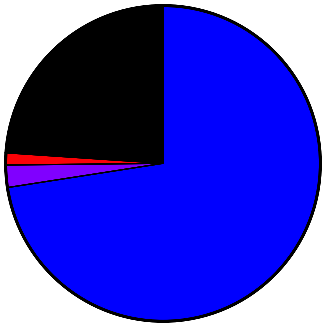

```
  Library: GSVKVKF???GEEKEVDTSKI??V?R?GK?V?F?YDDNGK?G?G?V?EKDAPKELLDMLARAEREKKL
   72.56%  GSVKVKFNGPGEEKEVDTSKILYVNRTGKVVHFLYDDNGKNGYGSVHEKDAPKELLDMLARAEREKKL
    2.28%  GSVKVKF***GEEKEVDTSKIRSVWRGGKWVDFSYDDNGKQGYGYVLEKDAPKELLDMLARAEREKKL
    1.23%  GSVKVKFRIDGEEKEVDTSKICYVYRIGKAVIFIYDDNGKRGLGLVFEKDAPKELLDMLARAEREKKL
   23.93%  Other Sequence Less than 1% Total Reads
```

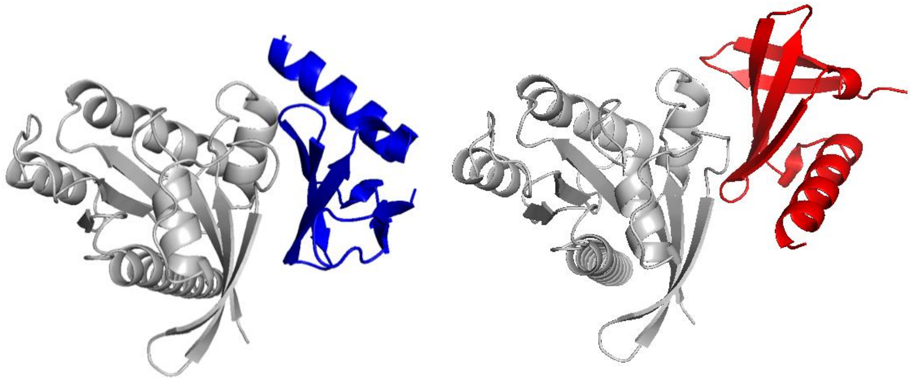

pLDDT = 77.4
pTM = 0.668
ipTM = 0.160

pLDDT = 82.5
pTM = 0.696
ipTM = 0.264

| | |
|---|---|
| Target | Calprotectin monomer s100a8 |
| Scaffold | Affitin |
| Total HTPS Reads | 9697 |
| Unique HTPS Reads | 1118 |
| Top Variant % of Total Reads | 48 |
| Final Phage Titer (PFU/mL) | 9.33E+08 |

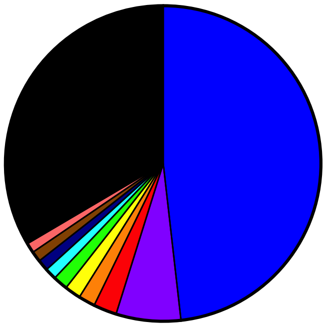

| Library: | GSVKVKF???GEEKEVDTSKI??V?R?GK?V?F?YDDNGK?G?G?V?EKDAPKELLDMLARAEREKKL |
|---|---|
| 48.21% | GSVKVKFISIGEEKEVDTSKIRVVFRAGKCVYFFYDDNGKLGFGGVPEKDAPKELLDMLARAEREKKL |
| 6.62% | GSVKVKFNTHGEEKEVDTSKIIYVNRRGKHVYFIYDDNGKLGGGVVDEKDAPKELLDMLARAEREKKL |
| 2.38% | GSVKVKFNTHGEEKEVDTSKIRVVFRAGKCVYFFYDDNGKLGFGGVPEKDAPKELLDMLARAEREKKL |
| 1.71% | GSVKVKFRSHGEEKEVDTSKILDVHRIGKSVFFIYDDNGKCGFGDVTEKDAPKELLDMLARAEREKKL |
| 1.64% | GSVKVKFISIGEEKEVDTSKIRVVFRAGKCVYFFYDDNGKLGGGVVDEKDAPKELLDMLARAEREKKL |
| 1.51% | GSVKVKFISIGEEKEVDTSKIIYVNRRGKHVYFIYDDNGKLGGGVVDEKDAPKELLDMLARAEREKKL |
| 1.19% | GSVKVKFLVHGEEKEVDTSKIFFVSRRGKAVYFIYDDNGKIGAGSVFEKDAPKELLDMLARAEREKKL |
| 1.14% | GSVKVKFRIDGEEKEVDTSKICYVYRIGKAVIFIYDDNGKRGLGLVFEKDAPKELLDMLARAEREKKL |
| 1.09% | GSVKVKFLVHGEEKEVDTSKIRVVFRAGKCVYFFYDDNGKLGFGGVPEKDAPKELLDMLARAEREKKL |
| 1.02% | GSVKVKF***GEEKEVDTSKIRSVWRGGKWVDFSYDDNGKQGYGYVLEKDAPKELLDMLARAEREKKL |
| 33.48% | Other Sequence Less than 1% Total Reads |

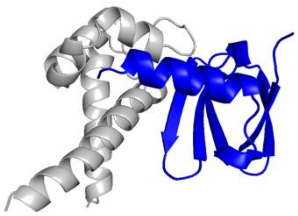

pLDDT = 87.8
pTM = 0.759
ipTM = 0.613

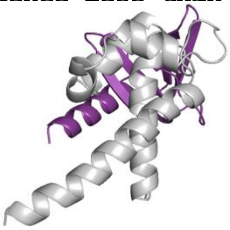

pLDDT = 78.5
pTM = 0.652
ipTM = 0.367

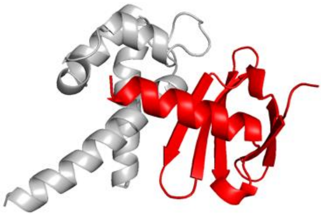

pLDDT = 86.2
pTM = 0.710
ipTM = 0.511

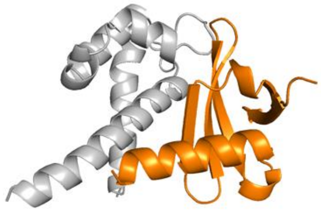

pLDDT = 80.4
pTM = 0.561
ipTM = 0.200

| | |
|---|---|
| Target | KRAS G12V |
| Scaffold | Affitin |
| Total HTPS Reads | 8588 |
| Unique HTPS Reads | 682 |
| Top Variant % of Total Reads | 78 |
| Final Phage Titer (PFU/mL) | 3.31E+08 |

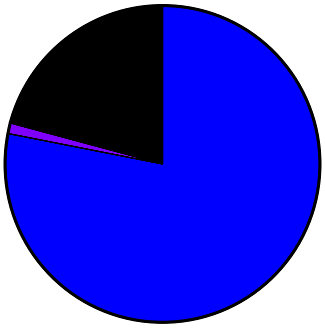

Library: GSVKVKF???GEEKEVDTSKI??V?R?GK?V?F?YDDNGK?G?G?V?EKDAPKELLDMLARAEREKKL

78.09% GSVKVKFNGPGEEKEVDTSKILYVNRTGKVVHFLYDDNGKNGYGSVHEKDAPKELLDMLARAEREKKL

1.09% GSVKVKFRIDGEEKEVDTSKICYVYRIGKAVIFIYDDNGKRGLGLVFEKDAPKELLDMLARAEREKKL

20.82% Other Sequence Less than 1% Total Reads

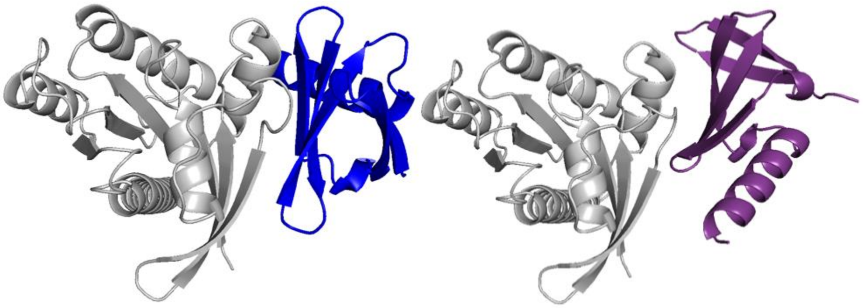

pLDDT = 77.6
pTM = 0.670
ipTM = 0.167

pLDDT = 82.4
pTM = 0.696
ipTM = 0.265

| Target | mCherry |
|---|---|
| Scaffold | Affitin |
| Total HTPS Reads | 11909 |
| Unique HTPS Reads | 1436 |
| Top Variant % of Total Reads | 33 |
| Final Phage Titer (PFU/mL) | 8.32E+08 |

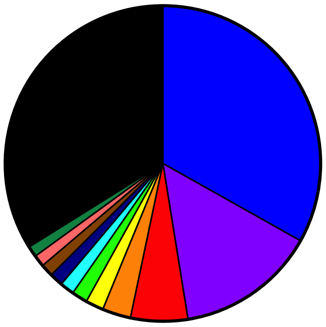

```
Library: GSVKVKF???GEEKEVDTSKI??V?R?GK?V?F?YDDNGK?G?G?V?EKDAPKELLDMLARAEREKKL
33.14%   GSVKVKFDFLGEEKEVDTSKIHLVYRHGKYVRFVYDDNGKGGYGNVFEKDAPKELLDMLARAEREKKL
14.32%   GSVKVKFGAGGEEKEVDTSKITDVTRRGKYVYFNYDDNGKSGYGFVYEKDAPKELLDMLARAEREKKL
5.84%    GSVKVKFVGFGEEKEVDTSKIRSVSRSGKTVYFHYDDNGKGGYGFVCEKDAPKELLDMLARAEREKKL
2.92%    GSVKVKFGAGGEEKEVDTSKIHLVYRHGKYVRFVYDDNGKGGYGNVFEKDAPKELLDMLARAEREKKL
1.85%    GSVKVKFVGFGEEKEVDTSKIHLVYRHGKYVRFVYDDNGKGGYGNVFEKDAPKELLDMLARAEREKKL
1.61%    GSVKVKFDFLGEEKEVDTSKITDVTRRGKYVYFNYDDNGKSGYGFVYEKDAPKELLDMLARAEREKKL
1.46%    GSVKVKF***GEEKEVDTSKIRSVWRGGKWVDFSYDDNGKQGYGYVLEKDAPKELLDMLARAEREKKL
1.37%    GSVKVKFDFLGEEKEVDTSKIHLVYRHGKYVRFVYDDNGKGGYGFVCEKDAPKELLDMLARAEREKKL
1.29%    GSVKVKFVGFGEEKEVDTSKIRSVSRSGKTVYFHYDDNGKGGYGNVFEKDAPKELLDMLARAEREKKL
1.19%    GSVKVKFDFLGEEKEVDTSKIRSVSRSGKTVYFHYDDNGKGGYGFVCEKDAPKELLDMLARAEREKKL
1.13%    GSVKVKFNGPGEEKEVDTSKILYVNRTGKVVHFLYDDNGKNGYGSVHEKDAPKELLDMLARAEREKKL
33.87%   Other Sequence Less than 1% Total Reads
```

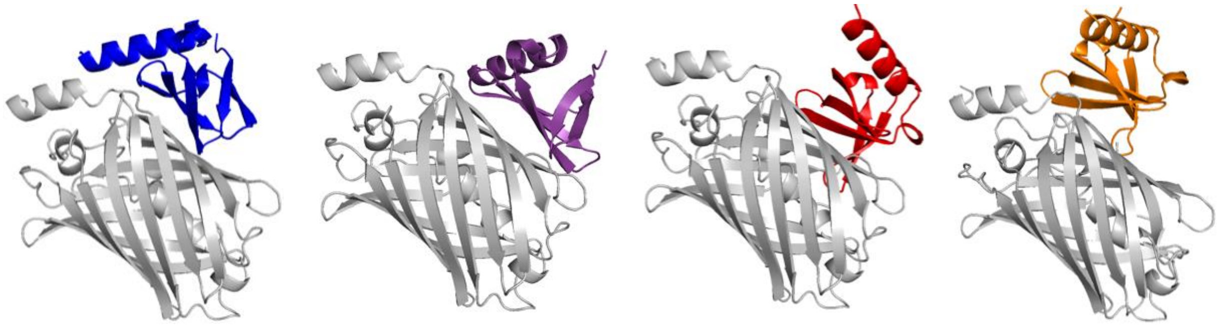

| | | | |
|---|---|---|---|
| pLDDT = 83.4 | pLDDT = 83.2 | pLDDT = 82.1 | pLDDT = 82.6 |
| pTM = 0.721 | pTM = 0.727 | pTM = 0.713 | pTM = 0.718 |
| ipTM = 0.152 | ipTM = 0.170 | ipTM = 0.150 | ipTM = 0.129 |

| Target | Myc DBD |
|---|---|
| Scaffold | Affitin |
| Total HTPS Reads | 12528 |
| Unique HTPS Reads | 1116 |
| Top Variant % of Total Reads | 49 |
| Final Phage Titer (PFU/mL) | 2.88E+08 |

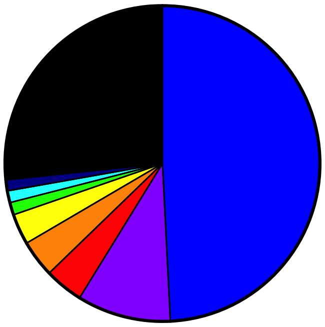

| | |
|---|---|
| Library: | GSVKVKF???GEEKEVDTSKI??V?R?GK?V?F?YDDNGK?G?G?V?EKDAPKELLDMLARAEREKKL |
| 49.19% | GSVKVKFLDAGEEKEVDTSKIRLVDRFGKIVIFTYDDNGKVGVGDVDEKDAPKELLDMLARAEREKKL |
| 9.63% | GSVKVKFNLHGEEKEVDTSKICLVDRIGKFVGFIYDDNGKDGFGAVFEKDAPKELLDMLARAEREKKL |
| 3.95% | GSVKVKFNLHGEEKEVDTSKIRLVDRFGKIVIFTYDDNGKVGVGDVDEKDAPKELLDMLARAEREKKL |
| 3.81% | GSVKVKFGTTGEEKEVDTSKITHVRRTGKGVVFRYDDNGKIGLGIVTEKDAPKELLDMLARAEREKKL |
| 3.14% | GSVKVKFLDAGEEKEVDTSKICLVDRIGKFVGFIYDDNGKDGFGAVFEKDAPKELLDMLARAEREKKL |
| 1.29% | GSVKVKFLDAGEEKEVDTSKIRLVDRFGKIVIFTYDDNGKDGFGAVFEKDAPKELLDMLARAEREKKL |
| 1.21% | GSVKVKFRIDGEEKEVDTSKICYVYRIGKAVIFIYDDNGKRGLGLVFEKDAPKELLDMLARAEREKKL |
| 1.13% | GSVKVKFGTTGEEKEVDTSKIRLVDRFGKIVIFTYDDNGKVGVGDVDEKDAPKELLDMLARAEREKKL |
| 26.65% | Other Sequence Less than 1% Total Reads |

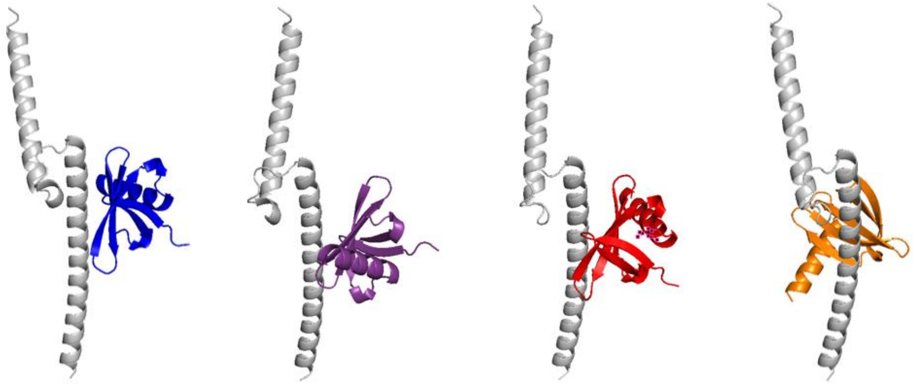

| | | | |
|---|---|---|---|
| pLDDT = 91.6 | pLDDT = 78.4 | pLDDT = 94.4 | pLDDT = 81.4 |
| pTM = 0.753 | pTM = 0.566 | pTM = 0.816 | pTM = 0.477 |
| ipTM = 0.744 | ipTM = 0.365 | ipTM = 0.850 | ipTM = 0.150 |

| Target | PP2A-B" (PR72) |
|---|---|
| Scaffold | Affitin |
| Total HTPS Reads | 15478 |
| Unique HTPS Reads | 1166 |
| Top Variant % of Total Reads | 77 |
| Final Phage Titer (PFU/mL) | 1.20E+09 |

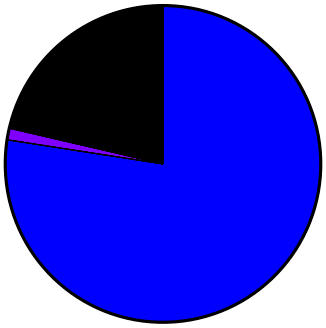

**Library:** **GSVKVKF???GEEKEVDTSKI??V?R?GK?V?F?YDDNGK?G?G?V?EKDAPKELLDMLARAEREKKL**

77.45% **GSVKVKFCPCGEEKEVDTSKIRLVLRYGKIVTFCYDDNGKYGRGFVNEKDAPKELLDMLARAEREKKL**

1.17% **GSVKVKFRIDGEEKEVDTSKICYVYRIGKAVIFIYDDNGKRGLGLVFEKDAPKELLDMLARAEREKKL**

21.38% **Other Sequence Less than 1% Total Reads**

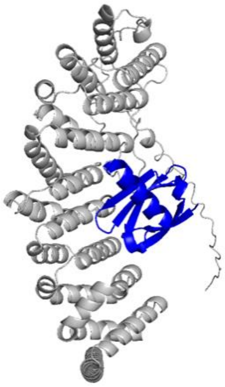

pLDDT = 78.2
pTM = 0.719
ipTM = 0.318

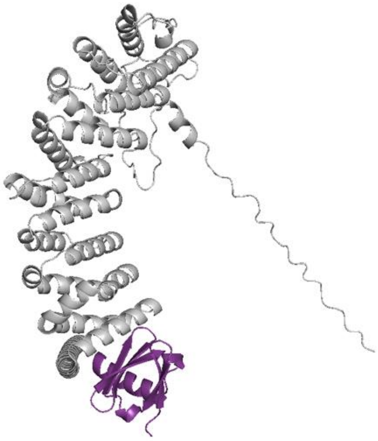

pLDDT = 82.2
pTM = 0.768
ipTM = 0.817

| Target | MBP |
|---|---|
| Scaffold | Affitin |
| Total HTPS Reads | 7972 |
| Unique HTPS Reads | 1561 |
| Top Variant % of Total Reads | 36 |
| Final Phage Titer (PFU/mL) | 3.93E+08 |

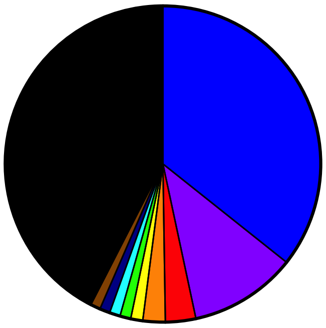

Library: GSVKVKF???GEEKEVDTSKI??V?R?GK?V?F?YDDNGK?G?G?V?EKDAPKELLDMLARAEREKKL

35.67% GSVKVKFSRLGEEKEVDTSKIHTVYRIGKSVGFNYDDNGKYGFGSVTEKDAPKELLDMLARAEREKKL

10.98% GSVKVKF***GEEKEVDTSKIRSVWRGGKWVDFSYDDNGKQGYGYVLEKDAPKELLDMLARAEREKKL

3.11% GSVKVKFHRLGEEKEVDTSKIHTVFRSGKSVVFDYDDNGKRGVGDVSEKDAPKELLDMLARAEREKKL

2.28% GSVKVKFLNFGEEKEVDTSKINIVYRIGKSVSFTYDDNGKRGFGDVYEKDAPKELLDMLARAEREKKL

1.23% GSVKVKFSRLGEEKEVDTSKINIVYRIGKSVSFTYDDNGKRGFGDVYEKDAPKELLDMLARAEREKKL

1.12% GSVKVKFPFSGEEKEVDTSKIHHVYRIGKGVSFNYDDNGKRGIGHVIEKDAPKELLDMLARAEREKKL

1.09% GSVKVKFCPCGEEKEVDTSKIRLVLRYGKIVTFCYDDNGKYGRGFVNEKDAPKELLDMLARAEREKKL

1.05% GSVKVKFRIDGEEKEVDTSKICYVYRIGKAVIFIYDDNGKRGLGLVFEKDAPKELLDMLARAEREKKL

1.02% GSVKVKFCRLGEEKEVDTSKIHVVYRIGKSVCFNYDDNGKRGFGNVTEKDAPKELLDMLARAEREKKL

42.45% Other Sequence Less than 1% Total Reads

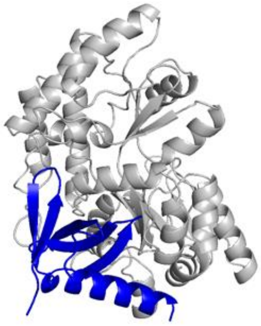

pLDDT = 90.1
pTM = 0.833
ipTM = 0.395

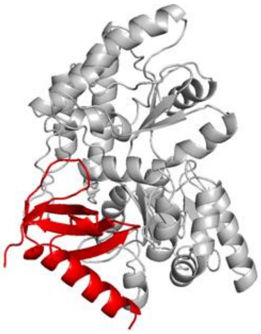

pLDDT = 88.3
pTM = 0.842
ipTM = 0.356

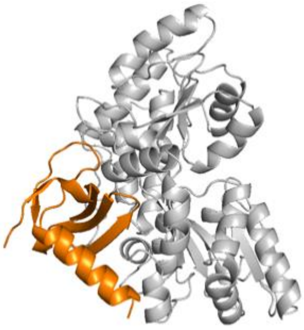

pLDDT = 88.3
pTM = 0.842
ipTM = 0.356

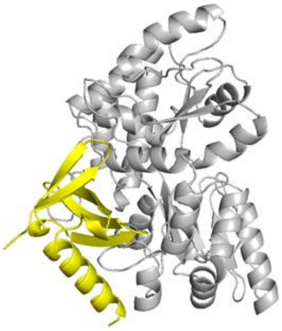

pLDDT = 89.5
pTM = 0.830
ipTM = 0.358

| Target | Fibronectin |
|---|---|
| Scaffold | Affitin |
| Total HTPS Reads | 12644 |
| Unique HTPS Reads | 1530 |
| Top Variant % of Total Reads | 26 |
| Final Phage Titer (PFU/mL) | 7.76E+08 |

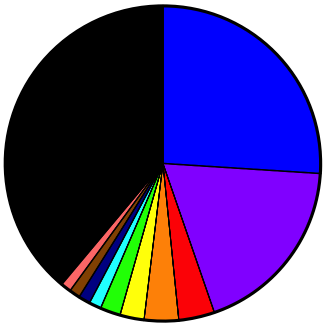

```
Library: GSVKVKF???GEEKEVDTSKI??V?R?GK?V?F?YDDNGK?G?G?V?EKDAPKELLDMLARAEREKKL
26.01%   GSVKVKFTNDGEEKEVDTSKIRNVTRHGKHVVFIYDDNGKSGLGTVIEKDAPKELLDMLARAEREKKL
18.70%   GSVKVKFSYTGEEKEVDTSKIFSVTRHGKYVVFCYDDNGKSGLGVVHEKDAPKELLDMLARAEREKKL
3.68%    GSVKVKFTNDGEEKEVDTSKIFSVTRHGKYVVFCYDDNGKSGLGVVHEKDAPKELLDMLARAEREKKL
3.52%    GSVKVKFSYTGEEKEVDTSKIRNVTRHGKHVVFIYDDNGKSGLGTVIEKDAPKELLDMLARAEREKKL
2.48%    GSVKVKFINIGEEKEVDTSKIFGVTRHGKYVVFAYDDNGKHGIGNVNEKDAPKELLDMLARAEREKKL
2.07%    GSVKVKFTNDGEEKEVDTSKIRNVTRHGKHVVFIYDDNGKSGLGVVHEKDAPKELLDMLARAEREKKL
1.26%    GSVKVKFRIDGEEKEVDTSKICYVYRIGKAVIFIYDDNGKRGLGLVFEKDAPKELLDMLARAEREKKL
1.19%    GSVKVKFSYTGEEKEVDTSKIFSVTRHGKYVVFCYDDNGKSGLGTVIEKDAPKELLDMLARAEREKKL
1.12%    GSVKVKFTNDGEEKEVDTSKIRNVTRHGKHVVFCYDDNGKSGLGVVHEKDAPKELLDMLARAEREKKL
1.04%    GSVKVKFTNDGEEKEVDTSKIRNVTRHGKYVVFCYDDNGKSGLGVVHEKDAPKELLDMLARAEREKKL
38.93%   Other Sequence Less than 1% Total Reads
```

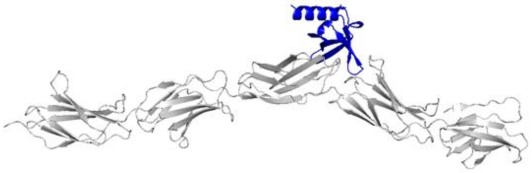

pLDDT = 83.8
pTM = 0.531
ipTM = 0.173

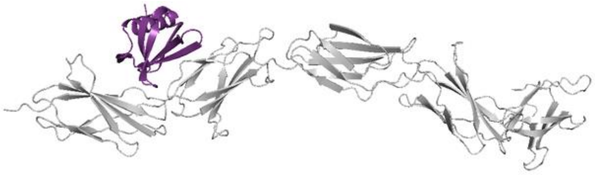

pLDDT = 83.5
pTM = 0.557
ipTM = 0.180

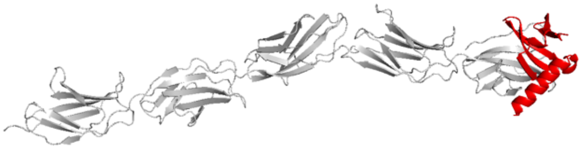

pLDDT = 89.4
pTM = 0.610
ipTM = 0.725

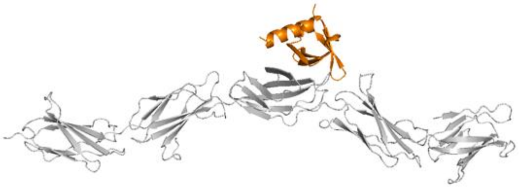

pLDDT = 86.7
pTM = 0.584
ipTM = 0.506

| Target | Calprotectin heterodimer 6aa linker |
|---|---|
| Scaffold | Affitin |
| Total HTPS Reads | 11907 |
| Unique HTPS Reads | 1924 |
| Top Variant % of Total Reads | 19 |
| Final Phage Titer (PFU/mL) | 1.45E+09 |

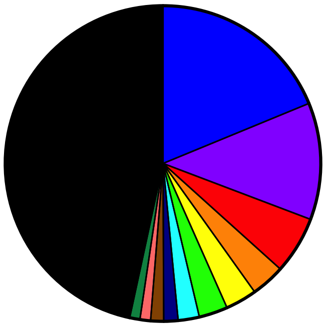

| | |
|---|---|
| Library: | GSVKVKF???GEEKEVDTSKI??V?R?GK?V?F?YDDNGK?G?G?V?EKDAPKELLDMLARAEREKKL |
| 18.79% | GSVKVKFVLSGEEKEVDTSKIIRVNRFGKVVYFSYDDNGKSGFGYVSEKDAPKELLDMLARAEREKKL |
| 11.98% | GSVKVKFNDDGEEKEVDTSKIFRVNRFGKFVYFVYDDNGKLGFGVVSEKDAPKELLDMLARAEREKKL |
| 5.89% | GSVKVKFPFHGEEKEVDTSKIPRVNRFGKCVYFPYDDNGKPGLGYVAEKDAPKELLDMLARAEREKKL |
| 3.49% | GSVKVKF***GEEKEVDTSKIRSVWRGGKWVDFSYDDNGKQGYGYVLEKDAPKELLDMLARAEREKKL |
| 3.21% | GSVKVKFHITGEEKEVDTSKILRVNRFGKLVYFNYDDNGKIGFGDVSEKDAPKELLDMLARAEREKKL |
| 2.96% | GSVKVKFNDDGEEKEVDTSKIIRVNRFGKVVYFSYDDNGKSGFGYVSEKDAPKELLDMLARAEREKKL |
| 2.17% | GSVKVKFVLSGEEKEVDTSKIFRVNRFGKFVYFVYDDNGKLGFGVVSEKDAPKELLDMLARAEREKKL |
| 1.47% | GSVKVKFVLSGEEKEVDTSKIPRVNRFGKCVYFPYDDNGKPGLGYVAEKDAPKELLDMLARAEREKKL |
| 1.33% | GSVKVKFRIDGEEKEVDTSKICYVYRIGKAVIFIYDDNGKRGLGLVFEKDAPKELLDMLARAEREKKL |
| 1.08% | GSVKVKFNGPGEEKEVDTSKILYVNRTGKVVHFLYDDNGKNGYGSVHEKDAPKELLDMLARAEREKKL |
| 1.05% | GSVKVKFNDDGEEKEVDTSKIPRVNRFGKCVYFPYDDNGKPGLGYVAEKDAPKELLDMLARAEREKKL |
| 46.60% | Other Sequence Less than 1% Total Reads |

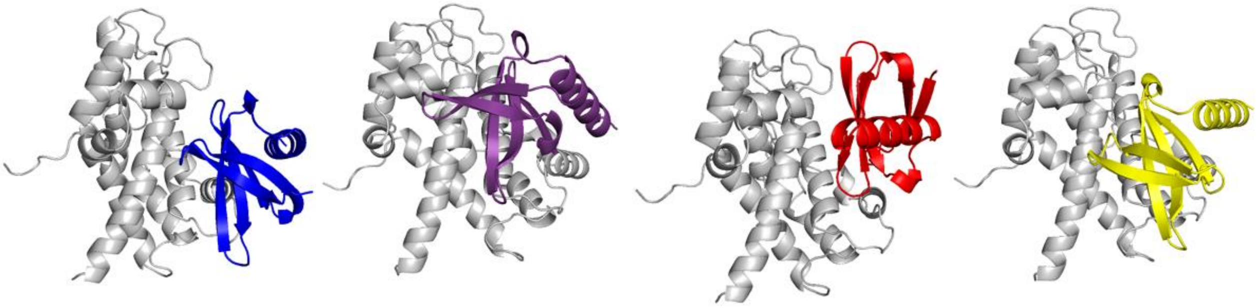

| | | | |
|---|---|---|---|
| pLDDT = 84.1 | pLDDT = 83.6 | pLDDT = 84.2 | pLDDT = 86.0 |
| pTM = 0.844 | pTM = 0.827 | pTM = 0.841 | pTM = 0.866 |
| ipTM = 0.764 | ipTM = 0.702 | ipTM = 0.720 | ipTM = 0.850 |

| Target | GABARAP |
| --- | --- |
| Scaffold | Affitin |
| Total HTPS Reads | 10315 |
| Unique HTPS Reads | 1614 |
| Top Variant % of Total Reads | 20 |
| Final Phage Titer (PFU/mL) | 5.50E+08 |

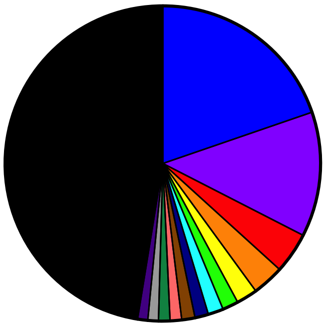

```
Library: GSVKVKF???GEEKEVDTSKI??V?R?GK?V?F?YDDNGK?G?G?V?EKDAPKELLDMLARAEREKKL
19.73%   GSVKVKFDGNGEEKEVDTSKIRSVYRDGKVVFFIYDDNGKRGAGYVFEKDAPKELLDMLARAEREKKL
12.81%   GSVKVKFDDFGEEKEVDTSKIYLVYRFGKYVYFIYDDNGKRGIGLVPEKDAPKELLDMLARAEREKKL
4.19%    GSVKVKFNCPGEEKEVDTSKINYVTRCGKYVYFDYDDNGKGGNGFVSEKDAPKELLDMLARAEREKKL
3.26%    GSVKVKFPDFGEEKEVDTSKIGVVYRLGKYVGFLYDDNGKSGDGLVNEKDAPKELLDMLARAEREKKL
2.14%    GSVKVKFHCPGEEKEVDTSKITYVLRLGKYVYFSYDDNGKDGDGFVIEKDAPKELLDMLARAEREKKL
1.65%    GSVKVKFDDFGEEKEVDTSKIRSVYRDGKVVFFIYDDNGKRGAGYVFEKDAPKELLDMLARAEREKKL
1.56%    GSVKVKFNITGEEKEVDTSKIIYVFRNGKFVIFDYDDNGKHGAGCVAEKDAPKELLDMLARAEREKKL
1.40%    GSVKVKFRIDGEEKEVDTSKICYVYRIGKAVIFIYDDNGKRGLGLVFEKDAPKELLDMLARAEREKKL
1.35%    GSVKVKFPTYGEEKEVDTSKINSVFRIGKYVCFDYDDNGKRGDGIVDEKDAPKELLDMLARAEREKKL
1.19%    GSVKVKFDGNGEEKEVDTSKIRSVYRDGKVVFFIYDDNGKRGIGLVPEKDAPKELLDMLARAEREKKL
1.17%    GSVKVKFPIFGEEKEVDTSKILAVFRTGKYVVFAYDDNGKTGSGYVAEKDAPKELLDMLARAEREKKL
1.09%    GSVKVKFPTFGEEKEVDTSKIHTVLRIGKFVFFDYDDNGKYGDGFVNEKDAPKELLDMLARAEREKKL
1.03%    GSVKVKFDGNGEEKEVDTSKIYLVYRFGKYVYFIYDDNGKRGIGLVPEKDAPKELLDMLARAEREKKL
47.45%   Other Sequence Less than 1% Total Reads
```

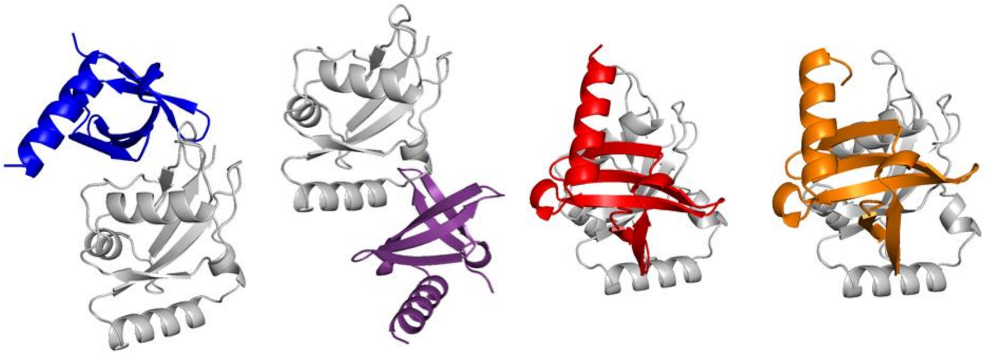

| | | | |
| --- | --- | --- | --- |
| pLDDT = 81.2 | pLDDT = 86.5 | pLDDT = 88.3 | pLDDT = 93.3 |
| pTM = 0.602 | pTM = 0.651 | pTM = 0.862 | pTM = 0.890 |
| ipTM = 0.129 | ipTM = 0.221 | ipTM = 0.824 | ipTM = 0.850 |

| Target | OmpF (ec) |
|---|---|
| Scaffold | Affitin |
| Total HTPS Reads | 4796 |
| Unique HTPS Reads | 833 |
| Top Variant % of Total Reads | 65 |
| Final Phage Titer (PFU/mL) | 1.07E+08 |

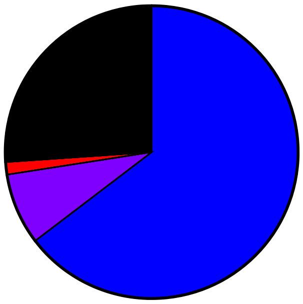

```
  Library: GSVKVKF???GEEKEVDTSKI??V?R?GK?V?F?YDDNGK?G?G?V?EKDAPKELLDMLARAEREKKL
  64.64%   GSVKVKFIISGEEKEVDTSKITNVYRIGKTVGFYYDDNGKLGGGSVPEKDAPKELLDMLARAEREKKL
  7.90%    GSVKVKF***GEEKEVDTSKIRSVWRGGKWVDFSYDDNGKQGYGYVLEKDAPKELLDMLARAEREKKL
  1.38%    GSVKVKFLIHGEEKEVDTSKIIFVNRYGKGVTFFYDDNGKYGLGHVTEKDAPKELLDMLARAEREKKL
  26.08%   Other Sequence less than 1%
```

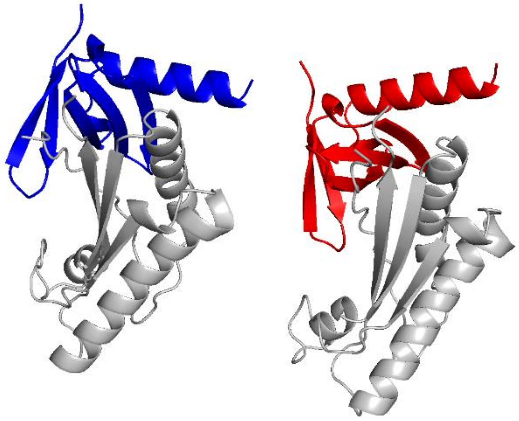

| | |
|---|---|
| pLDDT = 86.3 | pLDDT = 85.3 |
| pTM = 0.779 | pTM = 0.767 |
| ipTM = 0.653 | ipTM = 0.633 |

| Target | 14-3-3z |
|---|---|
| Scaffold | Affitin |
| Total HTPS Reads | 6084 |
| Unique HTPS Reads | 2721 |
| Top Variant % of Total Reads | 6 |
| Final Phage Titer (PFU/mL) | 1.23E+09 |

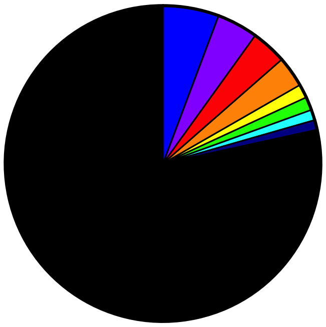

```
Library: GSVKVKF???GEEKEVDTSKI??V?R?GK?V?F?YDDNGK?G?G?V?EKDAPKELLDMLARAEREKKL
5.74%    GSVKVKFFDAGEEKEVDTSKIFPVPRYGKYVIFYYDDNGKVGRGSVNEKDAPKELLDMLARAEREKKL
4.21%    GSVKVKF***GEEKEVDTSKIRSVWRGGKWVDFSYDDNGKQGYGYVLEKDAPKELLDMLARAEREKKL
3.62%    GSVKVKFVVRGEEKEVDTSKISAVARAGKIVGFAYDDNGKLGVGHVHEKDAPKELLDMLARAEREKKL
3.16%    GSVKVKFHPNGEEKEVDTSKIYGVYRIGKIVNFIYDDNGKFGRGSVREKDAPKELLDMLARAEREKKL
1.40%    GSVKVKFRSCGEEKEVDTSKISIVNRSGKTVGFLYDDNGKFGIGIVLEKDAPKELLDMLARAEREKKL
1.36%    GSVKVKFANHGEEKEVDTSKIRHVLRRGKHVIFSYDDNGKSGLGLVHEKDAPKELLDMLARAEREKKL
1.08%    GSVKVKFYFRGEEKEVDTSKIYGVYRFGKCVSFIYDDNGKFGSGRVFEKDAPKELLDMLARAEREKKL
1.00%    GSVKVKFCPCGEEKEVDTSKIRLVLRYGKIVTFCYDDNGKYGRGFVNEKDAPKELLDMLARAEREKKL
78.44%   Other Sequence Less than 1% Total Reads
```

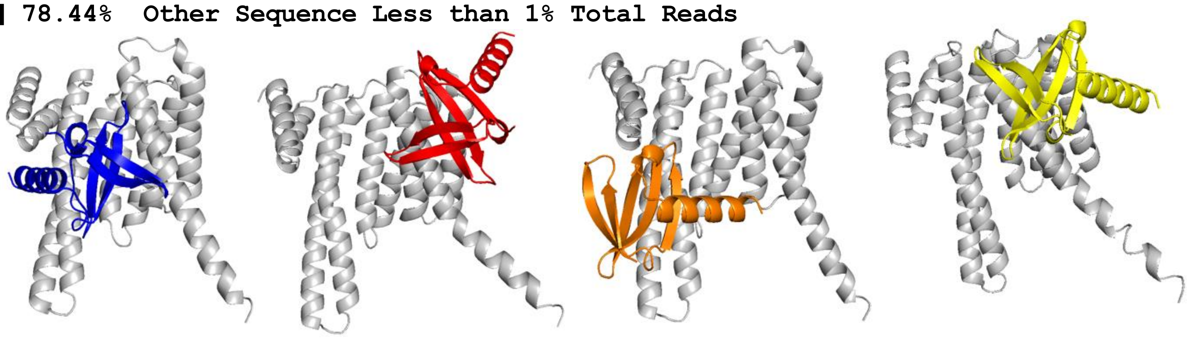

pLDDT = 79.4
pTM = 0.736
ipTM = 0.177

pLDDT = 91.0
pTM = 0.860
ipTM = 0.771

pLDDT = 93.3
pTM = 0.901
ipTM = 0.912

pLDDT = 91.3
pTM = 0.863
ipTM = 0.779

| | |
|---|---|
| Target | TEV |
| Scaffold | Affitin |
| Total HTPS Reads | 5592 |
| Unique HTPS Reads | 563 |
| Top Variant % of Total Reads | 79 |
| Final Phage Titer (PFU/mL) | 2.14E+07 |

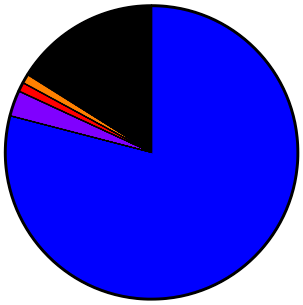

```
Library: GSVKVKF???GEEKEVDTSKI??V?R?GK?V?F?YDDNGK?G?G?V?EKDAPKELLDMLARAEREKKL
79.02%   GSVKVKFLIHGEEKEVDTSKIIFVNRYGKGVTFFYDDNGKYGLGHVTEKDAPKELLDMLARAEREKKL
2.84%    GSVKVKF***GEEKEVDTSKIRSVWRGGKWVDFSYDDNGKQGYGYVLEKDAPKELLDMLARAEREKKL
1.02%    GSVKVKFHIPGEEKEVDTSKIFIVLRYGKFVNFLYDDNGKAGYGYVHEKDAPKELLDMLARAEREKKL
1.02%    GSVKVKFSYSGEEKEVDTSKISIVHRNGKTVCFFYDDNGKCGCGIVNEKDAPKELLDMLARAEREKKL
16.09%   Other Sequence less than 1%
```

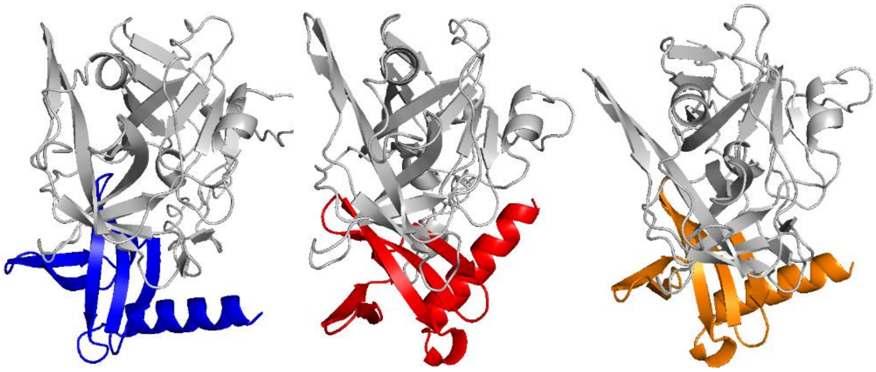

pLDDT = 79.1
pTM = 0.764
ipTM = 0.503

pLDDT = 78.7
pTM = 0.790
ipTM = 0.713

pLDDT = 76.1
pTM = 0.719
ipTM = 0.312

| Target | FnbpA |
|---|---|
| Scaffold | Affitin |
| Total HTPS Reads | 12202 |
| Unique HTPS Reads | 2358 |
| Top Variant % of Total Reads | 50 |
| Final Phage Titer (PFU/mL) | 6.76E+08 |

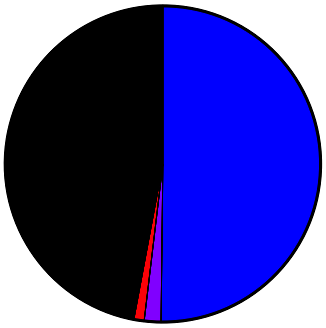

**Library: GSVKVKF???GEEKEVDTSKI??V?R?GK?V?F?YDDNGK?G?G?V?EKDAPKELLDMLARAEREKKL**

**50.20% GSVKVKFASIGEEKEVDTSKIFYVPRGGKPVHFYL***W*IRLRYCLEKDAPKELLDMLARAEREKKL**

**1.71% GSVKVKF***GEEKEVDTSKIRSVWRGGKWVDFSYDDNGKQGYGYVLEKDAPKELLDMLARAEREKKL**

**1.06% GSVKVKFRIDGEEKEVDTSKICYVYRIGKAVIFIYDDNGKRGLGLVFEKDAPKELLDMLARAEREKKL**

**47.03% Other Sequence Less than 1% Total Reads**

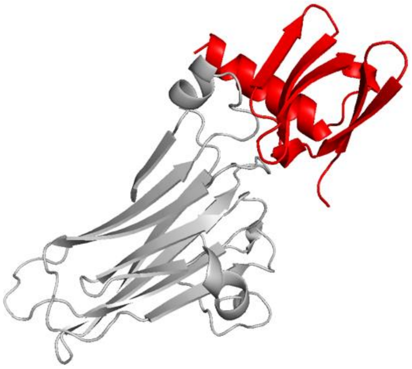

pLDDT = 80.4
pTM = 0.645
ipTM = 0.123

| | |
|---|---|
| Target | IBTK |
| Scaffold | Affitin |
| Total HTPS Reads | 16754 |
| Unique HTPS Reads | 1014 |
| Top Variant % of Total Reads | 82 |
| Final Phage Titer (PFU/mL) | 6.92E+08 |

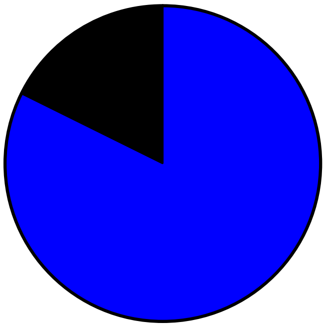

Library: GSVKVKF???GEEKEVDTSKI??V?R?GK?V?F?YDDNGK?G?G?V?EKDAPKELLDMLARAEREKKL

82.32% GSVKVKFRIDGEEKEVDTSKICYVYRIGKAVIFIYDDNGKRGLGLVFEKDAPKELLDMLARAEREKKL

17.68% Other Sequence Less than 1% Total Reads

**Insufficient Confidence to Predict Complex Structure (pLDDT, pTM)**

pLDDT = 49.8
pTM = 0.317
ipTM = 0.175

| Target | PAL |
|---|---|
| Scaffold | Affitin |
| Total HTPS Reads | 5671 |
| Unique HTPS Reads | 1889 |
| Top Variant % of Total Reads | 48 |
| Final Phage Titer (PFU/mL) | 4.79E+08 |

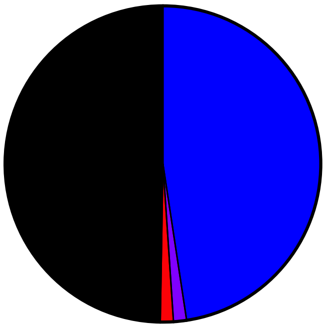

Library: GSVKVKF???GEEKEVDTSKI??V?R?GK?V?F?YDDNGK?G?G?V?EKDAPKELLDMLARAEREKKL

47.59% GSVKVKF***GEEKEVDTSKIRSVWRGGKWVDFSYDDNGKQGYGYVLEKDAPKELLDMLARAEREKKL

1.34% GSVKVKFNGPGEEKEVDTSKILYVNRTGKVVHFLYDDNGKNGYGSVHEKDAPKELLDMLARAEREKKL

1.34% GSVKVKFRIDGEEKEVDTSKICYVYRIGKAVIFIYDDNGKRGLGLVFEKDAPKELLDMLARAEREKKL

49.73% Other Sequence Less than 1% Total Reads

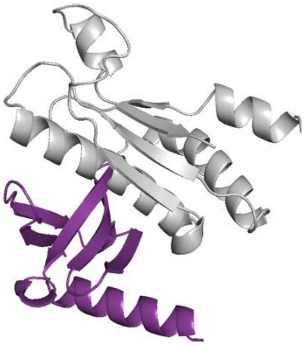

pLDDT = 91.4
pTM = 0.857
ipTM = 0.717

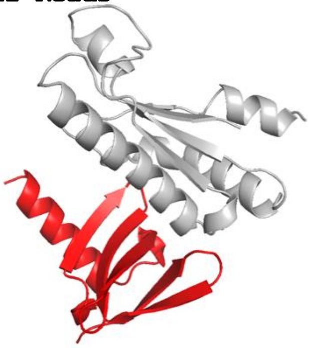

pLDDT = 88.1
pTM = 0.685
ipTM = 0.279

| Target | PIK3CB |
|---|---|
| Scaffold | Affitin |
| Total HTPS Reads | 9800 |
| Unique HTPS Reads | 1184 |
| Top Variant % of Total Reads | 36 |
| Final Phage Titer (PFU/mL) | 1.58E+09 |

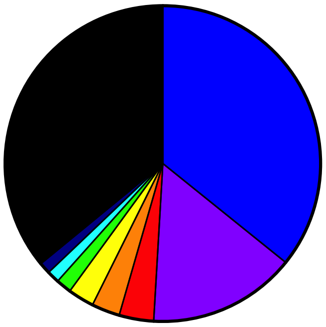

```
Library: GSVKVKF???GEEKEVDTSKI??V?R?GK?V?F?YDDNGK?G?G?V?EKDAPKELLDMLARAEREKKL
35.78%   GSVKVKFYNTGEEKEVDTSKISHVIRYGKFVCFFYDDNGKRGAGVVVEKDAPKELLDMLARAEREKKL
15.15%   GSVKVKFSRFGEEKEVDTSKILVVDRNGKLVYFGYDDNGKYGSGIVNEKDAPKELLDMLARAEREKKL
3.53%    GSVKVKFTDAGEEKEVDTSKIGVVIRHGKVVLFYYDDNGKLGRGYVLEKDAPKELLDMLARAEREKKL
2.90%    GSVKVKFYNTGEEKEVDTSKILVVDRNGKLVYFGYDDNGKYGSGIVNEKDAPKELLDMLARAEREKKL
2.67%    GSVKVKFSRFGEEKEVDTSKISHVIRYGKFVCFFYDDNGKRGAGVVVEKDAPKELLDMLARAEREKKL
1.58%    GSVKVKFLSVGEEKEVDTSKIIFVSRRGKCVFFLYDDNGKFGPGIVLEKDAPKELLDMLARAEREKKL
1.27%    GSVKVKFRIDGEEKEVDTSKICYVYRIGKAVIFIYDDNGKRGLGLVFEKDAPKELLDMLARAEREKKL
1.18%    GSVKVKF***GEEKEVDTSKIRSVWRGGKWVDFSYDDNGKQGYGYVLEKDAPKELLDMLARAEREKKL
35.94%   Other Sequence Less than 1% Total Reads
```

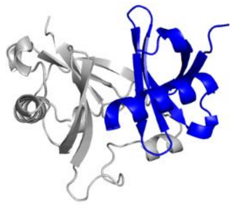

pLDDT = 78.2
pTM = 0.662
ipTM = 0.429

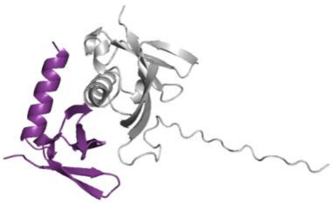

pLDDT = 85.1
pTM = 0.784
ipTM = 0.734

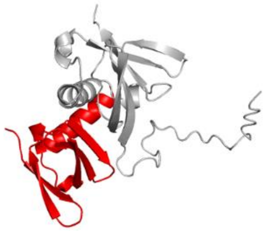

pLDDT = 85.1
pTM = 0.772
ipTM = 0.704

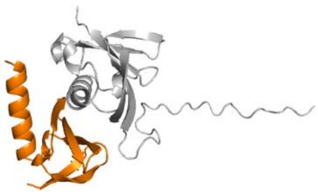

pLDDT = 85.8
pTM = 0.818
ipTM = 0.815

| Target | SortaseA |
|---|---|
| Scaffold | Affitin |
| Total HTPS Reads | 10380 |
| Unique HTPS Reads | 1131 |
| Top Variant % of Total Reads | 45 |
| Final Phage Titer (PFU/mL) | 1.78E+09 |

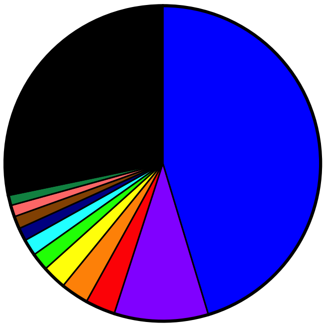

```
Library: GSVKVKF???GEEKEVDTSKI??V?R?GK?V?F?YDDNGK?G?G?V?EKDAPKELLDMLARAEREKKL
45.33%   GSVKVKFTAHGEEKEVDTSKIHLVCRGGKIVAFLYDDNGKLGHGSVGEKDAPKELLDMLARAEREKKL
9.65%    GSVKVKFPNPGEEKEVDTSKIVLVCRAGKIVHFYYDDNGKSG*WSR*EKDAPKELLDMLARAEREKKL
3.06%    GSVKVKFSSYGEEKEVDTSKIHIVCRDGKIVGFFYDDNGKPGYGRVGEKDAPKELLDMLARAEREKKL
2.84%    GSVKVKFPNPGEEKEVDTSKIHLVCRGGKIVAFLYDDNGKLGHGSVGEKDAPKELLDMLARAEREKKL
2.47%    GSVKVKFTAHGEEKEVDTSKIVLVCRAGKIVHFYYDDNGKSG*WSR*EKDAPKELLDMLARAEREKKL
1.84%    GSVKVKFSSYGEEKEVDTSKIHLVCRGGKIVAFLYDDNGKLGHGSVGEKDAPKELLDMLARAEREKKL
1.70%    GSVKVKFTAHGEEKEVDTSKIHIVCRDGKIVGFFYDDNGKPGYGRVGEKDAPKELLDMLARAEREKKL
1.34%    GSVKVKFTAHGEEKEVDTSKIHLVCRGGKIVAFLYDDNGKPGYGRVGEKDAPKELLDMLARAEREKKL
1.30%    GSVKVKF***GEEKEVDTSKIRSVWRGGKWVDFSYDDNGKQGYGYVLEKDAPKELLDMLARAEREKKL
1.18%    GSVKVKFRIDGEEKEVDTSKICYVYRIGKAVIFIYDDNGKRGLGLVFEKDAPKELLDMLARAEREKKL
1.04%    GSVKVKFNGPGEEKEVDTSKILYVNRTGKVVHFLYDDNGKNGYGSVHEKDAPKELLDMLARAEREKKL
28.26%   Other Sequence Less than 1% Total Reads
```

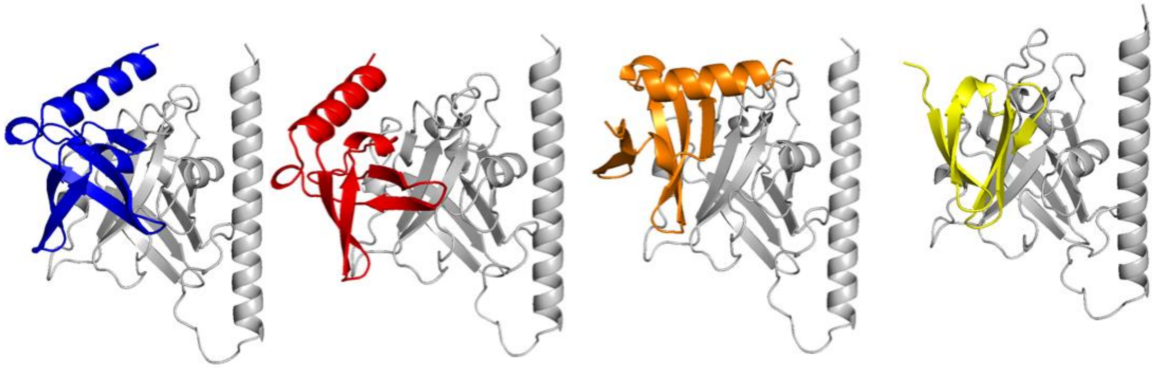

| pLDDT = 66.6<br>pTM = 0.579<br>ipTM = 0.149 | pLDDT = 70.9<br>pTM = 0.591<br>ipTM = 0.154 | pLDDT = 68.4<br>pTM = 0.594<br>ipTM = 0.159 | pLDDT = 64.6<br>pTM = 0.623<br>ipTM = 0.174 |
|---|---|---|---|

| Target | DltD (ef) |
|---|---|
| Scaffold | Affitin |
| Total HTPS Reads | 8084 |
| Unique HTPS Reads | 696 |
| Top Variant % of Total Reads | 76 |
| Final Phage Titer (PFU/mL) | 2.82E+08 |

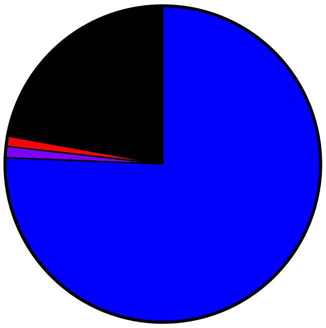

Library: GSVKVKF???GEEKEVDTSKI??V?R?GK?V?F?YDDNGK?G?G?V?EKDAPKELLDMLARAEREKKL

75.64% GSVKVKFTDCGEEKEVDTSKITYVIRFGKFVDFDYDDNGKIGSGAVREKDAPKELLDMLARAEREKKL

1.15% GSVKVKFNGPGEEKEVDTSKILYVNRTGKVVHFLYDDNGKNGYGSVHEKDAPKELLDMLARAEREKKL

1.04% GSVKVKFRIDGEEKEVDTSKICYVYRIGKAVIFIYDDNGKRGLGLVFEKDAPKELLDMLARAEREKKL

22.17% Other Sequence Less than 1% Total Reads

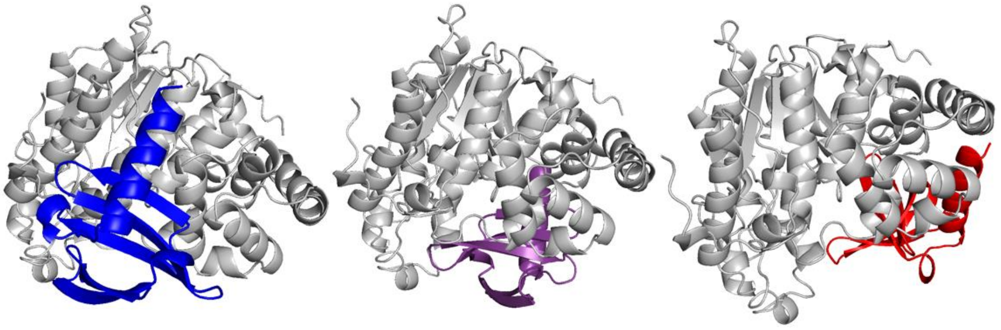

pLDDT = 92.6
pTM = 0.915
ipTM = 0.708

pLDDT = 86.4
pTM = 0.824
ipTM = 0.162

pLDDT = 86.4
pTM = 0.815
ipTM = 0.144

| Target | SH3BP5 |
|---|---|
| Scaffold | Affitin |
| Total HTPS Reads | 10940 |
| Unique HTPS Reads | 776 |
| Top Variant % of Total Reads | 79 |
| Final Phage Titer (PFU/mL) | 1.32E+09 |

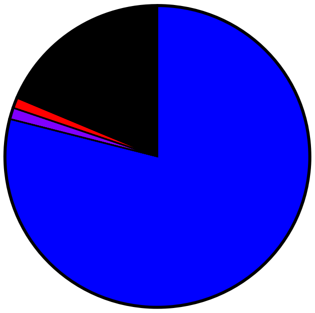

```
Library: GSVKVKF???GEEKEVDTSKI??V?R?GK?V?F?YDDNGK?G?G?V?EKDAPKELLDMLARAEREKKL
78.99%   GSVKVKFPIVGEEKEVDTSKIHYVYRYGKTVAFFYDDNGKGGYGSVLEKDAPKELLDMLARAEREKKL
1.23%    GSVKVKF***GEEKEVDTSKIRSVWRGGKWVDFSYDDNGKQGYGYVLEKDAPKELLDMLARAEREKKL
1.12%    GSVKVKFHGTGEEKEVDTSKISVVIRAGKTVYFSYDDNGKRGTGRVSEKDAPKELLDMLARAEREKKL
18.65%   Other Sequence Less than 1% Total Reads
```

**Insufficient Confidence to Predict Complex Structure (pTM)**

pLDDT = 75.3
pTM = 0.358
ipTM = 0.110

**Insufficient Confidence to Predict Complex Structure (pTM)**

pLDDT = 76.3
pTM = 0.352
ipTM = 0.090

| | |
|---|---|
| Target | OTUBI |
| Scaffold | Affitin |
| Total HTPS Reads | 11981 |
| Unique HTPS Reads | 885 |
| Top Variant % of Total Reads | 79 |
| Final Phage Titer (PFU/mL) | 2.40E+07 |

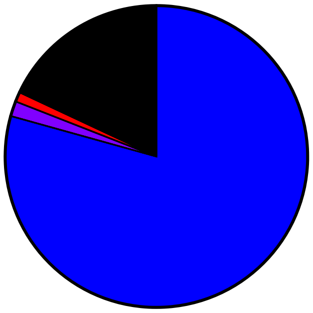

Library: GSVKVKF???GEEKEVDTSKI??V?R?GK?V?F?YDDNGK?G?G?V?EKDAPKELLDMLARAEREKKL

79.33% GSVKVKFDACGEEKEVDTSKIYFVRRNGKTVAFTYDDNGNTVAVTFLEKDAPKELLDMLARAEREKKL

1.62% GSVKVKF***GEEKEVDTSKIRSVWRGGKWVDFSYDDNGKQGYGYVLEKDAPKELLDMLARAEREKKL

1.04% GSVKVKFHGTGEEKEVDTSKISVVIRAGKTVYFSYDDNGKRGTGRVSEKDAPKELLDMLARAEREKKL

18.01% Other Sequence Less than 1% Total Reads

**Insufficient Confidence to Predict Complex Structure (pLDDT, pTM)**

pLDDT = 45.0
pTM = 0.400
ipTM = 0.175

**Insufficient Confidence to Predict Complex Structure (pLDDT, pTM)**

pLDDT = 49.1
pTM = 0.403
ipTM = 0.115

| Target | DltD (strep) |
|---|---|
| Scaffold | Affitin |
| Total HTPS Reads | 13357 |
| Unique HTPS Reads | 1100 |
| Top Variant % of Total Reads | 36 |
| Final Phage Titer (PFU/mL) | 4.57E+08 |

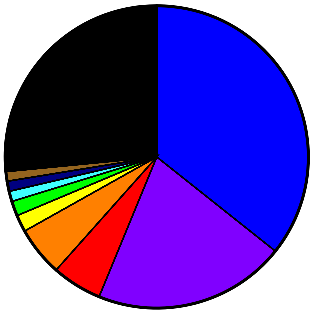

```
Library: GSVKVKF???GEEKEVDTSKI??V?R?GK?V?F?YDDNGK?G?G?V?EKDAPKELLDMLARAEREKKL
35.70%   GSVKVKFSHAGEEKEVDTSKIRIVCRAGKLVYFLYDDNGKRGCGLVYEKDAPKELLDMLARAEREKKL
20.51%   GSVKVKFFHTGEEKEVDTSKIRVVLRAGKLVRFFYDDNGKFGCGTVNEKDAPKELLDMLARAEREKKL
5.34%    GSVKVKFFHTGEEKEVDTSKIRIVCRAGKLVYFLYDDNGKRGCGLVYEKDAPKELLDMLARAEREKKL
5.29%    GSVKVKFSHAGEEKEVDTSKIRVVLRAGKLVRFFYDDNGKFGCGTVNEKDAPKELLDMLARAEREKKL
1.81%    GSVKVKFLIHGEEKEVDTSKIAVVIRAGKLVRFLYDDNGKPGCGLVNEKDAPKELLDMLARAEREKKL
1.59%    GSVKVKFSHAGEEKEVDTSKIRIVCRAGKLVYFLYDDNGKFGCGTVNEKDAPKELLDMLARAEREKKL
1.10%    GSVKVKFFHTGEEKEVDTSKIRVVLRAGKLVRFFYDDNGKRGCGLVYEKDAPKELLDMLARAEREKKL
1.09%   GSVKVKFHGTGEEKEVDTSKISVVIRAGKTVYFSYDDNGKRGTGRVSEKDAPKELLDMLARAEREKKL
1.03%    GSVKVKF***GEEKEVDTSKIRSVWRGGKWVDFSYDDNGKQGYGYVLEKDAPKELLDMLARAEREKKL
26.55%   Other Sequence Less than 1% Total Reads
```

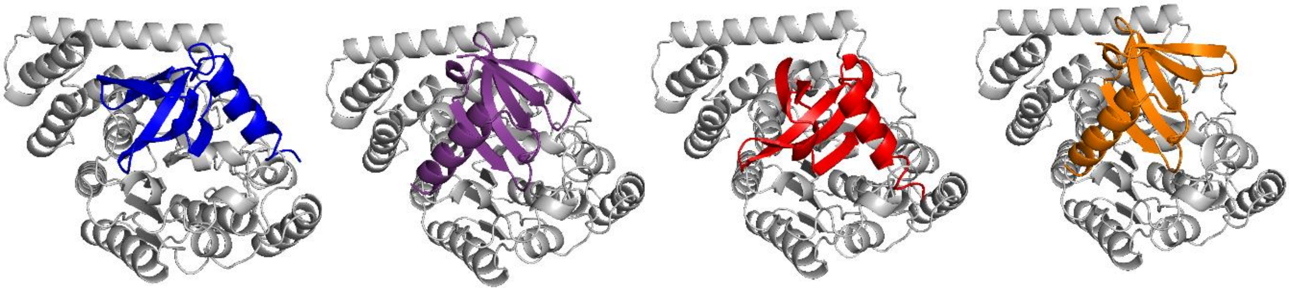

| | | | |
|---|---|---|---|
| pLDDT = 86.8 | pLDDT = 86.8 | pLDDT = 86.5 | pLDDT = 86.7 |
| pTM = 0.839 | pTM = 0.841 | pTM = 0.842 | pTM = 0.837 |
| ipTM = 0.302 | ipTM = 0.308 | ipTM = 0.319 | ipTM = 0.287 |

| Target | RB1 |
| --- | --- |
| Scaffold | Affitin |
| Total HTPS Reads | 5315 |
| Unique HTPS Reads | 350 |
| Top Variant % of Total Reads | 83 |
| Final Phage Titer (PFU/mL) | 2.95E+08 |

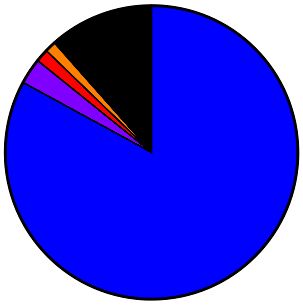

```
Library: GSVKVKF???GEEKEVDTSKI??V?R?GK?V?F?YDDNGK?G?G?V?EKDAPKELLDMLARAEREKKL
82.92%   GSVKVKFSYSGEEKEVDTSKISIVHRNGKTVCFFYDDNGKCGCGIVNEKDAPKELLDMLARAEREKKL
2.86%    GSVKVKFTYSGEEKEVDTSKIGSVIRNGKSVAFAYDDNGKLGYGYVSEKDAPKELLDMLARAEREKKL
1.43%    GSVKVKF***GEEKEVDTSKIRSVWRGGKWVDFSYDDNGKQGYGYVLEKDAPKELLDMLARAEREKKL
1.22%    GSVKVKFHIPGEEKEVDTSKIFIVLRYGKFVNFLYDDNGKAGYGYVHEKDAPKELLDMLARAEREKKL
11.57%   Other Sequence less than 1%
```

**Insufficient Confidence to Predict Complex Structure (pTM)**

pLDDT = 58.7
pTM = 0.449
ipTM = 0.294

**Insufficient Confidence to Predict Complex Structure (pTM)**

pLDDT = 53.6
pTM = 0.478
ipTM = 0.458

**Insufficient Confidence to Predict Complex Structure (pTM)**

pLDDT = 50.5
pTM = 0.431
ipTM = 0.171

| Target | SasG (930-1060) |
|---|---|
| Scaffold | Affitin |
| Total HTPS Reads | 7836 |
| Unique HTPS Reads | 1786 |
| Top Variant % of Total Reads | 31 |
| Final Phage Titer (PFU/mL) | 5.89E+08 |

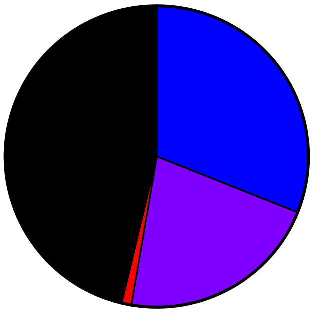

Library: GSVKVKF???GEEKEVDTSKI??V?R?GK?V?F?YDDNGK?G?G?V?EKDAPKELLDMLARAEREKKL

31.05% GSVKVKFHVNGEEKRS*YF*NSSRSPLW*RRSLSL***W*IRSWLRCEKDAPKELLDMLARAEREKKL

21.63% GSVKVKF***GEEKEVDTSKIRSVWRGGKWVDFSYDDNGKQGYGYVLEKDAPKELLDMLARAEREKKL

1.03% GSVKVKFHGTGEEKEVDTSKISVVIRAGKTVYFSYDDNGKRGTGRVSEKDAPKELLDMLARAEREKKL

46.28% Other Sequence Less than 1% Total Reads

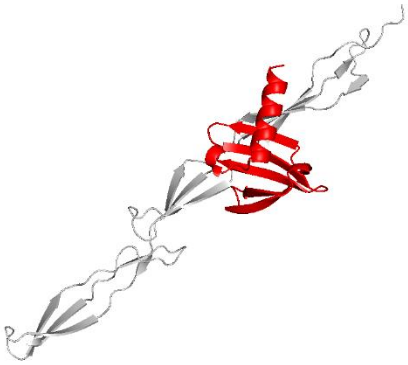

pLDDT = 78.6
pTM = 0.546
ipTM = 0.233

| Target | BRD4 |
|---|---|
| Scaffold | Affitin |
| Total HTPS Reads | 4849 |
| Unique HTPS Reads | 1491 |
| Top Variant % of Total Reads | 48 |
| Final Phage Titer (PFU/mL) | 5.75E+08 |

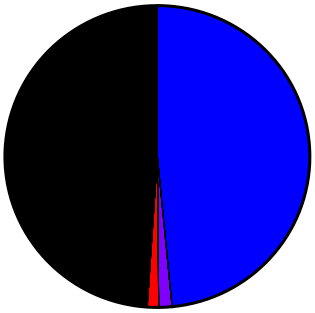

Library: GSVKVKF???GEEKEVDTSKI??V?R?GK?V?F?YDDNGK?G?G?V?EKDAPKELLDMLARAEREKKL

48.46% GSVKVKF***GEEKEVDTSKIRSVWRGGKWVDFSYDDNGKQGYGYVLEKDAPKELLDMLARAEREKKL

1.40% GSVKVKFHGTGEEKEVDTSKISVVIRAGKTVYFSYDDNGKRGTGRVSEKDAPKELLDMLARAEREKKL

1.20% GSVKVKFHGTGEEKEVDTSKISVVIRAGKTVYFSYDDNGKRGTGRVSEKDAPKELLDMLARAEREKKL

48.94% Other Sequence Less than 1% Total Reads

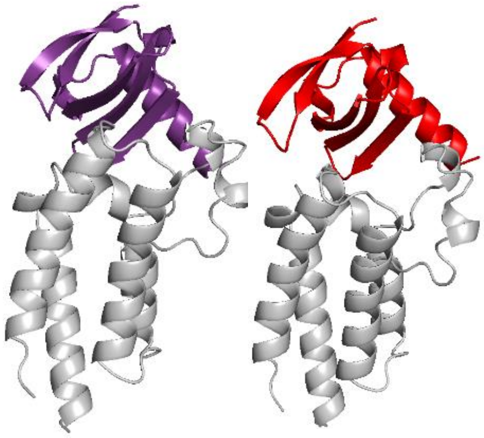

pLDDT = 79.1
pTM = 0.654
ipTM = 0.216

pLDDT = 79.1
pTM = 0.654
ipTM = 0.216

| Target | IsdB |
|---|---|
| Scaffold | Affitin |
| Total HTPS Reads | 13922 |
| Unique HTPS Reads | 1951 |
| Top Variant % of Total Reads | 36 |
| Final Phage Titer (PFU/mL) | 3.09E+08 |

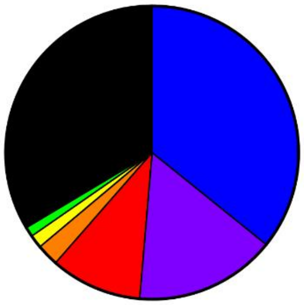

```
Library: GSVKVKF???GEEKEVDTSKI??V?R?GK?V?F?YDDNGK?G?G?V?EKDAPKELLDMLARAEREKKL
35.79%   GSVKVKFHVNGEEKRS*YF*NSSRSPLW*RRSLSL***W*IRSWLRCEKDAPKELLDMLARAEREKKL
15.57%   GSVKVKFHAIGEEKEVDTSKISHVPRPGKTVSFLYDDNGKPGFGIVNEKDAPKELLDMLARAEREKKL
10.11%   GSVKVKF***GEEKEVDTSKIRSVWRGGKWVDFSYDDNGKQGYGYVLEKDAPKELLDMLARAEREKKL
 2.43%   GSVKVKFTDHGEEKEVDTSKITVVHRNGKHVIFCYDDNGKSGSGSVLEKDAPKELLDMLARAEREKKL
 1.47%   GSVKVKFHAIGEEKEVDTSKIRRVPRYGKGVHFLYDDNGKSGPGCVVEKDAPKELLDMLARAEREKKL
 1.08%   GSVKVKFADLGEEKEVDTSKISHVVRSGKPVAFTYDDNGKPGHGSVHEKDAPKELLDMLARAEREKKL
33.56%   Other Sequence Less Than 1% Total Reads
```

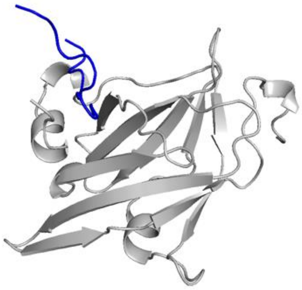

pLDDT = 84.8
pTM = 0.827
ipTM = 0.302

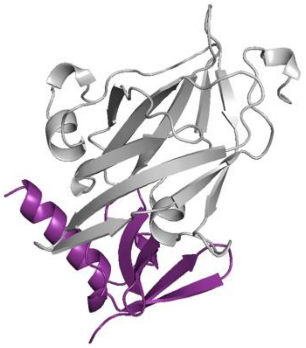

pLDDT = 74.5
pTM = 0.673
ipTM = 0.163

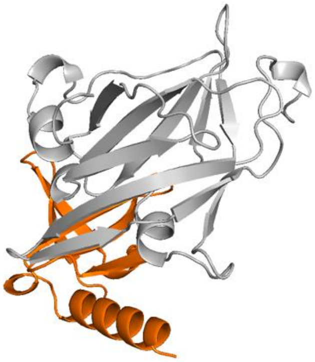

pLDDT = 84.0
pTM = 0.678
ipTM = 0.173

| Target | HEWL |
|---|---|
| Scaffold | Affitin |
| Total HTPS Reads | 10773 |
| Unique HTPS Reads | 794 |
| Top Variant % of Total Reads | 43 |
| Final Phage Titer (PFU/mL) | 2.19E+08 |

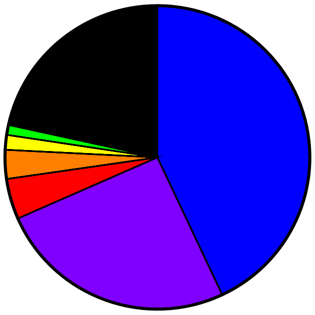

```
Library: GSVKVKF???GEEKEVDTSKI??V?R?GK?V?F?YDDNGK?G?G?V?EKDAPKELLDMLARAEREKKL
43.01%   GSVKVKFLAFGEEKEVDTSKIHLVRRFGKGVLFRYDDNGKRGSGIVYEKDAPKELLDMLARAEREKKL
25.40%   GSVKVKFNAAGEEKEVDTSKIDVVFRHGKCVFFLYDDNGKHGLGLVIEKDAPKELLDMLARAEREKKL
4.32%    GSVKVKFNAAGEEKEVDTSKIHLVRRFGKGVLFRYDDNGKRGSGIVYEKDAPKELLDMLARAEREKKL
3.08%    GSVKVKFLAFGEEKEVDTSKIDVVFRHGKCVFFLYDDNGKHGLGLVIEKDAPKELLDMLARAEREKKL
1.58%    GSVKVKFLAFGEEKEVDTSKIHLVRRFGKGVLFRYDDNGKHGLGLVIEKDAPKELLDMLARAEREKKL
1.06%    GSVKVKFHGTGEEKEVDTSKISVVIRAGKTVYFSYDDNGKRGTGRVSEKDAPKELLDMLARAEREKKL
21.56%   Other Sequence Less than 1% Total Reads
```

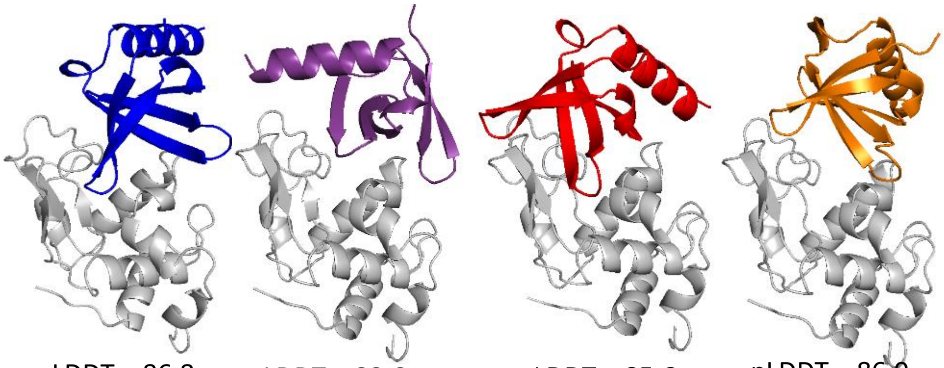

| | | | |
|---|---|---|---|
| pLDDT = 86.8 | pLDDT = 88.6 | pLDDT = 85.6 | pLDDT = 86.0 |
| pTM = 0.665 | pTM = 0.746 | pTM = 0.669 | pTM = 0.690 |
| ipTM = 0.173 | ipTM = 0.412 | ipTM = 0.179 | ipTM = 0.243 |

| Target | HSPB1 (90-171) |
|---|---|
| Scaffold | Affitin |
| Total HTPS Reads | 10892 |
| Unique HTPS Reads | 1656 |
| Top Variant % of Total Reads | 16 |
| Final Phage Titer (PFU/mL) | 4.68E+08 |

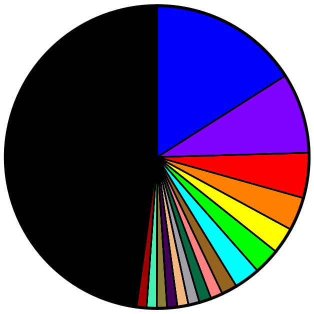

```
Library: GSVKVKF???GEEKEVDTSKI??V?R?GK?V?F?YDDNGK?G?G?V?EKDAPKELLDMLARAEREKKL
15.96%   GSVKVKFNPHGEEKEVDTSKIDPVFRAGKTVHFTYDDNGKFGCGPVNEKDAPKELLDMLARAEREKKL
8.58%    GSVKVKFTTPGEEKEVDTSKIDPVFRAGKTVLFTYDDNGKCGDGYVFEKDAPKELLDMLARAEREKKL
4.82%    GSVKVKFHPYGEEKEVDTSKIDPVFRAGKTVYFYYDDNGKDGNGYVCEKDAPKELLDMLARAEREKKL
3.57%    GSVKVKFLLPGEEKEVDTSKIDPVFRAGKTVAFIYDDNGKNGFGCVHEKDAPKELLDMLARAEREKKL
2.80%    GSVKVKFTTPGEEKEVDTSKIDPVFRAGKTVHFTYDDNGKFGCGPVNEKDAPKELLDMLARAEREKKL
2.79%    GSVKVKFHPYGEEKEVDTSKIDPVFRAGKTVHFTYDDNGKFGCGPVNEKDAPKELLDMLARAEREKKL
2.75%    GSVKVKFHPYGEEKEVDTSKIDPVFRAGKTVHFTYDDNGKFGCGPVNEKDAPKELLDMLARAEREKKL
1.67%    GSVKVKFNPHGEEKEVDTSKIDPVFRAGKTVYFYYDDNGKDGNGYVCEKDAPKELLDMLARAEREKKL
1.29%    GSVKVKFNPHGEEKEVDTSKIDPVFRAGKTVLFTYDDNGKCGDGYVFEKDAPKELLDMLARAEREKKL
1.29%    GSVKVKFTTPGEEKEVDTSKIDPVFRAGKTVHFTYDDNGKFGCGPVNEKDAPKELLDMLARAEREKKL
1.18%    GSVKVKFPTLGEEKEVDTSKIVIVNRIGKYVRFLYDDNGKHGGGYVLEKDAPKELLDMLARAEREKKL
1.18%    GSVKVKFNPHGEEKEVDTSKIDPVFRAGKTVHFTYDDNGKCGDGYVFEKDAPKELLDMLARAEREKKL
1.09%    GSVKVKFHGTGEEKEVDTSKISVVIRAGKTVYFSYDDNGKRGTGRVSEKDAPKELLDMLARAEREKKL
1.08%    GSVKVKFLLPGEEKEVDTSKIDPVFRAGKTVHFTYDDNGKFGCGPVNEKDAPKELLDMLARAEREKKL
1.08%    GSVKVKF***GEEKEVDTSKIRSVWRGGKWVDFSYDDNGKQGYGYVLEKDAPKELLDMLARAEREKKL
1.06%    GSVKVKFNPHGEEKEVDTSKIDPVFRAGKTVAFIYDDNGKNGFGCVHEKDAPKELLDMLARAEREKKL
47.81%   Other Sequence Less than 1% Total Reads
```

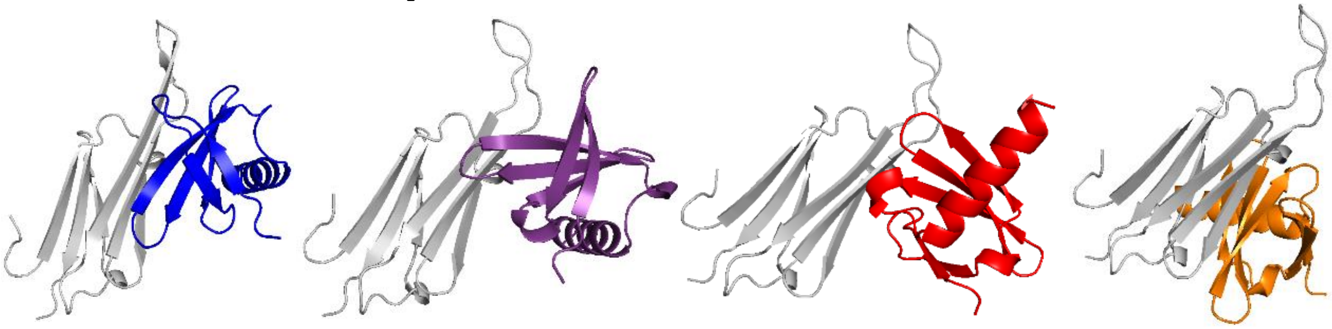

pLDDT = 77.9
pTM = 0.625
ipTM = 0.428

pLDDT = 82.3
pTM = 0.613
ipTM = 0.324

pLDDT = 81.4
pTM = 0.528
ipTM = 0.172

pLDDT = 72.8
pTM = 0.519
ipTM = 0.208

| Target | KRAS G12C |
|---|---|
| Scaffold | Affitin |
| Total HTPS Reads | 9880 |
| Unique HTPS Reads | 686 |
| Top Variant % of Total Reads | 79 |
| Final Phage Titer (PFU/mL) | 3.47E+08 |

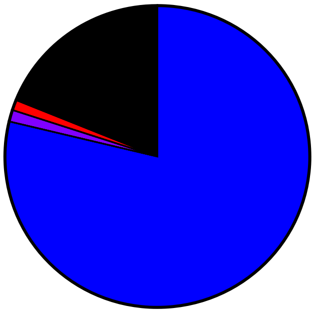

```
Library:  GSVKVKF???GEEKEVDTSKI??V?R?GK?V?F?YDDNGK?G?G?V?EKDAPKELLDMLARAEREKKL
  78.73%  GSVKVKFIFSGEEKEVDTSKIHAVFRNGKYVTFFYDDNGKHGFGSVLEKDAPKELLDMLARAEREKKL
  1.26%   GSVKVKF***GEEKEVDTSKIRSVWRGGKWVDFSYDDNGKQGYGYVLEKDAPKELLDMLARAEREKKL
  1.11%   GSVKVKFHGTGEEKEVDTSKISVVIRAGKTVYFSYDDNGKRGTGRVSEKDAPKELLDMLARAEREKKL
  18.90%  Other Sequence Less than 1% Total Reads
```

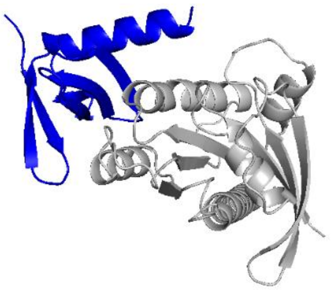

pLDDT = 86.6
pTM = 0.809
ipTM = 0.640

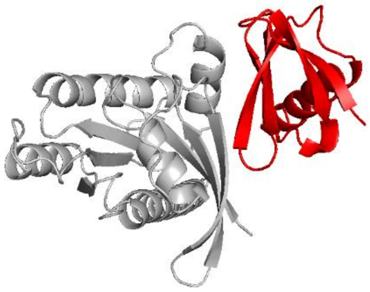

pLDDT = 80.2
pTM = 0.692
ipTM = 0.276

| | |
|---|---|
| Target | LMO2 |
| Scaffold | Affitin |
| Total HTPS Reads | 11742 |
| Unique HTPS Reads | 905 |
| Top Variant % of Total Reads | 53 |
| Final Phage Titer (PFU/mL) | 3.98E+08 |

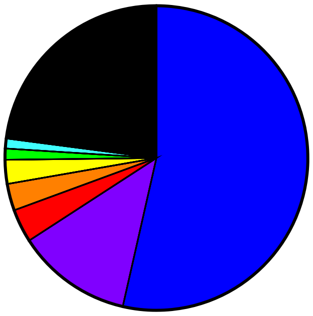

```
Library: GSVKVKF???GEEKEVDTSKI??V?R?GK?V?F?YDDNGK?G?G?V?EKDAPKELLDMLARAEREKKL
53.57%   GSVKVKFICRGEEKEVDTSKIYRVYRCGKAVSFDYDDNGKSGFGFVIEKDAPKELLDMLARAEREKKL
12.31%   GSVKVKFISRGEEKEVDTSKILRVTRIGKAVTFDYDDNGKCGPGFVDEKDAPKELLDMLARAEREKKL
3.54%    GSVKVKFICRGEEKEVDTSKILRVTRIGKAVTFDYDDNGKCGPGFVDEKDAPKELLDMLARAEREKKL
2.84%    GSVKVKFISRGEEKEVDTSKIYRVYRCGKAVSFDYDDNGKSGFGFVIEKDAPKELLDMLARAEREKKL
2.55%    GSVKVKFICRGEEKEVDTSKIYRVYRCGKAVSFDYDDNGKCGPGFVDEKDAPKELLDMLARAEREKKL
1.18%    GSVKVKFICRGEEKEVDTSKIYRVYRCGKAVTFDYDDNGKCGPGFVDEKDAPKELLDMLARAEREKKL
1.06%    GSVKVKFHGTGEEKEVDTSKISVVIRAGKTVYFSYDDNGKRGTGRVSEKDAPKELLDMLARAEREKKL
22.94%   Other Sequence Less than 1% Total Reads
```

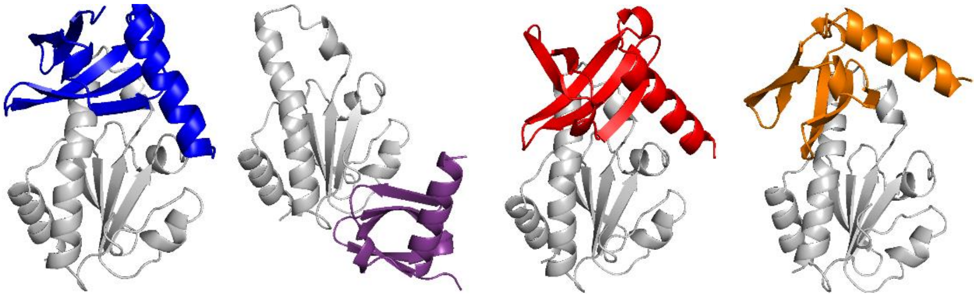

| | | | |
|---|---|---|---|
| pLDDT = 81.3 | pLDDT = 76.2 | pLDDT = 80.1 | pLDDT = 79.9 |
| pTM = 0.602 | pTM = 0.591 | pTM = 0.591 | pTM = 0.591 |
| ipTM = 0.182 | ipTM = 0.169 | ipTM = 0.160 | ipTM = 0.169 |

| Target | HIF1a |
|---|---|
| Scaffold | Affitin |
| Total HTPS Reads | 15079 |
| Unique HTPS Reads | 1161 |
| Top Variant % of Total Reads | 44 |
| Final Phage Titer (PFU/mL) | 4.47E+08 |

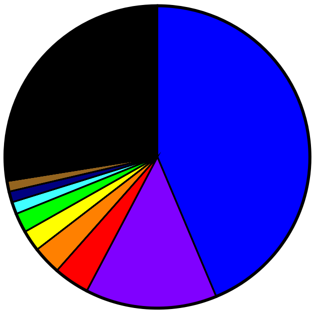

| | |
|---|---|
| Library: | GSVKVKF???GEEKEVDTSKI??V?R?GK?V?F?YYDDNGK?G?G?V?EKDAPKELLDMLARAEREKKL |
| 43.70% | GSVKVKFISHGEEKEVDTSKIRYVYRGGKLVYFYYDDNGKDGDGLVNEKDAPKELLDMLARAEREKKL |
| 13.97% | GSVKVKFGVSGEEKEVDTSKIYPVTRDGKSVHFYYDDNGKYGYGSVSEKDAPKELLDMLARAEREKKL |
| 3.79% | GSVKVKFTHSGEEKEVDTSKIYPVYRGGKSVHFFYDDNGKFGRGVVHEKDAPKELLDMLARAEREKKL |
| 3.04% | GSVKVKFTPGGEEKEVDTSKIYPVYRCGKIVHFFYDDNGKYGYGHVFEKDAPKELLDMLARAEREKKL |
| 2.26% | GSVKVKFGVSGEEKEVDTSKIRYVYRGGKLVYFYYDDNGKDGDGLVNEKDAPKELLDMLARAEREKKL |
| 2.10% | GSVKVKFISHGEEKEVDTSKIYPVTRDGKSVHFYYDDNGKYGYGSVSEKDAPKELLDMLARAEREKKL |
| 1.29% | GSVKVKFTHSGEEKEVDTSKIRYVYRGGKLVYFYYDDNGKDGDGLVNEKDAPKELLDMLARAEREKKL |
| 1.16% | GSVKVKFLATGEEKEVDTSKIHVVVRNGKHVHFFYDDNGKFGPGTVYEKDAPKELLDMLARAEREKKL |
| 1.13% | GSVKVKFTPGGEEKEVDTSKIYPVTRDGKSVHFYYDDNGKYGYGSVSEKDAPKELLDMLARAEREKKL |
| 27.56% | Other Sequence Less than 1% Total Reads |

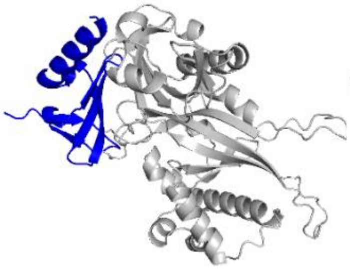

pLDDT = 69.8
pTM = 0.569
ipTM = 0.169

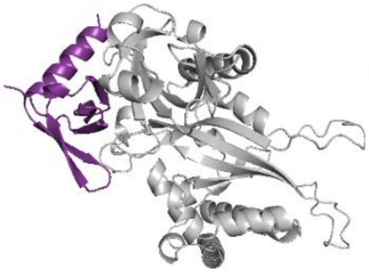

pLDDT = 72.6
pTM = 0.543
ipTM = 0.255

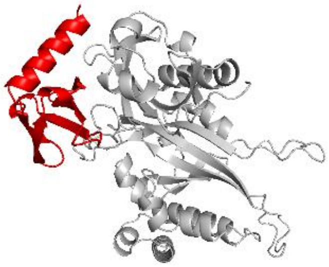

pLDDT = 71.4
pTM = 0.568
ipTM = 0.231

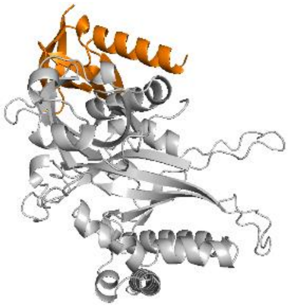

pLDDT = 69.2
pTM = 0.605
ipTM = 0.205

| Target | UEV |
|---|---|
| Scaffold | Affitin |
| Total HTPS Reads | 15123 |
| Unique HTPS Reads | 1191 |
| Top Variant % of Total Reads | 49 |
| Final Phage Titer (PFU/mL) | 1.55E+09 |

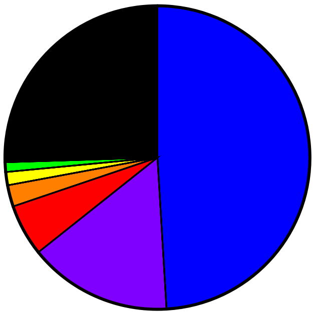

| | |
|---|---|
| Library: | GSVKVKF???GEEKEVDTSKI??V?R?GK?V?F?YDDNGK?G?G?V?EKDAPKELLDMLARAEREKKL |
| 49.05% | GSVKVKFSYSGEEKEVDTSKISIVHRNGKTVCFFYDDNGKCGCGIVNEKDAPKELLDMLARAEREKKL |
| 15.22% | GSVKVKFIYNGEEKEVDTSKILSVYRCGKDVRFLYDDNGKLGGGDVGEKDAPKELLDMLARAEREKKL |
| 5.51% | GSVKVKFSYSGEEKEVDTSKILSVYRCGKDVRFLYDDNGKLGGGDVGEKDAPKELLDMLARAEREKKL |
| 2.29% | GSVKVKFIYNGEEKEVDTSKISIVHRNGKTVCFFYDDNGKCGCGIVNEKDAPKELLDMLARAEREKKL |
| 1.41% | GSVKVKF***GEEKEVDTSKIRSVWRGGKWVDFSYDDNGKQGYGYVLEKDAPKELLDMLARAEREKKL |
| 1.03% | GSVKVKFTYSGEEKEVDTSKIGSVIRNGKSVAFAYDDNGKLGYGYVSEKDAPKELLDMLARAEREKKL |
| 25.49% | Other Sequence Less than 1% Total Reads |

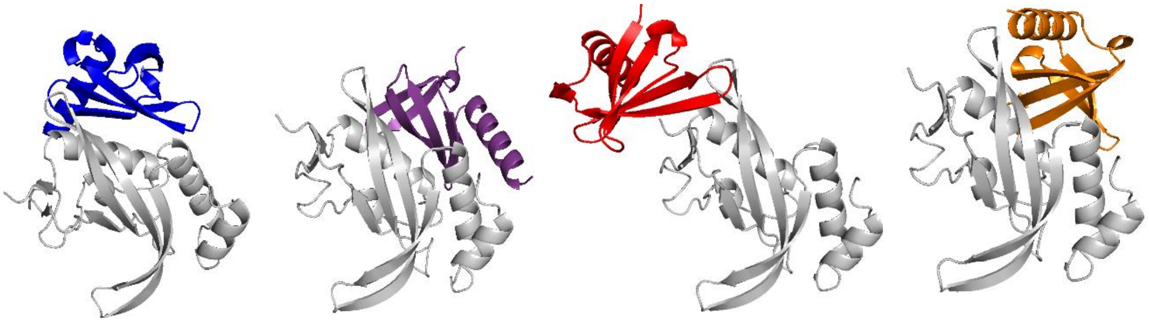

pLDDT = 82.5
pTM = 0.686
ipTM = 0.202

pLDDT = 84.4
pTM = 0.677
ipTM = 0.190

pLDDT = 93.1
pTM = 0.852
ipTM = 0.729

pLDDT = 83.3
pTM = 0.684
ipTM = 0.195

| Target | FRB |
|---|---|
| Scaffold | Affitin |
| Total HTPS Reads | 6751 |
| Unique HTPS Reads | 4216 |
| Top Variant % of Total Reads | 3 |
| Final Phage Titer (PFU/mL) | 5.75E+08 |

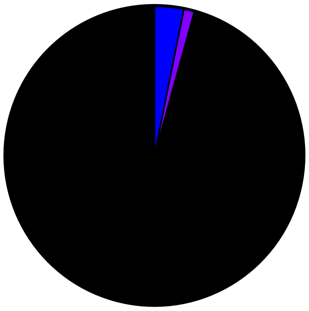

```
Library: GSVKVKF???GEEKEVDTSKI??V?R?GK?V?F?YDDNGK?G?G?V?EKDAPKELLDMLARAEREKKL
3.17%    GSVKVKF***GEEKEVDTSKIRSVWRGGKWVDFSYDDNGKQGYGYVLEKDAPKELLDMLARAEREKKL
1.16%    GSVKVKFHGTGEEKEVDTSKISVVIRAGKTVYFSYDDNGKRGTGRVSEKDAPKELLDMLARAEREKKL
95.67%   Other Sequence Less than 1% Total Reads
```

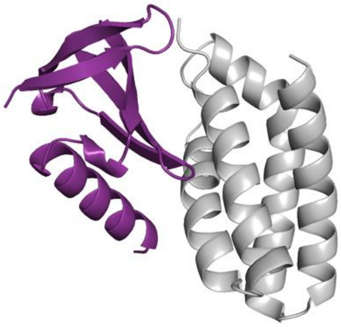

pLDDT = 85.6
pTM = 0.812
ipTM = 0.688

| Target | PD-1 |
|---|---|
| Scaffold | Affitin |
| Total HTPS Reads | 5055 |
| Unique HTPS Reads | 365 |
| Top Variant % of Total Reads | 85 |
| Final Phage Titer (PFU/mL) | 1.91E+08 |

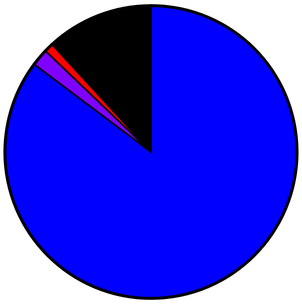

```
Library: GSVKVKF???GEEKEVDTSKI??V?R?GK?V?F?YDDNGK?G?G?V?EKDAPKELLDMLARAEREKKL
85.24%   GSVKVKFHIPGEEKEVDTSKIFIVLRYGKFVNFLYDDNGKAGYGYVHEKDAPKELLDMLARAEREKKL
1.90%    GSVKVKF***GEEKEVDTSKIRSVWRGGKWVDFSYDDNGKQGYGYVLEKDAPKELLDMLARAEREKKL
1.01%    GSVKVKFLIHGEEKEVDTSKIIFVNRYGKGVTFFYDDNGKYGLGHVTEKDAPKELLDMLARAEREKKL
11.85%   Other Sequence less than 1%
```

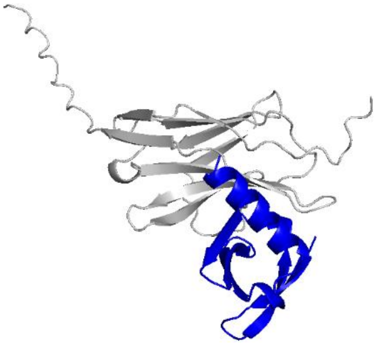

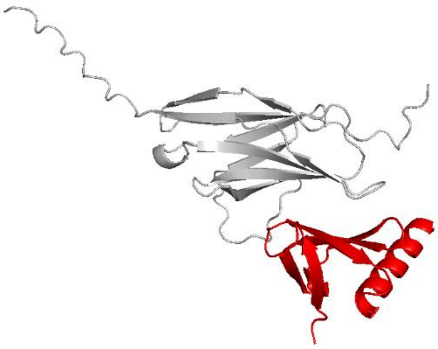

pLDDT = 72.2
pTM = 0.582
ipTM = 0.212

pLDDT = 77.0
pTM = 0.609
ipTM = 0.224

| Target | SasA |
|---|---|
| Scaffold | Affitin |
| Total HTPS Reads | 9987 |
| Unique HTPS Reads | 2668 |
| Top Variant % of Total Reads | 6 |
| Final Phage Titer (PFU/mL) | 7.41E+08 |

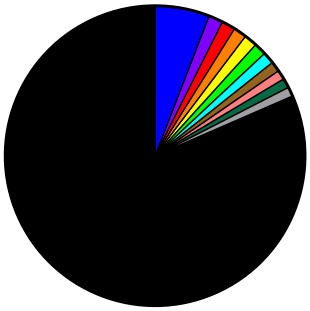

Library: GSVKVKF???GEEKEVDTSKI??V?R?GK?V?F?YDDNGK?G?G?V?EKDAPKELLDMLARAEREKKL

5.87% GSVKVKF***GEEKEVDTSKIRSVWRGGKWVDFSYDDNGKQGYGYVLEKDAPKELLDMLARAEREKKL

1.54% GSVKVKFIILGEEKEVDTSKIRSVWRGGKWVDFSYDDNGKQGYGYVLEKDAPKELLDMLARAEREKKL

1.48% GSVKVKFDILGEEKEVDTSKIRSVWRGGKWVDFSYDDNGKQGYGYVLEKDAPKELLDMLARAEREKKL

1.42% GSVKVKFPVRGEEKEVDTSKIRSVWRGGKWVDFSYDDNGKQGYGYVLEKDAPKELLDMLARAEREKKL

1.40% GSVKVKF-RRGEEKEVDTSKIRSVWRGGKWVDFSYDDNGKQGYGYVLEKDAPKELLDMLARAEREKKL

1.32% GSVKVKFARCGEEKEVDTSKIRSVWRGGKWVDFSYDDNGKQGYGYVLEKDAPKELLDMLARAEREKKL

1.27% GSVKVKFLLCGEEKEVDTSKIRSVWRGGKWVDFSYDDNGKQGYGYVLEKDAPKELLDMLARAEREKKL

1.09% GSVKVKFHVIGEEKEVDTSKIRSVWRGGKWVDFSYDDNGKQGYGYVLEKDAPKELLDMLARAEREKKL

1.04% GSVKVKFCILGEEKEVDTSKIRSVWRGGKWVDFSYDDNGKQGYGYVLEKDAPKELLDMLARAEREKKL

1.02% GSVKVKFHGTGEEKEVDTSKISVVIRAGKTVYFSYDDNGKRGTGRVSEKDAPKELLDMLARAEREKKL

1.02% GSVKVKFHCLGE-KEVDTSKIRSVWRGGKWVDFSYDDNGKQGYGYVLEKDAPKELLDMLARAEREKKL

81.52% Other Sequence Less than 1% Total Reads

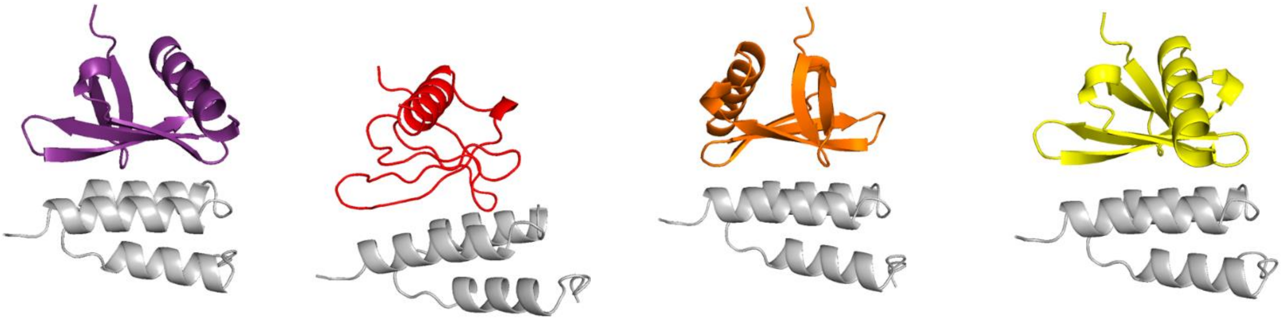

pLDDT = 80.2
pTM = 0.651
ipTM = 0.469

pLDDT = 67.2
pTM = 0.772
ipTM = 0.683

pLDDT = 87.1
pTM = 0.753
ipTM = 0.654

pLDDT = 78.5
pTM = 0.642
ipTM = 0.439

| | |
|---|---|
| Target | LC3B |
| Scaffold | Affitin |
| Total HTPS Reads | 15818 |
| Unique HTPS Reads | 1750 |
| Top Variant % of Total Reads | 36 |
| Final Phage Titer (PFU/mL) | 3.55E+08 |

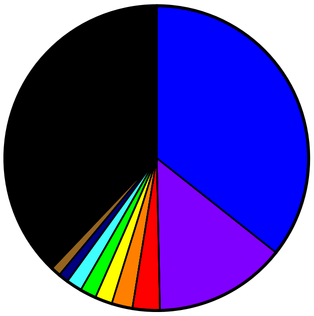

```
Library: GSVKVKF???GEEKEVDTSKI??V?R?GK?V?F?YDDNGK?G?G?V?EKDAPKELLDMLARAEREKKL
 35.64%  GSVKVKFATSGEEKEVDTSKIDPVARRGKIVAFFYDDNGKLGSGYVSEKDAPKELLDMLARAEREKKL
 14.03%  GSVKVKFLIHGEEKEVDTSKIHIVPRFGKYVLFAYDDNGKNGVGYVTEKDAPKELLDMLARAEREKKL
 2.91%   GSVKVKFASHGEEKEVDTSKIGCVPRTGKIVVFYYDDNGKLGSGTVNEKDAPKELLDMLARAEREKKL
 2.22%   GSVKVKFGGHGEEKEVDTSKISNVHRSGKCVYFTYDDNGKRGIGFVIEKDAPKELLDMLARAEREKKL
 1.83%   GSVKVKFATSGEEKEVDTSKIHIVPRFGKYVLFAYDDNGKNGVGYVTEKDAPKELLDMLARAEREKKL
 1.73%   GSVKVKFPTNGEEKEVDTSKIDPVRRAGKIVVFYYDDNGKLGCGHVTEKDAPKELLDMLARAEREKKL
 1.68%   GSVKVKFLIHGEEKEVDTSKIDPVARRGKIVAFFYDDNGKLGSGYVSEKDAPKELLDMLARAEREKKL
 1.07%   GSVKVKFHGTGEEKEVDTSKISVVIRAGKTVYFSYDDNGKRGTGRVSEKDAPKELLDMLARAEREKKL
 1.03%   GSVKVKFASHGEEKEVDTSKIDPVARRGKIVAFFYDDNGKLGSGYVSEKDAPKELLDMLARAEREKKL
 37.87%  Other Sequence Less than 1% Total Reads
```

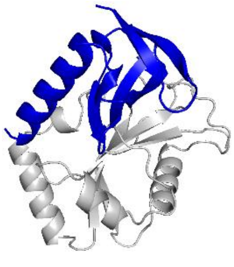

pLDDT = 77.9
pTM = 0.617
ipTM = 0.182

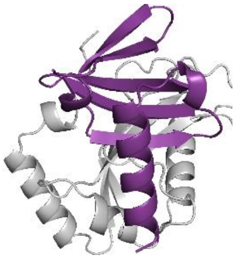

pLDDT = 78.6
pTM = 0.635
ipTM = 0.232

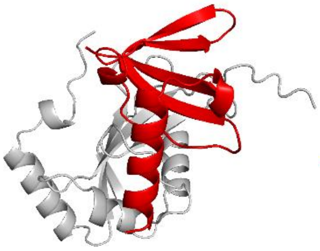

pLDDT = 77.6
pTM = 0.647
ipTM = 0.286

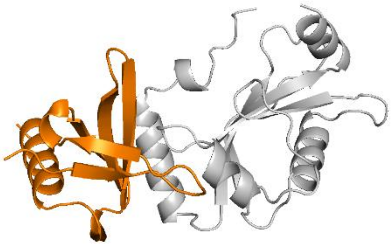

pLDDT = 80.2
pTM = 0.616
ipTM = 0.144

| | |
|---|---|
| Target | MST2 |
| Scaffold | Affitin |
| Total HTPS Reads | 13402 |
| Unique HTPS Reads | 1279 |
| Top Variant % of Total Reads | 46 |
| Final Phage Titer (PFU/mL) | 4.17E+08 |

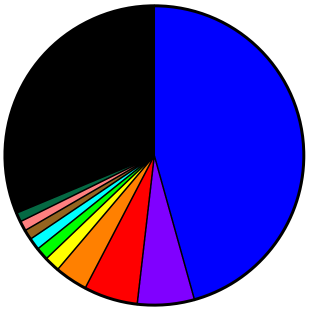

```
Library: GSVKVKF???GEEKEVDTSKI??V?R?GK?V?F?YDDNGK?G?G?V?EKDAPKELLDMLARAEREKKL
45.67%   GSVKVKFHVNGEEKRS*YF*NSSRSPLW*RRSLSL***W*IRSWLRCEKDAPKELLDMLARAEREKKL
6.16%    GSVKVKFIASGEEKEVDTSKISYVSRSGKSVSFNYDDNGKYGNGRVNEKDAPKELLDMLARAEREKKL
5.83%    GSVKVKFYGIGEEKEVDTSKIANVNRHGKPVPFGYDDNGKSGLGHVSEKDAPKELLDMLARAEREKKL
3.55%    GSVKVKFSLSGEEKEVDTSKINTVRRTGKHVHFTYDDNGKYGRGRVPEKDAPKELLDMLARAEREKKL
1.67%    GSVKVKFTAAGEEKEVDTSKIPSVIRYGKTVDFSYDDNGKPGNGSVSEKDAPKELLDMLARAEREKKL
1.39%    GSVKVKF***GEEKEVDTSKIRSVWRGGKWVDFSYDDNGKQGYGYVLEKDAPKELLDMLARAEREKKL
1.29%    GSVKVKFFRSGEEKEVDTSKISSVVRSGKPVAFYYDDNGKRGFGFVYEKDAPKELLDMLARAEREKKL
1.08%    GSVKVKFHVNGEEKRS*YF*NSSRSPLW*RRSLSL***W*IRSWSRQEKDAPKELLDMLARAEREKKL
1.07%    GSVKVKFYGIGEEKEVDTSKIRRVPRYGKGVHFLYDDNGKSGPGCVVEKDAPKELLDMLARAEREKKL
1.01%    GSVKVKFIASGEEKEVDTSKIRRVPRYGKGVHFLYDDNGKSGPGCVVEKDAPKELLDMLARAEREKKL
31.28%   Other Sequence Less than 1% Total Reads
```

**Insufficient Confidence to Predict Complex Structure (pLDDT, pTM)**

pLDDT = 46.8
pTM = 0.374
ipTM = 0.165

**Insufficient Confidence to Predict Complex Structure (pLDDT, pTM)**

pLDDT = 44.9
pTM = 0.347
ipTM = 0.159

**Insufficient Confidence to Predict Complex Structure (pLDDT, pTM)**

pLDDT = 46.8
pTM = 0.372
ipTM = 0.143

**Insufficient Confidence to Predict Complex Structure (pLDDT, pTM)**

pLDDT = 49.0
pTM = 0.369
ipTM = 0.161

| Target | BAX Peptide |
|---|---|
| Scaffold | Affitin |
| Total HTPS Reads | 8739 |
| Unique HTPS Reads | 2062 |
| Top Variant % of Total Reads | 19 |
| Final Phage Titer (PFU/mL) | 4.07E+08 |

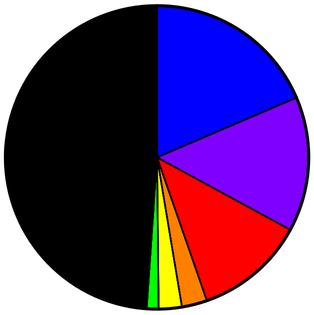

```
Library: GSVKVKF???GEEKEVDTSKI??V?R?GK?V?F?YDDNGK?G?G?V?EKDAPKELLDMLARAEREKKL
18.56%   GSVKVKF***GEEKEVDTSKIRSVWRGGKWVDFSYDDNGKQGYGYVLEKDAPKELLDMLARAEREKKL
14.36%   GSVKVKFHVNGEEKRS*YF*NSSRSPLW*RRSLSL***W*IRSWLRCEKDAPKELLDMLARAEREKKL
11.75%   GSVKVKFADLGEEKEVDTSKISHVVRSGKPVAFTYDDNGKPGHGSVHEKDAPKELLDMLARAEREKKL
2.70%    GSVKVKFYGIGEEKEVDTSKIANVNRHGKPVPFGYDDNGKSGLGHVSEKDAPKELLDMLARAEREKKL
2.46%    GSVKVKFTDHGEEKEVDTSKITVVHRNGKHVIFCYDDNGKSGSGSVLEKDAPKELLDMLARAEREKKL
1.26%    GSVKVKFHGTGEEKEVDTSKISVVIRAGKTVYFSYDDNGKRGTGRVSEKDAPKELLDMLARAEREKKL
48.91%   Other Sequence Less than 1% Total Reads
```

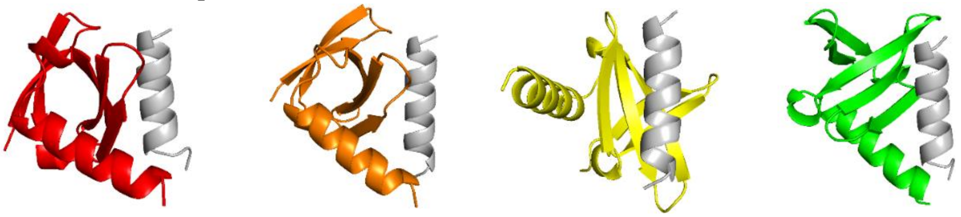

| pLDDT = 69.9 | pLDDT = 64.8 | pLDDT = 81.2 | pLDDT = 74.2 |
|---|---|---|---|
| pTM = 0.601 | pTM = 0.511 | pTM = 0.727 | pTM = 0.631 |
| ipTM = 0.172 | ipTM = 0.153 | ipTM = 0.315 | ipTM = 0.220 |

| Target | MAX |
|---|---|
| Scaffold | Affitin |
| Total HTPS Reads | 11205 |
| Unique HTPS Reads | 356 |
| Top Variant % of Total Reads | 91 |
| Final Phage Titer (PFU/mL) | 1.00E+09 |

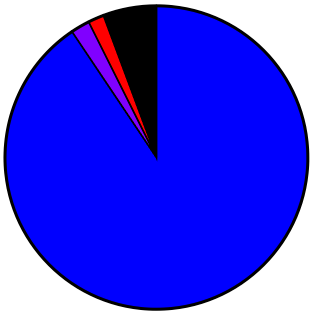

```
Library: GSVKVKF???GEEKEVDTSKI??V?R?GK?V?F?YDDNGK?G?G?V?EKDAPKELLDMLARAEREKKL
 90.58%  GSVKVKFSSAGEEKEVDTSKIFVVRRLGKYVGFTYDDNGKAGLGLVCEKDAPKELLDMLARAEREKKL
 2.03%   GSVKVKFLHSGEEKEVDTSKIPYVHRNGKHVIFLYDDNGKIGYGSVTEKDAPKELLDMLARAEREKKL
 1.70%   GSVKVKF***GEEKEVDTSKIRSVWRGGKWVDFSYDDNGKQGYGYVLEKDAPKELLDMLARAEREKKL
 5.69%   Other Sequence Less than 1% Total Reads
```

**Insufficient Confidence to Predict Complex Structure (pTM)**

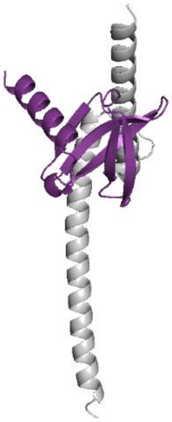

pLDDT = 83.3
pTM = 0.478
ipTM = 0.142

pLDDT = 84.9
pTM = 0.517
ipTM = 0.255

| Target | Keap1 |
|---|---|
| Scaffold | Affitin |
| Total HTPS Reads | 11196 |
| Unique HTPS Reads | 436 |
| Top Variant % of Total Reads | 80 |
| Final Phage Titer (PFU/mL) | 3.98E+08 |

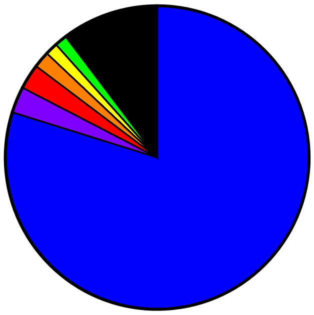

```
Library: GSVKVKF???GEEKEVDTSKI??V?R?GK?V?F?YDDNGK?G?G?V?EKDAPKELLDMLARAEREKKL
79.82%   GSVKVKFLHSGEEKEVDTSKIPYVHRNGKHVIFLYDDNGKIGYGSVTEKDAPKELLDMLARAEREKKL
2.80%    GSVKVKFRPRGEEKEVDTSKIPYVHRNGKHVIFLYDDNGKIGYGSVTEKDAPKELLDMLARAEREKKL
2.68%    GSVKVKFSSAGEEKEVDTSKIFVVRRLGKYVGFTYDDNGKAGLGLVCEKDAPKELLDMLARAEREKKL
1.75%    GSVKVKFRPRGEEKEVDTSKIPFVHRSGKNVIFLYDDNGKVGYGSVAEKDAPKELLDMLARAEREKKL
1.33%    GSVKVKFLHSGEEKEVDTSKIPYVHRNGKHVIFLYDDNGKVGYGSVAEKDAPKELLDMLARAEREKKL
1.31%    GSVKVKF***GEEKEVDTSKIRSVWRGGKWVDFSYDDNGKQGYGYVLEKDAPKELLDMLARAEREKKL
10.31%   Other Sequence Less than 1% Total Reads
```

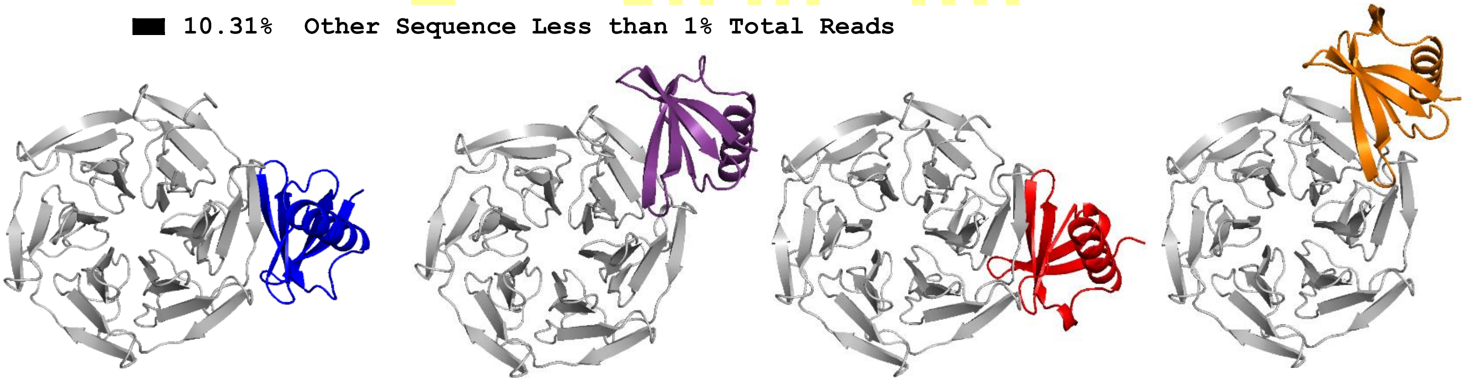

pLDDT = 87.1
pTM = 0.789
ipTM = 0.184

pLDDT = 90.2
pTM = 0.832
ipTM = 0.376

pLDDT = 87.2
pTM = 0.785
ipTM = 0.157

pLDDT = 90.9
pTM = 0.851
ipTM = 0.470

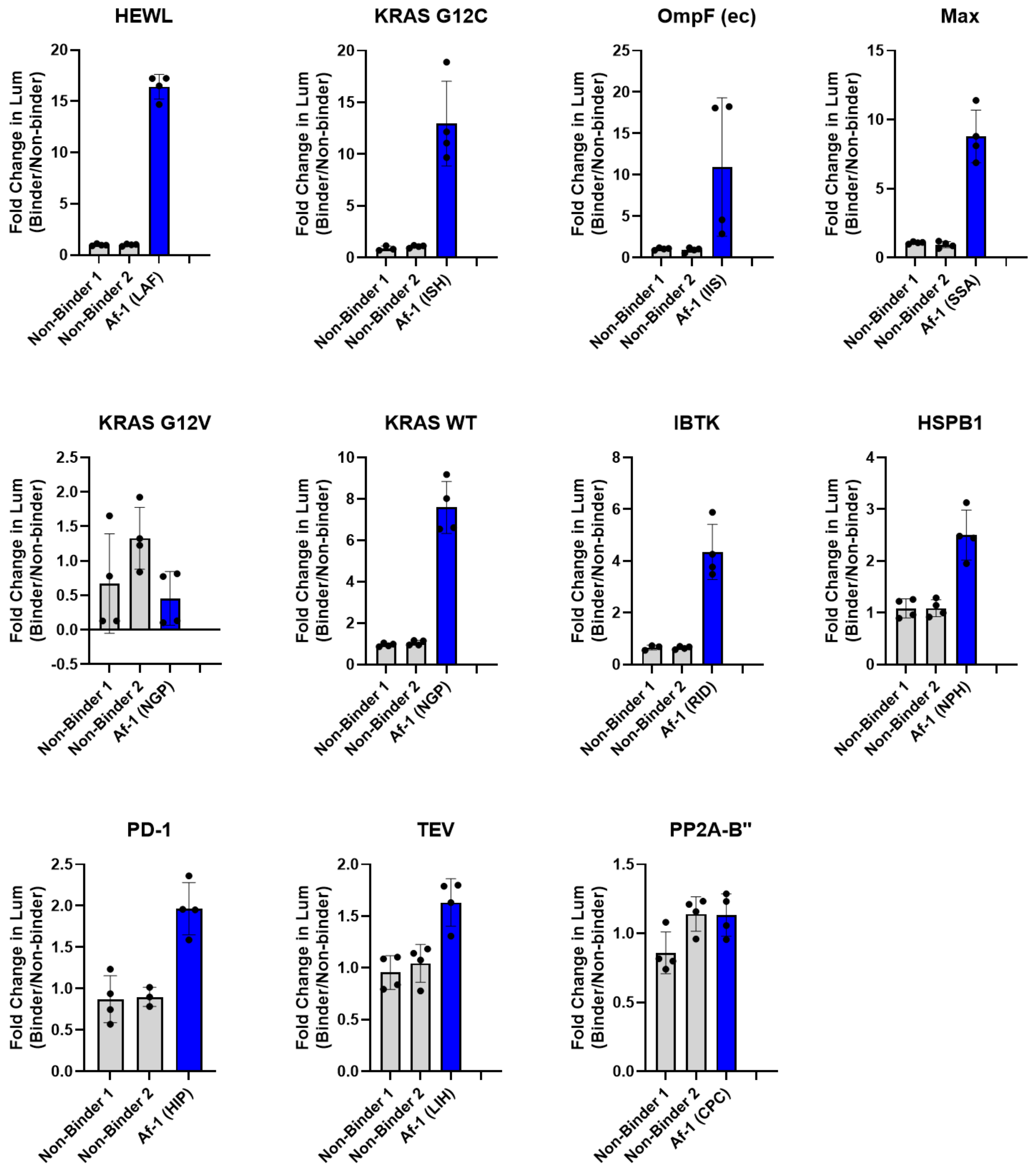


**Supplementary Fig. 23**: ***E. coli* luciferase validation assay for variants isolated from 95-panel target PANCS**.

Scaffolds from each hit were subcloned into Lux-N expression plasmids and three clones were sequenced prior to testing. If the top variant was not obtained, a second attempt was made; after which, the most dominant variant obtained was tested. Affibody hits are shown in red and affitin hits are shown in blue. Fold-change in luciferase signal for select variants from the *de novo* screens over two non-binding protein variants (Non-Binder 1 is an Affibody evolved to bind PD-L1[6] and Non-binder 2 is an Affitin evolved to bind SasA[5]; for SasA, a third Non-Binder variant was used ((Monobody (hSUMO1)[3]). Each condition has four independent data points and error bars indicate SD.

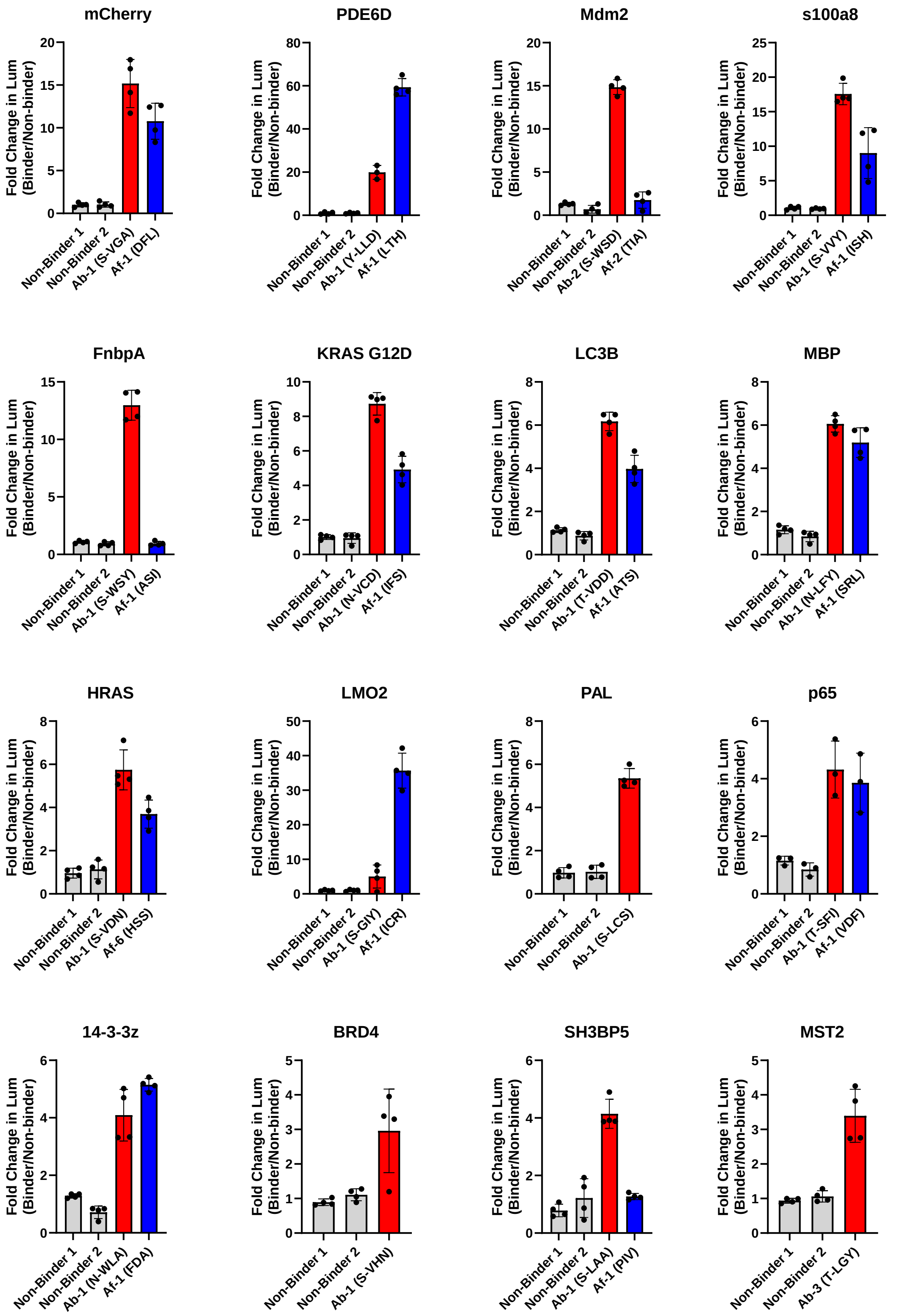
mCherry
Fold Change in Lum (Binder/Non-binder)
Non-Binder 1
Non-Binder 2
Ab-1 (S-VGA)
Af-1 (DFL)
PDE6D
Ab-1 (Y-LLD)
Af-1 (LTH)
Mdm2
Ab-2 (S-WSD)
Af-2 (TIA)
s100a8
Ab-1 (S-VVY)
Af-1 (ISH)
FnbpA
Ab-1 (S-WSY)
Af-1 (ASI)
KRAS G12D
Ab-1 (N-VCD)
Af-1 (IFS)
LC3B
Ab-1 (T-VDD)
Af-1 (ATS)
MBP
Ab-1 (N-LFY)
Af-1 (SRL)
HRAS
Ab-1 (S-VDN)
Af-6 (HSS)
LMO2
Ab-1 (S-GIY)
Af-1 (ICR)
PAL
Ab-1 (S-LCS)
p65
Ab-1 (T-SFI)
Af-1 (VDF)
14-3-3z
Ab-1 (N-WLA)
Af-1 (FDA)
BRD4
Ab-1 (S-VHN)
SH3BP5
Ab-1 (S-LAA)
Af-1 (PIV)
MST2
Ab-3 (T-LGY)

***Supplementary Fig. 23*** *(continued)*

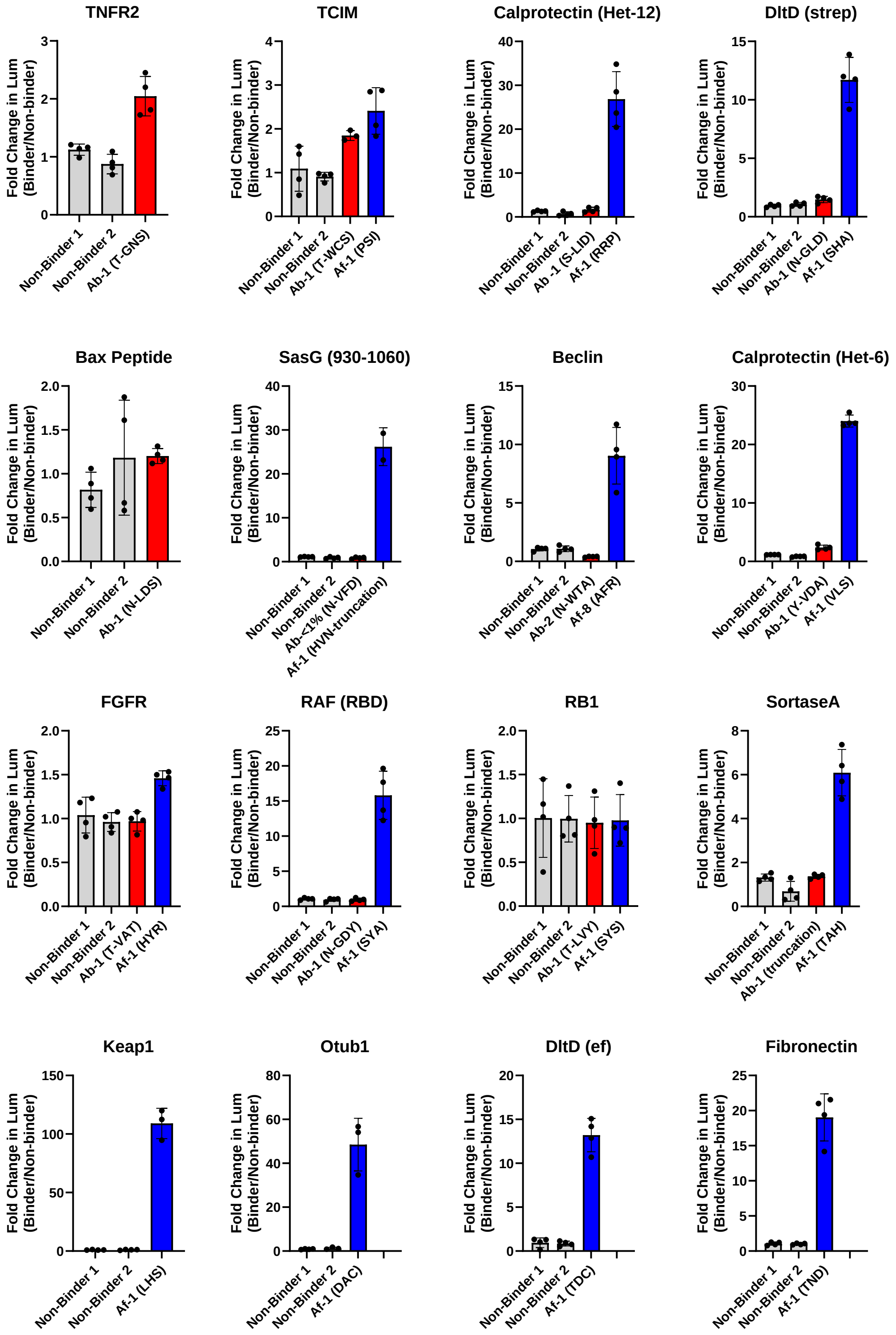

***Supplementary Fig. 23*** *(continued)*

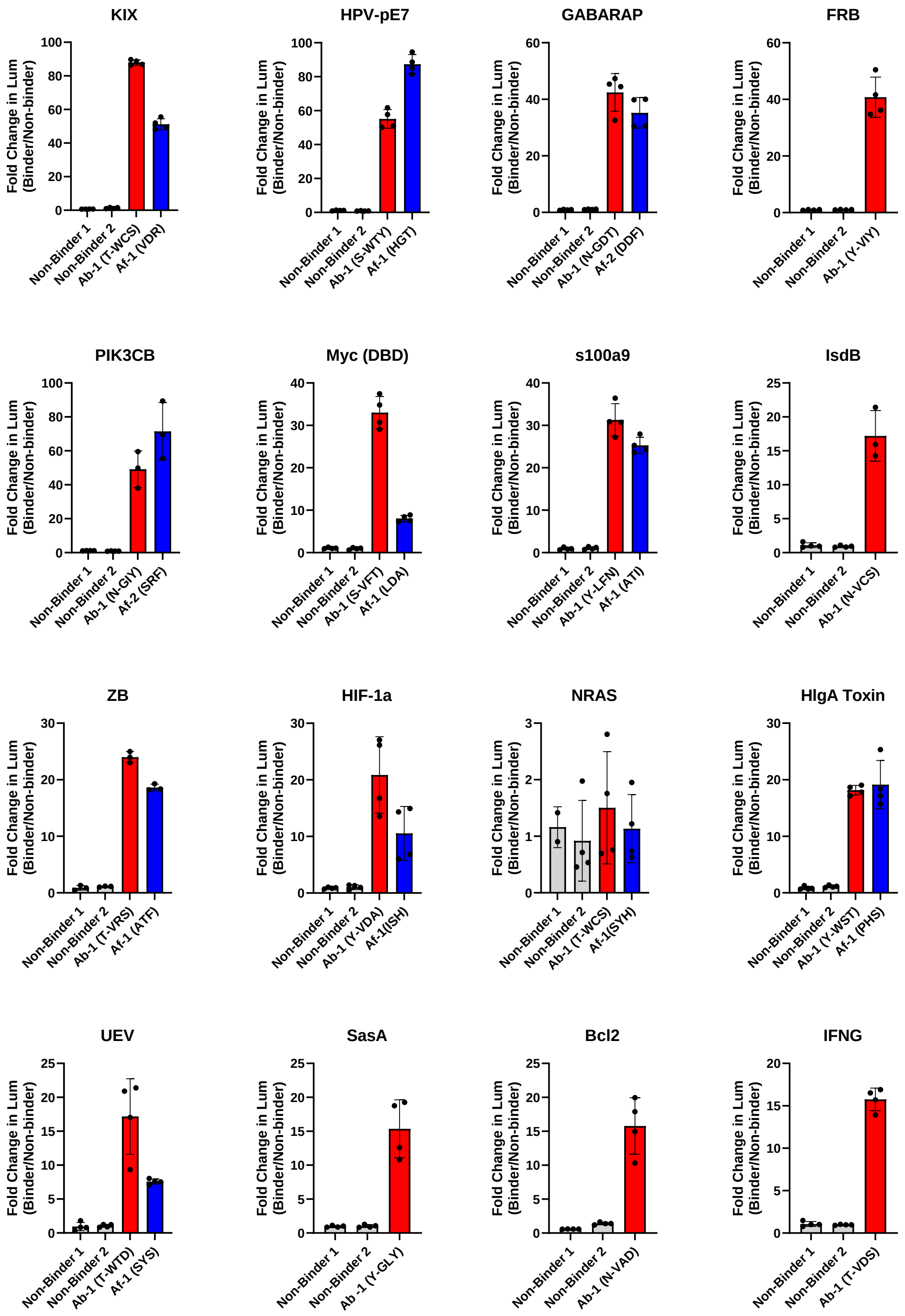

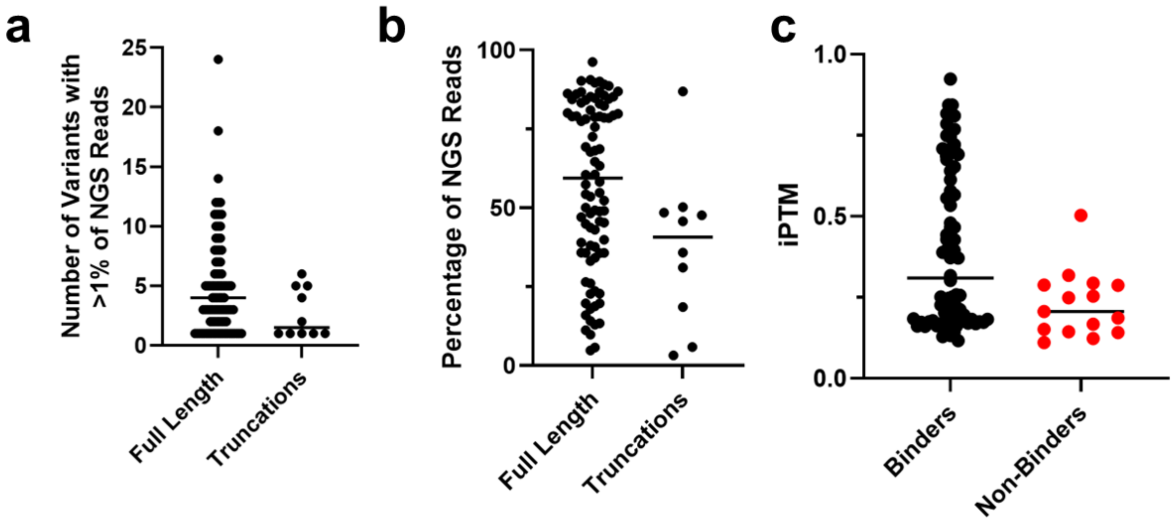


**Supplementary Fig. 24**: Summary of the 100 preliminary hit NGS results.

**a)** Number of variants with >1% of reads within a hit, grouped according to whether top variant was a full-length scaffold or a truncation. **b)** Percentage of reads of the most dominant variant within a hit, grouped according to whether the top variant was a full-length scaffold or a truncation. **c)** AlphaFold2 Multimers prediction iPTM for the top variant in all hits (excluding top hits that were truncated) shown as black for binders and red for non-binders as assessed by binder luciferase assay (**Supplementary Fig. 23**).

**Supplementary Fig. 25**: NGS data for select PANCS wells just below the hit criteria threshold.

| Target | KRAS WT |
|---|---|
| Scaffold | Affibody |
| Total HTPS Reads | 1365 |
| Unique HTPS Reads | 957 |
| Top Variant % of Total Reads | 3 |
| Final Phage Titer (PFU/mL) | 4.27E+05 |

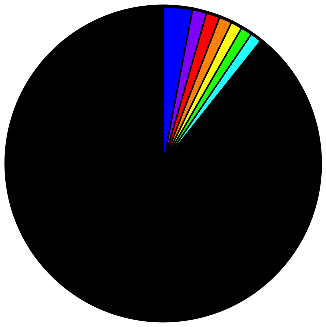

Library: VDNKF?KE???A???I??LPNLN??Q??AF??SL??DPSQSANLLAEAKKLNDAQAPK

3.00% VDNKFSKEVVYAVYQITGLPNLNLVQKFAFVTSLCDDPSQSANLLAEAKKLNDAQAPK

1.47% VDNKFNKEVADANSEIRYLPNLNLGQRLAFLGSLYNDPSQSANLLAEAKKLNDAQAPK

1.39% VDNKFNKEGDTAVLEIGHLPNLNLHQQYAFLHSLYNDPSQSANLLAEAKKLNDAQAPK

1.32% VDNKFTKEWCSAFCEICLLPNLNLSQMHAFVSSLLNDPSQSANLLAEAKKLNDAQAPK

1.17% VDNKFTKEVATAVCQICCLPNLNVDQMNAFISSLFDDPSQSANLLAEAKKLNDAQAPK

1.17% VDNKFTKESFI-E--IVNLPNLNLRQIDAFVSSLFDDPSQSANLLAEAKKLNDAQAPK

1.17% VDNKFYKEWSTAICEIVGLPNLNLSQQHAFIFSLLDDPSQSANLLAEAKKLNDAQAPK

89.30% Other Sequence Less than 1% Total Reads

| Target | FOXP3 |
|---|---|
| Scaffold | Affibody |
| Total HTPS Reads | 1512 |
| Unique HTPS Reads | 1062 |
| Top Variant % of Total Reads | 2 |
| Final Phage Titer (PFU/mL) | 2.88E+05 |

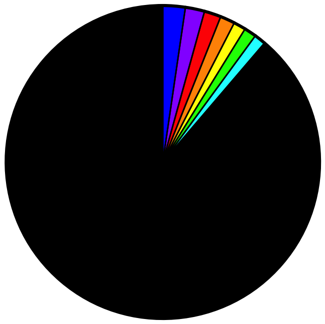

Library: VDNKF?KE???A???I??LPNLN??Q??AF??SL??DPSQSANLLAEAKKLNDAQAPK

2.31% VDNKFNKEVCDAYAQIGNLPNLNVYQIVAFIRSLDNDPSQSANLLAEAKKLNDAQAPK

1.98% VDNKFNKEVADANSEIRYLPNLNLGQRLAFLGSLYNDPSQSANLLAEAKKLNDAQAPK

1.72% VDNKFTKE--S-FIEIVNLPNLNLRQIDAFVSSLFDDPSQSANLLAEAKKLNDAQAPK

1.52% VDNKFNKEGDTAVLEIGHLPNLNLHQQYAFLHSLYNDPSQSANLLAEAKKLNDAQAPK

1.26% VDNKFTKEVATAVCQICCLPNLNVDQMNAFISSLFDDPSQSANLLAEAKKLNDAQAPK

1.26% VDNKFSKEVVYAVYQITGLPNLNLVQKFAFVTSLCDDPSQSANLLAEAKKLNDAQAPK

1.19% VDNKFTKEWCSAFCEICLLPNLNLSQMHAFVSSLLNDPSQSANLLAEAKKLNDAQAPK

88.76% Other Sequence Less than 1% Total Reads

| Target | Alpha amylase |
|---|---|
| Scaffold | Affibody |
| Total HTPS Reads | 1311 |
| Unique HTPS Reads | 799 |
| Top Variant % of Total Reads | 14 |
| Final Phage Titer (PFU/mL) | 2.14E+05 |

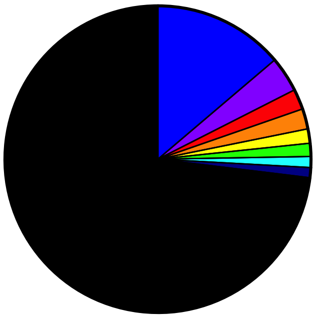

Library: VDNKF?KE???A???I??LPNLN??Q??AF??SL??DPSQSANLLAEAKKLNDAQAPK

13.81% VDNKFYKEVCDAFDEICDLPNLNLLQLDAFLLSLVDDPSQSANLLAEAKKLNDAQAPK

3.74% VDNKFSKEVFTAFDQIGGLPNLNLCQKDAFICSLVDDPSQSANLLAEAKKLNDAQAPK

2.14% VDNKFYKEVDAAASEICSLPNLNRNQRSAFICSLYNDPSQSANLLAEAKKLNDAQAPK

2.14% VDNKFSKEGIYAISQIVRLPNLNGFQEVAFLSSLPNDPSQSANLLAEAKKLNDAQAPK

1.53% VDNKFTKEWCSAFCEICLLPNLNLSQMHAFVSSLLNDPSQSANLLAEAKKLNDAQAPK

1.37% VDNKFYKEVIYAAAEIFGLPNLNVGQRSAFVCSLYDDPSQSANLLAEAKKLNDAQAPK

1.14% VDNKFNKEVCSAFNEIGNLPNLNLGQKIAFVRSLYNDPSQSANLLAEAKKLNDAQAPK

1.07% VDNKFNKELDSAYNQICHLPNLNANQRSAFLSSLYDDPSQSANLLAEAKKLNDAQAPK

73.07% Other Sequence Less than 1% Total Reads

| Target | No Fusion |
|---|---|
| Scaffold | Affitin |
| Total HTPS Reads | 2202 |
| Unique HTPS Reads | 964 |
| Top Variant % of Total Reads | 35 |
| Final Phage Titer (PFU/mL) | 1.55E+06 |

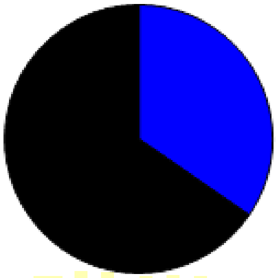

Library: GSVKVKF???GEEKEVDTSKI??V?R?GK?V?F?YDDNGK?G?G?V?EKDAPKELLDMLARAEREKKL

34.60% GSVKVKF***GEEKEVDTSKIRSVWRGGKWVDFSYDDNGKQGYGYVLEKDAPKELLDMLARAEREKKL

65.40% Other Sequence less than 1%

| | |
|---|---|
| Target | GFP |
| Scaffold | Affitin |
| Total HTPS Reads | 3145 |
| Unique HTPS Reads | 1061 |
| Top Variant % of Total Reads | 42 |
| Final Phage Titer (PFU/mL) | 2.04E+06 |

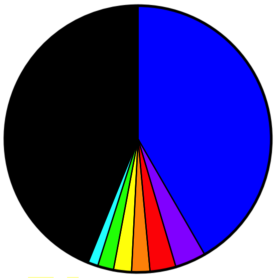

Library: GSVKVKF???GEEKEVDTSKI??V?R?GK?V?F?YDDNGK?G?G?V?EKDAPKELLDMLARAEREKKL

41.65% GSVKVKF***GEEKEVDTSKIRSVWRGGKWVDFSYDDNGKQGYGYVLEKDAPKELLDMLARAEREKKL

3.75% GSVKVKFATFGEEKEVDTSKIHDVDRYGKTVHFIYDDNGKCGLGYVNEKDAPKELLDMLARAEREKKL

3.15% GSVKVKFHYRGEEKEVDTSKIVHVHRDGKFVLFVYDDNGKNGVGGVYEKDAPKELLDMLARAEREKKL

2.23% GSVKVKFSYHGEEKEVDTSKINNVNRHGKHVLFDYDDNGKIGSGLVFEKDAPKELLDMLARAEREKKL

2.19% GSVKVKFIFSGEEKEVDTSKIHAVFRNGKYVTFFYDDNGKHGFGSVLEKDAPKELLDMLARAEREKKL

1.94% GSVKVKFVDFGEEKEVDTSKIRLVIRFGKSVHFIYDDNGKDGYGIVNEKDAPKELLDMLARAEREKKL

1.21% GSVKVKFRRPGEEKEVDTSKIGYVHRVGKFVAFFYDDNGKIGSGTVSEKDAPKELLDMLARAEREKKL

43.88% Other Sequence Less than 1% Total Reads

| | |
|---|---|
| Target | G3BP |
| Scaffold | Affitin |
| Total HTPS Reads | 1712 |
| Unique HTPS Reads | 760 |
| Top Variant % of Total Reads | 31 |
| Final Phage Titer (PFU/mL) | 3.24E+06 |

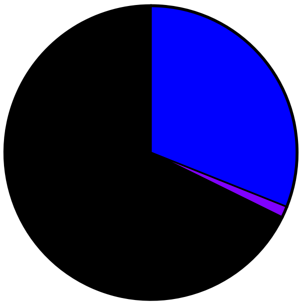

Library: GSVKVKF???GEEKEVDTSKI??V?R?GK?V?F?YDDNGK?G?G?V?EKDAPKELLDMLARAEREKKL

31.02% GSVKVKF***GEEKEVDTSKIRSVWRGGKWVDFSYDDNGKQGYGYVLEKDAPKELLDMLARAEREKKL

1.23% GSVKVKFPLLGEEKEVDTSKITCVHRTGKDVAFIMMIMVKTVLVAFPEKDAPKELLDMLARAEREKKL

67.76% Other Sequence less than 1%

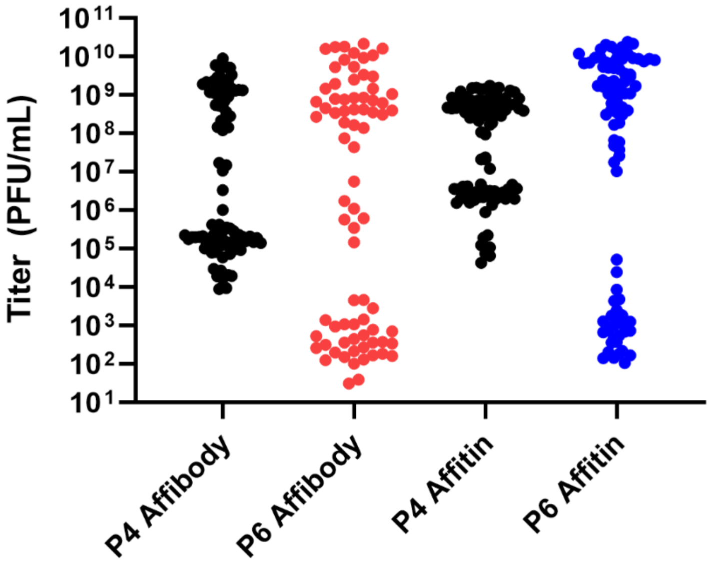


**Supplementary Fig. 26. Further passaging increases signal-to-noise of hits and deceases false positives.**

Additional passaging of passage 4 (P4) using a 5% transfer rate for 2 passages (P6). Each data point indicates the titer (measured by qPCR) of a single well in the deep 96-well PANCS plate.

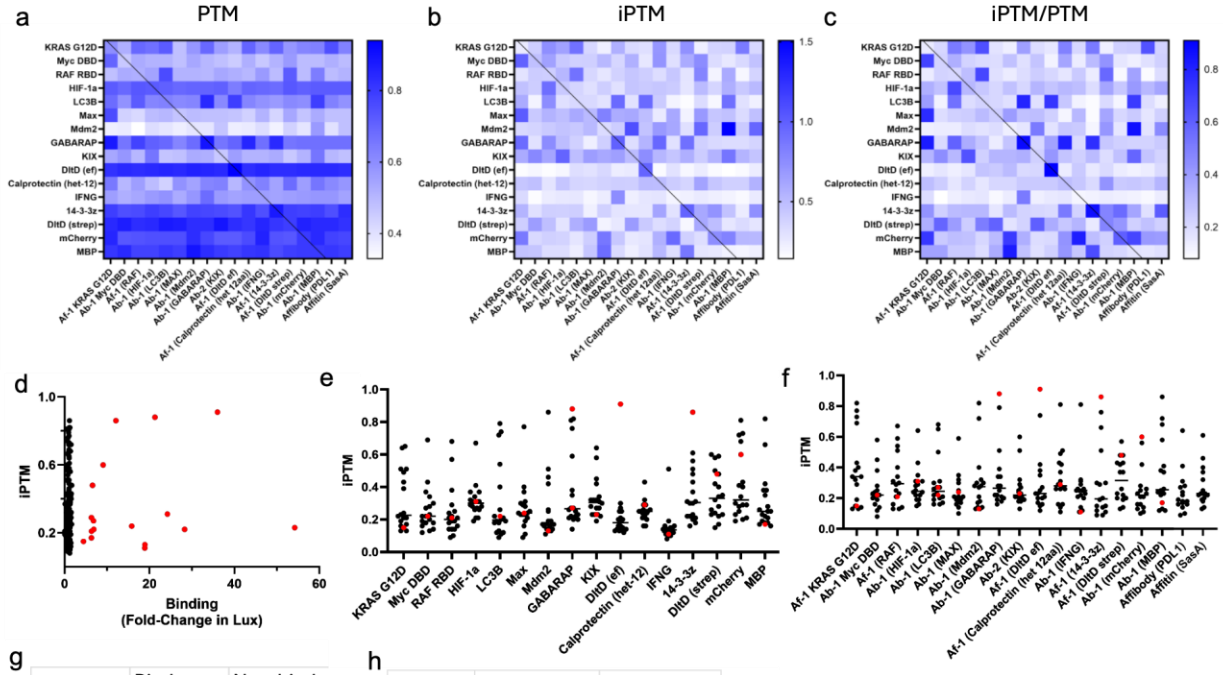


g

| | Binders | Non-binde |
|---|---|---|
| Average | 0.369412 | 0.291661 |
| STDEV | 0.273575 | 0.167486 |
| Max | 0.91 | 0.86 |
| Min | 0.11 | 0.08 |

h

| | False negatives | False positives |
|---|---|---|
| 0.5 cut-off | 76.5 | 11.5 |
| 0.6 cut-off | 76.5 | 7.6 |
| 0.7 cut-off | 82.4 | 3.8 |
| 0.8 cut-off | 82.4 | 1.7 |
| 0.9 cut -off | 94.1 | 0.0 |

**Supplementary Fig. 27. AlphaFold 3 predictions for every target-binder pairing in our specificity of binding luciferase dataset (Fig. 4c).**

**a)** pTM, **b)** iPTM, and **c)** iPTM/PTM heatmaps. In A-C, the diagonal indicating position of binder-target pairing is shown a black line. **d)** Correlation of iPTM and binding. iPTM separated by **e)** target and **f)** by binder. In D-F, black dotes indicate non-binders and red dots indicate binders (>3 fold change in luciferase signal). **g)** Statistical categorization of iPTM of predictions for variant-target pairings for binder and non-binder pairs. **h)** Percentage of false negatives and false positives within this data set if an iPTM threshold is set as a cut-off for filtering binders from nonbinders.

**Supplementary Table 6. Library size estimation.**

Cloning of large ~$10^{9-10}$ variant affibody and affitin libraries (referred to as $10^{10}$ libraries). Estimated number of variants is based on the average of 40- and 80-minute timepoints. Titers determined using activity independent plaque assays. # indicates different transformations that were kept separate.

| Library | # | 40 minute Total Titer | 80 minute Total Titer | 120 minute Total Titer | Estimated Number of Variants | Amplified Titer (PFU/mL) |
|---|---|---|---|---|---|---|
| Affibody | 1 | 3.75E+08 | 1.90E+09 | 1.30E+09 | 1.14E+09 | 2.50E+11 |
| Affibody | 2 | 2.05E+08 | 2.30E+09 | 2.10E+09 | 1.25E+09 | 1.95E+11 |
| Affibody | 3 | 4.95E+08 | 3.20E+09 | 2.50E+09 | 1.85E+09 | 1.15E+11 |
| Affibody | 4 | 2.00E+08 | 3.25E+09 | 3.50E+09 | 1.73E+09 | 7.75E+10 |
| Affibody | 5 | 3.10E+08 | 2.80E+09 | 1.80E+09 | 1.56E+09 | 1.18E+11 |
| Affibody | 6 | 2.20E+08 | 2.90E+09 | 1.65E+09 | 1.56E+09 | 1.43E+11 |
| Affibody | 7 | 1.85E+08 | 2.60E+09 | 1.10E+10 | 1.39E+09 | 1.78E+11 |
| Affibody | 8 | 1.70E+08 | 2.15E+09 | 1.90E+09 | 1.16E+09 | 1.20E+11 |
| | | | | **Total** | **1.16E+10** | |
| Affitin | 1 | 3.50E+07 | 5.50E+08 | 5.50E+08 | 2.93E+08 | 1.58E+11 |
| Affitin | 2 | 7.50E+07 | 6.50E+08 | 1.20E+09 | 3.63E+08 | 1.05E+11 |
| Affitin | 3 | 3.50E+07 | 6.00E+08 | 1.15E+09 | 3.18E+08 | 2.23E+11 |
| Affitin | 4 | 2.50E+07 | 4.50E+08 | 1.20E+09 | 2.38E+08 | 6.75E+10 |
| Affitin | 5 | 3.50E+07 | 6.00E+08 | 1.30E+09 | 3.18E+08 | 1.78E+11 |
| Affitin | 6 | 2.00E+07 | 6.50E+08 | 1.15E+09 | 3.35E+08 | 1.38E+11 |
| Affitin | 7 | 5.00E+07 | 5.50E+08 | 1.40E+09 | 3.00E+08 | 1.20E+11 |
| Affitin | 8 | 2.00E+08 | 5.00E+08 | 1.45E+09 | 3.50E+08 | 8.25E+10 |
| | | | | **Total** | **2.51E+09** | |

**Supplementary Table 7. Endpoint titers of large library selections.**

Passage 6 endpoint phage titers from PANCS with large, $10^{10}$, libraries; assessed using activity independent plaque assay. Bold indicates this library-target pair was successful at $10^8$ scale. Green shading indicates confirmed binders were enriched.

| Target | Titer (PFU/mL) for Affibody Library | Titer (PFU/mL) for Affitin Library | Target | Titer (PFU/mL) for Affibody Library | Titer (PFU/mL) for Affitin Library |
|---|---|---|---|---|---|
| KRAS G12D | **1.42E+11** | **2.38E+09** | TRIM21 | 2.20E+08 | <1.0E+05 |
| HRAS | **1.65E+09** | **4.44E+11** | ALKBH5 | 5.30E+06 | <1.0E+05 |
| RAF | 2.43E+10 | **1.45E+10** | PINK | <1.0E+05 | <1.0E+05 |
| IFNG | **1.93E+10** | 1.63E+10 | PIK3CA | 5.00E+07 | <1.0E+05 |
| VHL | 1.03E+10 | 2.75E+09 | BAX | <1.0E+05 | 3.00E+07 |
| PCNA | 9.25E+09 | 6.00E+09 | NIX | 4.00E+07 | <1.0E+05 |

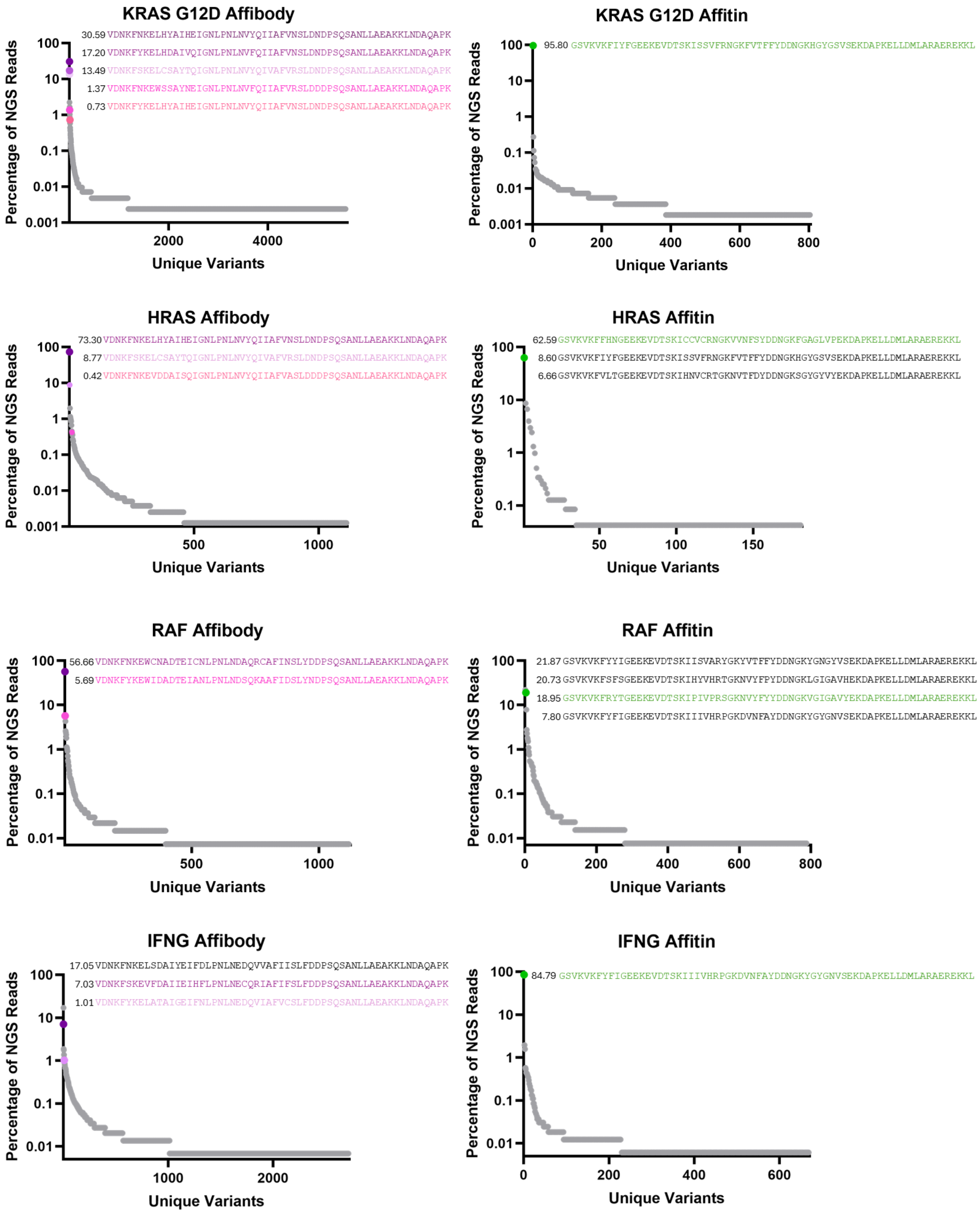
KRAS G12D Affibody
30.59 VDNKFNKELHYAIHEIGNLPNLNVYQIIAFVNSLDNDPSQSANLLAEAKKLNDAQAPK
17.20 VDNKFYKELHDAIVQIGNLPNLNVFQIIAFVNSLDNDPSQSANLLAEAKKLNDAQAPK
13.49 VDNKFSKELCSAYTQIGNLPNLNVYQIVAFVRSLDNDPSQSANLLAEAKKLNDAQAPK
1.37 VDNKFNKEWSSAYNEIGNLPNLNVFQIIAFVRSLDDDPSQSANLLAEAKKLNDAQAPK
0.73 VDNKFYKELHYAIHEIGNLPNLNVYQIIAFVNSLDNDPSQSANLLAEAKKLNDAQAPK
Percentage of NGS Reads
100
10
1
0.1
0.01
0.001
2000
4000
Unique Variants
KRAS G12D Affitin
95.80 GSVKVKFIYFGEEKEVDTSKISSVFRNGKFVTFFYDDNGKHGYGSVSEKDAPKELLDMLARAEREKKL
Percentage of NGS Reads
100
10
1
0.1
0.01
0.001
0
200
400
600
800
Unique Variants
HRAS Affibody
73.30 VDNKFNKELHYAIHEIGNLPNLNVYQIIAFVNSLDNDPSQSANLLAEAKKLNDAQAPK
8.77 VDNKFSKELCSAYTQIGNLPNLNVYQIVAFVRSLDNDPSQSANLLAEAKKLNDAQAPK
0.42 VDNKFNKEVDDAISQIGNLPNLNVYQIIAFVASLDDDPSQSANLLAEAKKLNDAQAPK
Percentage of NGS Reads
100
10
1
0.1
0.01
0.001
500
1000
Unique Variants
HRAS Affitin
62.59 GSVKVKFFHNGEEKEVDTSKICCVCRNGKVVNFSYDDNGKFGAGLVPEKDAPKELLDMLARAEREKKL
8.60 GSVKVKFIYFGEEKEVDTSKISSVFRNGKFVTFFYDDNGKHGYGSVSEKDAPKELLDMLARAEREKKL
6.66 GSVKVKFVLTGEEKEVDTSKIHNVCRTGKNVTFDYDDNGKSGYGYVYEKDAPKELLDMLARAEREKKL
Percentage of NGS Reads
100
10
1
0.1
50
100
150
Unique Variants
RAF Affibody
56.66 VDNKFNKEWCNADTEICNLPNLNDAQRCAFINSLYDDPSQSANLLAEAKKLNDAQAPK
5.69 VDNKFYKEWIDADTEIANLPNLNDSQKAAFIDSLYNDPSQSANLLAEAKKLNDAQAPK
Percentage of NGS Reads
100
10
1
0.1
0.01
500
1000
Unique Variants
RAF Affitin
21.87 GSVKVKFYYIGEEKEVDTSKIISVARYGKYVTFFYDDNGKYGNGYVSEKDAPKELLDMLARAEREKKL
20.73 GSVKVKFSFSGEEKEVDTSKIHYVHRTGKNVYFPYDDNGKLGIGAVHEKDAPKELLDMLARAEREKKL
18.95 GSVKVKFRYTGEEKEVDTSKIPIVPRSGKNVYFYYDDNGKVGIGAVYEKDAPKELLDMLARAEREKKL
7.80 GSVKVKFYFIGEEKEVDTSKIIIVHRPGKDVNFAYDDNGKYGYGNVSEKDAPKELLDMLARAEREKKL
Percentage of NGS Reads
100
10
1
0.1
0.01
0
200
400
600
800
Unique Variants
IFNG Affibody
17.05 VDNKFNKELSDAIYEIFDLPNLNEDQVVAFIISLFDDPSQSANLLAEAKKLNDAQAPK
7.03 VDNKFSKEVFDAIIEIHFLPNLNECQRIAFIFSLFDDPSQSANLLAEAKKLNDAQAPK
1.01 VDNKFYKELATAIGEIFNLPNLNEDQVIAFVCSLFDDPSQSANLLAEAKKLNDAQAPK
Percentage of NGS Reads
100
10
1
0.1
0.01
1000
2000
Unique Variants
IFNG Affitin
84.79 GSVKVKFYFIGEEKEVDTSKIIIVHRPGKDVNFAYDDNGKYGYGNVSEKDAPKELLDMLARAEREKKL
Percentage of NGS Reads
100
10
1
0.1
0.01
0
200
400
600
Unique Variants

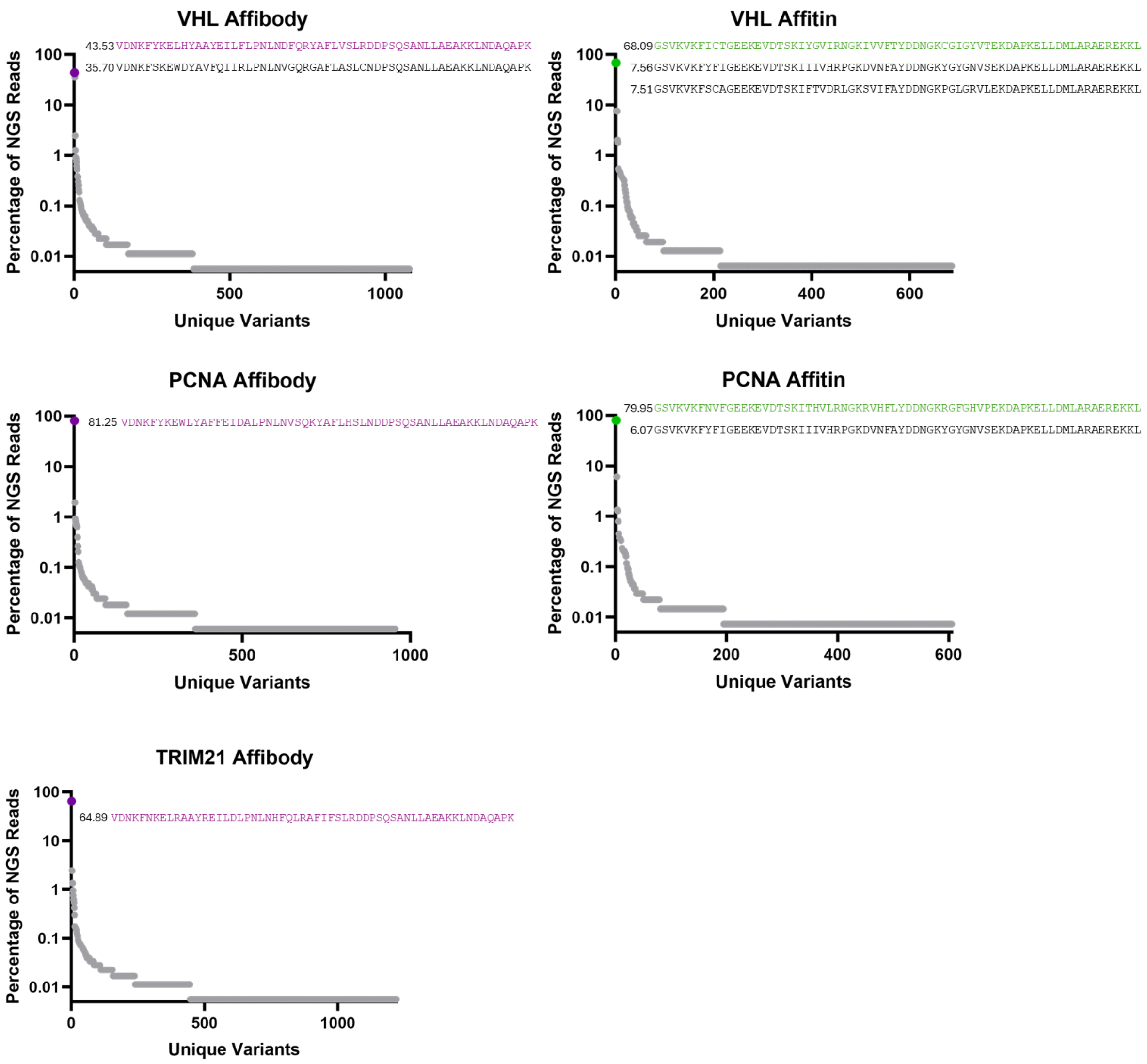


**Supplementary Fig. 28. Sequence prevalence plots for hits from the large, $10^{10}$, library PANCS.**

The percentage of NGS reads is plotted for each unique variant, and variants tested in the luminescence assay are highlighted in the color shown in **Supplementary Fig. 30**. The sequences of all variants with >5% of reads are listed next to their percentage of NGS reads (sequences below 5% included if we collected luminescence assay data for that variant).

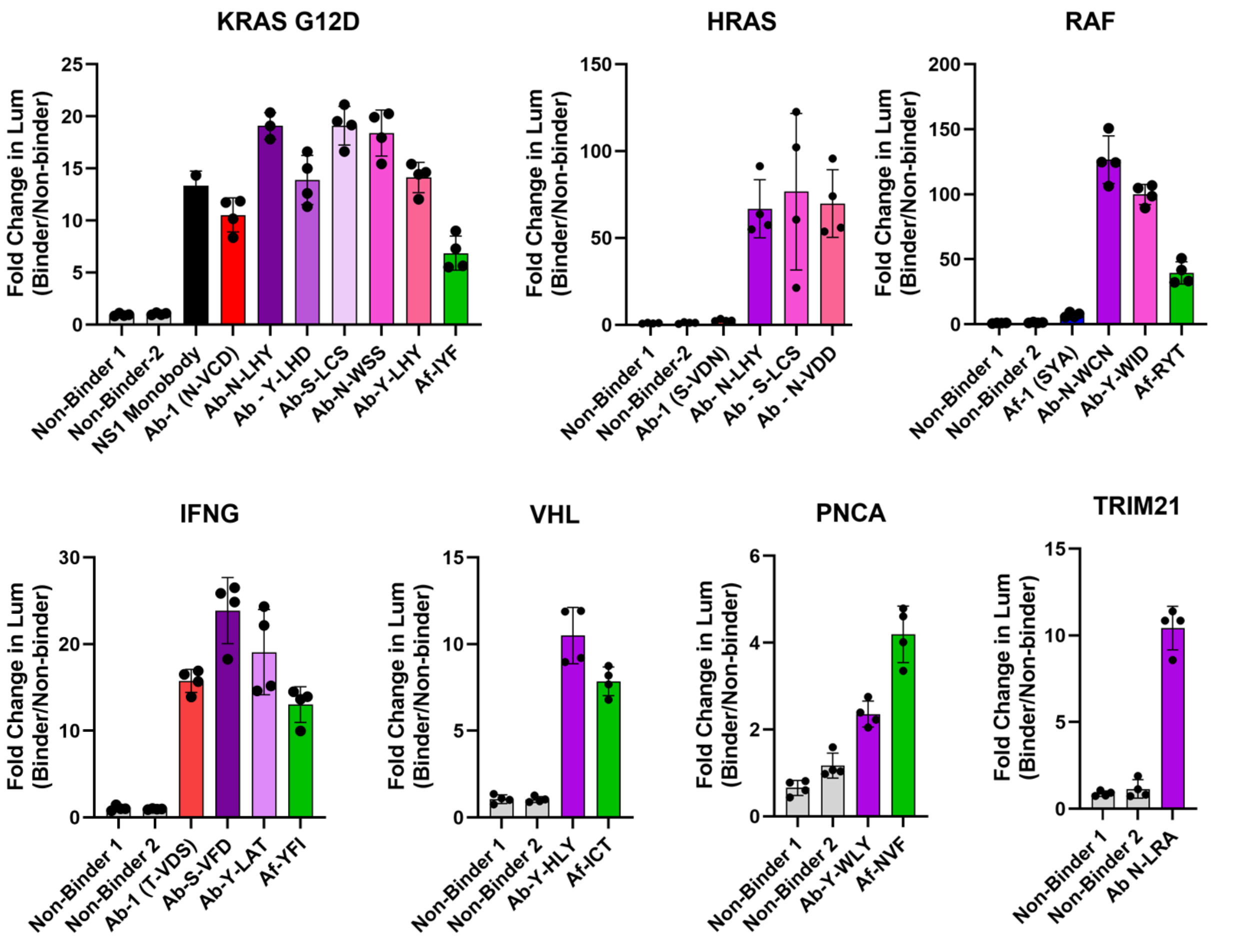


**Supplementary Fig. 29. Luciferase binding assays of variants from $10^{10}$ library PANCS.**

Original best variant from $10^8$ library PANCS hit shown in either red (affibody) or blue (affitin). New affibody hits are shown in purple, and new affitin hits are shown in green. Fold-change in luciferase signal for select variants from the *de novo* screens over two non-binding protein variants (Non-Binder 1 is an Affibody evolved to bind PD-L1[6] and Non-binder 2 is an Affitin evolved to bind SasA[5]). Each condition has four independent data points and error bars indicate SD.

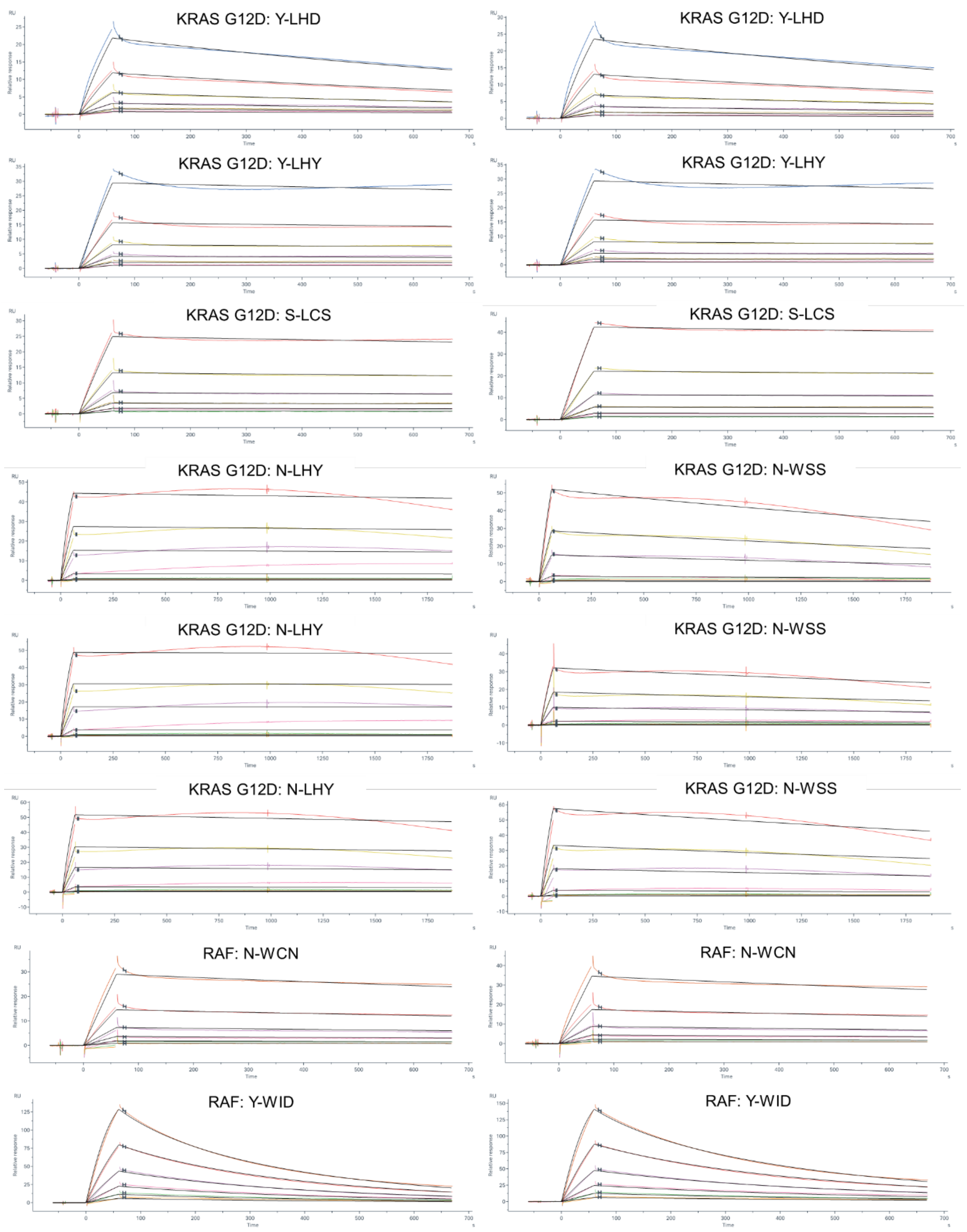


**Supplementary Fig. 30**: **Surface Plasmon Resonance (SPR) dose-response and kinetic model fits for $K_d$s shown in Fig. 5B** (**Supplementary Table 8** for details of model fits).

Each target-binder pair required a different concentration range: KRAS G12D with Y-LHD (400, 200, 100, 50, 25, 12.5 0 nM), Y-LHY (400, 200, 100, 50, 25, 12.5 0 nM), N-LHY (100, 50, 25, 5, 1, 0.2, 0 nM), S-LCS (200, 100, 50, 25, 12.5, 6.3, 0 nM), and N-WSS (100, 50, 25, 5, 1, 0.2, 0 nM); and RAF with N-WCN

(60, 30, 15, 7.5, 3.8, 1.9, 0 nM) and Y-WID (60, 30, 15, 7.5, 3.8, 1.9, 0 nM). All interactions were measured in duplicate and all data was fit to a 1:1 binding model.

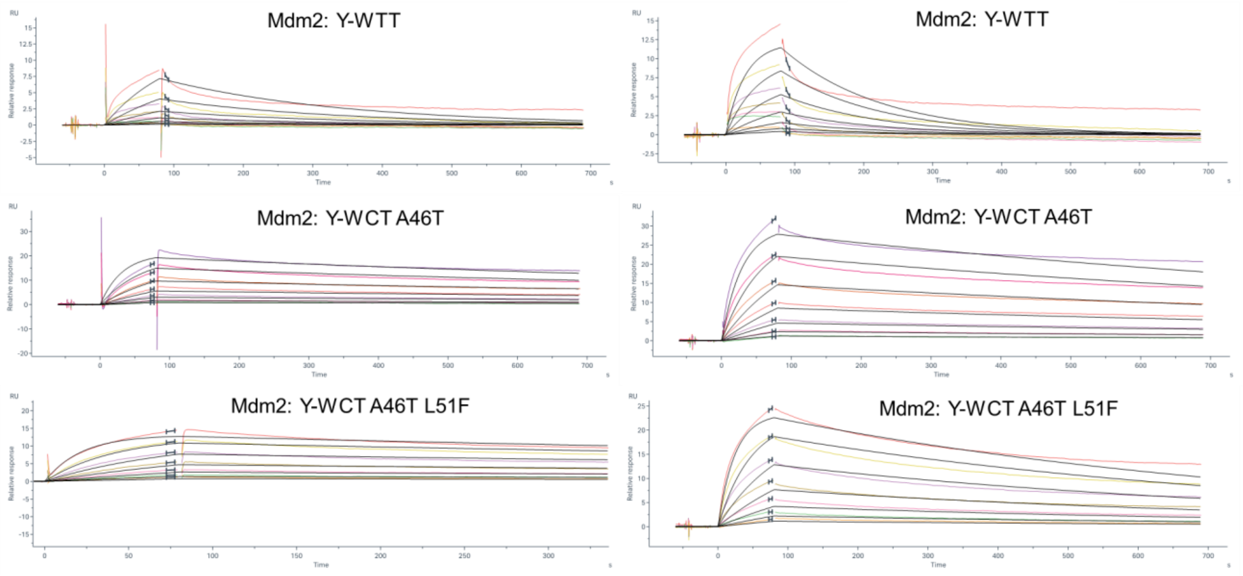


**Supplementary Fig. 31: Surface Plasmon Resonance (SPR) dose-response and kinetic model fits for $K_d$s shown in Fig. 5D** (**Supplementary Table 8** for details of model fits).

Each target-binder pair required a different concentration range: Mdm2 with Y-WTT (1000, 500, 250, 125, 62.5, 31.3, 15.6, 0 nM), Y-WCT A46T (200, 100, 50, 25, 6.3, 1.5, 0.4, 0 nM), and Y-WCT A46T L51F (1000, 500, 250, 125, 62.5, 31.3, 15.6, 0 nM). All interactions were measured in duplicate and all data was fit to a 1:1 binding model.

**Supplementary Table 8:** Summary of all SPR data model fits (**Supplementary Figs. 14, 28,** and **31**).

| Target | Binder | Model | $Chi^2$ | $k_a$ (1/Ms) | $k_d$ (1/s) | $K_d$ (nM) |
|---|---|---|---|---|---|---|
| KRAS G12D | Ab- N-VCD | 1:1 binding | 1.44E+01 | 4.76E+04 | 1.80E-02 | 378 |
| | | 1:1 binding | 1.99E+01 | 1.55E+04 | 6.25E-03 | 403 |
| | | | Average | **3.16E+04** | **1.21E-02** | **384** |
| KRAS G12D | Ab- Y-LHD | 1:1 binding | 2.19E-01 | 1.47E+04 | 8.87E-04 | 60.3 |
| | | 1:1 binding | 3.40E-01 | 1.86E+04 | 7.99E-04 | 43.0 |
| | | | 50.6Average | **1.67E+04** | **8.43E-04** | **50.6** |
| KRAS G12D | Ab- Y-LHY | 1:1 binding | 4.31E-01 | 1.14E+04 | 1.35E-04 | 11.8 |
| | | 1:1 binding | 4.17E-01 | 1.15E+04 | 1.54E-04 | 13.4 |
| | | | Average | **1.15E+04** | **1.45E-04** | **12.6** |
| KRAS G12D | Ab- N-LHY | 1:1 binding | 5.14E+00 | 1.59E+05 | 3.29E-05 | 0.207 |
| | | 1:1 binding | 5.85E+00 | 1.72E+05 | 6.89E-06 | 0.040 |
| | | 1:1 binding | 3.84E+00 | 1.16E+05 | 5.25E-05 | 0.453 |
| | | | Average | **1.49E+05** | **3.08E-05** | **0.206** |
| KRAS G12D | Ab- S-LCS | 1:1 binding | 9.98E-02 | 2.11E+04 | 1.21E-04 | 5.7 |
| | | 1:1 binding | 1.33E-01 | 8.33E+04 | 1.73E-04 | 2.1 |
| | | | Average | **5.22E+04** | **1.47E-04** | **2.8** |
| KRAS G12D | Ab- N-WSS | 1:1 binding | 1.94E+00 | 6.58E+04 | 2.37E-04 | 3.60 |
| | | 1:1 binding | 8.10E-01 | 1.05E+05 | 1.67E-04 | 1.59 |

| | | | | | | |
|---|---|---|---|---|---|---|
| | | 1:1 binding | 3.67E+00 | 1.08E+05 | 1.66E-04 | 1.54 |
| | | | Average | **9.29E+04** | **1.90E-04** | **2.04** |
| RAF | Ab- T-VPN | 1:1 binding | 1.13E+01 | 6.71E+04 | 1.79E-02 | 267 |
| | | 1:1 binding | 2.36E+01 | 5.11E+04 | 1.36E-02 | 266 |
| | | | 266.5Average | **5.91E+04** | **1.58E-02** | **267** |
| RAF | Ab- N-LFN | 1:1 binding | 4.24E+00 | 1.68E+02 | 1.52E-03 | 9048 |
| | | 1:1 binding | 1.38E+00 | 1.01E+03 | 2.08E-03 | 2059 |
| | | | 5.89E+021.80E-033056.0Average | **5.89E+02** | **1.80E-03** | **3056** |
| RAF | Ab- Y-LCS | 1:1 binding | 3.54E+00 | 2.87E+04 | 5.75E-02 | 2004 |
| | | 1:1 binding | 3.59E+00 | 2.60E+04 | 3.90E-02 | 1500 |
| | | | Average | **2.74E+04** | **4.83E-02** | **1764** |
| RAF | Ab- N-WCN | 1:1 binding | 4.29E-01 | 3.87E+03 | 3.19E-04 | 82.4 |
| | | 1:1 binding | 7.14E-01 | 1.15E+04 | 3.66E-04 | 31.8 |
| | | | Average | **7.69E+03** | **3.43E-04** | **44.6** |
| RAF | Ab- Y-WID | 1:1 binding | 1.75E+00 | 7.80E+05 | 5.86E-03 | 7.5 |
| | | 1:1 binding | 2.06E+00 | 8.99E+05 | 5.10E-03 | 5.7 |
| | | | Average | **8.40E+05** | **5.48E-03** | **6.5** |
| IFNG | Ab- S-VVD | 1:1 binding | 3.39E-01 | 6.89E+02 | 1.10E-03 | 1597 |
| | | 1:1 binding | 5.05E+00 | 1.84E+03 | 5.06E-04 | 275 |
| | | | Average | **1.26E+03** | **8.03E-04** | **635** |
| IFNG | Ab- S-VCD | 1:1 binding | 1.07E+01 | 3.12E+02 | 1.95E-03 | 6250 |
| | | 1:1 binding | 5.73E+00 | 7.67E+02 | 1.75E-03 | 2282 |
| | | | Average | **5.40E+02** | **1.85E-03** | **3429** |
| Mdm2 | Ab-S-WFY | 1:1 binding | 9.21E-01 | 1.06E+04 | 1.40E-03 | 132 |
| | | 1:1 binding | 8.51E-01 | 3.75E+03 | 1.13E-03 | 301 |
| | | | Average | **7.18E+03** | **1.27E-03** | **176** |
| Mdm2 | Ab-S-WYS | 1:1 binding | 2.07E+00 | 2.57E+04 | 6.47E-03 | 252 |
| | | 1:1 binding | 1.24E+00 | 2.20E+04 | 4.90E-03 | 223 |
| | | | Average | **2.39E+04** | **5.69E-03** | **238** |
| Mdm2 | Ab-T-WDN | 1:1 binding | 2.29E-01 | 6.44E+04 | 4.01E-02 | 623 |
| | | 1:1 binding | 2.88E+00 | 7.13E+02 | 8.77E-04 | 1230 |
| | | | Average | **3.26E+04** | **2.05E-02** | **629** |
| Mdm2 | Ab-Y-WTT | 1:1 binding | 4.13E-01 | 6.73E+03 | 3.84E-03 | 571 |
| | | 1:1 binding | 1.56E+00 | 2.99E+04 | 6.98E-03 | 233 |
| | | | Average | **1.83E+04** | **5.41E-03** | **295** |
| Mdm2 | Ab-Y-WCT | 1:1 binding | 7.16E-01 | 7.87E+04 | 6.69E-04 | 8.5 |
| | A46T | 1:1 binding | 9.40E-01 | 8.54E+04 | 7.16E-04 | 8.4 |
| | | | Average | **8.21E+04** | **6.93E-04** | **8.4** |
| Mdm2 | Ab-Y-WCT | 1:1 binding | 2.29E-01 | 4.52E+04 | 9.17E-04 | 20.3 |
| | A46T L51F | 1:1 binding | 6.78E-01 | 4.05E+04 | 1.29E-03 | 31.9 |
| | | | Average | **4.29E+04** | **1.10E-03** | **25.8** |

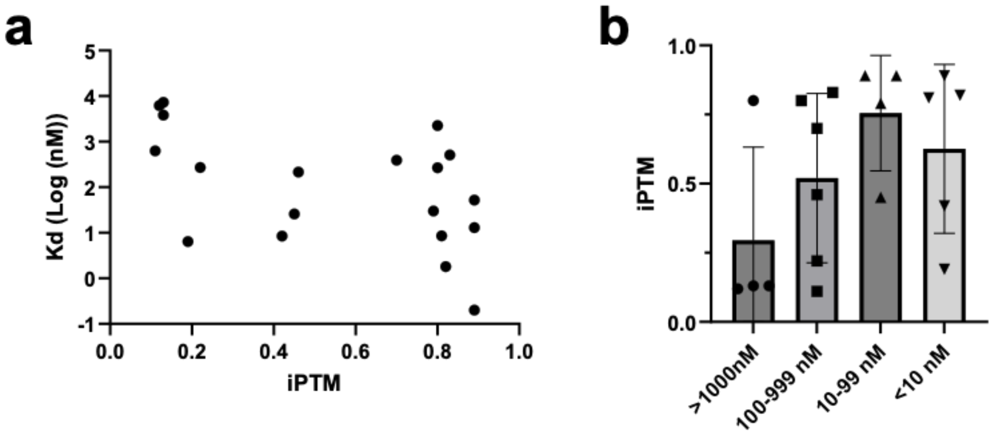


**Supplementary Fig. 32. AlphaFold cannot predict whether proteins bind**.

**a)** Correlation between the iPTM (AlphaFold 3 predictions) and the $K_d$ for every binder we determined the *in-vitro* binding affinity. Pearson Correlation of -0.50 (two tailed p = 0.028; 95% CI = -0.78 to -0.06). **b)** Grouping of iPTM values by affinity. No grouping is statistically different from any other group (all pairwise *t*-test are not significant at p = 0.05).

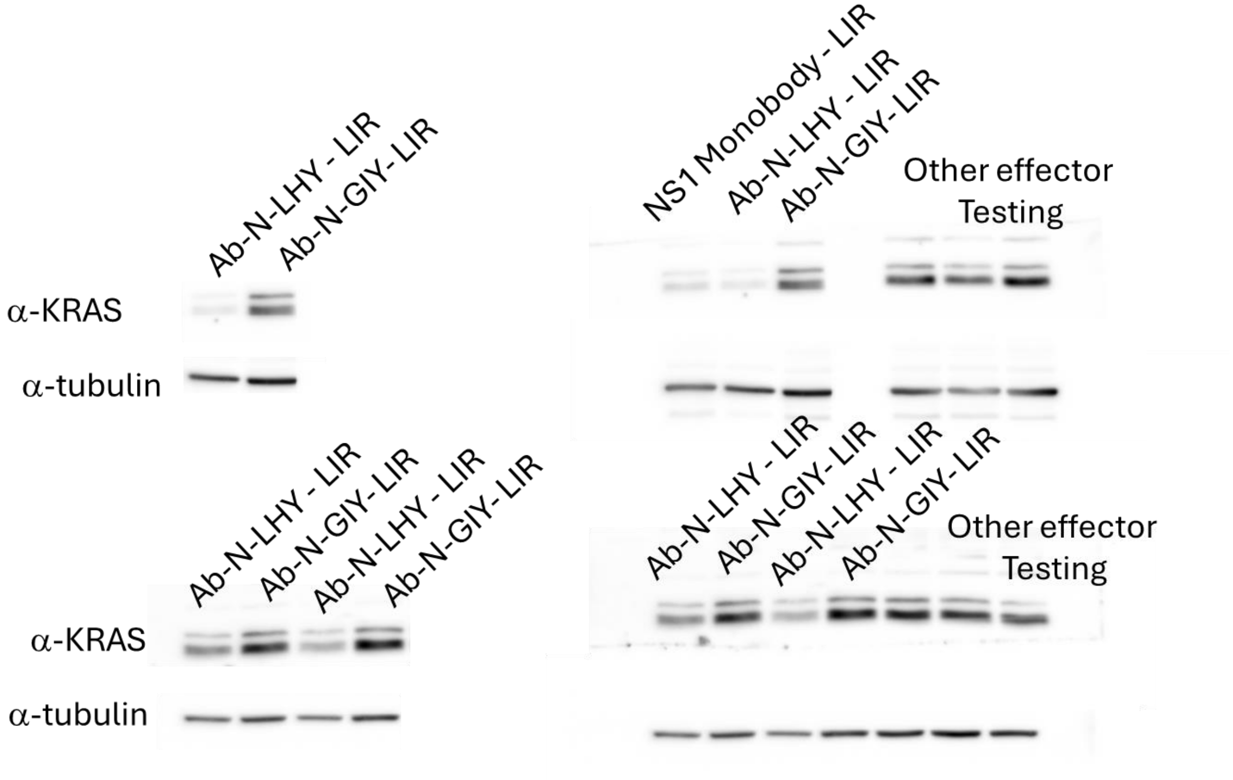


**Supplementary Fig. 33. Endogenous KRAS degradation.**

KRAS binder (Ab-N-LHY)-LIR fusion targeted degradation of endogenous KRAS in U2OS cells (**Fig 5 g,h**). An initial replicate, which included a positive control of NS1 Monobody (previously demonstrated to interact with KRAS in cells) was followed up by two additional replicate transfections. Ab-N-GIY binds to PIK3CB and not KRAS. Other bands in the full blots are from other attempts to screen effectors for degradation and are not related to LIR driven degradation.

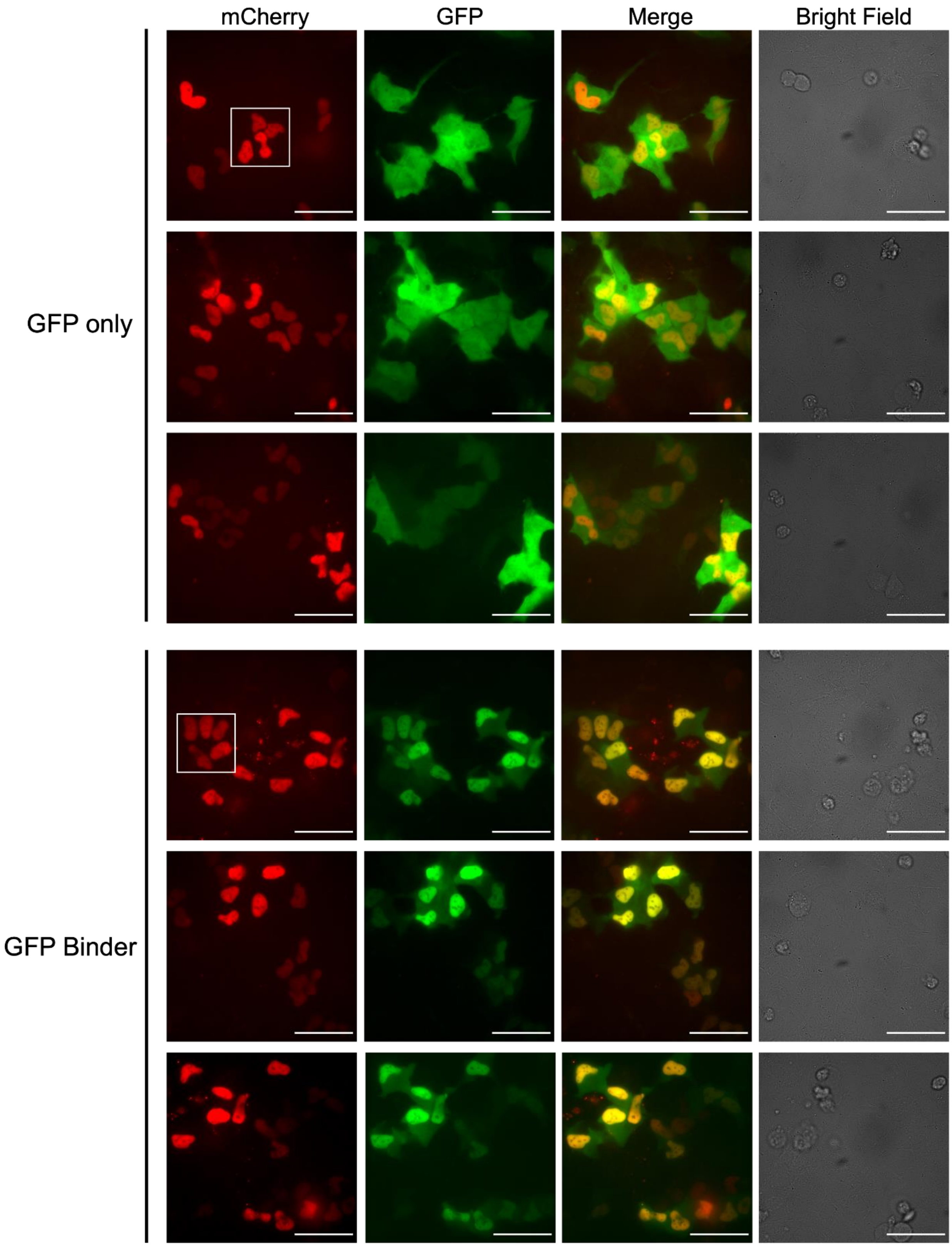


**Supplementary Fig. 34. Mdm2 co-localization.**

Mdm2-mCherry and either GFP or Mdm2 binder (**Fig. 5d**)-GFP fusions were co-transfected in HEK293T cells.

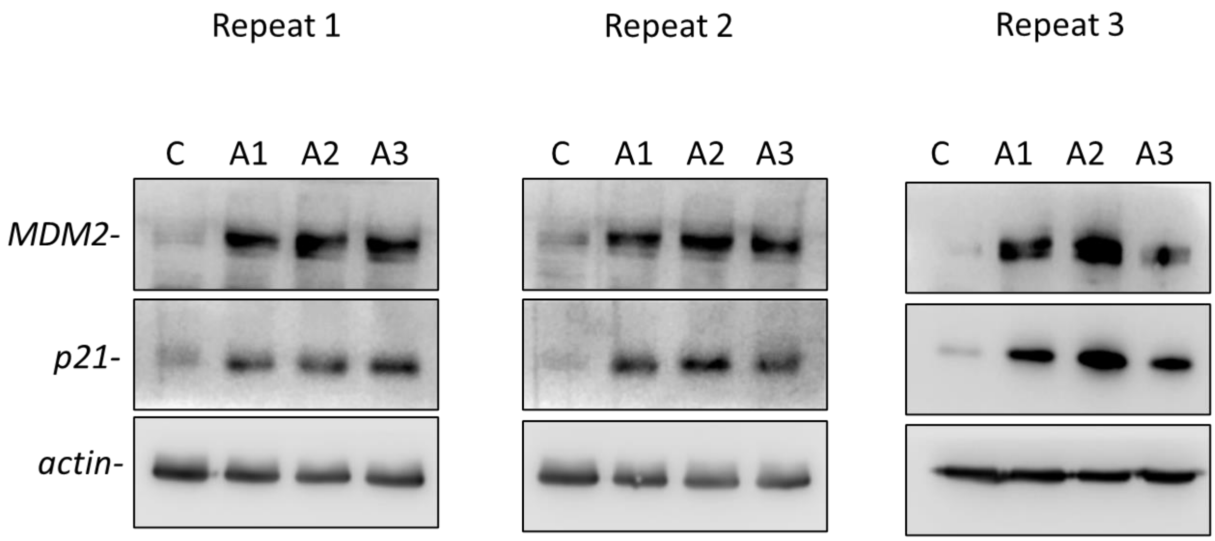


**Supplementary Fig. 35. Mdm2 inhibition.**

Mdm2 Binders (**Fig. 5d**) expressed in U2OS cells inhibit the Mdm2-p53 interaction resulting in p53 driven transcription of Mdm2 and p21 (**Fig. 5 j,k,l**).

**References**


1 Liu, Y. X. *et al.* TRIM21 is a druggable target for the treatment of metastatic colorectal cancer through ubiquitination and activation of MST2. *Cell chemical biology* **30**, 709-725.e706, doi:10.1016/j.chembiol.2023.05.009 (2023).

2 Grimm, S., Salahshour, S. & Nygren, P. A. Monitored whole gene in vitro evolution of an anti-hRaf-1 affibody molecule towards increased binding affinity. *N Biotechnol* **29**, 534-542, doi:10.1016/j.nbt.2011.10.008 (2012).

3 Koide, A., Wojcik, J., Gilbreth, R. N., Hoey, R. J. & Koide, S. Teaching an old scaffold new tricks: monobodies constructed using alternative surfaces of the FN3 scaffold. *J Mol Biol* **415**, 393-405, doi:10.1016/j.jmb.2011.12.019 (2012).

4 Spencer-Smith, R. *et al.* Inhibition of RAS function through targeting an allosteric regulatory site. *Nature chemical biology* **13**, 62-68, doi:10.1038/nchembio.2231 (2017).

5 Behar, G. *et al.* Whole-bacterium ribosome display selection for isolation of highly specific anti-Staphyloccocus aureus Affitins for detection- and capture-based biomedical applications. *Biotechnol Bioeng* **116**, 1844-1855, doi:10.1002/bit.26989 (2019).

6 Jing, L. *et al.* Screening and production of an affibody inhibiting the interaction of the PD-1/PD-L1 immune checkpoint. *Protein Expr Purif* **166**, 105520, doi:10.1016/j.pep.2019.105520 (2020).
